# EOMES and IL-10 regulate anti-tumor activity of PD-1^+^ CD4^+^ T-cells in B-cell Non-Hodgkin lymphoma

**DOI:** 10.1101/2020.03.09.983098

**Authors:** Philipp M. Roessner, Laura Llaó Cid, Ekaterina Lupar, Tobias Roider, Marie Bordas, Christoph Schifflers, Ann-Christin Gaupel, Fabian Kilpert, Marit Krötschel, Sebastian J. Arnold, Leopold Sellner, Stephan Stilgenbauer, Sascha Dietrich, Peter Lichter, Ana Izcue, Martina Seiffert

**Affiliations:** Molecular Genetics, German Cancer Research Center (DKFZ), Heidelberg, Germany; Faculty of Biosciences, University of Heidelberg, Heidelberg, Germany; Max-Planck-Institute of Immunobiology and Epigenetics, Freiburg, Germany; Department of Medicine V, Hematology, Oncology and Rheumatology, University of Heidelberg, Heidelberg, Germany; Immunotherapy and Immunoprevention, German Cancer Research Center (DKFZ), Heidelberg, Germany and Cell Biology Research Unit (URBC) – Namur Research Institute of Life Science (Narilis), University of Namur, Namur, Belgium; Essen University Hospital, Institute of Human Genetics, Genomeinformatics, Essen, Germany; BioMed X Innovation Center, Heidelberg, Germany; Institute of Experimental and Clinical Pharmacology and Toxicology, Faculty of Medicine, University of Freiburg, Freiburg, Germany; Signalling Research Centres BIOSS and CIBSS, University of Freiburg, Freiburg, Germany; Internal Medicine III, University of Ulm, Ulm, Germany; Center for Chronic Immunodeficiency, University Medical Center Freiburg and University of Freiburg, Freiburg, Germany; Institute of Molecular Medicine, University Hospital RWTH Aachen, Aachen, Germany

**Keywords:** CLL, CD4^+^ T-cells, Eomes, IL-10, Exhaustion

## Abstract

The transcription factor Eomesodermin (EOMES) promotes IL-10 production of CD4^+^ T-cells, which has been linked to immunosuppressive and cytotoxic activities. We detected EOMES-expressing CD4^+^ T-cells in lymph node samples of patients with chronic lymphocytic leukemia (CLL) or diffuse large B-cell lymphoma. This was in line with an observed expansion of EOMES-positive CD4^+^ T-cells in leukemic Eµ-TCL1 mice, a well-established model of CLL, and upon adoptive transfer of TCL1 leukemia in mice. Transcriptome and flow cytometry analyses revealed that EOMES does not only drive the transcription of IL-10, but rather controls a unique differentiation program in CD4^+^ T-cells. Moreover, EOMES was necessary for the accumulation of a specific CD4^+^ T-cell subset that expresses IFNγ and IL-10, as well as inhibitory receptors, like PD-1 and LAG3. T-cell transfer studies in leukopenic *Rag2*^*-/-*^ mice showed that EOMES-deficient CD4^+^ T-cells were inferior in controlling TCL1 leukemia development compared to wildtype T-cells, even though expansion of *Eomes*^*-/-*^ CD4^+^ T-cells was observed. We further showed that control of TCL1 leukemia was driven by IL-10 receptor-mediated signals, as *Il10rb*-deficient CD4^+^ T-cells showed impaired anti-leukemia activity. Altogether, our data suggest that IL-10 producing PD-1^+^ CD4^+^ T-cells contribute to CLL control in an EOMES- and IL-10R-dependent manner.

## Introduction

Despite abundant data characterizing CD4^+^ T-cells and their subsets in B-cell Non-Hodgkin lymphoma (B-NHL) (1), and in particularly chronic lymphocytic leukemia (CLL) (2-6), their role in disease development and progression is poorly understood. Besides well-known T helper (Th) cell subsets (5), interleukin (IL-)10 producing, FOXP3^-^ conventional CD4^+^ T-cells, named type 1 regulatory (T_R_1) cells, are gaining attention in mouse models as well as patients harboring chronic inflammatory conditions such as inflammatory bowel disease (7-10). T_R_1 cells were initially described as IL-10-induced cells that produce IL-10 and IFNγ and harbor cytotoxic activity, but also express several co-inhibitory receptors such as programmed cell death protein-1 (PD-1).

In B-NHL, an increased expression of PD-1 in blood-derived CD4^+^ T-cells was reported for diffuse large B-cell lymphoma (DLBCL) (1, 11) as well as for CLL (4, 12, 13). In a preclinical study, a blocking antibody against the PD-1 ligand 1 (PD-L1) showed high activity in controlling CLL progression in the Eµ-TCL1 mouse model (14). These findings were the basis for clinical trials using immune checkpoint inhibitors targeting the PD-1/PD-L1 axis, which lead to varying but in general disappointing clinical results with none of the included CLL patients achieving remission in response to therapy and only a subgroup of patients, harboring a more aggressive Richter’s transformation, benefitting from this treatment (NCT02332980) (15). It was hypothesized that the lack of clinical success of PD-1 blockade in CLL was due to the fact that tumor infiltrating T-cells in CLL have a lower expression of PD-1 in comparison to other B-NHL entities, including DLBCL (11). In DLBCL, PD-1 expression was shown to correlate with better survival (16, 17). However, immune checkpoint blockade resulted in an overall response rate of only about 10% in DLBCL patients (NCT02038933) (18).

Recently, the transcription factor Eomesodermin (EOMES) has been shown to promote IL-10 production in T_R_1 cells (7-9). EOMES belongs to the T-box transcription factor family, which is expressed in many organs including the immune system (19). Redundantly with its paralogue T-BET, EOMES has been shown to promote IFNγ production and cytotoxicity in natural killer (NK) cells (20-22), as well as in CD8^+^ (21-23) and CD4^+^ T-cells (7, 8, 21, 24, 25). EOMES has also a non-redundant role in promoting maturation of the classical NK cell subset (26), and in the accumulation of central memory (22, 27) and exhausted, PD-1 expressing CD8^+^ T-cells (28). Co-expression of PD-1 and EOMES of CD8^+^ T-cells was confirmed by us in a mouse model of CLL (29).

In contrast to NK cells and CD8^+^ T lymphocytes, very few CD4^+^ T-cells express EOMES without immunological challenge (30). However, we and others have shown that EOMES can be upregulated in CD4^+^ T-cells upon activation, which affects their differentiation into helper cell lineages. To this end, it has been shown that EOMES promotes IFNγ expression by Th1 T-cells (31-33) and inhibits differentiation of Th17 (8, 33, 34) and FOXP3^+^ regulatory T-cells (Treg) (30).

In cooperation with other factors like PR domain zinc finger protein 1 (BLIMP1), EOMES is a driver of IL-10 production in T_R_1 T-cells in mice (7) as well as humans (8). Whether EOMES only promotes *Il10* expression by T_R_1 cells or is a lineage-defining transcription factor is still unclear (35).

As investigations of EOMES and IL-10-producing CD4^+^ T-cells in B-NHL, in particularly CLL and DLBCL, are scarce, we quantified EOMES-expressing CD4^+^ T-cells in samples of CLL and DLBCL patients and the Eµ-TCL1 mouse model of CLL. Using isolated cells of *Eomes-GFP* reporter mice, we characterized the transcriptional profile of EOMES^+^ CD4^+^ T-cells and elucidated the regulatory role of EOMES for this profile, identifying IL-10 as a main target in these cells that depends on EOMES. By performing co-transfer experiments of CD4^+^ T-cells and Eµ-TCL1 leukemia cells in immunodeficient mice, we further unravelled the importance of EOMES and IL-10 in CD4^+^ T-cells in controlling leukemia development.

## Methods

### Patient samples

Patient samples were obtained after approval of study protocols by local ethics committees from the Department of Internal Medicine III of the University Clinic Ulm and the Department of Medicine V of the University Clinic Heidelberg according to the declaration of Helsinki, and after obtaining informed consent of patients. Patients met standard diagnosis criteria for CLL or DLBCL, respectively. Patient characteristics such as age, mutational state and Binet stage are provided in Supplementary Tables 1-3. Healthy, age-matched controls were obtained from Biomex GmbH (Heidelberg, Germany) after informed consent.

### Tumor models and adoptive CD4^+^ T-cell transfer

Adoptive transfer of mouse leukemic cells was performed by i.p. or i.v. transplantation of 1-2*10^7^ Eμ-TCL1 splenocytes into C57BL/6 N or J wildtype (WT) animals. Splenocytes were enriched in CD19^+^ cells using EasySep™ Mouse Pan-B Cell Isolation Kit (Stemcell Technologies, Vancouver, Canada) yielding a purity above 95% of CD5^+^ CD19^+^ cells. For CD4^+^ co-transfer experiments, CD4^+^ T-cells were isolated from splenocytes using EasySep™ Mouse CD4^+^ T Cell Isolation Kit resulting in a purity of about 95% CD4^+^ T-cells of total T-cells. *Rag2*^*-/-*^ mice (DKFZ central animal facility) were i.v. transplanted with 2*10^5^ CD4^+^ T-cells or PBS as control. The following day, 1*10^6^ purified TCL1 leukemic cells were transferred i.v. into recipients.

Adoptive transfer of naïve CD4^+^ T-cells was performed as previously described (30). In brief, *Rag2*^*−/−*^ recipient mice received 4*10^5^ FACS-sorted CD4^+^ CD45RB^high^ T-cells by i.p. injection. Three weeks post transfer, mice were sacrificed and spleens analyzed.

All animal experiments were carried out according to institutional and governmental guidelines approved by the local authorities (Regierungspräsidium Karlsruhe, permit numbers: G36/14, G98/16, G123/14, and Regierungspräsidium Freiburg, permit number: 35-9185.81/G-13/73).

### Statistical analysis

Samples of different groups were compared using non-parametric Mann-Whitney test. Comparison of matched samples was performed using Wilcoxon matched-pairs signed rank test Values of p < 0.05 were considered statistically significant.

## Results

### EOMES-expressing PD-1^+^ CD4^+^ T-cells are present in B-NHL patients

CLL is associated with elevated numbers of CD4^+^ and CD8^+^ T-cells in blood (5, 36, 37), and increased expression of inhibitory molecules in T-cells has been reported in several studies for CLL (4, 13, 37) as well as DLBCL patients (1, 11). To explore the relevance of T-cell accumulation for disease progression, we first quantified PD-1 expressing CD4^+^ T-cells in blood samples of patients with CLL or DLBCL, which represent an indolent and more aggressive B-NHL, respectively, in comparison to healthy controls (HC). Confirming previously published data, we detected higher percentages of PD-1-expressing CD4^+^ T-cells in both patient groups in comparison to age-matched healthy controls (Figure 1A, C), as well as higher absolute numbers of these cells in CLL, but not DLBCL samples (Figure 1B, D).

**Figure 1:**
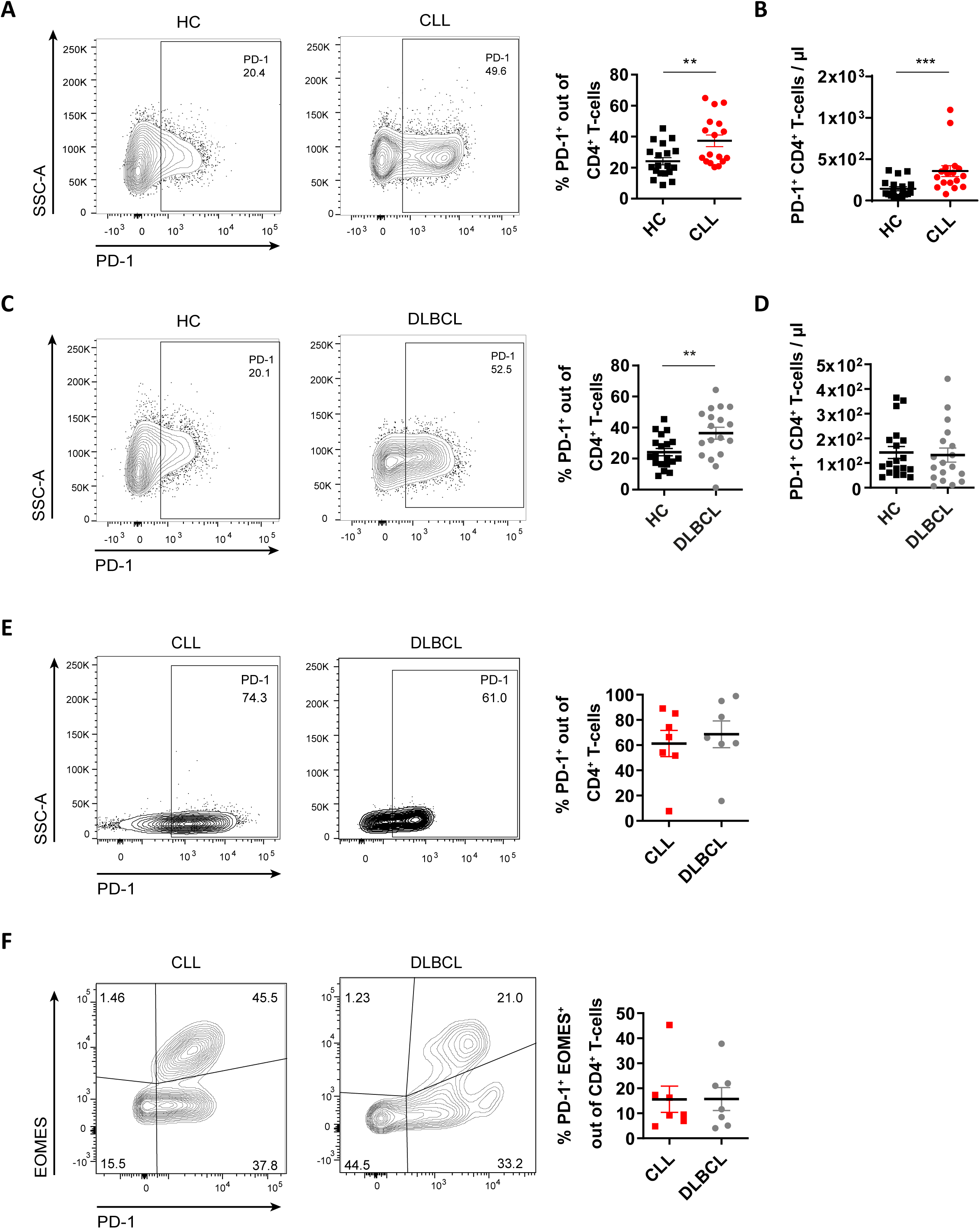
EOMES-expressing PD-1^+^ CD4^+^ T-cells are abundant in B-cell Non-Hodgkin lymphoma patients. **A-D)** Blood samples of CLL and DLBCL patients were stained for flow cytometry. **A)** Representative contour plots as well as percentage, and **B)** absolute numbers of PD-1^+^ CD4^+^ T-cells per µl blood of CLL patients and age-matched healthy controls (HC). **C)** Representative contour plots as well as percentage, and **D)** absolute numbers of PD-1^+^ CD4^+^ T-cells per µl blood of of DLBCL patients and HC. **E-F)** Single-cell suspensions from lymph node samples of CLL as well as DLBCL patients were analyzed by flow cytometry. **E)** Representative contour plots as well as percentage of PD-1^+^ out of CD4^+^ T-cells. **F)** Representative contour plots as well as frequency of EOMES^+^ PD-1^+^ cells out of CD4^+^ T-cells. All graphs show mean ± SEM. Each dot represents data of an individual patient. Statistical analysis was performed using Mann-Whitney test. *p<0.05, **p<0.01.

Malignant B-cells in CLL (38) and DLBCL proliferate and expand in secondary lymphoid organs, which are also the sites of T-cell activation (39). Along this line, we have recently noted that the phenotype of peripheral blood T-cells and lymphoid organ-derived T-cells is distinct in CLL (37). Therefore, we analyzed lymph node (LN) samples of CLL and DLBCL patients and detected a substantial proportion of CD4^+^ T-cells that express PD-1 in most samples (mean: 61% in CLL and 69% in DLBCL) with a high variability of the percentages ranging from 8 to 95% (Figure 1E; gating strategy: Supplementary Figure 1). We further observed that approximately 20% of all CD4^+^ T-cells in CLL and DLBCL LN samples co-expressed PD-1 and the transcription factor EOMES (Figure 1F).

Hence, EOMES-expressing PD-1^+^ CD4^+^ T-cells are present in CLL and DLBCL patients.

### Leukemia development in the Eµ-TCL1 mouse model is associated with an accumulation of EOMES^+^ PD-1^+^ CD4^+^ T-cells

To further investigate the role of EOMES-expressing CD4^+^ T-cells in B-NHL, we analyzed the presence of these cells in the Eµ-TCL1 (TCL1) mouse model of CLL. Interestingly, analysis of age- and sex-matched leukemic TCL1 mice and wildtype (WT) littermates revealed a higher abundance of PD-1^+^ CD4^+^ T-cells in the spleen of TCL1 mice (Figure 2A, Gating strategy Supplementary Figure 2A). Of note, the majority of these PD-1^+^ cells also expressed EOMES (Figure 2B) as well as the inhibitory receptor LAG3 (Supplementary Figure 2B). To overcome long latency of CLL development in this mouse model, leukemic splenocytes of TCL1 mice were retrieved and adoptively transfered into syngeneic WT mice (TCL1 AT), as previously described (6, 37, 40). Upon leukemia development in the TCL1 AT model, we observed an accumulation of antigen-experienced CD4^+^ T-cells (Supplementary Figure 2C) that show signs of activation as measured by CD69 (Supplementary Figure 2D). Moreover, a higher frequency of PD-1-expressing CD4^+^ T-cells (Figure 2C) was detected, which showed a high co-expression rate of EOMES (Figure 2D) as well as LAG3 (Supplementary Figure 2E). Interestingly, the frequency of EOMES^+^ PD-1^+^ CD4^+^ T-cells was higher in aging Eµ-TCL1 mice compared to the younger mice of the TCL1 AT (Figure 2B, D), which is in line with our previous data showing that EOMES^+^ CD4^+^ T-cells accumulate with age (30).

**Figure 2:**
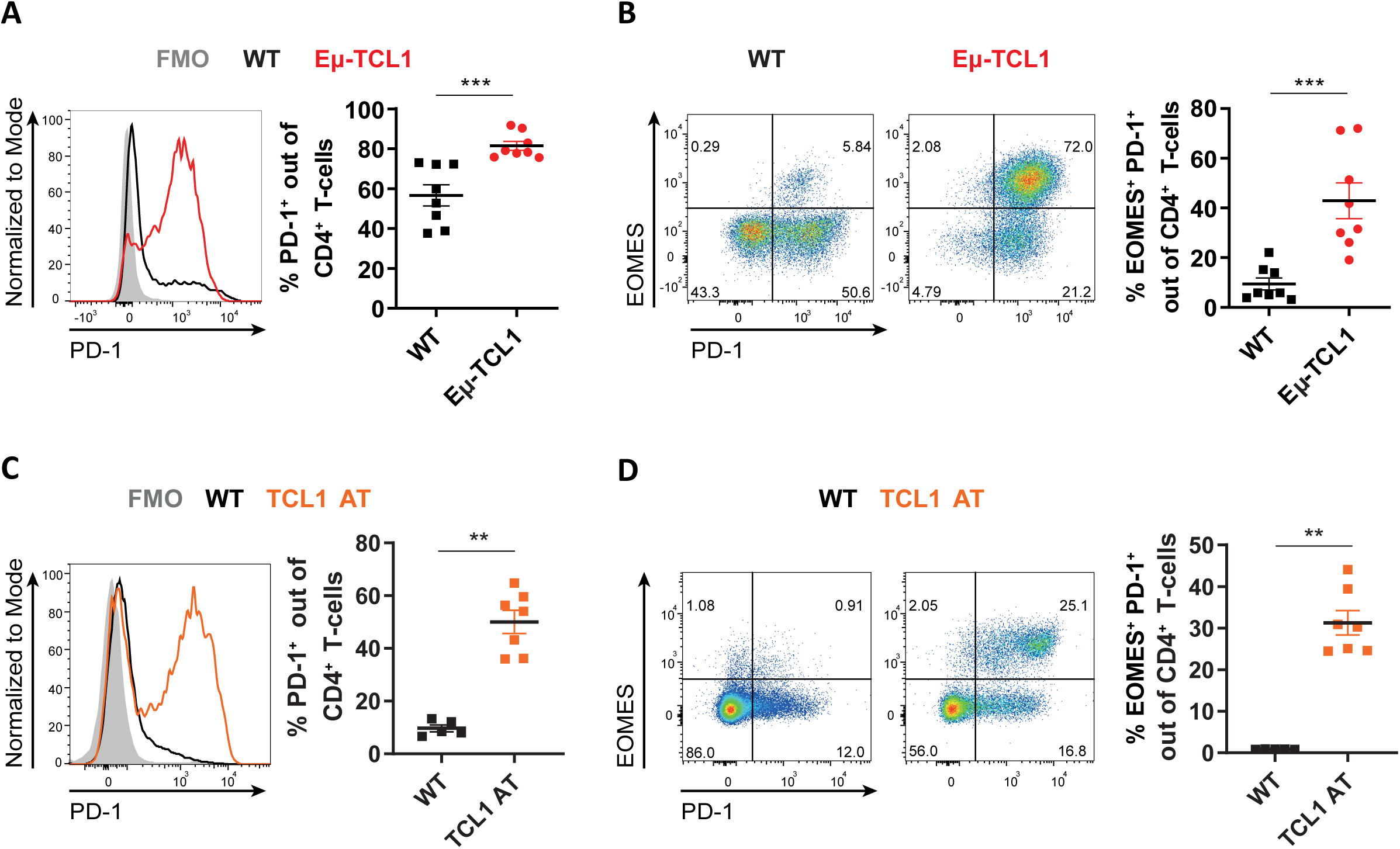
PD-1^+^ CD4^+^ T-cells are enriched in the Eµ-TCL1 mouse model of CLL and co-express EOMES. **A)-B)** Splenocytes of hemizygous Eµ-TCL1 (TCL1) leukemic mice and WT littermates at a median age of 65 weeks were analyzed by flow cytometry. **A)** Representative histogram as well as percentage of PD-1^+^ out of CD4^+^ T-cells. **B)** Representative flow cytometry plots and frequency of EOMES^+^ PD-1^+^ cells of CD4^+^ T-cells. **C)-D)** Leukemic cells of TCL1 mice were transplanted into syngenic WT mice (TCL1 AT) and splenocytes were analyzed by flow cytometry, 4 weeks after transfer of cells. Representative histogram or flow cytometry plots, and quantification of the frequency of **C)** PD-1^+^ out of CD4^+^ T-cells, as well as **D)** EOMES^+^ PD-1^+^ out of CD4^+^ T-cells. All graphs show mean ± SEM. Each dot represents data of an individual mouse. Statistical analysis was performed using Mann-Whitney test. **p<0.01, ***p<0.001.

Altogether, CLL development in the Eµ-TCL1 and TCL1 AT models is associated with an enrichment of EOMES-expressing PD-1^+^ CD4^+^ T-cells and therefore, these models are useful tools to investigate the role of this cell type in CLL.

### EOMES is indispensable for CD4^+^ T-cell mediated control of leukemia development in TCL1 AT mice

In order to decipher the role of EOMES^+^ PD-1^+^ CD4^+^ T-cells in CLL development, we transferred CD4^+^ T-cells into *Rag2*^-/-^ mice, which lack mature B- and T-cells (41). Subsequently, the mice were transplanted with TCL1 leukemia cells, as previously described (6, 37, 40). We detected an expansion of CD4^+^ T-cells in *Rag2*^*-/-*^ mice, which was similar with or without leukemia cell transfer (Supplementary Figure 3A). We further observed that CD4^+^ T-cells controlled leukemia progression, as indicated by lower CD5^+^ CD19^+^ CLL cell counts in blood (Figure 3A) as well as lower weight and leukemia cell content per spleen compared to mice without T-cell transfer (Supplementary Figure 3B-C). Of interest, CD4^+^ T-cells from leukemia-bearing *Rag2*^*-/-*^ mice showed a higher frequency of PD-1-positive cells (Supplementary Figure 3D) as well as EOMES^+^ PD-1^+^ CD4^+^ T-cells (Supplementary Figure 3E) in comparison to non-leukemic control mice.

**Figure 3:**
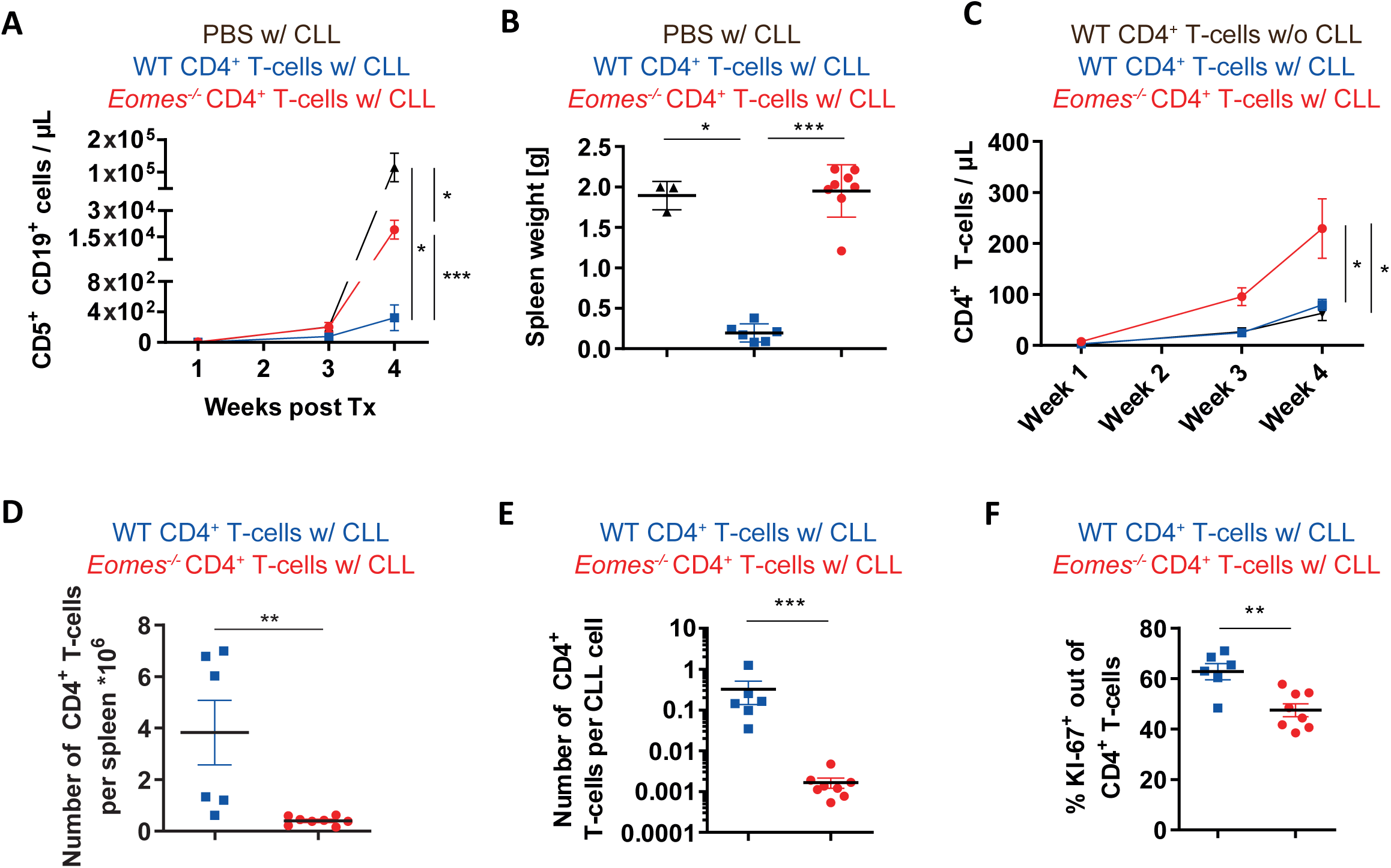
EOMES is crucial for CLL control by CD4^+^ T-cells. *Rag2*^*-/-*^ mice were transplanted i.v. with PBS or 2× 10^5^ CD4^+^ T-cells of WT or *Eomes*^*-/-*^ origin on day −1 and the following day with 1 × 10^6^ leukemic cells from Eµ-TCL1 mice, and analyzed at indicated timepoints by flow cytometry. **A)** Absolute numbers of CD5^+^ CD19^+^ CLL cells in peripheral blood are shown over time. **B)** Spleen weight at endpoint, 4 weeks after transfer of leukemic cells. **C)** Absolute numbers of CD4^+^ T-cells cells in peripheral blood are depicted over time. **D)** Number of CD4^+^ T-cells per spleen, as well as **E)** number of CD4^+^ T-cells per CD5^+^ CD19^+^ CLL cell are shown. **F)** Percentage of KI-67^+^ out of CD4^+^ T-cells. All graphs show mean ± SEM. In B) and D)-F), each dot represents data of an individual mouse. Statistical analysis was performed using Mann-Whitney test. *p<0.05, **p<0.01, ***p<0.001.

To analyze the role of EOMES in controlling CLL progression, we next transplanted wildtype (WT) or *Eomes* knock-out (*Eomes*^-/-^) CD4^+^ T-cells, expressing fluorescent reporter proteins for *Foxp3* (RFP) and *Il10* (GFP), into *Rag2*^-/-^ mice followed by adoptive transfer of TCL1 leukemia cells. Analysis of CLL progression in these mice showed that *Eomes*^-/-^ CD4^+^ T-cells failed to control CLL development as evidenced by higher numbers of CD5^+^ CD19^+^ CLL cells in blood (Figure 3A) as well as higher spleen weights (Figure 3B). We further monitored CD4^+^ T-cell expansion in these mice over time and observed higher T-cell numbers in recipient mice of *Eomes*^-/-^ CD4^+^ T-cells compared to WT T-cells (Figure 3C). However, in the spleen of these animals, a lower absolute number of CD4^+^ T-cells per spleen (Figure 3D) as well as per CLL cell (Figure 3E) was noted in the *Eomes*^-/-^ in comparison to the WT group. In line with this, *Eomes*^-/-^ T-cells in the spleen showed a lower proliferation rate based on KI-67 staining compared to WT T-cells (Figure 3F).

To sum up, CD4^+^ T-cells lacking EOMES show an impaired control of TCL1 leukemia progression in *Rag2*^-/-^ mice in comparison to EOMES-proficient T-cells. Although *Eomes*^-/-^ CD4^+^ T-cells are more highly abundant in blood compared to WT T-cells, they proliferate less and are significantly lower in numbers in the spleen of these mice, which might explain the reduced leukemia control in the *Eomes*^-/-^ group.

### EOMES positive CD4^+^ T-cells express inhibitory receptors

To gain mechanistic insights into how EOMES regulates CD4^+^ T-cell differentiation and function, we performed comparative transcriptome analyses of GFP^+^ EOMES^+^ and GFP^-^ EOMES^-^ CD4^+^ T-cells from *Eomes-GFP* reporter mice (*Eomes*^*+/GFP*^) (42). Moreover, we obtained gene expression data of GFP^+^ versus GFP^-^ CD4^+^ T-cells isolated from *Eomes*^*ΔT*/GFP^ knock-out mice, in which one *Eomes* allele is disrupted by *GFP* insertion and the other allele is deleted using a T-cell-specific *cre* recombinase. Deletion of the floxed exons of *Eomes* in these mice was confirmed by RNA sequencing (Supplementary Figure 4). To expand EOMES-expressing cells for these analyses, we transferred naïve CD25^-^ CD45RB^high^ CD4^+^ T-cells isolated from *Eomes*^*+/GFP*^ reporter or *Eomes*^*ΔT/GFP*^ knock-out donor animals into *Rag2*^*-/-*^ mice as previously described (30), and three weeks later, sorted GFP^+^ and GFP^-^ CD4^+^ T-cell populations from both *Eomes*^*+/GFP*^ and *Eomes*^*ΔT/GFP*^ donor mice for RNA sequencing (see Figure 4A for experimental setup).

**Figure 4:**
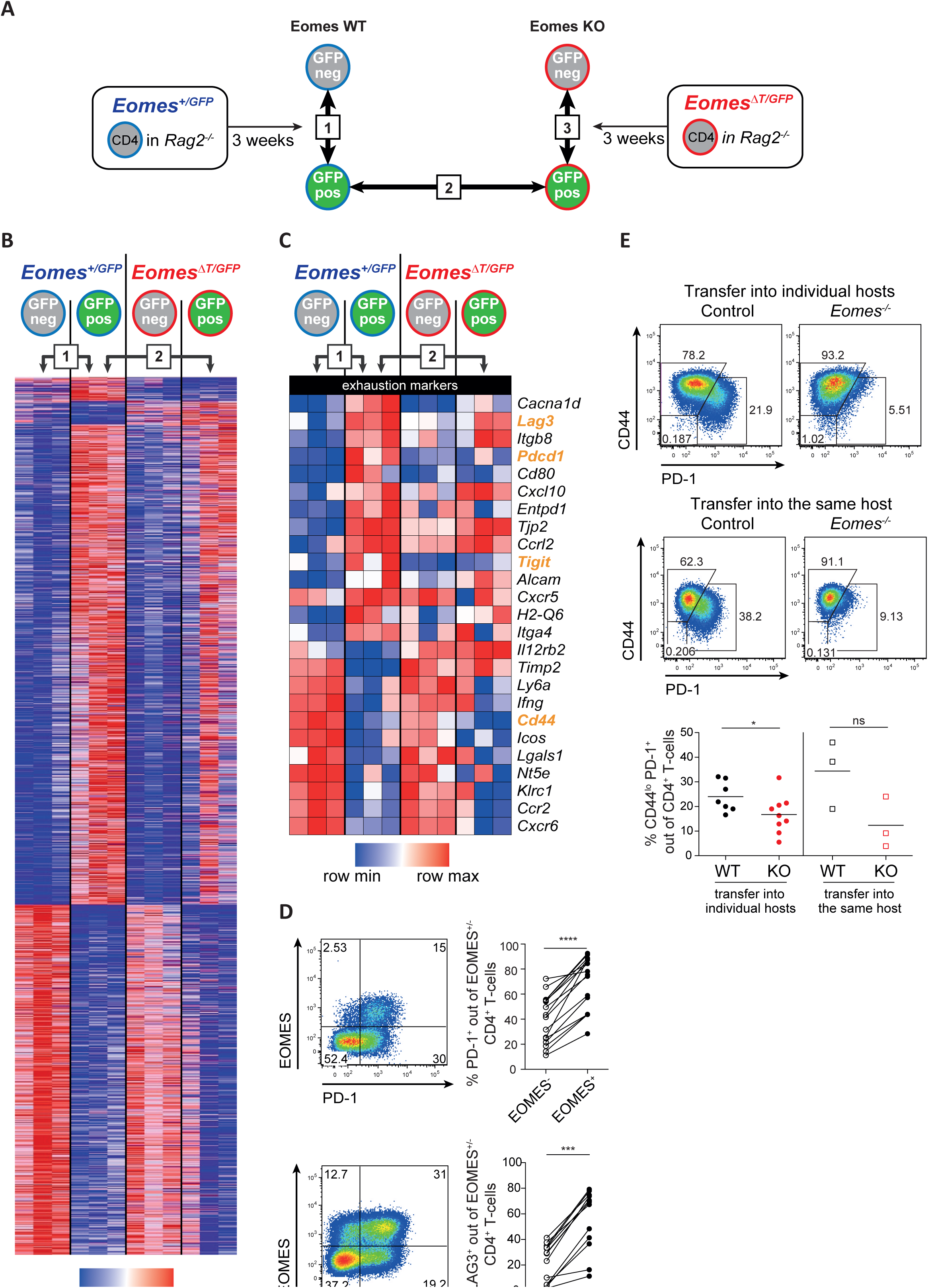
EOMES drives inhibitory co-receptor expression in CD4^+^ T-cells. Naïve CD25^-^ CD45RB^High^ CD4^+^ T-cells of *Eomes-GFP* reporter mice (*Eomes*^*+/GFP*^), as well as of mice with a T-cell-specific deletion of *Eomes* in combination with a GFP reporter (*Eomes*^*ΔT/GFP*^) were transferred into *Rag2*^*-/-*^ mice, and three weeks after adoptive transfer, RNA sequencing of sorted, splenic CD4^+^ T-cell subsets was performed. **A)** Strategy of RNA sequencing analysis of GFP^+^ and GFP^-^ cells of *Eomes*^*+/GFP*^ (comparison 1) as well as *Eomes*^*ΔT/GFP*^ mice, depicting also the comparison of GFP^+^ cells from *Eomes*^*+/GFP*^ and *Eomes*^*ΔT/GFP*^ mice (comparison 2). **B)** Heatmap of all differentially expressed genes. **C)** Heatmap showing differentially expressed, exhaustion-associated surface receptor genes (selected as differentially expressed in exhausted T-cells (43) filtered by surface expression (GO:0009986)). **D)** Representative flow cytometry plots of PD-1 and LAG-3 expression in EOMES^+^ and EOMES^-^ CD4^+^ T-cells and quantification of data. **D)** Flow cytometric analysis showing representative graphs and quantification of the frequency of the CD44^lo^ PD-1^+^ population after transfer of *Eomes*^*+/+*^ or *Eomes* ^*-/-*^ CD4^+^ T-cells into different or the same *Rag2*^*-/-*^ hosts. Gene expression in heatmaps in C) and D) was row-normalized. Each dot represents data of an individual mouse. Lines in D) link data of EOMES^-^ and EOMES^+^ cells from the same animal. Statistical analysis was performed using Mann-Whitney test. Comparison of matched samples was performed using Wilcoxon matched-pairs signed rank test. *p<0.05, ***p<0.001, ****p<0.0001.

Comparing EOMES^+^ versus EOMES^-^ T-cells from *Eomes*^*+/GFP*^ reporter mice (Figure 4A, comparison 1), we identified a signature transcriptome of EOMES^+^ CD4^+^ T-cells with 1,395 differentially expressed genes (Figure 4B; Supplementary Figure 5A; Supplementary Table 4). In contrast, only 77 genes (Figure 4B; Supplementary Figure 5A; Supplementary Table 4) were dependent on the presence of EOMES, as they were differentially expressed between EOMES^+^ GFP^+^ cells from *Eomes*^*+/GFP*^ reporter mice and EOMES^-^ GFP^+^ cells from *Eomes*^*ΔT*/GFP^ knock-out mice (Figure 4A, comparison 2). Intersecting these two sets of differentially expressed genes (Figure 4A, comparison 1, 2) we identified a small number of 37 genes (Supplementary Figure 5A; Supplementary Table 4) as being dependent on the transcriptional activity of EOMES. While this small difference can be partially attributed to a high variability of data among the three samples of *Eomes* knock-out origin, a heatmap of the differentially expressed genes in the two comparisons shows that most of the transcriptional signature of *Eomes*^*+/GFP*^ CD4^+^ T-cells is not dependent on EOMES (Figure 4B).

As previously shown (30), both EOMES^+^ and EOMES^-^ CD4^+^ T-cells expressed Th1-like transcripts, such as *Tbx21, Ifng* and *Cxcr3* (Supplementary Figure 5B). In line with the redundant functions of EOMES and T-BET in Th1 differentiation (32), the Th1-like profile was EOMES-independent as these genes were also expressed in GFP^+^ CD4^+^ T-cells of *Eomes*^*ΔT/GFP*^ knock-out mice (Supplementary Figure 5B). Neither Th2-, Th17-, or other Th-, nor Treg-associated genes were expressed by GFP^+^ cells of *Eomes*^*+/GFP*^ reporter or *Eomes*^*ΔT/GFP*^ knock-out mice (Supplementary Figure 5B), respectively, suggesting that these CD4^+^ T-cell fates are not regulated by EOMES.

Gene set enrichement analysis (GSEA) identified similarities of the transcriptional profile of EOMES^+^ CD4^+^ T-cells with human T_R_1 cells (8) (Supplementary Figure 5C) which is indicative for shared functional properties of these two cell populations.

Interestingly, the identified signature of EOMES^+^ CD4^+^ T-cells contained several transcripts of co-inhibitory receptors, like LAG3, PD-1 (encoded by the *Pdcd1 gene*), and TIGIT which were upregulated in EOMES^+^ CD4^+^ T-cells (Figure 4C), as similarily reported for exhausted CD4^+^ T-cells in chronic viral infections (43). We further validated the increased expression of PD-1 and LAG3 in EOMES^+^ versus EOMES^-^ CD4^+^ T-cells by flow cytometry (Figure 4D). Of note, also EOMES^-^ CD4^+^ T-cells expressed these inhibitory receptors, albeit to a significantly lower frequency of cells.

As EOMES^+^ CD4^+^ T-cells showed a low gene expression of *Cd44* (Figure 4C), we investigated whether EOMES-expressing CD4^+^ T-cells can be defined by the expression of PD-1 and low expression of CD44 (CD44^lo^). Adoptive transfer of WT or *Eomes*^-/-^ CD4^+^ T-cells in *Rag2*^*-/-*^ mice revealed a lower frequency of the PD-1^+^ CD44^lo^ CD4^+^ T-cell population in the *Eomes* knock-out setting (Figure 4E). This finding was reproduced after co-transplantation of WT and *Eomes*^-/-^ CD4^+^ T-cells into the same recipients (Figure 4E). Whereas PD-1^+^ CD44^lo^ was suitable to identify EOMES^+^ CD4^+^ T-cells in T-cell transfer experiments in *Rag2*^*-/-*^ mice, this population was not distinct in aged mice (Supplementary Figure 5D), suggesting that this marker combination is not suitable to identify EOMES^+^ CD4^+^ T-cells under more physiological conditions.

In summary, EOMES drives the expression of a small gene signature in CD4^+^ T-cells, which includes PD-1 and several other co-inhibitory receptors.

### EOMES drives IL-10 production and IL-10 receptor expression in CD4^+^ T-cells

In addition to the high expression of inhibitory receptors by EOMES^+^ CD4^+^ T-cells, we identified three cytokines among the upregulated genes (*Il10, Il1b, Tnfsf13b*), of which *Il10* showed an EOMES-dependent expression (Figure 5A; Supplementary Table 4), which is in agreement with previous reports (7, 8). We further observed a significantly higher expression of the IL-10 receptor alpha and beta chain genes, *Il10ra* and *Il10rb*, in GFP^+^ versus GFP^-^ CD4^+^ T-cells of *Eomes*^*+/GFP*^ reporter mice (Figure 4A, comparison 1), which was less pronounced in the *Eomes*^*ΔT/GFP*^ knock-out mouse line (Figure 5A), suggesting that expression of these genes might be EOMES-dependent.

**Figure 5:**
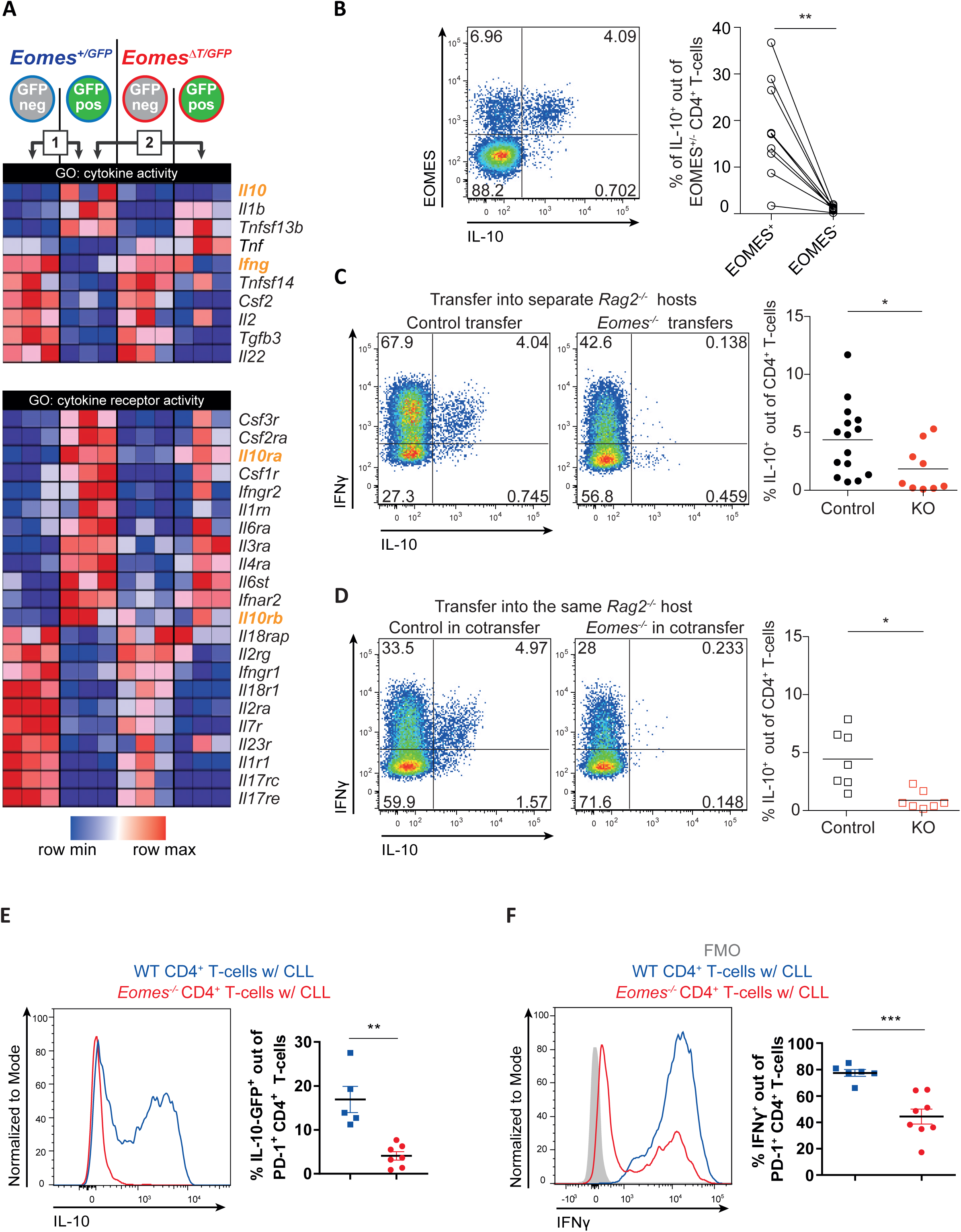
EOMES drives IL-10 production in CD4^+^ T-cells. Cells for analyses were prepared as described in Figure 4. **A)** RNA sequencing of sorted, splenic CD4^+^ T-cell subsets was performed. Depicted heatmaps focus on differentially expressed genes related to GO terms cytokine activity (GO:005125) and cytokine receptor activity (GO:0004896). **B)** Representative plot with quantification showing IL-10 production by EOMES^+^ and EOMES^-^ CD4^+^ T-cells 3 weeks post transfer into *Rag2*^-/-^ hosts as analyzed by intracellular flow cytometry after stimulation with PMA/ionomycin *ex vivo*. **C-D)** *Eomes*^*+/+*^ or *Eomes*^*-/-*^ CD4^+^ T-cells were transferred into **C)** separate, or **D)** the same *Rag2*^*-/-*^ hosts and stimulated as described in B). *Eomes*^*+/+*^ and *Eomes*^*-/-*^ cells were distinguished by expression of the congenic marker CD45.1. Graphs show representative expression of IL-10 and IFNγ, as well as quantification of IL-10-producing CD4^+^ T-cells. **E-F)** *Rag2*^*-/-*^ mice were transplanted with CD4^+^ T-cells of WT or *Eomes*^*-/-*^ origin on day −1 and the following day with leukemic cells of Eµ-TCL1 mice, and analyzed by flow cytometry. **F)** Intrinsic production of IL-10-GFP out of PD-1^+^ CD4^+^ T-cells shown as representative histogram and quantification. **G)** Splenocytes were stimulated *ex vivo* with PMA/ionomycin and cytokine expression was analyzed by intracellular flow cytometry. The graph shows percentage of IFNγ-producing cells out of PD-1^+^ CD4^+^ T-cells. Gene expression in heatmaps in A) was row-normalized. Each dot represents data of an individual mouse. Lines in B) link data of EOMES^-^ and EOMES^+^ cells from the same mouse. E)-F) show mean ± SEM. Statistical analysis was performed using Mann-Whitney test. Comparison of matched samples was performed using Wilcoxon matched-pairs signed rank test. *p<0.05, **p<0.01, ***p<0.001.

Transplantation of T-cells into *Rag2*^*-/-*^ mice, confirmed that the majority of IL-10-producing CD4^+^ T-cells co-express EOMES (Figure 5B). Importantly, most IL-10-producing cells also co-expressed IFNγ (Figure 5C) which was dependent on EOMES but not on microenvironmental factors, as similar results were seen after transfer of WT and *Eomes*^-/-^ CD4^+^ T-cells in separate and the same *Rag2*^*-/-*^ hosts (Figure 5C, D).

IL-10 has been shown to exert tumor-supporting (44) as well as anti-tumoral functions (45). To analyze whether IL-10 was also produced by PD-1^+^ CD4^+^ T-cells in the setting of TCL1 leukemia, we subsequently analyzed IL-10 and IFNγ production in the TCL1 AT mouse model. Indeed, we observed an increased percentage of CD4^+^ T-cells that produce IL-10 after transfer of TCL1 leukemia which was consiberably higher compared to unchallenged mice (Supplementary Figure 6). In addition, we confirmed a higher production of IFNγ of CD4^+^ T-cells in this transfer model, as previously described by us (5). Transfer of CD4^+^ T-cells into *Rag2*^*-/-*^ mice showed that lack of EOMES in CD4^+^ T-cells ablated the expression of IL-10 also in the TCL1 leukemia setting (Figure 5E) and caused a reduced production of IFNγ in PD-1^+^ CD4^+^ T-cells after *ex vivo* stimulation (Figure 5F).

In sum, these results confirm that EOMES drives IL-10 production in CD4^+^ T-cells, and show that CLL development strongly enhances IL-10 and IFNγ production in CD4^+^ T-cells.

### IL-10R signaling maintains moderate EOMES expression in CD4^+^ T-cells and allows them to control CLL

Since EOMES^+^ CD4^+^ T-cells showed increased expression of IL-10 receptor genes (*Il10ra* and *Il10rb*, Figure 5A), we investigated the role of IL-10R-mediated signaling in EOMES^+^ PD-1^+^ CD4^+^ T-cells and its impact on control of TCL1 leukemia. Either *Il10rb*^*+/+*^ (WT) or *Il10rb*^*-/-*^ CD4^+^ T-cells were injected into *Rag2*^-/-^ mice followed by transplantation of TCL1 leukemia. Of interest, *Il10rb*-deficient CD4^+^ T-cells showed a reduced control of CLL as measured by CD5^+^ CD19^+^ CLL counts in blood over time (Figure 6A). Accordingly, mice that received *Il10rb*^*-/-*^ T-cells had a higher spleen weight in comparison to control mice receiving WT T-cells (Figure 6B). To evaluate whether a reduced expansion of *Il10rb*^*-/-*^ CD4^+^ T-cells contributes to the diminished CLL control, T-cell counts were monitored in blood over time. Three and four weeks after transfer of leukemic cells, a higher number of *Il10Rb*^*-/-*^ versus WT CD4^+^ T-cells was seen in blood (Figure 6C). CD4^+^ T-cell counts in spleen showed a trend towards a higher number of *Il10rb*^*-/-*^ CD4^+^ T-cells compared to WT T-cells per spleen (Figure 6D), which is likely a reflection of the bigger spleen sizes in the *Il10rb*^*-/-*^ group, as the number of CD4^+^ T-cells per CLL cell was reduced in these mice in comparison to the WT group (Figure 6E). Nevertheless, proliferation of CD4^+^ T-cells, as measured by KI-67, did not differ between the two groups (Figure 6F). Hence, IL-10R-mediated signaling in CD4^+^ T-cells is required for their efficient control of CLL development which is not primarily due to an impact on T-cell expansion.

**Figure 6:**
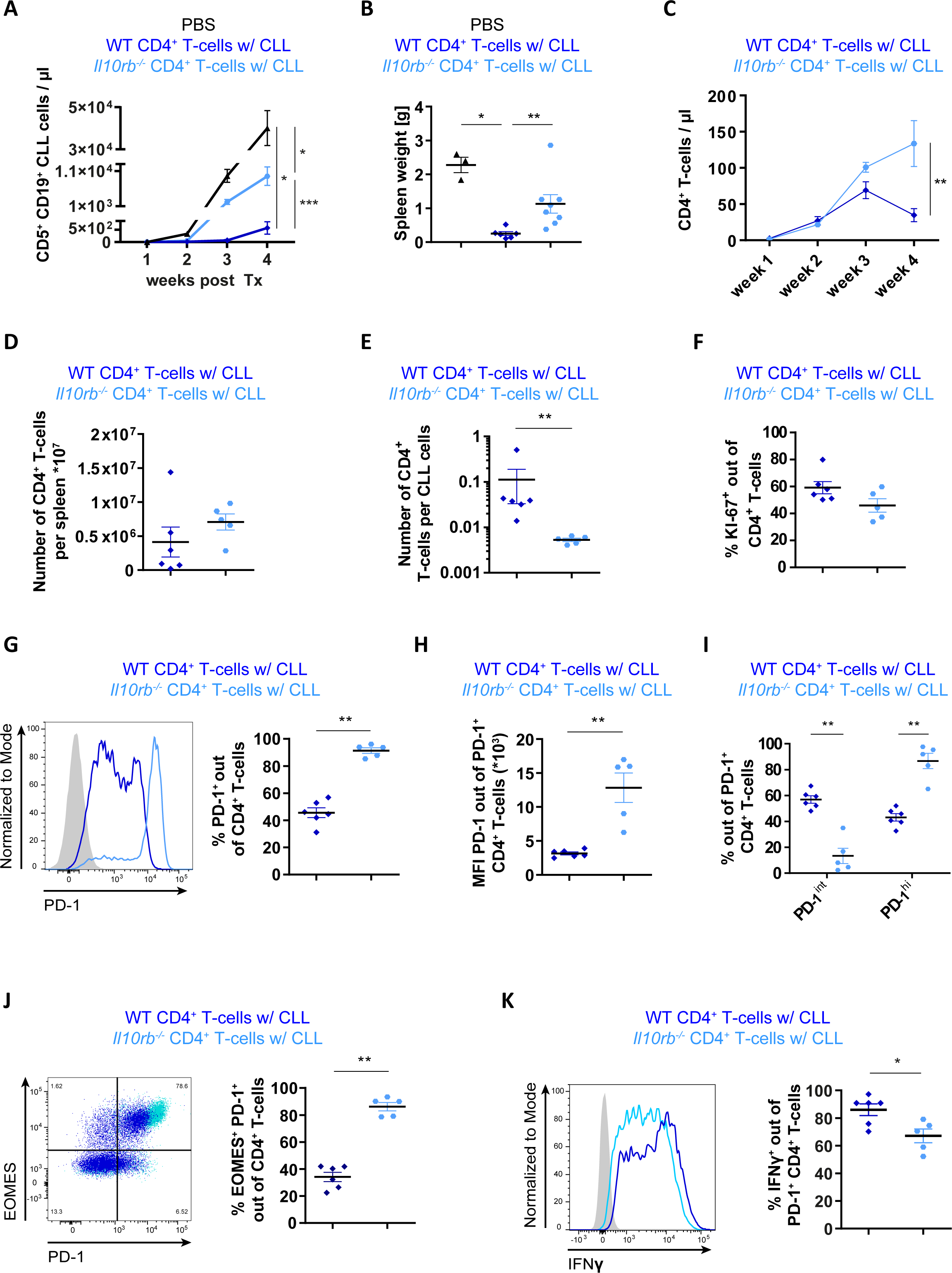
IL-10R signalling controls CLL development by altering EOMES expression of CD4^+^ T-cells. *Rag2*^*-/-*^ mice were transplanted with PBS or CD4^+^ T-cells of WT or *IL10Rb*^*-/-*^ origin on day −1 and the following day with leukemic cells of Eµ-TCL1 mice, and analyzed by flow cytometry. **A)** Absolute numbers of CD5^+^ CD19^+^ CLL cells in peripheral blood are shown over time. **B)** Spleen weight at endpoint, 4 weeks after transfer of leukemic cells. **C)** Absolute numbers of CD4^+^ T-cells cells in peripheral blood are depicted over time. **D)** Number of CD4^+^ T-cells per spleen, as well as **E)** number of CD4^+^ T-cells per CD5^+^ CD19^+^ CLL cell are shown. **F)** Percentage of KI-67^+^ cells out of CD4^+^ T-cells. **G)** Frequency of PD-1^+^ of CD4^+^ T-cells depicted as representative histogram and quantification. **H)** MFI of PD-1 of PD-1-expressing CD4^+^ T-cells. **I)** PD-1^+^ T-cell subsets based on high (PD-1^hi^) or intermediate (PD-1^int^) expression of PD-1 were quantified, and percentages of subsets out of total PD-1^+^ CD4^+^ T-cells are depicted. **J)** Representative graph and frequency of EOMES^+^ PD-1^+^ CD4^+^ T-cells. **K)** Splenocytes were stimulated *ex vivo* with PMA/ionomycin and cytokine expression was analyzed by intracellular flow cytometry. Percentage of IFNγ-producing cells out of PD-1^+^ CD4^+^ T-cells. All graphs show mean ± SEM. In B) and D-K), each dot represents data of an individual mouse. Statistical analysis was performed using Mann-Whitney. *p<0.05, **p<0.01, ***p<0.001. MFI = median fluorescence intensity.

IL-10Rα signaling was shown to be dispensable for the differentiation of T_R_1 T-cells, but not for the function of this cell type (46). Similarly, we investigated the effect of *Il10Rb*^-/-^ on EOMES^+^ PD-1^+^ CD4^+^ T-cells. Intriguingly, loss of *Il10rb* signaling in CD4^+^ T-cells resulted in a higher percentage of PD-1-expressing cells (Figure 6G). Moreover, the expression level of PD-1 was significantly higher in *Il10rb*^*-/-*^ CD4^+^ T-cells compared to WT T-cells (Figure 6H), with almost all *Il10rb*^*-/-*^ CD4^+^ T-cells expressing high levels of PD-1, whereas WT CD4^+^ T-cells showed similar percentages of cells with high or intermediate PD-1 expression (Figure 6I). Furthermore, *Il10rb*^*-/-*^ CD4^+^ T-cells showed a higher frequency of EOMES^+^ PD-1^+^ CD4^+^ T-cells than WT T-cells (Figure 6J). Finally, *Il10rb* deficieny resulted in a reduced capacity of these T-cells to produce IFNγ after *ex vivo* stimulation (Figure 6K).

Taken together, our data suggests that EOMES regulates IL10R signaling in CD4^+^ T-cells and that EOMES and IL10R are necessary to efficiently limit CLL progression.

## Discussion

An altered frequency of CD4^+^ T-cell subsets in B-NHL patients is widely described (1, 3, 5, 6). Among these subsets, particularly IL-10-expressing Tregs were of interest during investigations, as they are thought to mediate immunosuppressive functions and thus contribute to disease progression (6). By analyzing blood samples of CLL and DLBCL patients, we observed a higher proportion of PD-1-expressing CD4^+^ T-cells in both patient groups compared to healthy controls thus confirming published data (1, 4, 11-13). Moreover, we noted a population of EOMES^+^ PD-1^+^ CD4^+^ T-cells in LN samples of CLL as well as DLBCL patients. In the follicles of secondary lymphoid organs, tight interactions of T- and malignant B-cells take place, which lead to activation and, in case of persistent exposure to antigens, to T-cell exhaustion (37, 47) which might contribute to the observed phenotype of T-cells in CLL and DLBCL LNs.

CD4^+^ T-cells expressing high amounts of the degranulation marker protein CD107a and effector molecules like perforines and granzymes were found in blood of B-NHL patients (48-50). Moreover, CD4^+^ T-cells that were co-cultured with autologous B-cells, either from CLL patients or healthy controls, had the capacity to kill autologous B-cells (48). Similar to our approach, co-transfer of naïve, tumor-specific CD4^+^ T-cells in a transplantation mouse model of B16 melanoma significantly prolonged survival of mice, which was further enhanced by anti-CTLA-4 treatment (51). A cytotoxic activity of these CD4^+^ T-cells was suggested as they acquired an effector phenotype with high cytokine production (51). Similar results were obtained in *Rag2*^*-/-*^ mice, indicating that CD4^+^ T-cell-mediated tumor control was independent of endogenous B- and T-cells (51), which is in line with our data in the TCL1 AT mouse model of CLL.

In our study, EOMES was shown to be essential for the control of leukemia progression. EOMES is a transcription factor that is crucial in memory formation of CD8^+^ T-cells (27), but not necessary to induce their effector function during viral infections (21). In contrast, high EOMES expression was shown to result in terminal differentiation of CD8^+^ T-cells (28). Here, we provide evidence that EOMES is indispensable for CLL control by CD4^+^ T-cells most likely due to its role in their effector function, which is in line with reports demonstrating EOMES-dependency for granzyme B production of cytotoxic CD4^+^ T-cells (24, 25, 52).

Besides its role in regulating cytotoxic activity of CD4^+^ T-cells, the importance of EOMES in the generation of T_R_1 T-cells was recently shown (7, 8). T_R_1 T-cells do not constitutively express FOXP3, produce the immunosuppressive cytokine IL-10, express co-inhibitory receptors such as PD-1, and are able to suppress the function of effector immune cells (10). The expression of IL-10 and other cytokines and their receptors, in combination with increased expression levels of inhibitory receptors are shared features of CD4^+^ EOMES^+^ T-cells in B-NHL and T_R_1 cells.

In line with published data demonstrating that EOMES regulates IL-10 expression (7, 8, 53), our data in the TCL1 mouse model of CLL show that IL-10 production of CD4^+^ T-cells is dependent on EOMES. In addition, IL-10 receptor expression is higher in EOMES-positive than -negative CD4^+^ T-cells, which is in accordance with data of CD8^+^ T-cells (53). Since IL-10-driven signaling via *p38 MAPK* was shown to be important to maintain IL-10 production in T_R_1 CD4^+^ T-cells (46), we investigated the role of IL-10R signaling in CD4^+^ T-cells. Intriguingly, *Il10rb*^*-/-*^ CD4^+^ T-cells showed a reduced CLL control alongside with a high expression of PD-1 as well as EOMES. This increase in PD-1 and EOMES expression was accompanied by a reduction in IFNγ production, suggesting that IL-10R signaling is additionally involved in the regulation of the effector activity of CD4^+^ T-cells. This is in line with published results for CD8^+^ T-cells, showing that overexpression of EOMES resulted in an increased expression of exhaustion molecules such as *CD244, Havcr2* as well as *Il10ra*, implicating a role for IL-10-mediated signaling in regulating T-cell exhaustion (54). Moreover, during murine chronic viral infections, CD4^+^ T-cells upregulate EOMES as well as inhibitory receptors that are associated with T-cell exhaustion (43). Studies in such infection models as well as from murine and human cancer showed that the expression level of PD-1 in CD8^+^ T-cells determines their state of exhaustion and potential for reinvigoration by PD-1 blockade (55). Very similar to these data, we detected CD4^+^ T-cells with intermediate or high PD-1 expression in the TCL1 mouse model. Interestingly, loss of IL-10R signaling in these cells resulted in an accumulation of PD-1^hi^ cells with reduced IFNγ production and impaired CLL control. This suggests that IL-10-mediated signals are important to maintain CD4^+^ T-cell effector function.

Genome-wide assosciation studies (GWAS) showed that single nucleotide polymorphisms (SNPs) in proximitiy of the *EOMES* gene, which is located at chromosome 3p24.1, are associated with a higher risk of CLL (rs9880772) (56, 57), DLBCL (rs6773363) (58) as well as Hodgkin’s Lymphoma (rs3806624) (59). The higher likelihood of B-NHL in individuals carrying these SNPs was thought to be caused by a deregulated immune function, which could at least partially be explained by the reduced control of CLL development in the absence of CD8^+^ T-cells. Alongside, preclinical data in mouse models (60), as well as data of a phase 1 basket trial (NCT02009449) using pegylated IL-10 for treatment of solid cancer demonstrated that IL-10 helps to maintain CD8^+^ T-cell mediated tumor control and improves patients’ responses to PD-1 blockade (45).

In summary, this report highlights the presence of EOMES-expressing PD-1^+^ CD4^+^ T-cells in LNs of CLL and DLBCL patients as well as in the TCL1 mouse model of CLL. Our data in this animal model clearly show that EOMES is crucial for CD4^+^ T-cell-mediated disease control. As EOMES regulates IL-10 production, we further demonstrate a role for IL-10-mediated signaling in EOMES-expressing CD4^+^ T-cells, which mediate effector activity and thus control of leukemia.

## Acknowledgments

This study was supported by the German Research Foundation project EV-RNA (SE 2331/2-1) and by the German José Carreras Foundation (grant 13R/2018) to MS. PMR was supported by the German Cancer Aid grant number 112069. EL and AI were supported by Bundesministerium für Bildung und Forschung (grant BMBF 01 EO 1303) and EL, FK, MK and AI were supported by the Max Planck Society. SJA was supported by the German Research Foundation (AR 732/2-1, AR 732/3-1), project A03 of SFB 850 (project ID 89986987) and Germany’s Excellence Strategy (CIBSS – EXC-2189 – Project ID 390939984). St.St. was supported by DFG SFB1074 subproject B1.

## Author contributions

PMR, LLC and EL designed the study, performed experiments, analyzed and interpreted data, prepared figures, and wrote the manuscript. TR, MB, CS, MK and ACG performed experiments. FK performed bioinformatics analysis. LS, StSt and SD provided clinical samples and information. SJA provided mice and advised the study. PL critically advised the study and reviewed the manuscript. AI and MS designed and supervised the study, interpreted data, and wrote the manuscript.

## References

1. Zhang L, Du H, Xiao TW, Liu JZ, Liu GZ, Wang JX, et al. Prognostic value of PD-1 and TIM-3 on CD3+ T cells from diffuse large B-cell lymphoma. Biomed Pharmacother. 2015;75:83–7.

2. Kocher T, Asslaber D, Zaborsky N, Flenady S, Denk U, Reinthaler P, et al. CD4+ T cells, but not non-classical monocytes, are dispensable for the development of chronic lymphocytic leukemia in the TCL1-tg murine model. Leukemia. 2016;30(6):1409–13.

3. Bagnara D, Kaufman MS, Calissano C, Marsilio S, Patten PE, Simone R, et al. A novel adoptive transfer model of chronic lymphocytic leukemia suggests a key role for T lymphocytes in the disease. Blood. 2011;117(20):5463–72.

4. Brusa D, Serra S, Coscia M, Rossi D, D’Arena G, Laurenti L, et al. The PD-1/PD-L1 axis contributes to T-cell dysfunction in chronic lymphocytic leukemia. Haematologica. 2013;98(6):953–63.

5. Roessner PM, Hanna BS, Ozturk S, Schulz R, Llao Cid L, Yazdanparast H, et al. TBET-expressing Th1 CD4(+) T cells accumulate in chronic lymphocytic leukaemia without affecting disease progression in Emicro-TCL1 mice. British journal of haematology. 2019.

6. Hanna BS, Roessner PM, Scheffold A, Jebaraj BMC, Demerdash Y, Ozturk S, et al. PI3Kdelta inhibition modulates regulatory and effector T-cell differentiation and function in chronic lymphocytic leukemia. Leukemia. 2019;33(6):1427–38.

7. Zhang P, Lee JS, Gartlan KH, Schuster IS, Comerford I, Varelias A, et al. Eomesodermin promotes the development of type 1 regulatory T (TR1) cells. Sci Immunol. 2017;2(10).

8. Gruarin P, Maglie S, De Simone M, Haringer B, Vasco C, Ranzani V, et al. Eomesodermin controls a unique differentiation program in human IL-10 and IFN-gamma coproducing regulatory T cells. European journal of immunology. 2019;49(1):96–111.

9. Brockmann L, Soukou S, Steglich B, Czarnewski P, Zhao L, Wende S, et al. Molecular and functional heterogeneity of IL-10-producing CD4(+) T cells. Nat Commun. 2018;9(1):5457.

10. Roncarolo MG, Gregori S, Bacchetta R, Battaglia M, Gagliani N. The Biology of T Regulatory Type 1 Cells and Their Therapeutic Application in Immune-Mediated Diseases. Immunity. 2018;49(6):1004–19.

11. Muenst S, Hoeller S, Willi N, Dirnhofera S, Tzankov A. Diagnostic and prognostic utility of PD-1 in B cell lymphomas. Disease markers. 2010;29(1):47–53.

12. Palma M, Gentilcore G, Heimersson K, Mozaffari F, Nasman-Glaser B, Young E, et al. T cells in chronic lymphocytic leukemia display dysregulated expression of immune checkpoints and activation markers. Haematologica. 2017;102(3):562–72.

13. Riches JC, Davies JK, McClanahan F, Fatah R, Iqbal S, Agrawal S, et al. T cells from CLL patients exhibit features of T-cell exhaustion but retain capacity for cytokine production. Blood. 2013;121(9):1612–21.

14. McClanahan F, Hanna B, Miller S, Clear AJ, Lichter P, Gribben JG, et al. PD-L1 checkpoint blockade prevents immune dysfunction and leukemia development in a mouse model of chronic lymphocytic leukemia. Blood. 2015;126(2):203–11.

15. Ding W, LaPlant BR, Call TG, Parikh SA, Leis JF, He R, et al. Pembrolizumab in patients with CLL and Richter transformation or with relapsed CLL. Blood. 2017;129(26):3419–27.

16. Fang X, Xiu B, Yang Z, Qiu W, Zhang L, Zhang S, et al. The expression and clinical relevance of PD-1, PD-L1, and TP63 in patients with diffuse large B-cell lymphoma. Medicine (Baltimore). 2017;96(15):e6398.

17. Ahearne MJ, Bhuller K, Hew R, Ibrahim H, Naresh K, Wagner SD. Expression of PD-1 (CD279) and FoxP3 in diffuse large B-cell lymphoma. Virchows Arch. 2014;465(3):351–8.

18. Zhang J, Medeiros LJ, Young KH. Cancer Immunotherapy in Diffuse Large B-Cell Lymphoma. Frontiers in oncology. 2018;8:351.

19. Naiche LA, Harrelson Z, Kelly RG, Papaioannou VE. T-box genes in vertebrate development. Annu Rev Genet. 2005;39:219–39.

20. Cruz-Guilloty F, Pipkin ME, Djuretic IM, Levanon D, Lotem J, Lichtenheld MG, et al. Runx3 and T-box proteins cooperate to establish the transcriptional program of effector CTLs. The Journal of experimental medicine. 2009;206(1):51–9.

21. Pearce EL, Mullen AC, Martins GA, Krawczyk CM, Hutchins AS, Zediak VP, et al. Control of effector CD8+ T cell function by the transcription factor Eomesodermin. Science. 2003;302(5647):1041–3.

22. Intlekofer AM, Takemoto N, Wherry EJ, Longworth SA, Northrup JT, Palanivel VR, et al. Effector and memory CD8+ T cell fate coupled by T-bet and eomesodermin. Nat Immunol. 2005;6(12):1236–44.

23. Hegel JK, Knieke K, Kolar P, Reiner SL, Brunner-Weinzierl MC. CD152 (CTLA-4) regulates effector functions of CD8+ T lymphocytes by repressing Eomesodermin. European journal of immunology. 2009;39(3):883–93.

24. Qui HZ, Hagymasi AT, Bandyopadhyay S, St Rose MC, Ramanarasimhaiah R, Menoret A, et al. CD134 plus CD137 dual costimulation induces Eomesodermin in CD4 T cells to program cytotoxic Th1 differentiation. Journal of immunology (Baltimore, Md: 1950). 2011;187(7):3555–64.

25. Curran MA, Geiger TL, Montalvo W, Kim M, Reiner SL, Al-Shamkhani A, et al. Systemic 4-1BB activation induces a novel T cell phenotype driven by high expression of Eomesodermin. The Journal of experimental medicine. 2013;210(4):743–55.

26. Gordon SM, Chaix J, Rupp LJ, Wu J, Madera S, Sun JC, et al. The transcription factors T-bet and Eomes control key checkpoints of natural killer cell maturation. Immunity. 2012;36(1):55–67.

27. Banerjee A, Gordon SM, Intlekofer AM, Paley MA, Mooney EC, Lindsten T, et al. Cutting edge: The transcription factor eomesodermin enables CD8+ T cells to compete for the memory cell niche. Journal of immunology (Baltimore, Md: 1950). 2010;185(9):4988–92.

28. Paley MA, Kroy DC, Odorizzi PM, Johnnidis JB, Dolfi DV, Barnett BE, et al. Progenitor and terminal subsets of CD8+ T cells cooperate to contain chronic viral infection. Science. 2012;338(6111):1220–5.

29. Llao Cid L, Hanna BS, Iskar M, Roessner PM, Ozturk S, Lichter P, et al. CD8(+) T-cells of CLL-bearing mice acquire a transcriptional program of T-cell activation and exhaustion. Leuk Lymphoma. 2020;61(2):351–6.

30. Lupar E, Brack M, Garnier L, Laffont S, Rauch KS, Schachtrup K, et al. Eomesodermin Expression in CD4+ T Cells Restricts Peripheral Foxp3 Induction. Journal of immunology (Baltimore, Md: 1950). 2015;195(10):4742–52.

31. Suto A, Wurster AL, Reiner SL, Grusby MJ. IL-21 inhibits IFN-gamma production in developing Th1 cells through the repression of Eomesodermin expression. Journal of immunology (Baltimore, Md: 1950). 2006;177(6):3721–7.

32. Steiner DF, Thomas MF, Hu JK, Yang Z, Babiarz JE, Allen CD, et al. MicroRNA-29 regulates T-box transcription factors and interferon-gamma production in helper T cells. Immunity. 2011;35(2):169–81.

33. Yang Y, Xu J, Niu Y, Bromberg JS, Ding Y. T-bet and eomesodermin play critical roles in directing T cell differentiation to Th1 versus Th17. Journal of immunology (Baltimore, Md: 1950). 2008;181(12):8700–10.

34. Mazzoni A, Maggi L, Siracusa F, Ramazzotti M, Rossi MC, Santarlasci V, et al. Eomes controls the development of Th17-derived (non-classic) Th1 cells during chronic inflammation. European journal of immunology. 2019;49(1):79–95.

35. Dejean AS, Joulia E, Walzer T. The role of Eomes in human CD4 T cell differentiation: A question of context. European journal of immunology. 2019;49(1):38–41.

36. Catovsky D, Miliani E, Okos A, Galton DA. Clinical significance of T-cells in chronic lymphocytic leukaemia. Lancet (London, England). 1974;2(7883):751–2.

37. Hanna BS, Roessner PM, Yazdanparast H, Colomer D, Campo E, Kugler S, et al. Control of chronic lymphocytic leukemia development by clonally-expanded CD8(+) T-cells that undergo functional exhaustion in secondary lymphoid tissues. Leukemia. 2019;33(3):625–37.

38. Herishanu Y, Perez-Galan P, Liu D, Biancotto A, Pittaluga S, Vire B, et al. The lymph node microenvironment promotes B-cell receptor signaling, NF-kappaB activation, and tumor proliferation in chronic lymphocytic leukemia. Blood. 2011;117(2):563–74.

39. Mempel TR, Henrickson SE, Von Andrian UH. T-cell priming by dendritic cells in lymph nodes occurs in three distinct phases. Nature. 2004;427(6970):154–9.

40. Ozturk S, Roessner PM, Schulze-Edinghausen L, Yazdanparast H, Kalter V, Lichter P, et al. Rejection of adoptively transferred Emicro-TCL1 chronic lymphocytic leukemia cells in C57BL/6 substrains or knockout mouse lines. Leukemia. 2019;33(6):1514–39.

41. Shinkai Y, Rathbun G, Lam KP, Oltz EM, Stewart V, Mendelsohn M, et al. RAG-2-deficient mice lack mature lymphocytes owing to inability to initiate V(D)J rearrangement. Cell. 1992;68(5):855–67.

42. Arnold SJ, Sugnaseelan J, Groszer M, Srinivas S, Robertson EJ. Generation and analysis of a mouse line harboring GFP in the Eomes/Tbr2 locus. Genesis (New York, NY: 2000). 2009;47(11):775–81.

43. Crawford A, Angelosanto JM, Kao C, Doering TA, Odorizzi PM, Barnett BE, et al. Molecular and transcriptional basis of CD4(+) T cell dysfunction during chronic infection. Immunity. 2014;40(2):289–302.

44. Sun Z, Fourcade J, Pagliano O, Chauvin JM, Sander C, Kirkwood JM, et al. IL10 and PD-1 Cooperate to Limit the Activity of Tumor-Specific CD8+ T Cells. Cancer Res. 2015;75(8):1635–44.

45. Naing A, Infante JR, Papadopoulos KP, Chan IH, Shen C, Ratti NP, et al. PEGylated IL-10 (Pegilodecakin) Induces Systemic Immune Activation, CD8(+) T Cell Invigoration and Polyclonal T Cell Expansion in Cancer Patients. Cancer Cell. 2018;34(5):775–91 e3.

46. Brockmann L, Gagliani N, Steglich B, Giannou AD, Kempski J, Pelczar P, et al. IL-10 Receptor Signaling Is Essential for TR1 Cell Function In Vivo. Journal of immunology (Baltimore, Md: 1950). 2017;198(3):1130–41.

47. de Weerdt I, Hofland T, de Boer R, Dobber JA, Dubois J, van Nieuwenhuize D, et al. Distinct immune composition in lymph node and peripheral blood of CLL patients is reshaped during venetoclax treatment. Blood advances. 2019;3(17):2642–52.

48. Lindqvist CA, Christiansson LH, Thorn I, Mangsbo S, Paul-Wetterberg G, Sundstrom C, et al. Both CD4+ FoxP3+ and CD4+ FoxP3-T cells from patients with B-cell malignancy express cytolytic markers and kill autologous leukaemic B cells in vitro. Immunology. 2011;133(3):296–306.

49. Porakishvili N, Roschupkina T, Kalber T, Jewell AP, Patterson K, Yong K, et al. Expansion of CD4+ T cells with a cytotoxic phenotype in patients with B-chronic lymphocytic leukaemia (B-CLL). Clin Exp Immunol. 2001;126(1):29–36.

50. Yang ZZ, Kim HJ, Villasboas JC, Chen YP, Price-Troska T, Jalali S, et al. Expression of LAG-3 defines exhaustion of intratumoral PD-1(+) T cells and correlates with poor outcome in follicular lymphoma. Oncotarget. 2017;8(37):61425–39.

51. Quezada SA, Simpson TR, Peggs KS, Merghoub T, Vider J, Fan X, et al. Tumor-reactive CD4(+) T cells develop cytotoxic activity and eradicate large established melanoma after transfer into lymphopenic hosts. The Journal of experimental medicine. 2010;207(3):637–50.

52. Patil VS, Madrigal A, Schmiedel BJ, Clarke J, O’Rourke P, de Silva AD, et al. Precursors of human CD4(+) cytotoxic T lymphocytes identified by single-cell transcriptome analysis. Sci Immunol. 2018;3(19).

53. Reiser J, Sadashivaiah K, Furusawa A, Banerjee A, Singh N. Eomesodermin driven IL-10 production in effector CD8(+) T cells promotes a memory phenotype. Cellular immunology. 2019;335:93–102.

54. Li J, He Y, Hao J, Ni L, Dong C. High Levels of Eomes Promote Exhaustion of Anti-tumor CD8(+) T Cells. Frontiers in immunology. 2018;9(2981):2981.

55. Im SJ, Hashimoto M, Gerner MY, Lee J, Kissick HT, Burger MC, et al. Defining CD8+ T cells that provide the proliferative burst after PD-1 therapy. Nature. 2016;537(7620):417–21.

56. Berndt SI, Camp NJ, Skibola CF, Vijai J, Wang Z, Gu J, et al. Meta-analysis of genome-wide association studies discovers multiple loci for chronic lymphocytic leukemia. Nat Commun. 2016;7(1):10933.

57. Law PJ, Sud A, Mitchell JS, Henrion M, Orlando G, Lenive O, et al. Genome-wide association analysis of chronic lymphocytic leukaemia, Hodgkin lymphoma and multiple myeloma identifies pleiotropic risk loci. Scientific reports. 2017;7(1):41071.

58. Kleinstern G, Yan H, Hildebrandt MAT, Vijai J, Berndt SI, Ghesquieres H, et al. Inherited variants at 3q13.33 and 3p24.1 are associated with risk of diffuse large B-cell lymphoma and implicate immune pathways. Human molecular genetics. 2020;29(1):70–9.

59. Frampton M, da Silva Filho MI, Broderick P, Thomsen H, Forsti A, Vijayakrishnan J, et al. Variation at 3p24.1 and 6q23.3 influences the risk of Hodgkin’s lymphoma. Nat Commun. 2013;4:2549.

60. Mumm JB, Emmerich J, Zhang X, Chan I, Wu L, Mauze S, et al. IL-10 elicits IFNgamma-dependent tumor immune surveillance. Cancer Cell. 2011;20(6):781–96.

