## Supplementary Materials and Methods for "EOMES and IL-10 regulate anti-tumor activity of PD-1^+^ CD4^+^ T-cells in B-cell Non-Hodgkin lymphoma"

### Supplementary Material and Methods

*Processing of patient samples*

Peripheral blood (PB) was drawn using ethylene diamine tetraacetic acid (EDTA)-coated tubes (Sarstedt, Nümbrecht, Germany). PB mononuclear cells (PBMCs) were isolated by Ficoll (Biochrom, Berlin, Germany) density gradient centrifugation. If necessary, PBMCs were viably frozen and when needed, frozen PBMCs were thawed and rested for three hours until further processing. Lymph node (LN) samples were processed as previously described (1).

*Collection of murine tissue samples and preparation of cell suspensions*

Mice were euthanized by increasing concentrations of carbon dioxide ($CO_2$). PB was drawn from the submandibular vein or via cardiac puncture and collected in EDTA-coated tubes (Sarstedt). Single-cell suspensions from spleens, bone marrow and inguinal LNs were prepared as previously described (2, 3).

*Mice*

*Eomesodermin* (*Eomes*) conditional allele mice that harbour loxP sites flanking the exons 2-5 encoding the T-box DNA-binding domain, were previously described (4). In order to achieve a T-cell-specific deletion of *Eomes*, these mice were crossed to a transgenic *Cre* line in which the expression of *Cre* recombinase is regulated by the mouse proximal *Lck* promoter (5). Mice carrying a *GFP* reporter allele of *Eomes* were previously described (6). This mouse line was crossed with the *Eomes*$^{fl/fl}$ *x Lck-cre* strain to generate *Eomes*$^{\Delta T/GFP}$ knock-out mice. Moreover, *Lck-cre x Eomes*$^{fl/fl}$ *x Foxp3-IRES-mRFP (FIR)* (7) x *Il10-GFP (tiger)* (8) (*Eomes*$^{-/-}$ *knock-out* mice) or *Eomes*$^{fl/fl}$ *x FIR x tige*r (WT) mice were used for TCL1 leukemia transfer. All mouse strains described above were maintained under conventional or specific pathogen-free conditions at Max-Planck Institute of Immunobiology and Epigenetics (Freiburg, Germany). Eμ-TCL1 (C. Croce, OH, USA) mice were held at specific pathogen-free conditions on a pure C57BL/6N or J background at the central animal facility of the German Cancer Research Center (DKFZ).

*Flow cytometric analysis and cell sorting*

Whole blood stainings were performed by addition of the surface antibody cocktail to a defined volume of blood. Red blood cell lysis was performed by addition of 1x Red blood cell lysis buffer (BD Bioscience, Heidelberg, Germany or ThermoFisher Scientific, Dreieich, Germany) and subsequent incubation at room temperature. After centrifugation, cell suspension was resuspended in 1x Red blood cell lysis buffer and 123count eBeads™ Counting Beads (ThermoFisher Scientific) were added to determine cell concentrations by the formula: Absolute Count (cells/μl) = (Cell count x eBead™ Volume) / (eBead™ Count x Cell Volume) x (eBead™ concentration).

Single cell suspensions were stained in phosphate-buffered saline (PBS) with addition Fixable Viability Dye eFluor® 506 (ThermoFisher Scientific) at a concentration of 1:1000 for 30 min at 4°C. Cells were fixed using eBioscience™ Foxp3 / Transcription Factor Staining Buffer Set (ThermoFisher Scientific) for 30 min at room temperature. Subsequently, cells were permeabilized with eBioscience™ Permeabilization Buffer (ThermoFisher Scientific) and stained intracellularly for 30 min at room temperature. Samples were stored at 4°C in the dark until acquisition.

Cytokine stainings were performed as previously described (3, 9, 10). Briefly, cells were stimulated using either 1x PMA/ionomycin-based eBioscience™ Cell Stimulation and Inhibitor Cocktail (ThermoFisher Scientific) or a stimulation cocktail containing 0.1 µg/ml of PMA (Enzo Life Sciences, Lausen, Switzerland), 1 µg/ml of ionomycin (SERVA Electrophoresis, Heidelberg, Germany) and 1x protein transport inhibitor Monensin (BioLegend, London, United Kingdom) for 4-6 hours at 37°C and 5% $CO_2$. Subsequently, cells were washed, surface stained, fixed and permeabilized as detailed above.

Flow cytometry data was acquired using a BD FACS Canto II, BD LSR II or BD LSR Fortessa (BD Biosciences) FACS analyzer and analyzed using FlowJo X 10.0.7 software (FlowJo, Ashland, OR, USA).

Fluorescence-activated cell sorting of naive $CD25^-$ $CD45RB^{hi}$ $CD4^+$ T-cells or of activated $GFP^+$ and $GFP^-$ $CD4^+$ splenic T-cells (purity typically >98%) after 3 weeks of adoptive transfer was performed using BD FACSAria™ III or BD FACSAria™ Fusion (BD Biosciences) instruments as previously described (3).

Data analysis and graphical display were performed using Prism 7 GraphPad software (GraphPad Software, La Jolla, USA).

*RNA sequencing and analysis*

For RNA sequencing, total RNA was extracted with TRI reagent (Sigma-Aldrich, Munich, Germany) according to the manufacturer's instructions. Depletion of ribosomal RNA and library preparation was performed with TruSeq® Stranded Total RNA Gold kit (former name TruSeq® Stranded Total RNA LT - (with Ribo-Zero TM GOLD), Illumina, San Diego, USA). The obtained RNA was quality-controlled in a Fragment Analyzer (Agilent Technologies, Santa Clara, USA) and sequenced with an Illumina HiSeq2500 in a 2 x 75 bp paired end, multiplexing run, aiming for ~25 million reads per sample. Raw data from the Illumina HiSeq 2500 sequencing machine was demultiplexed and converted into FASTQ files using Illumina bcl2fastq2 (version 1.8.4, http://support.illumina.com/downloads/bcl2fastq_conversion_software_184.html). Data was screened for contamination with fastq_screen (version 0.5.1, (http://www.bioinformatics.babraham.ac.uk/projects/fastq_screen/). Automatic detection and trimming of adaptors, quality control and mapping was performed with an rna-seq-qc pipeline that is available on GitHub (https://github.com/maxplanck-ie/rna-seq-qc). We used TopHat2 (version 2.0.13) (11) to align the reads against mouse genome build mm10. Reads were counted with featureCounts (version 1.5.0-p1). Differential gene expression analysis was performed using the DESeq2 (version 1.6.1) (12). We considered genes with adjusted p-values less than 0.05 as significantly differentially expressed. Heatmaps were created with Genepattern (13). Heatmap “exhaustion markers” was created according to genes expressed by exhausted $CD4^+$ T-cells (14) and overlapping with the GO term cell surface (GO:0009986). Heatmaps for cytokine activity (GO:005125) and cytokine receptor activity (GO:0004896) were prepared according to the respective GO terms. Genes not corresponding to the definition of cytokine activity or cytokine receptor activity were manually removed (listed in Supplementary Table 5). Comparison of RNA-Seq data with the published human $T_R1$ signature (15) was performed with gene set enrichment analysis (GSEA) as described (16, 17). The dataset is available on Gene Expression Omnibus (GEO) under the accession numbers GSE145145.

### References


1. Roider T, Seufert J, Uvarovskii A, Frauhammer F, Bordas M, Abedpour N, et al. Dissecting intratumor heterogeneity of nodal B cell lymphomas on the transcriptional, genetic, and drug response level. bioRxiv. 2019:850438.
2. Hanna BS, McClanahan F, Yazdanparast H, Zaborsky N, Kalter V, Rossner PM, et al. Depletion of CLL-associated patrolling monocytes and macrophages controls disease development and repairs immune dysfunction in vivo. Leukemia. 2016;30(3):570-9.
3. Lupar E, Brack M, Garnier L, Laffont S, Rauch KS, Schachtrup K, et al. Eomesodermin Expression in CD4+ T Cells Restricts Peripheral Foxp3 Induction. Journal of immunology (Baltimore, Md : 1950). 2015;195(10):4742-52.
4. Arnold SJ, Hofmann UK, Bikoff EK, Robertson EJ. Pivotal roles for eomesodermin during axis formation, epithelium-to-mesenchyme transition and endoderm specification in the mouse. Development. 2008;135(3):501-11.
5. Orban PC, Chui D, Marth JD. Tissue- and site-specific DNA recombination in transgenic mice. Proceedings of the National Academy of Sciences of the United States of America. 1992;89(15):6861-5.
6. Arnold SJ, Sugnaseelan J, Groszer M, Srinivas S, Robertson EJ. Generation and analysis of a mouse line harboring GFP in the Eomes/Tbr2 locus. Genesis (New York, NY : 2000). 2009;47(11):775-81.
7. Wan YY, Flavell RA. Identifying Foxp3-expressing suppressor T cells with a bicistronic reporter. Proceedings of the National Academy of Sciences of the United States of America. 2005;102(14):5126-31.
8. Kamanaka M, Kim ST, Wan YY, Sutterwala FS, Lara-Tejero M, Galan JE, et al. Expression of interleukin-10 in intestinal lymphocytes detected by an interleukin-10 reporter knockin tiger mouse. Immunity. 2006;25(6):941-52.
9. Hanna BS, Roessner PM, Scheffold A, Jebaraj BMC, Demerdash Y, Ozturk S, et al. PI3Kdelta inhibition modulates regulatory and effector T-cell differentiation and function in chronic lymphocytic leukemia. Leukemia. 2019;33(6):1427-38.
10. Hanna BS, Roessner PM, Yazdanparast H, Colomer D, Campo E, Kugler S, et al. Control of chronic lymphocytic leukemia development by clonally-expanded CD8(+) T-cells that undergo functional exhaustion in secondary lymphoid tissues. Leukemia. 2019;33(3):625-37.
11. Kim D, Pertea G, Trapnell C, Pimentel H, Kelley R, Salzberg SL. TopHat2: accurate alignment of transcriptomes in the presence of insertions, deletions and gene fusions. Genome Biol. 2013;14(4):R36.
12. Anders S, Pyl PT, Huber W. HTSeq--a Python framework to work with high-throughput sequencing data. Bioinformatics (Oxford, England). 2015;31(2):166-9.
13. Reich M, Liefeld T, Gould J, Lerner J, Tamayo P, Mesirov JP. GenePattern 2.0. Nature genetics. 2006;38(5):500-1.
14. Crawford A, Angelosanto JM, Kao C, Doering TA, Odorizzi PM, Barnett BE, et al. Molecular and transcriptional basis of CD4(+) T cell dysfunction during chronic infection. Immunity. 2014;40(2):289-302.
15. Gruarin P, Maglie S, De Simone M, Haringer B, Vasco C, Ranzani V, et al. Eomesodermin controls a unique differentiation program in human IL-10 and IFN-gamma coproducing regulatory T cells. European journal of immunology. 2019;49(1):96-111.
16. Subramanian A, Tamayo P, Mootha VK, Mukherjee S, Ebert BL, Gillette MA, et al. Gene set enrichment analysis: a knowledge-based approach for interpreting genome-wide expression profiles. Proceedings of the National Academy of Sciences of the United States of America. 2005;102(43):15545-50.
17. Mootha VK, Lindgren CM, Eriksson KF, Subramanian A, Sihag S, Lehar J, et al. PGC-1alpha-responsive genes involved in oxidative phosphorylation are coordinately downregulated in human diabetes. Nature genetics. 2003;34(3):267-73.

**Supplementary Table 1: Clinical information for whole blood samples of CLL patients and healthy controls**

| | HC | CLL |
|---|---|---|
| **Number of samples** | 19 | 17 |
| **Sex** | 26.3% female (5/19) | 41.2% female (7/17) |
| **Age (years)** | mean: 61.2<br>median: 61 | mean: 60.4<br>median: 60 |
| **CMV IgG positivity** | 55.6% CMV $IgG^{+}$ (10/18) | 40.0% CMV $IgG^{+}$ (6/15) |
| **Binet stage** | | 70.6% A (12/17)<br>17.6% B (3/17)<br>11.8% C (2/17) |
| **Mutational state of *IGHV*** | | 58.8% mutated (10/17) |
| **Chromosomal aberration** | | 64.7% del13q14.3 (11/17)<br>23.5% normal karyotype (4/17)<br>5.9% trisomy 12 (1/17)<br>5.9% IGH translocation |
| **TP53 mutation** | | 20.0% (2/10) |
| **Prior treatment** | | 0.0% (0/17) |

**Supplementary Table 2: Clinical information for whole blood samples of DLBCL patients and healthy controls**

| | HC | DLBCL |
|---|---|---|
| **Number of samples** | 19 | 18 |
| **Age (years)** | mean: 61.2<br>median: 61 | mean: 66.29 (17/18)<br>median: 65 (17/18) |
| **Relapsed** | | 5.6% (1/18) |

**Supplementary Table 3: Clinical information for lymph node samples of CLL and DLBCL patients**

| | CLL | DLBCL |
|---|---|---|
| **Number of samples** | 7 | 7 |
| **Sub-classification** | | Germinal center B-cell:<br>12.5% (1/7)<br>Non-germinal center B-cell:<br>85.7% (6/7) |

### Supplementary Figures

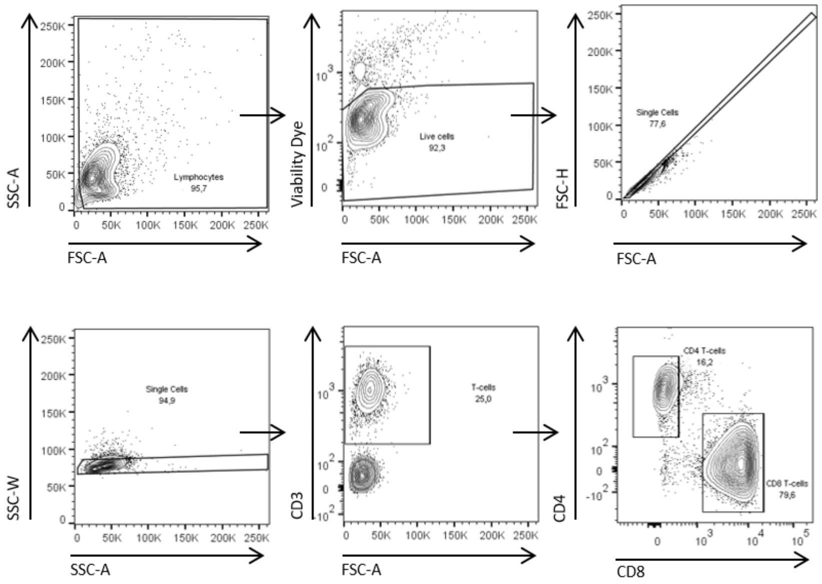


**Suppl. Figure 1: Representative gating strategy to define CD4⁺ T-cells**

Representative gating strategy for flow cytometry analysis to define viable, single CD4⁺ T-cells in lymph node samples of patients.

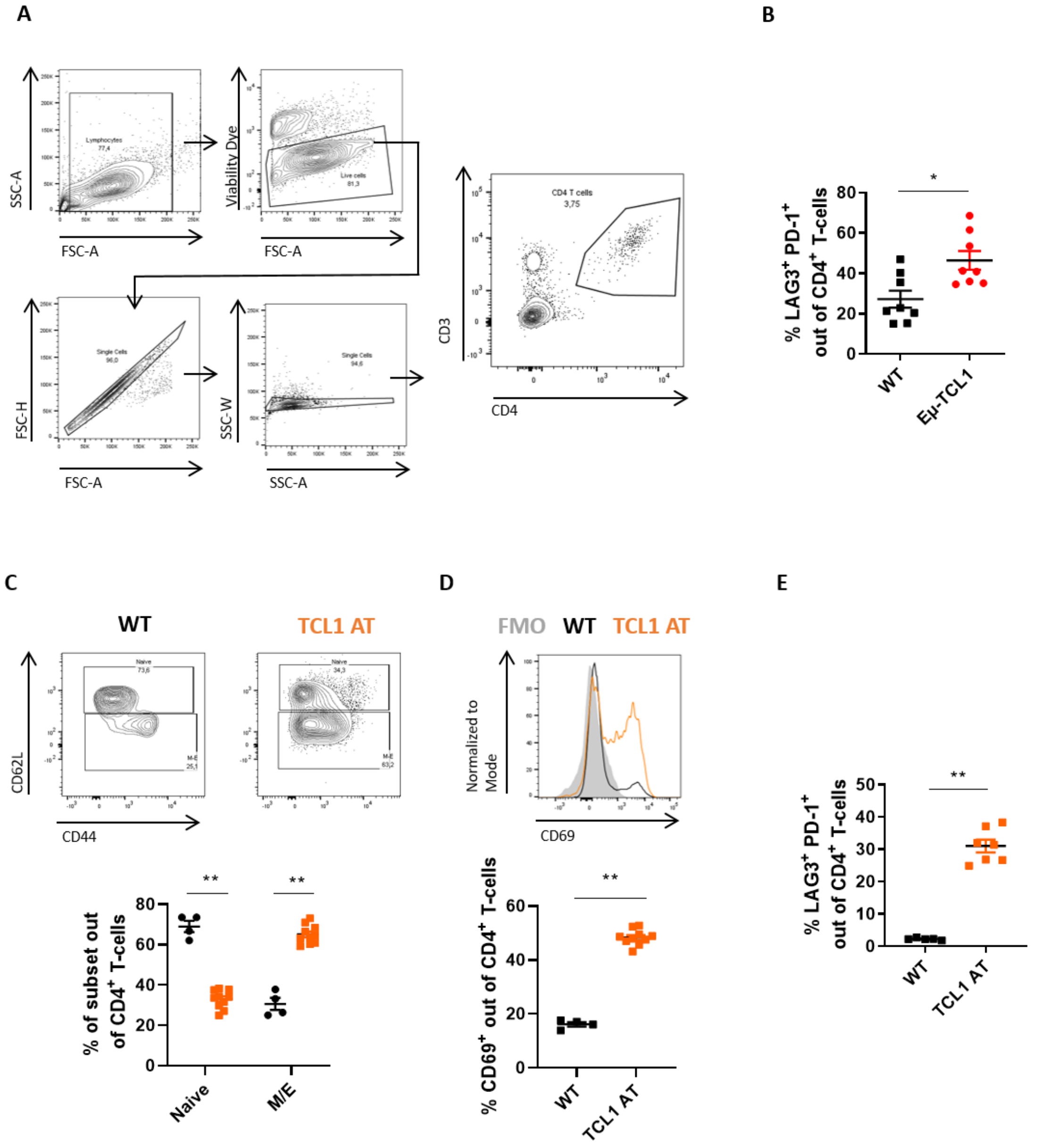


**Suppl. Figure 2: PD-1⁺ CD4⁺ T-cells in mouse models of CLL**

**A)** Representative gating strategy for flow cytometry analysis to define viable, single $CD4^+$ T-cells in murine splenocytes. **B)** Splenocytes of hemizygous Eμ-TCL1 leukemic mice and sex-matched WT littermates at a median age of 65 weeks were analyzed by flow cytometry. Percentage of $LAG3^+$ $PD\text{-}1^+$ co-expressing cells out of $CD4^+$ T-cells. **C-E)** Leukemic cells of Eμ-TCL1 mice were transplanted into syngeneic WT mice (TCL1 AT) and splenocytes were analyzed by flow cytometry, 4 weeks after transfer of cells. **C)** Frequency of $CD62L^{hi}$ $CD44^{low}$ naïve and $CD62L^{low}$ $CD44^{hi}$ antigen-experienced memory/effector $CD4^+$ T-cells. **D)** Percentage of CD69-expressing $CD4^+$ T-cells after TCL1 AT. **E)** Percentage of $LAG3^+$ $PD\text{-}1^+$ co-expressing cells out of $CD4^+$ T-cells.

All graphs show mean ± SEM. Each dot in B)-E) represents data of an individual mouse. Statistical analysis was performed using Mann-Whitney test. FMO = FMO = fluorescence-minus-one control.

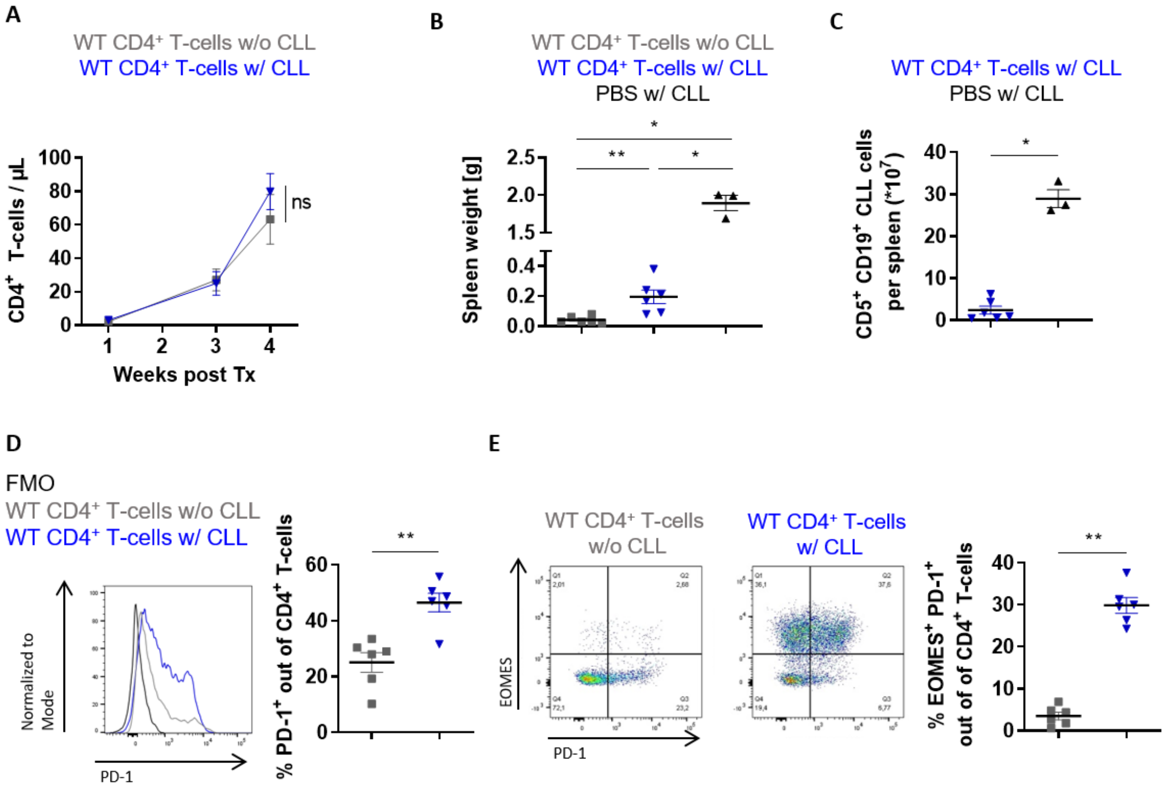


**Suppl. Figure 3:** **TCL1 leukemia induces PD-1 and EOMES co-expression of CD4+ T-cells**

*Rag2$^{-/-}$* mice were transplanted i.v. with CD4$^+$ T-cells on day -1 and the following day with (w/) or without (w/o) leukemic splenocytes of Eµ-TCL1 mice. **A)** Numbers of CD4$^+$ T-cells per µl blood over time. **B)** Spleen weight at endpoint, 4 weeks after transfer of leukemic cells. **C)** Number of CD5$^+$ CD19$^+$ CLL cells per spleen. **D)** Representative histogram and percentage of PD-1$^+$ out of CD4$^+$ T-cells. **E)** Representative graph and frequency of EOMES$^+$ PD-1$^+$ co-expressing CD4$^+$ T-cells.

All graphs show mean ± SEM. In B)-E), each dot represents data of an individual mouse. Statistical analysis was performed using Mann-Whitney test. Tx = transplantation.

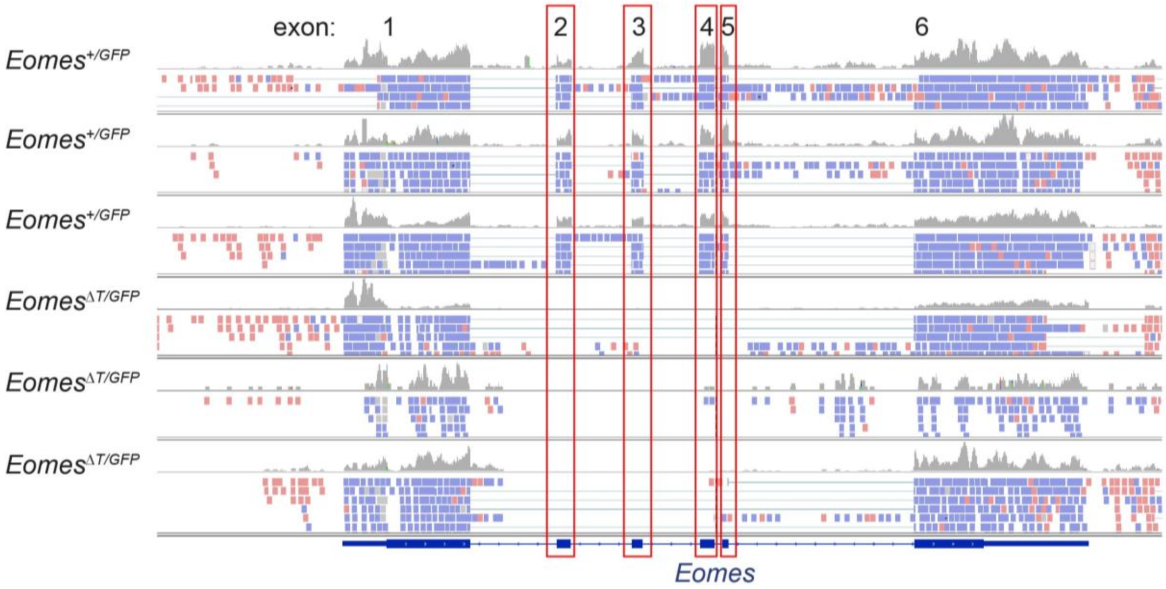


**Suppl. Figure 4:** **Efficient deletion of *Eomes* exons 2-5 in *Eomes*$^{\Delta T/GFP}$ CD4$^{+}$ T-cells**

Alignment of RNA sequencing reads to the *Eomes* gene locus. Top three samples retrieved GFP$^{+}$ CD4$^{+}$ T-cells cells of *Eomes*$^{+/GFP}$ reporter mice. Lower three samples are obtained of GFP$^{+}$ T-cells of *Eomes*$^{\Delta T/GFP}$ knock-out mice. Deletion of exons 2-5 in knock-out mice is highlighted in red.

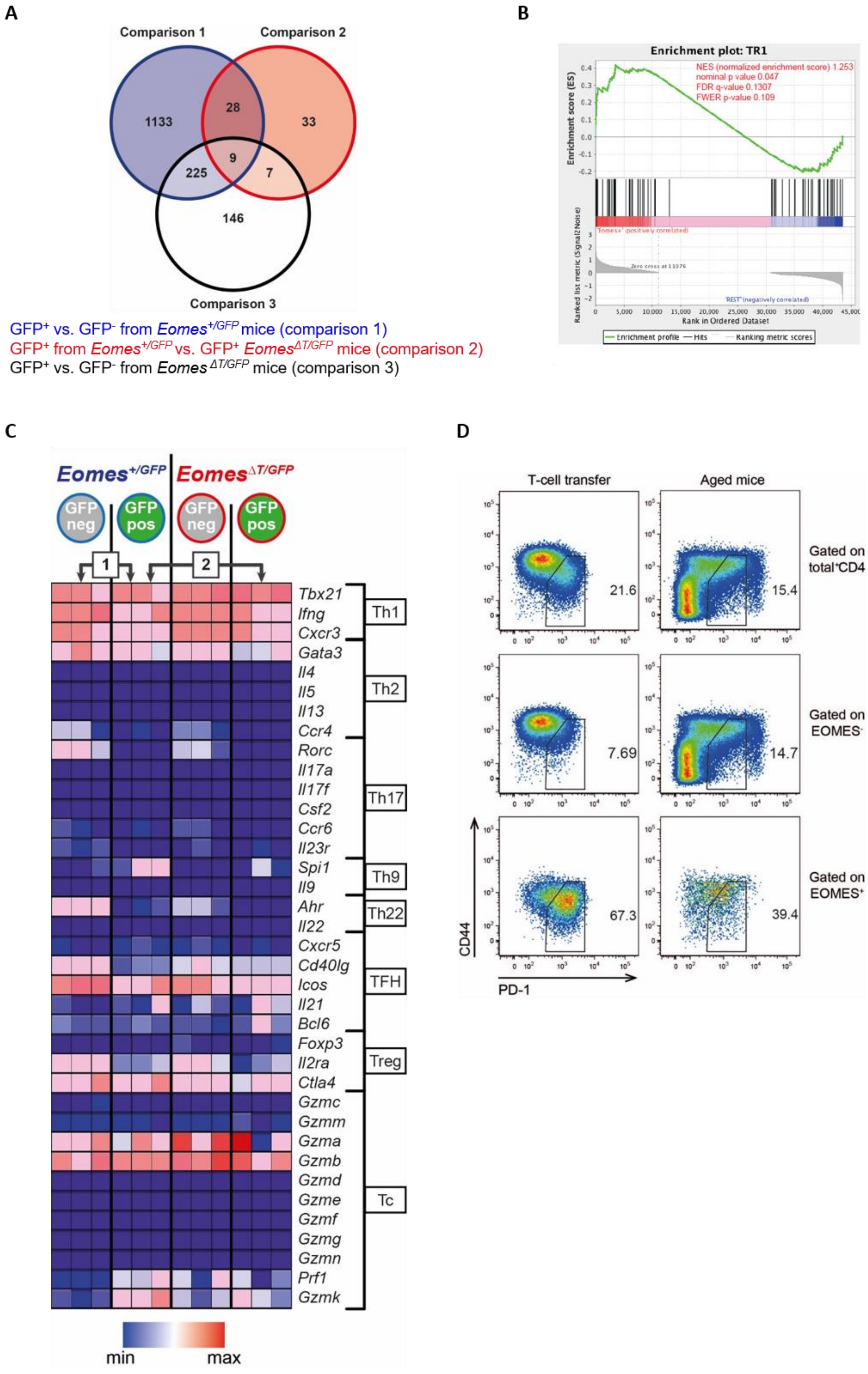
A
Comparison 1
Comparison 2
Comparison 3
1133
28
33
9
225
7
146
GFP+ vs. GFP- from Eomes+/GFP mice (comparison 1)
GFP+ from Eomes+/GFP vs. GFP+ EomesΔT/GFP mice (comparison 2)
GFP+ vs. GFP- from EomesΔT/GFP mice (comparison 3)
B
Enrichment plot: TR1
NES (normalized enrichment score) 1.253
nominal p value 0.047
FDR q-value 0.1307
FWER p-value 0.109
Enrichment score (ES)
Ranked list metric (Signal2Noise)
Rank in Ordered Dataset
Enrichment profile
Hits
Ranking metric scores
C
Eomes+/GFP
EomesΔT/GFP
GFP neg
GFP pos
GFP neg
GFP pos
1
2
Tbx21
Ifng
Cxcr3
Gata3
Il4
Il5
Il13
Ccr4
Rorc
Il17a
Il17f
Csf2
Ccr6
Il23r
Spi1
Il9
Ahr
Il22
Cxcr5
Cd40lg
Icos
Il21
Bcl6
Foxp3
Il2ra
Ctla4
Gzmc
Gzmm
Gzma
Gzmb
Gzmd
Gzme
Gzmf
Gzmg
Gzmn
Prf1
Gzmk
Th1
Th2
Th17
Th9
Th22
TFH
Treg
Tc
min
max
D
T-cell transfer
Aged mice
Gated on total+CD4
Gated on EOMES-
Gated on EOMES+
21.6
15.4
7.69
14.7
67.3
39.4
CD44
PD-1

**Suppl. Figure 5: Analysis of RNA-seq data of GFP$^{+}$ and GFP$^{-}$ CD4$^{+}$ T-cells from *Eomes*$^{+/GFP}$ or *Eomes*$^{\Delta T/GFP}$ mice**

**A)** Venn diagram showing overlap of differentially expressed genes revealed by RNA-sequencing as described in Figure 4A. **B)** Heatmap showing expression of genes characteristic for T helper (Th)1, Th2, Th17, Th9, Th22, TFH, Treg and Tc subsets. **C)** Gene set enrichment analysis (GSEA) of differentially expressed genes between *Eomes*$^{+}$ GFP$^{+}$ and *Eomes*$^{-}$ GFP$^{-}$ CD4$^{+}$ T-cells (comparison 1 in Figure 4A reveals an enrichment of previously described human $T_R$1 genes (15). **D)** Representative flow cytometric analysis of PD-1 and CD44 of CD4$^{+}$ T-cells after adoptive T-cell transfer in *Rag2*$^{-/-}$ mice (left) or in aged mice (right) gated either on total CD4$^{+}$ T-cells, EOMES$^{+}$ or EOMES$^{-}$ CD4$^{+}$ T-cells.

Gene expression in heatmap in C) was normalized to graph. TFH = follicular helper T-cell, Treg = regulatory T-cell, Tc = cytotoxic T-cell

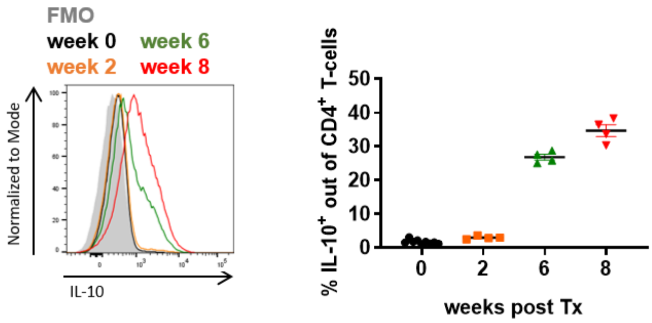


**Suppl. Figure 6:** **IL-10-producing CD4$^+$ T-cells accumulate over time in TCL1 AT mice**

Leukemic cells of Eµ-TCL1 mice were transplanted into syngeneic WT mice (TCL1 AT). Longitudinal analysis of frequency of IL-10-expressing cells out of CD4$^+$ T-cells in untransplanted (week 0) or leukemic TCL1 AT mice at indicated time points. Splenocytes of these mice were stimulated *ex vivo* with PMA/ionomycin and expression of IL-10 was analyzed by intracellular flow cytometry.

Each dot represents data of an individual mouse. FMO = fluorescence-minus-one control; Tx = transplantation.
