## Supplementary Table 4 for "EOMES and IL-10 regulate anti-tumor activity of PD-1^+^ CD4^+^ T-cells in B-cell Non-Hodgkin lymphoma"

| Comparison 1: EOMES$^{+}$ GFP$^{+}$ vs EOMES$^{-}$ GFP$^{-}$ of *Eomes* $^{+/GFP}$ reporter mice | | | | | | | |
|---|---|---|---|---|---|---|---|
| ENSEMBL ID | baseMean | log2FoldChange | lfcSE | stat | pvalue | padj | gene_names |
| ENSMUSG00000053965.6 | 347.5376204 | -4.518772307 | 0.281826982 | -16.03385267 | 7.41E-58 | 1.02E-53 | Pde5a |
| ENSMUSG00000026072.8 | 336.495343 | -3.471746784 | 0.27472138 | -12.63733744 | 1.31E-36 | 9.07E-33 | Il1r1 |
| ENSMUSG00000029810.11 | 329.728992 | -3.966843934 | 0.321857985 | -12.32482683 | 6.66E-35 | 3.06E-31 | Tmem176b |
| ENSMUSG00000033849.3 | 142.5193331 | -4.879593948 | 0.406063351 | -12.01682922 | 2.90E-33 | 1.00E-29 | B3galt2 |
| ENSMUSG00000023367.10 | 261.2449583 | -4.088683913 | 0.342589624 | -11.93464025 | 7.81E-33 | 2.16E-29 | Tmem176a |
| ENSMUSG00000053702.12 | 680.5658998 | -2.94038145 | 0.263569641 | -11.15599442 | 6.69E-29 | 1.54E-25 | Nebl |
| ENSMUSG00000043088.12 | 226.367437 | -4.235497251 | 0.412180847 | -10.27582257 | 9.06E-25 | 1.79E-21 | Il17re |
| ENSMUSG00000032446.10 | 548.5533969 | 3.805099655 | 0.376946626 | 10.09453166 | 5.84E-24 | 1.01E-20 | Eomes |
| ENSMUSG00000041272.7 | 727.6301705 | 3.79030092 | 0.387077992 | 9.792085826 | 1.22E-22 | 1.87E-19 | Tox |
| ENSMUSG00000001270.8 | 205.3658877 | -2.730611721 | 0.279744392 | -9.761095469 | 1.65E-22 | 2.28E-19 | Ckb |
| ENSMUSG00000020617.9 | 59.48228349 | -3.966988241 | 0.410257383 | -9.669510908 | 4.06E-22 | 5.10E-19 | 1700012B07Rik |
| ENSMUSG00000048521.7 | 2827.411361 | -2.555326196 | 0.264768072 | -9.651187078 | 4.86E-22 | 5.59E-19 | Cxcr6 |
| ENSMUSG00000023927.11 | 3530.075137 | -1.876170455 | 0.203438297 | -9.222307109 | 2.91E-20 | 3.09E-17 | Satb1 |
| ENSMUSG00000030281.12 | 62.3716093 | -4.042750778 | 0.443470197 | -9.116172409 | 7.78E-20 | 7.68E-17 | Il17rc |
| ENSMUSG00000052821.3 | 55.02848201 | -3.603498358 | 0.419578585 | -8.588375296 | 8.82E-18 | 8.12E-15 | Cysltr1 |
| ENSMUSG00000024646.9 | 451.2730295 | -1.675939969 | 0.196900021 | -8.511629189 | 1.72E-17 | 1.48E-14 | Cyb5a |
| ENSMUSG00000005802.8 | 278.1955796 | -2.76723975 | 0.327873768 | -8.439954698 | 3.17E-17 | 2.58E-14 | Slc30a4 |
| ENSMUSG00000033066.11 | 546.7513853 | 1.645414623 | 0.195713565 | 8.407258976 | 4.20E-17 | 3.22E-14 | Gas7 |
| ENSMUSG00000003882.4 | 2965.983947 | -1.832253561 | 0.219192952 | -8.359089754 | 6.32E-17 | 4.59E-14 | Il7r |
| ENSMUSG00000103779.1 | 441.400015 | -1.931859566 | 0.238535784 | -8.098824973 | 5.55E-16 | 3.83E-13 | Gm36931 |
| ENSMUSG00000078942.6 | 140.0653708 | 2.239338337 | 0.277858842 | 8.059266055 | 7.68E-16 | 5.05E-13 | Naip6 |
| ENSMUSG00000102973.1 | 220.6370693 | -2.14224063 | 0.266083041 | -8.051022795 | 8.21E-16 | 5.15E-13 | E430014B02Rik |
| ENSMUSG00000015133.12 | 573.8024898 | 2.226002211 | 0.282035199 | 7.892639701 | 2.96E-15 | 1.70E-12 | Lrrk1 |
| ENSMUSG00000020183.7 | 920.442726 | -2.131303995 | 0.269936239 | -7.89558305 | 2.89E-15 | 1.70E-12 | Cpm |
| ENSMUSG00000076472.2 | 154.146867 | -3.034778818 | 0.385065691 | -7.881197649 | 3.24E-15 | 1.79E-12 | Trbv15 |
| ENSMUSG00000030283.5 | 265.6191703 | -2.782597905 | 0.354392918 | -7.851731122 | 4.10E-15 | 2.10E-12 | St8sia1 |
| ENSMUSG00000040229.7 | 46.93313147 | -3.396101753 | 0.432450129 | -7.853163922 | 4.06E-15 | 2.10E-12 | Gpr34 |
| ENSMUSG00000026770.5 | 444.9558994 | -1.702031773 | 0.217040678 | -7.841994351 | 4.43E-15 | 2.11E-12 | Il2ra |
| ENSMUSG00000042385.10 | 440.3802572 | 2.552625405 | 0.32536075 | 7.845523478 | 4.31E-15 | 2.11E-12 | Gzmk |
| ENSMUSG00000028246.9 | 63.68034071 | 2.836323499 | 0.362513469 | 7.824049975 | 5.12E-15 | 2.35E-12 | Faxc |
| ENSMUSG00000028150.10 | 178.1360844 | -3.300343391 | 0.422243453 | -7.816209752 | 5.44E-15 | 2.42E-12 | Rorc |
| ENSMUSG00000043807.6 | 87.565277 | -2.803686993 | 0.359760336 | -7.793207624 | 6.53E-15 | 2.82E-12 | Ly6g5b |
| ENSMUSG00000024247.10 | 63.99761885 | 2.444236009 | 0.322387368 | 7.581674258 | 3.41E-14 | 1.43E-11 | Pkdcc |
| ENSMUSG00000086968.4 | 55.37796287 | -3.010496557 | 0.398344178 | -7.557526182 | 4.11E-14 | 1.67E-11 | 4933431E20Rik |
| ENSMUSG00000051354.9 | 51.75437764 | 2.98593293 | 0.398417332 | 7.494485538 | 6.66E-14 | 2.63E-11 | Samd3 |
| ENSMUSG00000038679.12 | 809.4081776 | 1.704573875 | 0.230941326 | 7.380982454 | 1.57E-13 | 6.03E-11 | Trps1 |
| ENSMUSG00000023830.9 | 2165.337168 | 1.110236969 | 0.151407035 | 7.332796453 | 2.25E-13 | 8.41E-11 | Igf2r |
| ENSMUSG00000032089.12 | 1197.722813 | 2.137466396 | 0.293678233 | 7.278259516 | 3.38E-13 | 1.23E-10 | Il10ra |
| ENSMUSG00000049103.9 | 3924.696823 | -2.326574879 | 0.319843131 | -7.274112382 | 3.49E-13 | 1.23E-10 | Ccr2 |
| ENSMUSG00000102212.1 | 421.0598169 | -1.858157136 | 0.255671764 | -7.267744808 | 3.66E-13 | 1.26E-10 | C230085N15Rik |
| ENSMUSG00000030336.10 | 331.6836066 | 2.585461671 | 0.35979078 | 7.186014238 | 6.67E-13 | 2.25E-10 | Cd27 |
| ENSMUSG00000025997.9 | 818.75674 | 2.402502984 | 0.335343205 | 7.16431092 | 7.82E-13 | 2.51E-10 | Ikzf2 |
| ENSMUSG00000029408.9 | 223.8798525 | 2.075185814 | 0.289572077 | 7.166387854 | 7.70E-13 | 2.51E-10 | Abcb9 |
| ENSMUSG00000102744.1 | 139.7924404 | -2.182697658 | 0.3049406 | -7.157779759 | 8.20E-13 | 2.57E-10 | 5830444F18Rik |
| ENSMUSG00000071203.6 | 78.97809544 | 2.178141687 | 0.30761286 | 7.080788777 | 1.43E-12 | 4.40E-10 | Naip5 |
| ENSMUSG00000103560.1 | 47.89763704 | -3.098894921 | 0.443107949 | -6.993543969 | 2.68E-12 | 8.05E-10 | Gm38070 |
| ENSMUSG00000015222.13 | 39.0752359 | 2.838341435 | 0.408263833 | 6.952223553 | 3.60E-12 | 1.06E-09 | Map2 |
| ENSMUSG00000039943.12 | 227.0110544 | -2.33244525 | 0.337962595 | -6.901489351 | 5.15E-12 | 1.48E-09 | Plcb4 |
| ENSMUSG00000035735.6 | 21.08600328 | 3.351764823 | 0.486123127 | 6.894888638 | 5.39E-12 | 1.52E-09 | Dagla |
| ENSMUSG00000015950.9 | 809.5909473 | 2.079940588 | 0.304104992 | 6.839547666 | 7.94E-12 | 2.19E-09 | Ncf1 |
| ENSMUSG00000026830.9 | 245.42401 | -2.086575272 | 0.305240635 | -6.835837155 | 8.15E-12 | 2.21E-09 | Ermn |
| ENSMUSG00000020644.8 | 3152.724258 | -1.613456093 | 0.236160761 | -6.832024452 | 8.37E-12 | 2.22E-09 | Id2 |
| ENSMUSG00000062044.9 | 45.52070707 | 2.550183074 | 0.374056097 | 6.817648732 | 9.25E-12 | 2.41E-09 | Lmtk3 |
| ENSMUSG00000044340.7 | 198.3359436 | 1.798815342 | 0.26471432 | 6.795308029 | 1.08E-11 | 2.76E-09 | Phlpp1 |
| ENSMUSG00000025491.10 | 756.5996143 | -2.9667851 | 0.436898881 | -6.790553217 | 1.12E-11 | 2.80E-09 | Ifitm1 |
| ENSMUSG00000052374.10 | 149.6203317 | -2.68483159 | 0.395594618 | -6.786825372 | 1.15E-11 | 2.83E-09 | Actn2 |
| ENSMUSG00000022657.9 | 1101.214728 | -1.50229203 | 0.221987003 | -6.767477432 | 1.31E-11 | 3.17E-09 | Cd96 |
| ENSMUSG00000020593.10 | 860.1416659 | 1.343555224 | 0.200349031 | 6.706072995 | 2.00E-11 | 4.76E-09 | Lpin1 |
| ENSMUSG00000103216.1 | 306.4032458 | -1.758667012 | 0.263475862 | -6.674869562 | 2.47E-11 | 5.79E-09 | Gm37248 |
| ENSMUSG00000045573.9 | 59.80202469 | 2.638978268 | 0.39662872 | 6.653522891 | 2.86E-11 | 6.59E-09 | Penk |

| | | | | | | | |
|---|---|---|---|---|---|---|---|
| ENSMUSG00000040249.11 | 380.5385895 | 2.499234106 | 0.376987319 | 6.629491184 | 3.37E-11 | 7.63E-09 | Lrp1 |
| ENSMUSG00000009588.9 | 527.1678862 | -2.004436084 | 0.303490577 | -6.604607305 | 3.99E-11 | 8.88E-09 | St6galnac1 |
| ENSMUSG00000000682.7 | 3209.662915 | -1.504496412 | 0.229061226 | -6.568097257 | 5.10E-11 | 1.12E-08 | Cd52 |
| ENSMUSG00000026070.11 | 5366.576568 | -1.349091275 | 0.2054698 | -6.565885973 | 5.17E-11 | 1.12E-08 | Il18r1 |
| ENSMUSG00000031933.13 | 73.43957176 | 2.17056149 | 0.331761498 | 6.542535843 | 6.05E-11 | 1.28E-08 | Izumo1r |
| ENSMUSG00000024451.8 | 106.1885205 | 2.681512288 | 0.41183682 | 6.511103807 | 7.46E-11 | 1.56E-08 | Arap3 |
| ENSMUSG00000027073.5 | 28.734156 | 3.188574833 | 0.494270207 | 6.45107633 | 1.11E-10 | 2.29E-08 | Prg2 |
| ENSMUSG00000031785.11 | 50.00380427 | 2.638185417 | 0.409412733 | 6.443828454 | 1.16E-10 | 2.37E-08 | Adgrg1 |
| ENSMUSG00000033213.12 | 227.1579367 | -2.609570132 | 0.406168245 | -6.424850204 | 1.32E-10 | 2.64E-08 | AA467197 |
| ENSMUSG00000032020.11 | 445.1276983 | 1.860030512 | 0.2899508 | 6.414986651 | 1.41E-10 | 2.78E-08 | Ubash3b |
| ENSMUSG00000061175.7 | 104.9053036 | 1.737970163 | 0.272831884 | 6.370113848 | 1.89E-10 | 3.67E-08 | Fnip2 |
| ENSMUSG00000100815.1 | 60.49736911 | 2.661249313 | 0.418689379 | 6.356142404 | 2.07E-10 | 3.97E-08 | Gm29112 |
| ENSMUSG00000029869.7 | 196.170893 | 1.952312119 | 0.307464991 | 6.349705421 | 2.16E-10 | 4.08E-08 | Ephb6 |
| ENSMUSG00000034117.3 | 32.82839961 | -3.134459394 | 0.494324985 | -6.34088806 | 2.28E-10 | 4.26E-08 | Ptgdr2 |
| ENSMUSG00000015355.9 | 1954.194329 | -1.48356936 | 0.234694341 | -6.321283053 | 2.59E-10 | 4.78E-08 | Cd48 |
| ENSMUSG00000015968.12 | 42.61359652 | 3.161140762 | 0.503042014 | 6.284049193 | 3.30E-10 | 5.99E-08 | Cacna1d |
| ENSMUSG00000070407.5 | 430.1918269 | -2.145311722 | 0.342171334 | -6.269700319 | 3.62E-10 | 6.49E-08 | Hs3st3b1 |
| ENSMUSG00000015709.8 | 133.8170262 | -1.708286232 | 0.274201648 | -6.230036338 | 4.66E-10 | 8.26E-08 | Arnt2 |
| ENSMUSG00000034220.7 | 689.7194678 | -1.492472482 | 0.241021407 | -6.192281852 | 5.93E-10 | 1.04E-07 | Gpc1 |
| ENSMUSG00000028435.8 | 74.82520558 | -2.40586497 | 0.38955058 | -6.176001503 | 6.57E-10 | 1.13E-07 | Aqp3 |
| ENSMUSG00000049410.8 | 62.07359261 | -2.392861254 | 0.388234495 | -6.163443188 | 7.12E-10 | 1.21E-07 | Zfp683 |
| ENSMUSG00000031132.1 | 317.078733 | -1.555918163 | 0.253353524 | -6.14129275 | 8.19E-10 | 1.38E-07 | Cd40lg |
| ENSMUSG00000029026.12 | 82.36065508 | 2.158473779 | 0.352799325 | 6.118134661 | 9.47E-10 | 1.58E-07 | Trp73 |
| ENSMUSG00000035725.9 | 827.743543 | -1.224833163 | 0.201962898 | -6.064644418 | 1.32E-09 | 2.17E-07 | Prkx |
| ENSMUSG00000002190.9 | 77.13209784 | 2.122008459 | 0.350843326 | 6.048307898 | 1.46E-09 | 2.35E-07 | Clgn |
| ENSMUSG00000019256.13 | 216.5532654 | -1.918198404 | 0.317307175 | -6.045241182 | 1.49E-09 | 2.35E-07 | Ahr |
| ENSMUSG00000026285.7 | 518.6135975 | 1.831999825 | 0.302921466 | 6.047771548 | 1.47E-09 | 2.35E-07 | Pdcd1 |
| ENSMUSG00000074570.9 | 151.6740749 | -1.74457701 | 0.288632441 | -6.044285965 | 1.50E-09 | 2.35E-07 | Cass4 |
| ENSMUSG00000020614.9 | 72.0643366 | -2.026584676 | 0.338391484 | -5.988876109 | 2.11E-09 | 3.28E-07 | Fam20a |
| ENSMUSG00000020865.12 | 46.2698475 | -2.227208218 | 0.372119626 | -5.985194173 | 2.16E-09 | 3.32E-07 | Abcc3 |
| ENSMUSG00000005947.7 | 47.67519566 | -2.1729357 | 0.364344657 | -5.963956542 | 2.46E-09 | 3.71E-07 | Itgae |
| ENSMUSG00000059901.8 | 42.43737372 | 2.770253141 | 0.464560439 | 5.963170582 | 2.47E-09 | 3.71E-07 | Adamts14 |
| ENSMUSG00000001663.6 | 64.74533338 | -2.116393165 | 0.356649213 | -5.934103008 | 2.95E-09 | 4.34E-07 | Gstt1 |
| ENSMUSG00000030167.11 | 386.0743588 | -2.201494822 | 0.370938284 | -5.934935578 | 2.94E-09 | 4.34E-07 | Klrc1 |
| ENSMUSG00000032690.12 | 134.4388152 | 2.029341969 | 0.343896097 | 5.901032284 | 3.61E-09 | 5.25E-07 | Oas2 |
| ENSMUSG00000017754.9 | 114.8034791 | -1.498677591 | 0.254315358 | -5.892988931 | 3.79E-09 | 5.46E-07 | Pltp |
| ENSMUSG00000030208.11 | 603.6245533 | -1.985093277 | 0.337134446 | -5.888135424 | 3.91E-09 | 5.56E-07 | Emp1 |
| ENSMUSG00000064147.6 | 143.3057364 | 2.465755761 | 0.419233124 | 5.881586215 | 4.06E-09 | 5.73E-07 | Rab44 |
| ENSMUSG00000046207.10 | 76.28938678 | 2.434681137 | 0.414819747 | 5.869250811 | 4.38E-09 | 6.11E-07 | Pik3r6 |
| ENSMUSG00000033174.13 | 106.7474609 | -1.984519578 | 0.33824362 | -5.86713085 | 4.43E-09 | 6.12E-07 | Mgll |
| ENSMUSG00000032336.13 | 1623.496701 | -1.236368805 | 0.21094633 | -5.861058623 | 4.60E-09 | 6.29E-07 | Nptn |
| ENSMUSG00000024812.9 | 287.0256541 | 1.296673415 | 0.221486743 | 5.854406445 | 4.79E-09 | 6.48E-07 | Tjp2 |
| ENSMUSG00000038872.9 | 67.66252164 | 2.274381274 | 0.388691129 | 5.851384569 | 4.87E-09 | 6.54E-07 | Zfhx3 |
| ENSMUSG00000097796.1 | 62.50121363 | 2.17601813 | 0.372892152 | 5.835516024 | 5.36E-09 | 7.12E-07 | Gm16702 |
| ENSMUSG00000051212.7 | 488.5450529 | -1.577389318 | 0.271766111 | -5.804216404 | 6.47E-09 | 8.50E-07 | Gpr183 |
| ENSMUSG00000074604.5 | 110.8387294 | 1.651090854 | 0.285381796 | 5.785550716 | 7.23E-09 | 9.42E-07 | Mgst2 |
| ENSMUSG00000082399.1 | 97.66639561 | -1.705480681 | 0.295414282 | -5.773182898 | 7.78E-09 | 1.00E-06 | Gm14036 |
| ENSMUSG00000000204.11 | 1065.652998 | 2.464844257 | 0.427187383 | 5.769936934 | 7.93E-09 | 1.01E-06 | Slfn4 |
| ENSMUSG00000020027.14 | 195.4831702 | -1.791074805 | 0.312525597 | -5.730969945 | 9.99E-09 | 1.27E-06 | Socs2 |
| ENSMUSG00000034730.12 | 14.76846412 | 2.830746153 | 0.4961053 | 5.705938142 | 1.16E-08 | 1.45E-06 | Adgrb1 |
| ENSMUSG00000035891.12 | 204.4610975 | -1.269813584 | 0.222595789 | -5.70457146 | 1.17E-08 | 1.45E-06 | Cerk |
| ENSMUSG00000021728.7 | 2260.260871 | -1.363464835 | 0.239258321 | -5.698714379 | 1.21E-08 | 1.49E-06 | Emb |
| ENSMUSG00000040613.10 | 165.9119935 | 1.592327262 | 0.279827805 | 5.690382555 | 1.27E-08 | 1.55E-06 | Apobec1 |
| ENSMUSG00000051339.8 | 81.40216183 | 2.449219481 | 0.431010558 | 5.682504607 | 1.33E-08 | 1.61E-06 | 2900026A02Rik |
| ENSMUSG00000026475.7 | 160.7130708 | 2.367377708 | 0.416966416 | 5.677622024 | 1.37E-08 | 1.64E-06 | Rgs16 |
| ENSMUSG00000026193.11 | 407.962143 | 2.359518065 | 0.416167609 | 5.66963409 | 1.43E-08 | 1.70E-06 | Fn1 |
| ENSMUSG00000061577.7 | 226.5612229 | -1.992113139 | 0.35142395 | -5.668689175 | 1.44E-08 | 1.70E-06 | Adgrg5 |
| ENSMUSG00000034591.5 | 71.98614068 | -2.011757451 | 0.355641137 | -5.656706283 | 1.54E-08 | 1.81E-06 | Slc41a2 |
| ENSMUSG00000034271.11 | 105.8204387 | 1.610742584 | 0.285664827 | 5.63857512 | 1.71E-08 | 1.99E-06 | Jdp2 |
| ENSMUSG00000032021.9 | 82.07758611 | 2.243435866 | 0.398652988 | 5.627540581 | 1.83E-08 | 2.10E-06 | Crtam |
| ENSMUSG00000052736.11 | 197.0528462 | -2.001181369 | 0.355979465 | -5.621620252 | 1.89E-08 | 2.14E-06 | Klrc2 |
| ENSMUSG00000103546.1 | 21.98703848 | 2.527306075 | 0.449526559 | 5.622150741 | 1.89E-08 | 2.14E-06 | Gm37666 |

| | | | | | | | |
|---|---|---|---|---|---|---|---|
| ENSMUSG00000045094.7 | 34.23909833 | 2.482101425 | 0.442392637 | 5.610630054 | 2.02E-08 | 2.26E-06 | Arhgef37 |
| ENSMUSG00000053716.9 | 318.4114094 | -1.230897524 | 0.219425124 | -5.609647149 | 2.03E-08 | 2.26E-06 | Dusp7 |
| ENSMUSG00000028859.10 | 352.9288115 | 2.354122304 | 0.420096002 | 5.603772211 | 2.10E-08 | 2.32E-06 | Csf3r |
| ENSMUSG00000021281.11 | 274.3286534 | 2.136712095 | 0.381720967 | 5.597575923 | 2.17E-08 | 2.38E-06 | Tnfaip2 |
| ENSMUSG00000030653.12 | 522.3813259 | 1.277258155 | 0.228216271 | 5.596700663 | 2.18E-08 | 2.38E-06 | Pde2a |
| ENSMUSG00000060012.7 | 1218.619431 | 1.584689293 | 0.284147329 | 5.57699873 | 2.45E-08 | 2.64E-06 | Kif13b |
| ENSMUSG00000037202.5 | 212.4957254 | 1.876364368 | 0.336987319 | 5.568056313 | 2.58E-08 | 2.76E-06 | Prf1 |
| ENSMUSG00000062995.8 | 81.5872782 | 1.925245889 | 0.345946723 | 5.565151401 | 2.62E-08 | 2.78E-06 | Ica1 |
| ENSMUSG00000026019.11 | 362.3443753 | -1.354043712 | 0.243611348 | -5.55821279 | 2.73E-08 | 2.87E-06 | Wdr12 |
| ENSMUSG00000022013.3 | 426.9167359 | -1.795819843 | 0.323792926 | -5.54619851 | 2.92E-08 | 3.05E-06 | Dnajc15 |
| ENSMUSG00000040957.10 | 39.22869415 | 2.385975258 | 0.430911007 | 5.537048759 | 3.08E-08 | 3.19E-06 | Cables1 |
| ENSMUSG00000048120.12 | 546.9639724 | 1.296902416 | 0.235238195 | 5.513145581 | 3.52E-08 | 3.63E-06 | Entpd1 |
| ENSMUSG00000079056.8 | 62.78302438 | 1.961043191 | 0.356063491 | 5.507566035 | 3.64E-08 | 3.72E-06 | Kcnip3 |
| ENSMUSG00000041268.12 | 196.3048718 | 2.315148362 | 0.421223642 | 5.496245058 | 3.88E-08 | 3.94E-06 | Dmxl2 |
| ENSMUSG00000040183.9 | 84.13397046 | -1.872494372 | 0.34092446 | -5.492402546 | 3.97E-08 | 4.00E-06 | Ankrd6 |
| ENSMUSG00000009687.10 | 2286.536414 | -1.044358925 | 0.190544836 | -5.480909099 | 4.23E-08 | 4.21E-06 | Fxyd5 |
| ENSMUSG00000025017.9 | 290.4983012 | 1.276066686 | 0.232830666 | 5.480664144 | 4.24E-08 | 4.21E-06 | Pik3ap1 |
| ENSMUSG00000041912.8 | 43.63631295 | -2.751532123 | 0.502907722 | -5.471246522 | 4.47E-08 | 4.41E-06 | Tdrkh |
| ENSMUSG00000019302.12 | 83.9265116 | 2.261436992 | 0.414677044 | 5.453489706 | 4.94E-08 | 4.84E-06 | Atp6v0a1 |
| ENSMUSG00000035711.4 | 84.97767399 | 2.237535746 | 0.410427372 | 5.451721542 | 4.99E-08 | 4.85E-06 | Dok3 |
| ENSMUSG00000083950.1 | 29.77347946 | 2.671189321 | 0.490843251 | 5.442041455 | 5.27E-08 | 5.09E-06 | Gm14466 |
| ENSMUSG00000043252.8 | 275.8488783 | -1.541762572 | 0.284667083 | -5.416019844 | 6.09E-08 | 5.84E-06 | Tmem64 |
| ENSMUSG00000018398.14 | 230.5447884 | -1.356628813 | 0.251625324 | -5.391463744 | 6.99E-08 | 6.66E-06 | Sept8 |
| ENSMUSG00000032656.10 | 138.7607279 | -1.546160909 | 0.287238609 | -5.382844988 | 7.33E-08 | 6.93E-06 | March3 |
| ENSMUSG00000028460.6 | 372.4840594 | -1.292085143 | 0.240110562 | -5.381209099 | 7.40E-08 | 6.95E-06 | Sit1 |
| ENSMUSG00000059326.6 | 201.6398504 | 2.266784071 | 0.423218997 | 5.356054634 | 8.51E-08 | 7.94E-06 | Csf2ra |
| ENSMUSG00000001020.7 | 4466.748751 | -1.533601578 | 0.287817579 | -5.328380506 | 9.91E-08 | 9.18E-06 | S100a4 |
| ENSMUSG00000031012.13 | 74.36926284 | 2.022964877 | 0.380297606 | 5.319425741 | 1.04E-07 | 9.58E-06 | Cask |
| ENSMUSG00000021360.11 | 321.116458 | -1.874724497 | 0.353051694 | -5.310056661 | 1.10E-07 | 1.00E-05 | Gcnt2 |
| ENSMUSG00000043004.9 | 1328.685799 | -1.014052221 | 0.191014868 | -5.308760673 | 1.10E-07 | 1.00E-05 | Gng2 |
| ENSMUSG00000030844.7 | 163.6042584 | -1.61477595 | 0.306946431 | -5.260774473 | 1.43E-07 | 1.29E-05 | Rgs10 |
| ENSMUSG00000070056.5 | 463.5896982 | -1.064693493 | 0.202730885 | -5.251757719 | 1.51E-07 | 1.35E-05 | Mfhas1 |
| ENSMUSG00000022377.12 | 1161.817367 | 1.332859843 | 0.254055855 | 5.246326026 | 1.55E-07 | 1.38E-05 | Asap1 |
| ENSMUSG00000004446.8 | 146.6758589 | 1.449549607 | 0.276586766 | 5.240849481 | 1.60E-07 | 1.41E-05 | Bid |
| ENSMUSG00000076745.1 | 12.16117575 | -2.724246628 | 0.52047246 | -5.234180171 | 1.66E-07 | 1.46E-05 | Tcrg-V4 |
| ENSMUSG00000035283.4 | 20.32866468 | 2.560602917 | 0.489758457 | 5.228297501 | 1.71E-07 | 1.50E-05 | Adrb1 |
| ENSMUSG00000030124.2 | 338.7920668 | 2.032357784 | 0.389104844 | 5.223162383 | 1.76E-07 | 1.53E-05 | Lag3 |
| ENSMUSG00000026919.1 | 51.51270423 | -1.943887456 | 0.372372184 | -5.220281047 | 1.79E-07 | 1.54E-05 | Lcn4 |
| ENSMUSG00000022106.10 | 723.9812976 | -0.943537019 | 0.18088688 | -5.216171667 | 1.83E-07 | 1.56E-05 | Rcbtb2 |
| ENSMUSG00000099954.1 | 89.25189826 | 2.487541922 | 0.476974747 | 5.215248688 | 1.84E-07 | 1.56E-05 | Gm28112 |
| ENSMUSG00000061533.11 | 387.9657736 | 1.211108634 | 0.232344224 | 5.212561825 | 1.86E-07 | 1.58E-05 | Cep128 |
| ENSMUSG00000056394.13 | 707.6727552 | 0.989246217 | 0.189918677 | 5.208788469 | 1.90E-07 | 1.60E-05 | Lig1 |
| ENSMUSG00000036503.9 | 333.0303783 | -1.420630582 | 0.272981173 | -5.20413392 | 1.95E-07 | 1.63E-05 | Rnf13 |
| ENSMUSG00000056413.12 | 656.7456797 | 1.032312723 | 0.198768222 | 5.193550106 | 2.06E-07 | 1.72E-05 | Adap1 |
| ENSMUSG00000025531.10 | 496.6766216 | -1.083430216 | 0.208681 | -5.191800958 | 2.08E-07 | 1.72E-05 | Chm |
| ENSMUSG00000009292.13 | 149.7169236 | 2.12475774 | 0.409674461 | 5.186453984 | 2.14E-07 | 1.76E-05 | Trpm2 |
| ENSMUSG00000056091.8 | 75.22225734 | 2.172354178 | 0.418961943 | 5.18508713 | 2.16E-07 | 1.76E-05 | St3gal5 |
| ENSMUSG00000060591.8 | 517.775375 | -2.234850793 | 0.431385597 | -5.180633767 | 2.21E-07 | 1.80E-05 | Ifitm2 |
| ENSMUSG00000026826.9 | 292.7453581 | 2.318260688 | 0.447712673 | 5.178009976 | 2.24E-07 | 1.80E-05 | Nr4a2 |
| ENSMUSG00000056501.3 | 214.6846962 | 1.506524424 | 0.290952434 | 5.177906245 | 2.24E-07 | 1.80E-05 | Cebpb |
| ENSMUSG00000043940.10 | 144.3182802 | 2.105509444 | 0.407698408 | 5.164379851 | 2.41E-07 | 1.93E-05 | Wdfy3 |
| ENSMUSG00000025507.9 | 187.7767081 | 1.148058234 | 0.222773449 | 5.153478735 | 2.56E-07 | 2.03E-05 | Pidd1 |
| ENSMUSG00000027035.6 | 502.1487743 | 1.518931359 | 0.29480807 | 5.152271986 | 2.57E-07 | 2.03E-05 | Cers6 |
| ENSMUSG00000029516.15 | 1035.057666 | 1.301211705 | 0.253001423 | 5.143100345 | 2.70E-07 | 2.12E-05 | Cit |
| ENSMUSG00000102594.1 | 19.02630242 | 2.487266304 | 0.483888081 | 5.140168564 | 2.74E-07 | 2.14E-05 | Gm38381 |
| ENSMUSG00000025330.6 | 106.5892755 | 2.179204627 | 0.424057771 | 5.138933357 | 2.76E-07 | 2.14E-05 | Padi4 |
| ENSMUSG00000021253.6 | 62.80830922 | -1.775172381 | 0.345656564 | -5.135653605 | 2.81E-07 | 2.17E-05 | Tgfb3 |
| ENSMUSG00000049093.9 | 60.84547467 | -2.061418944 | 0.401602841 | -5.132978991 | 2.85E-07 | 2.19E-05 | Il23r |
| ENSMUSG00000047821.12 | 250.0078801 | -1.406363449 | 0.274371469 | -5.125764193 | 2.96E-07 | 2.26E-05 | Trim16 |
| ENSMUSG00000063450.10 | 811.5670204 | 1.080904805 | 0.211211932 | 5.117631351 | 3.09E-07 | 2.35E-05 | Syne2 |
| ENSMUSG00000028977.12 | 78.7735388 | 2.051646293 | 0.401347812 | 5.11189106 | 3.19E-07 | 2.41E-05 | Casz1 |
| ENSMUSG00000021457.10 | 338.3621816 | 2.143533076 | 0.420016451 | 5.10345028 | 3.34E-07 | 2.50E-05 | Syk |

| | | | | | | | |
|---|---|---|---|---|---|---|---|
| ENSMUSG00000038473.10 | 28.6550399 | 2.372463682 | 0.465014456 | 5.101913827 | 3.36E-07 | 2.51E-05 | Nos1ap |
| ENSMUSG00000028078.10 | 224.8033429 | 1.477352935 | 0.289917005 | 5.09577883 | 3.47E-07 | 2.56E-05 | Dclk2 |
| ENSMUSG00000030589.11 | 181.7676974 | 2.124230018 | 0.41681308 | 5.09636122 | 3.46E-07 | 2.56E-05 | Rasgrp4 |
| ENSMUSG00000103233.1 | 63.03655878 | 1.956112066 | 0.383943238 | 5.094794935 | 3.49E-07 | 2.56E-05 | Gm37159 |
| ENSMUSG00000039852.12 | 497.726598 | 1.413070002 | 0.277557562 | 5.091088102 | 3.56E-07 | 2.60E-05 | Rere |
| ENSMUSG00000078247.3 | 84.74185725 | 2.135057137 | 0.419416893 | 5.090536821 | 3.57E-07 | 2.60E-05 | Airn |
| ENSMUSG00000006342.10 | 386.2420601 | -1.726081602 | 0.339315175 | -5.086956702 | 3.64E-07 | 2.63E-05 | Susd2 |
| ENSMUSG00000029561.13 | 171.1645224 | 1.800684575 | 0.354361352 | 5.081492557 | 3.74E-07 | 2.67E-05 | Oasl2 |
| ENSMUSG00000033720.8 | 35.10737117 | 1.907616649 | 0.37541825 | 5.081310381 | 3.75E-07 | 2.67E-05 | Sfxn5 |
| ENSMUSG00000052776.10 | 89.71340621 | 1.386515948 | 0.272869439 | 5.081243072 | 3.75E-07 | 2.67E-05 | Oas1a |
| ENSMUSG00000031304.14 | 1316.822393 | -1.065507126 | 0.209925268 | -5.075649709 | 3.86E-07 | 2.73E-05 | Il2rg |
| ENSMUSG00000038623.5 | 263.7182712 | 1.218697097 | 0.240217082 | 5.073315704 | 3.91E-07 | 2.75E-05 | Tm6sf1 |
| ENSMUSG00000024011.12 | 72.63905425 | 2.218223887 | 0.438317831 | 5.060765792 | 4.18E-07 | 2.93E-05 | Pi16 |
| ENSMUSG00000030761.11 | 59.62538023 | 1.819122005 | 0.360068588 | 5.052154135 | 4.37E-07 | 3.05E-05 | Myo7a |
| ENSMUSG00000036273.11 | 248.7690583 | 2.052150384 | 0.406485209 | 5.048524122 | 4.45E-07 | 3.09E-05 | Lrrk2 |
| ENSMUSG00000001995.8 | 95.74847485 | 1.617340748 | 0.320778498 | 5.041923815 | 4.61E-07 | 3.18E-05 | Sipa1l2 |
| ENSMUSG00000050989.9 | 87.87262914 | 1.417715943 | 0.28236078 | 5.020937898 | 5.14E-07 | 3.53E-05 | Sepn1 |
| ENSMUSG00000066026.10 | 165.693021 | -1.194342506 | 0.238065588 | -5.016863274 | 5.25E-07 | 3.59E-05 | Dhrs3 |
| ENSMUSG00000087265.1 | 15.22783347 | -2.53944327 | 0.507012583 | -5.008639539 | 5.48E-07 | 3.73E-05 | Gm12349 |
| ENSMUSG00000104475.1 | 34.85983981 | 2.08002889 | 0.417512979 | 4.981950246 | 6.29E-07 | 4.26E-05 | D630036G22Rik |
| ENSMUSG00000045071.9 | 176.3690255 | 1.63719987 | 0.328897496 | 4.977842307 | 6.43E-07 | 4.33E-05 | E130308A19Rik |
| ENSMUSG00000019951.9 | 1341.56714 | -1.070502111 | 0.215567845 | -4.965963791 | 6.84E-07 | 4.58E-05 | Uhrf1bp1l |
| ENSMUSG00000028525.12 | 1089.371903 | -1.427196807 | 0.28764027 | -4.961742001 | 6.99E-07 | 4.66E-05 | Pde4b |
| ENSMUSG00000055629.4 | 350.591638 | -1.611114834 | 0.324967016 | -4.957779572 | 7.13E-07 | 4.73E-05 | B4galnt4 |
| ENSMUSG00000030775.9 | 441.0072925 | -1.275196781 | 0.257372029 | -4.954682863 | 7.24E-07 | 4.79E-05 | Trat1 |
| ENSMUSG00000093661.1 | 334.542358 | -1.01651658 | 0.205417087 | -4.948549287 | 7.48E-07 | 4.92E-05 | Eif4e3 |
| ENSMUSG00000000686.11 | 97.6240591 | -1.396358393 | 0.282311857 | -4.946155676 | 7.57E-07 | 4.95E-05 | Abhd15 |
| ENSMUSG00000001156.7 | 989.7584465 | 1.211458462 | 0.244976311 | 4.945206563 | 7.61E-07 | 4.95E-05 | Mxd1 |
| ENSMUSG00000049807.12 | 62.66925843 | 1.990064331 | 0.402503804 | 4.944212485 | 7.65E-07 | 4.96E-05 | Arhgap23 |
| ENSMUSG00000027374.8 | 261.4751546 | -1.432775291 | 0.29059657 | -4.930461807 | 8.20E-07 | 5.29E-05 | Mrps5 |
| ENSMUSG00000003623.4 | 1139.576557 | -0.967744607 | 0.196378147 | -4.927964862 | 8.31E-07 | 5.34E-05 | Crot |
| ENSMUSG00000028456.13 | 21.47030431 | 2.371850638 | 0.481443127 | 4.926543774 | 8.37E-07 | 5.35E-05 | Unc13b |
| ENSMUSG00000076470.1 | 41.50205596 | -2.256839513 | 0.458708876 | -4.919982216 | 8.66E-07 | 5.51E-05 | Trbv13-3 |
| ENSMUSG00000024235.6 | 545.5725251 | -0.929410547 | 0.189320685 | -4.909186483 | 9.15E-07 | 5.79E-05 | Map3k8 |
| ENSMUSG00000059316.2 | 484.3609569 | 1.233168127 | 0.251792784 | 4.897551508 | 9.70E-07 | 6.11E-05 | Slc27a4 |
| ENSMUSG00000079186.2 | 20.94506215 | -2.361771202 | 0.482305689 | -4.896834636 | 9.74E-07 | 6.11E-05 | Gzmc |
| ENSMUSG00000026893.4 | 72.06690292 | 2.012640928 | 0.412937726 | 4.873957506 | 1.09E-06 | 6.83E-05 | Gca |
| ENSMUSG00000035900.14 | 779.5451792 | 1.112330609 | 0.228459246 | 4.868836024 | 1.12E-06 | 6.98E-05 | Gramd4 |
| ENSMUSG00000080538.1 | 204.1939653 | -1.989008912 | 0.409467386 | -4.857551495 | 1.19E-06 | 7.36E-05 | Gm25541 |
| ENSMUSG00000022900.10 | 185.3814296 | 1.306197974 | 0.268983092 | 4.856059786 | 1.20E-06 | 7.38E-05 | Ildr1 |
| ENSMUSG00000054555.7 | 56.64238906 | -2.210704963 | 0.45599105 | -4.848132355 | 1.25E-06 | 7.65E-05 | Adam12 |
| ENSMUSG00000025877.10 | 299.4849519 | 2.034577075 | 0.42066291 | 4.836597254 | 1.32E-06 | 8.07E-05 | Hk3 |
| ENSMUSG00000032420.7 | 211.8899234 | -1.618981246 | 0.335306057 | -4.828368634 | 1.38E-06 | 8.37E-05 | Nt5e |
| ENSMUSG00000025809.11 | 3567.745921 | -1.701721465 | 0.352680231 | -4.825111576 | 1.40E-06 | 8.47E-05 | Itgb1 |
| ENSMUSG00000096054.2 | 1092.866839 | 1.300249471 | 0.269952388 | 4.816588142 | 1.46E-06 | 8.81E-05 | Syne1 |
| ENSMUSG00000005087.13 | 2734.327683 | -1.001334649 | 0.208200311 | -4.809477204 | 1.51E-06 | 9.09E-05 | Cd44 |
| ENSMUSG00000034586.10 | 421.5806861 | -1.281380942 | 0.266990074 | -4.799357985 | 1.59E-06 | 9.52E-05 | Hid1 |
| ENSMUSG00000055013.10 | 101.6014036 | 2.094336 | 0.436910106 | 4.793516951 | 1.64E-06 | 9.75E-05 | Agap1 |
| ENSMUSG00000030792.7 | 74.91476921 | -1.749397769 | 0.365439679 | -4.787104055 | 1.69E-06 | 0.000100281 | Dkkl1 |
| ENSMUSG00000034573.10 | 837.7670725 | 1.207683314 | 0.252616576 | 4.780697031 | 1.75E-06 | 0.000103029 | Ptpn13 |
| ENSMUSG00000066278.5 | 344.4186717 | 1.2651138 | 0.264670545 | 4.779956901 | 1.75E-06 | 0.000103029 | Vps37b |
| ENSMUSG00000005534.9 | 190.724084 | 1.193885338 | 0.249934234 | 4.77679796 | 1.78E-06 | 0.000104016 | Insr |
| ENSMUSG00000038594.8 | 182.3933834 | 1.112237953 | 0.232905709 | 4.775485999 | 1.79E-06 | 0.000104016 | Cep85l |
| ENSMUSG00000055044.8 | 148.4245025 | -1.312370499 | 0.274786618 | -4.775962193 | 1.79E-06 | 0.000104016 | Pdlim1 |
| ENSMUSG00000021831.8 | 1046.096595 | -1.476155984 | 0.30925898 | -4.773203305 | 1.81E-06 | 0.000104763 | Ero1l |
| ENSMUSG00000045362.7 | 248.2450465 | -1.288103159 | 0.27061862 | -4.759846744 | 1.94E-06 | 0.000111473 | Tnfrsf26 |
| ENSMUSG00000040703.7 | 59.67405253 | -1.582358958 | 0.332803595 | -4.754633006 | 1.99E-06 | 0.000113914 | Cyp2s1 |
| ENSMUSG00000025429.8 | 36.00984392 | 2.119864064 | 0.446636941 | 4.746280189 | 2.07E-06 | 0.000118228 | Pstpip2 |
| ENSMUSG00000089525.1 | 22.02989177 | 2.316466558 | 0.489069595 | 4.736476322 | 2.17E-06 | 0.00012358 | Gm23833 |
| ENSMUSG00000028874.10 | 219.0701877 | 1.809431633 | 0.382292463 | 4.733108315 | 2.21E-06 | 0.000124831 | Fgr |
| ENSMUSG00000041598.7 | 47.88918974 | 1.699028546 | 0.358992408 | 4.73277013 | 2.21E-06 | 0.000124831 | Cdc42ep4 |
| ENSMUSG00000087497.3 | 184.2411836 | -1.509048338 | 0.319059129 | -4.72968238 | 2.25E-06 | 0.00012623 | 2810001G20Rik |

| | | | | | | | |
|---|---|---|---|---|---|---|---|
| ENSMUSG00000009628.10 | 56.26516914 | 1.563672322 | 0.330921263 | 4.725209574 | 2.30E-06 | 0.000128518 | Tex15 |
| ENSMUSG00000025375.11 | 35.59590553 | 2.259784838 | 0.479118805 | 4.716543823 | 2.40E-06 | 0.000133571 | Aatk |
| ENSMUSG00000096900.2 | 20.80604015 | 2.295314485 | 0.486799731 | 4.715110417 | 2.42E-06 | 0.000133975 | Trav9-1 |
| ENSMUSG00000021614.12 | 219.8745679 | 1.973194176 | 0.418872325 | 4.710729404 | 2.47E-06 | 0.00013634 | Vcan |
| ENSMUSG00000017485.6 | 3285.72589 | -0.857870029 | 0.183297629 | -4.680202533 | 2.87E-06 | 0.000157671 | Top2b |
| ENSMUSG00000022237.12 | 23.57451001 | 2.127939265 | 0.454920522 | 4.677606657 | 2.90E-06 | 0.000159046 | Ankrd33b |
| ENSMUSG00000026950.12 | 541.7655748 | 1.712080701 | 0.367368577 | 4.660389612 | 3.16E-06 | 0.000172264 | Neb |
| ENSMUSG00000025427.10 | 18.41180292 | 2.426090957 | 0.52083749 | 4.65805746 | 3.19E-06 | 0.000173541 | Rnf165 |
| ENSMUSG00000027398.9 | 153.739144 | 2.139866413 | 0.459632813 | 4.655599762 | 3.23E-06 | 0.00017466 | Il1b |
| ENSMUSG00000049804.9 | 31.28198887 | 1.922880209 | 0.413139266 | 4.654314817 | 3.25E-06 | 0.00017466 | Armcx4 |
| ENSMUSG00000055725.7 | 52.05717027 | 1.429507371 | 0.307130129 | 4.654402928 | 3.25E-06 | 0.00017466 | Paqr3 |
| ENSMUSG00000039021.11 | 44.88728697 | 1.695541246 | 0.36449916 | 4.651701385 | 3.29E-06 | 0.000176203 | Ttc16 |
| ENSMUSG00000063410.7 | 2691.392386 | -0.908381035 | 0.195496942 | -4.646522986 | 3.38E-06 | 0.000179984 | Stk24 |
| ENSMUSG00000042082.6 | 1384.722787 | 1.03671568 | 0.223209349 | 4.644588962 | 3.41E-06 | 0.000180772 | Arsb |
| ENSMUSG00000045078.8 | 408.78075 | 1.058068534 | 0.227833931 | 4.644034061 | 3.42E-06 | 0.000180772 | Rnf216 |
| ENSMUSG00000070291.4 | 31.32964807 | 2.211154101 | 0.477218465 | 4.633421088 | 3.60E-06 | 0.000189569 | Ddx43 |
| ENSMUSG00000017466.5 | 221.1707683 | -0.974314506 | 0.210643272 | -4.625424278 | 3.74E-06 | 0.000196283 | Timp2 |
| ENSMUSG00000079138.3 | 67.22560444 | -1.664160952 | 0.360264584 | -4.619274345 | 3.85E-06 | 0.000201425 | Gm8818 |
| ENSMUSG00000024053.10 | 169.2642821 | 2.04767238 | 0.444556705 | 4.606099415 | 4.10E-06 | 0.000213801 | Emilin2 |
| ENSMUSG00000058099.11 | 242.5452174 | 1.993458461 | 0.433091318 | 4.602859438 | 4.17E-06 | 0.000216339 | Nfam1 |
| ENSMUSG00000025461.10 | 145.5555277 | -1.642869284 | 0.357079854 | -4.600845619 | 4.21E-06 | 0.000216811 | Cd163l1 |
| ENSMUSG00000037003.11 | 132.9948355 | -1.476843966 | 0.320967838 | -4.601221032 | 4.20E-06 | 0.000216811 | Tns2 |
| ENSMUSG00000019312.6 | 61.39259995 | 1.591733992 | 0.346637597 | 4.591925415 | 4.39E-06 | 0.000225449 | Grb7 |
| ENSMUSG00000029528.13 | 1057.4495 | 0.896422292 | 0.195327038 | 4.589340539 | 4.45E-06 | 0.000226053 | Pxn |
| ENSMUSG00000040451.13 | 1225.062963 | -0.942410435 | 0.2053606 | -4.589051822 | 4.45E-06 | 0.000226053 | Sgms1 |
| ENSMUSG00000049625.5 | 48.22290115 | 1.845451917 | 0.402131136 | 4.589179376 | 4.45E-06 | 0.000226053 | Tifab |
| ENSMUSG00000052234.2 | 10.07443378 | 2.456312819 | 0.535545338 | 4.586563724 | 4.51E-06 | 0.000227925 | Epx |
| ENSMUSG00000045092.7 | 2116.900441 | -1.08890778 | 0.237534206 | -4.584214613 | 4.56E-06 | 0.000229661 | S1pr1 |
| ENSMUSG00000047507.8 | 2371.156195 | 1.825759939 | 0.398351171 | 4.583292512 | 4.58E-06 | 0.000229838 | Baiap3 |
| ENSMUSG00000021061.11 | 95.04326642 | 2.116221718 | 0.461973597 | 4.580828283 | 4.63E-06 | 0.00023172 | Sptb |
| ENSMUSG00000031824.10 | 214.0457984 | 1.958675858 | 0.428788335 | 4.567931776 | 4.93E-06 | 0.000244668 | 6430548M08Rik |
| ENSMUSG00000074607.7 | 138.1121201 | 1.978908726 | 0.433201333 | 4.568103964 | 4.92E-06 | 0.000244668 | Tox2 |
| ENSMUSG00000024124.5 | 26.47713303 | -1.980929682 | 0.433887941 | -4.565532931 | 4.98E-06 | 0.000246509 | Prss30 |
| ENSMUSG00000038213.7 | 453.7741059 | 1.232453184 | 0.269987322 | 4.564855775 | 5.00E-06 | 0.000246509 | Tapbpl |
| ENSMUSG00000022861.12 | 40.24051374 | 2.206129778 | 0.483448074 | 4.56332313 | 5.04E-06 | 0.000247433 | Dgkg |
| ENSMUSG00000026928.10 | 87.85527082 | 1.867786968 | 0.409498541 | 4.561156593 | 5.09E-06 | 0.000249113 | Card9 |
| ENSMUSG00000042035.7 | 19.89934716 | 2.221851089 | 0.487652499 | 4.556217993 | 5.21E-06 | 0.000253911 | Igsf3 |
| ENSMUSG00000065232.1 | 1878.212839 | -1.261416208 | 0.276889585 | -4.555665066 | 5.22E-06 | 0.000253911 | Gm22973 |
| ENSMUSG00000045973.14 | 770.7323285 | -0.797805007 | 0.175505036 | -4.545767031 | 5.47E-06 | 0.000265208 | Slc25a51 |
| ENSMUSG00000020889.11 | 28.8340876 | -2.204009245 | 0.485257031 | -4.54194191 | 5.57E-06 | 0.000269123 | Nr1d1 |
| ENSMUSG00000034023.12 | 268.1142488 | 0.90451604 | 0.19957013 | 4.532321754 | 5.83E-06 | 0.000280698 | Fancd2 |
| ENSMUSG00000015533.8 | 337.756251 | -1.430528701 | 0.316066943 | -4.526030747 | 6.01E-06 | 0.000283657 | Itga2 |
| ENSMUSG00000016087.9 | 1427.068186 | -0.931119666 | 0.205738958 | -4.525733364 | 6.02E-06 | 0.000283657 | Fli1 |
| ENSMUSG00000026980.11 | 966.3734477 | 0.840676803 | 0.185684589 | 4.527445202 | 5.97E-06 | 0.000283657 | Ly75 |
| ENSMUSG00000035199.6 | 1224.568418 | -0.883359333 | 0.195167269 | -4.52616536 | 6.01E-06 | 0.000283657 | Arl6ip5 |
| ENSMUSG00000039512.11 | 113.2376934 | 1.157451229 | 0.255676766 | 4.527009807 | 5.98E-06 | 0.000283657 | Uhrf1bp1 |
| ENSMUSG00000042724.7 | 149.2985829 | 2.078184031 | 0.45885641 | 4.529050887 | 5.92E-06 | 0.000283657 | Map3k9 |
| ENSMUSG00000022126.6 | 53.69310302 | 2.079485566 | 0.459603005 | 4.524525613 | 6.05E-06 | 0.000284311 | Irg1 |
| ENSMUSG00000033147.12 | 368.0752728 | 1.05537291 | 0.233992161 | 4.510291729 | 6.47E-06 | 0.000303042 | Slc22a15 |
| ENSMUSG00000025321.10 | 23.49783995 | 1.999582228 | 0.443846191 | 4.505124223 | 6.63E-06 | 0.000309462 | Itgb8 |
| ENSMUSG00000028221.3 | 129.1075093 | -1.242745259 | 0.276047656 | -4.501922876 | 6.73E-06 | 0.000313104 | Tmem55a |
| ENSMUSG00000030854.11 | 21.04929964 | 1.957830933 | 0.435164724 | 4.499057077 | 6.83E-06 | 0.000316289 | Ptpn5 |
| ENSMUSG00000033400.10 | 103.7187281 | 1.734528362 | 0.38612546 | 4.492136731 | 7.05E-06 | 0.000325653 | Agl |
| ENSMUSG00000020601.7 | 897.1089488 | -1.032547998 | 0.229969808 | -4.489928521 | 7.12E-06 | 0.00032795 | Trib2 |
| ENSMUSG00000045671.13 | 171.4436071 | 1.172192425 | 0.261673196 | 4.479604489 | 7.48E-06 | 0.000343076 | Spred2 |
| ENSMUSG00000024164.11 | 855.9257937 | 1.940586455 | 0.433437588 | 4.477199273 | 7.56E-06 | 0.000344672 | C3 |
| ENSMUSG00000031834.11 | 69.52737758 | 1.278653072 | 0.285548498 | 4.477884069 | 7.54E-06 | 0.000344672 | Pik3r2 |
| ENSMUSG00000002835.8 | 392.4497014 | 0.874753128 | 0.195553601 | 4.473214112 | 7.71E-06 | 0.000350006 | Chaf1a |
| ENSMUSG00000030852.11 | 68.48198766 | 1.714508208 | 0.383454892 | 4.471212247 | 7.78E-06 | 0.000351188 | Tacc2 |
| ENSMUSG00000040760.6 | 991.1615348 | -0.920326028 | 0.205839225 | -4.471091597 | 7.78E-06 | 0.000351188 | Appl1 |
| ENSMUSG00000019982.10 | 263.2337903 | 1.473928411 | 0.329757852 | 4.469729534 | 7.83E-06 | 0.000352281 | Myb |
| ENSMUSG00000039153.12 | 1723.919398 | -0.802602684 | 0.17991808 | -4.460934017 | 8.16E-06 | 0.000364679 | Runx2 |

| | | | | | | | |
|---|---|---|---|---|---|---|---|
| ENSMUSG00000040907.11 | 94.22857234 | 1.846315527 | 0.413878783 | 4.461005501 | 8.16E-06 | 0.000364679 | Atp1a3 |
| ENSMUSG00000047798.11 | 203.1435857 | 1.996294868 | 0.447592822 | 4.460068999 | 8.19E-06 | 0.000364973 | Cd300lf |
| ENSMUSG00000022788.12 | 105.7295915 | 1.942920167 | 0.436030132 | 4.455930964 | 8.35E-06 | 0.000370889 | Fgd4 |
| ENSMUSG00000086040.4 | 8.798460051 | 2.366098154 | 0.531265343 | 4.453703189 | 8.44E-06 | 0.000373559 | Wipf3 |
| ENSMUSG00000087066.1 | 17.54346058 | -2.162966662 | 0.486162613 | -4.449060058 | 8.62E-06 | 0.000380506 | Gm15518 |
| ENSMUSG00000050621.6 | 315.9569957 | -1.684506125 | 0.379043312 | -4.444099325 | 8.83E-06 | 0.000388151 | Rps27rt |
| ENSMUSG00000018925.3 | 262.0490431 | 1.888206991 | 0.425061137 | 4.442200957 | 8.90E-06 | 0.000390349 | Heatr9 |
| ENSMUSG00000068220.5 | 5149.453123 | -1.232529589 | 0.277678052 | -4.438700066 | 9.05E-06 | 0.000395496 | Lgals1 |
| ENSMUSG00000050075.8 | 502.3509046 | -0.951940557 | 0.21465449 | -4.43475726 | 9.22E-06 | 0.000401533 | Gpr171 |
| ENSMUSG00000043733.10 | 1388.195514 | 0.70477422 | 0.158979712 | 4.433107917 | 9.29E-06 | 0.000403346 | Ptpn11 |
| ENSMUSG00000021716.10 | 166.6456298 | -1.195543839 | 0.269928984 | -4.429105105 | 9.46E-06 | 0.000409616 | Srek1ip1 |
| ENSMUSG00000041801.5 | 44.37879418 | 1.644985073 | 0.371530331 | 4.427592947 | 9.53E-06 | 0.000409927 | Phlda3 |
| ENSMUSG00000102662.1 | 68.20523079 | 1.852910777 | 0.418480895 | 4.427706972 | 9.52E-06 | 0.000409927 | Gm38377 |
| ENSMUSG00000042747.8 | 311.3836707 | -1.143974339 | 0.258426732 | -4.426687324 | 9.57E-06 | 0.000410373 | Krtcap2 |
| ENSMUSG00000034833.9 | 319.5733554 | -0.966948236 | 0.218903735 | -4.417230421 | 1.00E-05 | 0.000427411 | Tespa1 |
| ENSMUSG00000009739.12 | 280.4839046 | 1.586657655 | 0.359701945 | 4.411034404 | 1.03E-05 | 0.000437191 | Pou6f1 |
| ENSMUSG00000020009.8 | 4554.66895 | -1.24766106 | 0.282852216 | -4.410999771 | 1.03E-05 | 0.000437191 | Ifngr1 |
| ENSMUSG00000044320.10 | 31.67903263 | 2.029717881 | 0.46039565 | 4.40863827 | 1.04E-05 | 0.000440628 | 1700001O22Rik |
| ENSMUSG00000044505.4 | 13.2909103 | -2.291801607 | 0.521378018 | -4.395662122 | 1.10E-05 | 0.000466362 | Lingo4 |
| ENSMUSG00000052062.10 | 10.50815717 | 2.264747077 | 0.515308645 | 4.394933208 | 1.11E-05 | 0.000466503 | Pard3b |
| ENSMUSG00000040964.12 | 36.31303856 | 2.032762893 | 0.463077098 | 4.389685656 | 1.14E-05 | 0.000476451 | Arhgef10l |
| ENSMUSG00000017861.7 | 325.9562623 | 1.10830344 | 0.252743762 | 4.385087219 | 1.16E-05 | 0.000480783 | Mybl2 |
| ENSMUSG00000031397.7 | 17.69587419 | -2.287277043 | 0.521579042 | -4.385293235 | 1.16E-05 | 0.000480783 | Tktl1 |
| ENSMUSG00000058290.3 | 563.0619038 | 1.058512038 | 0.241318777 | 4.386364172 | 1.15E-05 | 0.000480783 | Espl1 |
| ENSMUSG00000064797.1 | 130.3560315 | -1.514845114 | 0.345339521 | -4.386538527 | 1.15E-05 | 0.000480783 | Gm24357 |
| ENSMUSG00000070604.3 | 56.39944039 | 1.541048742 | 0.351548338 | 4.383604118 | 1.17E-05 | 0.00048262 | Vsig10l |
| ENSMUSG00000018476.7 | 662.9295442 | 1.198371354 | 0.273457472 | 4.382295149 | 1.17E-05 | 0.00048408 | Kdm6b |
| ENSMUSG00000028961.11 | 589.3471091 | 1.062676741 | 0.242702665 | 4.378512863 | 1.19E-05 | 0.00049109 | Pgd |
| ENSMUSG00000028028.7 | 28.5672946 | 1.861357816 | 0.42539135 | 4.375636255 | 1.21E-05 | 0.000496136 | Alpk1 |
| ENSMUSG00000079298.5 | 26.63964866 | -1.96284733 | 0.448859095 | -4.37296994 | 1.23E-05 | 0.00050075 | Klrb1b |
| ENSMUSG00000026594.10 | 350.1512616 | -0.975730417 | 0.223330944 | -4.368988901 | 1.25E-05 | 0.000508461 | Ralgps2 |
| ENSMUSG00000063810.5 | 344.2245514 | 0.880185127 | 0.201523378 | 4.36765767 | 1.26E-05 | 0.000510064 | Alms1 |
| ENSMUSG00000024317.10 | 586.4658538 | -1.104115617 | 0.253464224 | -4.356100435 | 1.32E-05 | 0.000534594 | Rnf138 |
| ENSMUSG00000047898.6 | 113.4718436 | -1.720662892 | 0.394952805 | -4.35662912 | 1.32E-05 | 0.000534594 | Ccr4 |
| ENSMUSG00000024621.11 | 246.361924 | 1.749659502 | 0.401993071 | 4.352461843 | 1.35E-05 | 0.000541962 | Csf1r |
| ENSMUSG00000032815.11 | 382.3768688 | 1.313024628 | 0.301728414 | 4.351677091 | 1.35E-05 | 0.000542325 | Fanca |
| ENSMUSG00000030263.9 | 987.9462166 | 1.061248854 | 0.243945281 | 4.350356155 | 1.36E-05 | 0.000544021 | Lrmp |
| ENSMUSG00000026384.9 | 1255.531294 | -0.773077653 | 0.177860795 | -4.346532081 | 1.38E-05 | 0.000551988 | Ptpn4 |
| ENSMUSG00000003484.4 | 37.44517908 | 2.112969103 | 0.486410876 | 4.344000526 | 1.40E-05 | 0.000556782 | Cyp4f18 |
| ENSMUSG00000030748.8 | 603.121645 | 0.93710647 | 0.216182159 | 4.334800223 | 1.46E-05 | 0.000578917 | Il4ra |
| ENSMUSG00000012123.11 | 82.91260164 | -1.440487754 | 0.332397957 | -4.33362397 | 1.47E-05 | 0.000580352 | Aim1l |
| ENSMUSG00000037346.4 | 22.50697254 | -2.2527129 | 0.519975461 | -4.332344638 | 1.48E-05 | 0.000582068 | Hrh4 |
| ENSMUSG00000086425.3 | 42.25756206 | 1.902678538 | 0.439312551 | 4.331036151 | 1.48E-05 | 0.00058387 | F730016J06Rik |
| ENSMUSG00000102151.1 | 262.8022882 | 1.282355561 | 0.296571152 | 4.323938967 | 1.53E-05 | 0.000601271 | Gm37472 |
| ENSMUSG00000040274.7 | 1367.538489 | 0.969114849 | 0.224179626 | 4.322939012 | 1.54E-05 | 0.000602293 | Cdk6 |
| ENSMUSG00000034930.11 | 69.39340939 | 1.953542661 | 0.452502945 | 4.317193252 | 1.58E-05 | 0.000616435 | Rtkn |
| ENSMUSG00000029163.7 | 65.39298313 | 1.718682326 | 0.398706133 | 4.310649335 | 1.63E-05 | 0.000631739 | Emilin1 |
| ENSMUSG00000029406.11 | 562.8139946 | 1.185546061 | 0.275034908 | 4.310529415 | 1.63E-05 | 0.000631739 | Pitpnm2 |
| ENSMUSG00000027506.11 | 126.7021774 | 1.603867695 | 0.372229089 | 4.308818801 | 1.64E-05 | 0.000634861 | Tpd52 |
| ENSMUSG00000040322.6 | 1092.716212 | -0.777571897 | 0.18049502 | -4.307996399 | 1.65E-05 | 0.000635446 | Slc25a24 |
| ENSMUSG00000032228.12 | 1145.09275 | -0.685366826 | 0.159168561 | -4.305918352 | 1.66E-05 | 0.000639654 | Tcf12 |
| ENSMUSG00000104262.1 | 44.39236284 | 1.63063151 | 0.378806303 | 4.304657812 | 1.67E-05 | 0.00064152 | Gm37747 |
| ENSMUSG00000020949.8 | 330.6687716 | -1.055377688 | 0.245814136 | -4.293397061 | 1.76E-05 | 0.000673083 | Fkbp3 |
| ENSMUSG00000084956.3 | 24.34176422 | 2.110090054 | 0.491823734 | 4.290337995 | 1.78E-05 | 0.000680537 | Gm16194 |
| ENSMUSG00000080984.1 | 12.0115008 | 2.266723636 | 0.528590139 | 4.288244272 | 1.80E-05 | 0.00068509 | Gm14469 |
| ENSMUSG00000014932.11 | 41.15422694 | -1.7010941 | 0.396757936 | -4.287486015 | 1.81E-05 | 0.000685543 | Yes1 |
| ENSMUSG00000046711.11 | 273.3946532 | 0.899975226 | 0.210053524 | 4.284504301 | 1.83E-05 | 0.000692898 | Hmga1 |
| ENSMUSG00000035258.10 | 10.77847269 | -2.249035602 | 0.525545251 | -4.279432834 | 1.87E-05 | 0.000705012 | Abi3bp |
| ENSMUSG00000097006.3 | 40.77750022 | 1.576812268 | 0.368433365 | 4.279775991 | 1.87E-05 | 0.000705012 | 9530082P21Rik |
| ENSMUSG00000020486.14 | 16.18929605 | 2.143958319 | 0.501222819 | 4.27745553 | 1.89E-05 | 0.000709369 | Sept4 |
| ENSMUSG00000059920.9 | 579.5654573 | -0.958009634 | 0.224156592 | -4.273841003 | 1.92E-05 | 0.000719019 | 4930453N24Rik |
| ENSMUSG00000095574.2 | 82.54107359 | -1.861739749 | 0.435891593 | -4.271107264 | 1.95E-05 | 0.000725924 | Trbv12-1 |

| | | | | | | | |
|---|---|---|---|---|---|---|---|
| ENSMUSG00000050229.3 | 817.0227343 | -0.736795599 | 0.172562735 | -4.26972601 | 1.96E-05 | 0.000728465 | Pigm |
| ENSMUSG00000004610.3 | 385.3143903 | 0.874567333 | 0.2049245 | 4.267753889 | 1.97E-05 | 0.000732957 | Etfb |
| ENSMUSG00000025058.4 | 123.641133 | -1.475779688 | 0.34608026 | -4.264270055 | 2.01E-05 | 0.000740505 | 5430427O19Rik |
| ENSMUSG00000062157.6 | 34.50641437 | 2.045123695 | 0.479582352 | 4.264384809 | 2.00E-05 | 0.000740505 | Ifnlr1 |
| ENSMUSG00000076467.3 | 54.76204313 | -1.719618199 | 0.403420747 | -4.262592367 | 2.02E-05 | 0.000744099 | Trbv13-1 |
| ENSMUSG00000033350.7 | 509.2385412 | 1.075539743 | 0.252405689 | 4.26115492 | 2.03E-05 | 0.00074691 | Chst2 |
| ENSMUSG00000022724.11 | 723.114422 | -1.141831512 | 0.268325595 | -4.255395434 | 2.09E-05 | 0.000764366 | Mina |
| ENSMUSG00000098178.1 | 36092.72949 | 1.746290092 | 0.411046247 | 4.24840296 | 2.15E-05 | 0.000786528 | Yam1 |
| ENSMUSG00000049985.10 | 11.64285217 | 2.209456151 | 0.520293102 | 4.246560532 | 2.17E-05 | 0.000790928 | Ankrd55 |
| ENSMUSG00000028602.8 | 29.75434348 | 1.740219286 | 0.410076634 | 4.243644092 | 2.20E-05 | 0.000799174 | Tnfrsf8 |
| ENSMUSG00000033767.10 | 961.6141259 | 0.898347632 | 0.211828118 | 4.240927229 | 2.23E-05 | 0.000806787 | D930015E06Rik |
| ENSMUSG00000025647.12 | 4791.414608 | -0.875697506 | 0.206834443 | -4.233808907 | 2.30E-05 | 0.000830139 | Shisa5 |
| ENSMUSG00000026480.8 | 319.4402527 | 1.810241241 | 0.427615118 | 4.233342476 | 2.30E-05 | 0.000830139 | Ncf2 |
| ENSMUSG00000018899.12 | 3578.509004 | 0.859783659 | 0.203161034 | 4.232030327 | 2.32E-05 | 0.000832824 | Irf1 |
| ENSMUSG00000037366.10 | 30.20540628 | 1.685394369 | 0.398439136 | 4.229992023 | 2.34E-05 | 0.000838223 | Pafah2 |
| ENSMUSG00000051177.12 | 22.78079107 | 2.196402795 | 0.519327835 | 4.22931845 | 2.34E-05 | 0.000838558 | Plcb1 |
| ENSMUSG00000017737.2 | 396.4431154 | 1.783583707 | 0.422095992 | 4.225540497 | 2.38E-05 | 0.000850549 | Mmp9 |
| ENSMUSG00000046949.11 | 323.5156453 | -1.331203926 | 0.315219019 | -4.22310789 | 2.41E-05 | 0.000855364 | Nqo2 |
| ENSMUSG00000079020.5 | 248.5734793 | 0.93187146 | 0.220657091 | 4.223165713 | 2.41E-05 | 0.000855364 | Slc45a4 |
| ENSMUSG00000034312.9 | 658.3417457 | 1.205814568 | 0.285606324 | 4.221946317 | 2.42E-05 | 0.000857581 | Iqsec1 |
| ENSMUSG00000036377.14 | 18.26092046 | 1.920597347 | 0.454981461 | 4.221265062 | 2.43E-05 | 0.000857977 | C530008M17Rik |
| ENSMUSG00000032841.11 | 88.22996141 | 1.512432352 | 0.358377324 | 4.220223352 | 2.44E-05 | 0.000859753 | Prr5l |
| ENSMUSG00000060112.1 | 145.8797144 | -1.619723243 | 0.383993049 | -4.218105639 | 2.46E-05 | 0.000865657 | Olfr60 |
| ENSMUSG00000017897.14 | 57.21328653 | -1.809072684 | 0.429140675 | -4.215570296 | 2.49E-05 | 0.000873219 | Eya2 |
| ENSMUSG00000043008.8 | 1308.338412 | -1.452131107 | 0.344714244 | -4.212564844 | 2.52E-05 | 0.000882682 | Klhl6 |
| ENSMUSG00000037826.3 | 349.8988898 | -1.014449052 | 0.240888569 | -4.211279326 | 2.54E-05 | 0.00088548 | Ppm1k |
| ENSMUSG00000008999.7 | 26.38324866 | 2.076987548 | 0.493363025 | 4.209856518 | 2.56E-05 | 0.00088883 | Bmp7 |
| ENSMUSG00000009418.11 | 65.51823352 | -1.585368294 | 0.376898037 | -4.206358583 | 2.60E-05 | 0.000900424 | Nav1 |
| ENSMUSG00000005686.12 | 90.71773364 | 1.754197105 | 0.417494572 | 4.201724336 | 2.65E-05 | 0.000916757 | Ampd3 |
| ENSMUSG00000042063.11 | 423.4212755 | -0.885499866 | 0.210895795 | -4.198755443 | 2.68E-05 | 0.000926536 | Zfp386 |
| ENSMUSG00000045991.14 | 35.0638126 | -2.132728067 | 0.508188566 | -4.196725801 | 2.71E-05 | 0.000932543 | Onecut2 |
| ENSMUSG00000023075.9 | 663.5526532 | -0.901155381 | 0.214819834 | -4.194935656 | 2.73E-05 | 0.0009376 | Akirin1 |
| ENSMUSG00000042129.6 | 124.127278 | 1.73011492 | 0.412531756 | 4.193895127 | 2.74E-05 | 0.000939577 | Rassf4 |
| ENSMUSG00000020841.5 | 641.9998297 | -1.600649574 | 0.381805913 | -4.192312161 | 2.76E-05 | 0.000941486 | Cpd |
| ENSMUSG00000030061.12 | 366.7696156 | -0.857038451 | 0.204411248 | -4.192716691 | 2.76E-05 | 0.000941486 | Uba3 |
| ENSMUSG00000043740.10 | 146.9240196 | 1.74864499 | 0.417864787 | 4.18471488 | 2.86E-05 | 0.000971134 | B430306N03Rik |
| ENSMUSG00000041959.10 | 3010.816077 | -1.250891537 | 0.299029824 | -4.183166479 | 2.87E-05 | 0.000975372 | S100a10 |
| ENSMUSG00000035042.2 | 2079.72564 | 1.444404487 | 0.346036377 | 4.174140592 | 2.99E-05 | 0.001012366 | Ccl5 |
| ENSMUSG00000052632.11 | 133.1284096 | 1.492858813 | 0.357939199 | 4.170705012 | 3.04E-05 | 0.001025238 | Asap2 |
| ENSMUSG00000000157.11 | 285.8323907 | 1.839392666 | 0.442091474 | 4.1606608 | 3.17E-05 | 0.001066176 | Itgb2l |
| ENSMUSG00000058881.8 | 106.05358 | 1.908163091 | 0.458608389 | 4.16076796 | 3.17E-05 | 0.001066176 | Zfp516 |
| ENSMUSG00000069601.9 | 22.7253661 | 1.952404965 | 0.469486337 | 4.158598047 | 3.20E-05 | 0.001073237 | Ank3 |
| ENSMUSG00000032238.13 | 4232.725363 | -0.787452505 | 0.18979538 | -4.148955065 | 3.34E-05 | 0.001116745 | Rora |
| ENSMUSG00000006169.15 | 2106.459267 | -0.637956687 | 0.153825775 | -4.147267838 | 3.36E-05 | 0.001119582 | Clint1 |
| ENSMUSG00000075122.4 | 169.3201736 | 1.433726614 | 0.345703547 | 4.147271926 | 3.36E-05 | 0.001119582 | Cd80 |
| ENSMUSG00000043102.2 | 71.15965672 | -1.506600939 | 0.363372684 | -4.14615904 | 3.38E-05 | 0.00112231 | Qrfp |
| ENSMUSG00000032462.10 | 158.9182478 | 1.086302564 | 0.262514041 | 4.138074134 | 3.50E-05 | 0.001158704 | Pik3cb |
| ENSMUSG00000033705.12 | 77.28133915 | 1.702763441 | 0.411574559 | 4.137193137 | 3.52E-05 | 0.001158704 | Stard9 |
| ENSMUSG00000034522.9 | 56.42593617 | 1.621144105 | 0.391828698 | 4.137379711 | 3.51E-05 | 0.001158704 | Zfp395 |
| ENSMUSG00000022895.10 | 139.7172501 | 1.212757815 | 0.293493184 | 4.132149851 | 3.59E-05 | 0.00118161 | Ets2 |
| ENSMUSG00000027490.13 | 131.6633979 | 1.069843669 | 0.259118634 | 4.128779359 | 3.65E-05 | 0.001196214 | E2f1 |
| ENSMUSG00000036882.6 | 106.4775294 | 1.338457454 | 0.324298267 | 4.127242076 | 3.67E-05 | 0.001201383 | Arhgap33 |
| ENSMUSG00000041180.9 | 108.9021541 | -1.265190152 | 0.306714474 | -4.124976999 | 3.71E-05 | 0.001210399 | Hectd2 |
| ENSMUSG00000089809.4 | 25.40640393 | 2.177746705 | 0.52819461 | 4.123000621 | 3.74E-05 | 0.001217956 | A930011G23Rik |
| ENSMUSG00000092251.2 | 21.89205939 | 2.036456846 | 0.494001336 | 4.122371129 | 3.75E-05 | 0.001218416 | Trav9n-1 |
| ENSMUSG00000049577.10 | 691.3340339 | 1.015720063 | 0.246667274 | 4.117773902 | 3.83E-05 | 0.00123715 | Zfpm1 |
| ENSMUSG00000097636.3 | 380.3158529 | 1.217192947 | 0.295589679 | 4.117846567 | 3.82E-05 | 0.00123715 | 5830416P10Rik |
| ENSMUSG00000026335.12 | 46.40417357 | 1.736990864 | 0.422142382 | 4.114703801 | 3.88E-05 | 0.001250801 | Pam |
| ENSMUSG00000044026.2 | 313.4056748 | -0.93370173 | 0.227080553 | -4.111764382 | 3.93E-05 | 0.001263884 | Slc35g1 |
| ENSMUSG00000087651.2 | 135.0324505 | -1.451692356 | 0.354152393 | -4.09906126 | 4.15E-05 | 0.001332181 | 1500009L16Rik |
| ENSMUSG00000005640.7 | 49.23980463 | 1.991945005 | 0.486041106 | 4.098305638 | 4.16E-05 | 0.001333435 | Insrr |
| ENSMUSG00000002111.8 | 172.7801632 | 1.819981652 | 0.444168085 | 4.097506579 | 4.18E-05 | 0.001334386 | Spi1 |

| | | | | | | | |
|---|---|---|---|---|---|---|---|
| ENSMUSG00000031788.9 | 9.563302437 | 2.146963138 | 0.524024205 | 4.097068645 | 4.18E-05 | 0.001334386 | Kifc3 |
| ENSMUSG00000040747.9 | 3189.959484 | -1.298478377 | 0.317237364 | -4.093081476 | 4.26E-05 | 0.001354421 | Cd53 |
| ENSMUSG00000066894.10 | 26.79014267 | 1.775201897 | 0.434137868 | 4.089027997 | 4.33E-05 | 0.001375137 | Vsig10 |
| ENSMUSG00000013707.3 | 418.1264149 | -0.928594545 | 0.227422646 | -4.083122601 | 4.44E-05 | 0.001404752 | Tnfaip8l2 |
| ENSMUSG00000022014.10 | 1460.32667 | -0.882104548 | 0.216042351 | -4.083016787 | 4.45E-05 | 0.001404752 | Epsti1 |
| ENSMUSG00000097365.3 | 44.843372 | -1.821820285 | 0.446437591 | -4.080794989 | 4.49E-05 | 0.00141501 | C030034L19Rik |
| ENSMUSG00000080848.3 | 81.69076234 | -1.358433048 | 0.333321899 | -4.075438941 | 4.59E-05 | 0.00144468 | Gm9385 |
| ENSMUSG00000001517.10 | 600.3724847 | 0.890081401 | 0.218527559 | 4.073085351 | 4.64E-05 | 0.001452745 | Foxm1 |
| ENSMUSG00000045629.7 | 14.72602307 | 1.921969154 | 0.471844148 | 4.073313536 | 4.63E-05 | 0.001452745 | Sh3tc2 |
| ENSMUSG00000032018.9 | 301.8713676 | -0.940736755 | 0.231092676 | -4.070820298 | 4.68E-05 | 0.001457032 | Sc5d |
| ENSMUSG00000035640.14 | 41.1877951 | 1.489988994 | 0.365953014 | 4.071530874 | 4.67E-05 | 0.001457032 | Dos |
| ENSMUSG00000042655.4 | 13.66219906 | -2.143557587 | 0.526564107 | -4.070838782 | 4.68E-05 | 0.001457032 | Fam159b |
| ENSMUSG00000061411.8 | 41.52820773 | -1.747992243 | 0.42956415 | -4.069222821 | 4.72E-05 | 0.001463761 | Nol4l |
| ENSMUSG00000026274.7 | 175.5140177 | 0.919742058 | 0.22607592 | 4.068288476 | 4.74E-05 | 0.001466346 | Pask |
| ENSMUSG00000027500.10 | 14.86669645 | -2.07900056 | 0.511548773 | -4.064129697 | 4.82E-05 | 0.001489395 | Stmn2 |
| ENSMUSG00000003283.9 | 240.9203741 | 1.783137433 | 0.438812004 | 4.063556638 | 4.83E-05 | 0.001489726 | Hck |
| ENSMUSG00000026009.10 | 1289.098787 | -1.19421698 | 0.294213549 | -4.059014226 | 4.93E-05 | 0.001515617 | Icos |
| ENSMUSG00000021302.9 | 259.1133309 | -0.996188441 | 0.245707035 | -4.054374924 | 5.03E-05 | 0.001542574 | Ggps1 |
| ENSMUSG00000030474.8 | 49.34652935 | 1.764908983 | 0.435821895 | 4.049610642 | 5.13E-05 | 0.001570825 | Siglece |
| ENSMUSG00000062014.8 | 1207.273842 | -0.799183476 | 0.19737468 | -4.049067868 | 5.14E-05 | 0.001570989 | Gmfb |
| ENSMUSG00000028480.10 | 2431.056467 | -1.008995935 | 0.249418335 | -4.045395996 | 5.22E-05 | 0.001578358 | Glipr2 |
| ENSMUSG00000028702.11 | 248.6401471 | 0.885129671 | 0.218709748 | 4.047051768 | 5.19E-05 | 0.001578358 | Rad54l |
| ENSMUSG00000031543.14 | 82.65784077 | 1.896197096 | 0.468571249 | 4.046763643 | 5.19E-05 | 0.001578358 | Ank1 |
| ENSMUSG00000043629.8 | 15.14864376 | 2.100632621 | 0.519155395 | 4.046250201 | 5.20E-05 | 0.001578358 | 1700019D03Rik |
| ENSMUSG00000100354.1 | 11.82176441 | 2.039496434 | 0.504096886 | 4.045842158 | 5.21E-05 | 0.001578358 | Gm29113 |
| ENSMUSG00000031428.7 | 65.64602354 | 1.523192145 | 0.376734494 | 4.043144887 | 5.27E-05 | 0.001590116 | Zcchc18 |
| ENSMUSG00000033446.7 | 300.6309287 | -1.2820821 | 0.31722217 | -4.04159048 | 5.31E-05 | 0.001594555 | Lpar6 |
| ENSMUSG00000036944.5 | 982.8765229 | -0.780769237 | 0.193189424 | -4.041469883 | 5.31E-05 | 0.001594555 | Tmem71 |
| ENSMUSG00000000782.11 | 1086.223095 | -0.941620627 | 0.233059916 | -4.040251299 | 5.34E-05 | 0.001599386 | Tcf7 |
| ENSMUSG00000024539.13 | 508.8554549 | -0.861853298 | 0.213512303 | -4.036550992 | 5.42E-05 | 0.001621294 | Ptpn2 |
| ENSMUSG00000079523.4 | 3850.196562 | -0.926871207 | 0.229687652 | -4.035354968 | 5.45E-05 | 0.001626056 | Tmsb10 |
| ENSMUSG00000031295.9 | 259.9948786 | 0.879825973 | 0.218525464 | 4.026194273 | 5.67E-05 | 0.001687045 | Phka2 |
| ENSMUSG00000044408.6 | 349.3224083 | -1.122664661 | 0.279189862 | -4.02115125 | 5.79E-05 | 0.001719871 | Sptssa |
| ENSMUSG00000052534.11 | 97.81449425 | 1.863836889 | 0.463941018 | 4.017400524 | 5.88E-05 | 0.001743717 | Pbx1 |
| ENSMUSG00000004609.7 | 83.42874295 | 1.839144686 | 0.457885068 | 4.016607694 | 5.90E-05 | 0.001744871 | Cd33 |
| ENSMUSG00000019876.11 | 43.87482339 | -1.861725695 | 0.463559243 | -4.016154838 | 5.92E-05 | 0.001744871 | Pkib |
| ENSMUSG00000057719.6 | 12.74654011 | 2.072108349 | 0.515997725 | 4.015731563 | 5.93E-05 | 0.001744871 | Sh3rf2 |
| ENSMUSG00000032724.5 | 796.3705375 | 1.166040625 | 0.290405889 | 4.015209982 | 5.94E-05 | 0.001745014 | Abtb2 |
| ENSMUSG00000032249.10 | 770.2848349 | -0.998993546 | 0.249303047 | -4.007145356 | 6.15E-05 | 0.001801827 | Anp32a |
| ENSMUSG00000074695.3 | 9.760902931 | -2.132643819 | 0.532441678 | -4.005403606 | 6.19E-05 | 0.00181131 | Il22 |
| ENSMUSG00000058297.12 | 120.7627679 | 1.614976176 | 0.403693345 | 4.000502349 | 6.32E-05 | 0.001845331 | Spock2 |
| ENSMUSG00000027947.7 | 108.332738 | 1.418937258 | 0.354794172 | 3.999325154 | 6.35E-05 | 0.00185062 | Il6ra |
| ENSMUSG00000025854.11 | 33.06731847 | 1.80016487 | 0.4505843 | 3.995178863 | 6.46E-05 | 0.001879345 | Fam20c |
| ENSMUSG00000059708.8 | 87.15580499 | -1.20946428 | 0.30328612 | -3.987865585 | 6.67E-05 | 0.001934142 | Akap17b |
| ENSMUSG00000078817.3 | 22.92068035 | 1.965509857 | 0.493134349 | 3.985749237 | 6.73E-05 | 0.001947374 | Nlrp12 |
| ENSMUSG00000023915.4 | 64.22054334 | 1.839898246 | 0.462028647 | 3.982216813 | 6.83E-05 | 0.001972418 | Tnfrsf21 |
| ENSMUSG00000038725.7 | 16.08247391 | 1.915875917 | 0.481224545 | 3.981251453 | 6.86E-05 | 0.001976312 | Pkhd1l1 |
| ENSMUSG00000102600.1 | 28.11957953 | -1.758504252 | 0.441900184 | -3.979415072 | 6.91E-05 | 0.00198749 | Gm37266 |
| ENSMUSG00000023473.7 | 34.11851519 | 2.038416428 | 0.512409369 | 3.978101397 | 6.95E-05 | 0.001994345 | Celsr3 |
| ENSMUSG00000041498.9 | 491.5966246 | 0.771600538 | 0.194067977 | 3.975929201 | 7.01E-05 | 0.002008464 | Kif14 |
| ENSMUSG00000089769.1 | 10.93309775 | 2.050404613 | 0.51605523 | 3.973227078 | 7.09E-05 | 0.00202719 | Gm16574 |
| ENSMUSG00000023908.7 | 243.7678883 | 0.904132764 | 0.227594518 | 3.972559492 | 7.11E-05 | 0.002028681 | Pkmyt1 |
| ENSMUSG00000045414.7 | 257.2384077 | -0.936501101 | 0.235788744 | -3.971780346 | 7.13E-05 | 0.002031133 | 1190002N15Rik |
| ENSMUSG00000089872.6 | 136.2876172 | 0.987422285 | 0.248710801 | 3.970162465 | 7.18E-05 | 0.002040767 | Rps6kc1 |
| ENSMUSG00000020642.8 | 150.0209183 | 1.459149759 | 0.367716646 | 3.968136265 | 7.24E-05 | 0.002053966 | Rnf144a |
| ENSMUSG00000020262.11 | 51.15160112 | -1.514698306 | 0.381850265 | -3.966733678 | 7.29E-05 | 0.002061852 | Adarb1 |
| ENSMUSG00000042364.10 | 536.5836891 | 0.849744062 | 0.214410864 | 3.963157678 | 7.40E-05 | 0.002088714 | Snx18 |
| ENSMUSG00000009647.9 | 309.7052947 | 0.749869629 | 0.189338668 | 3.960467442 | 7.48E-05 | 0.002108076 | Mcu |
| ENSMUSG00000057596.9 | 111.0428052 | 1.140565132 | 0.288125106 | 3.958576011 | 7.54E-05 | 0.00212051 | Trim30d |
| ENSMUSG00000026797.11 | 36.28714516 | 1.566153044 | 0.39572369 | 3.957693422 | 7.57E-05 | 0.002124032 | Stxbp1 |
| ENSMUSG00000029925.9 | 28.32808901 | 1.780527779 | 0.450206854 | 3.954910418 | 7.66E-05 | 0.002143054 | Tbxas1 |
| ENSMUSG00000037706.12 | 112.7055621 | 1.664557973 | 0.420917674 | 3.954592731 | 7.67E-05 | 0.002143054 | Cd81 |

| | | | | | | | |
|---|---|---|---|---|---|---|---|
| ENSMUSG00000000355.9 | 178.1700873 | -0.864035819 | 0.21860973 | -3.952412445 | 7.74E-05 | 0.00215592 | Mcts1 |
| ENSMUSG00000073643.7 | 468.2145435 | -0.700744161 | 0.17730506 | -3.95219493 | 7.74E-05 | 0.00215592 | Wdfy1 |
| ENSMUSG00000030786.14 | 634.579772 | 1.683315468 | 0.426027751 | 3.951187371 | 7.78E-05 | 0.002160661 | Itgam |
| ENSMUSG00000058818.9 | 248.1525621 | 1.745446496 | 0.441961349 | 3.949319319 | 7.84E-05 | 0.002173219 | Pirb |
| ENSMUSG00000027669.10 | 159.6315508 | 1.150690015 | 0.291406764 | 3.948741612 | 7.86E-05 | 0.002174104 | Gnb4 |
| ENSMUSG00000072621.9 | 179.083308 | 0.888561155 | 0.225121757 | 3.947024784 | 7.91E-05 | 0.002185367 | Slfn10-ps |
| ENSMUSG00000005824.6 | 223.9292968 | -1.185017871 | 0.300353593 | -3.945409339 | 7.97E-05 | 0.002195762 | Tnfsf14 |
| ENSMUSG00000001911.12 | 45.92851923 | 1.378149076 | 0.349412839 | 3.944185564 | 8.01E-05 | 0.002202608 | Nfix |
| ENSMUSG00000025006.11 | 84.57914462 | 1.157891528 | 0.293820199 | 3.940816642 | 8.12E-05 | 0.002229335 | Sorbs1 |
| ENSMUSG00000073434.7 | 319.2621962 | 0.976870997 | 0.248268186 | 3.934740952 | 8.33E-05 | 0.00228195 | Wdr90 |
| ENSMUSG00000025287.11 | 399.3760114 | -0.949727012 | 0.241451181 | -3.93341217 | 8.37E-05 | 0.002290064 | Acot9 |
| ENSMUSG00000024074.7 | 205.3288071 | 1.610065174 | 0.409566043 | 3.931149083 | 8.45E-05 | 0.002307163 | Crim1 |
| ENSMUSG00000087075.1 | 28.70880279 | 1.894470782 | 0.481990831 | 3.930512075 | 8.48E-05 | 0.002308721 | A230065H16Rik |
| ENSMUSG00000034751.11 | 256.0323569 | -1.228056104 | 0.31249135 | -3.929888315 | 8.50E-05 | 0.002310162 | Mast4 |
| ENSMUSG00000026832.8 | 4465.835579 | -0.753110817 | 0.191686058 | -3.928876331 | 8.53E-05 | 0.002310807 | Cytip |
| ENSMUSG00000034401.12 | 216.4837276 | -1.05395561 | 0.268244793 | -3.929081338 | 8.53E-05 | 0.002310807 | Spata6 |
| ENSMUSG00000042046.11 | 99.2560131 | 1.135966252 | 0.289213737 | 3.927774189 | 8.57E-05 | 0.002316876 | Dstyk |
| ENSMUSG00000044033.12 | 20.87557295 | 1.856420525 | 0.472729847 | 3.927022029 | 8.60E-05 | 0.002319591 | Ccdc141 |
| ENSMUSG00000022892.10 | 272.2291269 | 1.624244939 | 0.414260484 | 3.920830015 | 8.82E-05 | 0.002371093 | App |
| ENSMUSG00000028618.7 | 572.8430794 | -1.005666542 | 0.256495521 | -3.920795717 | 8.83E-05 | 0.002371093 | Tmem59 |
| ENSMUSG00000027508.11 | 1488.322194 | -0.740081147 | 0.188863751 | -3.918598158 | 8.91E-05 | 0.002388165 | Pag1 |
| ENSMUSG00000027360.5 | 197.0930098 | 1.76301672 | 0.450276767 | 3.91540681 | 9.03E-05 | 0.002415288 | Hdc |
| ENSMUSG00000020599.9 | 23.98424041 | 1.626746609 | 0.415798409 | 3.912344479 | 9.14E-05 | 0.002432215 | Rgs9 |
| ENSMUSG00000020898.14 | 238.0613656 | 0.808884118 | 0.206759335 | 3.912201189 | 9.15E-05 | 0.002432215 | Ctc1 |
| ENSMUSG00000025555.10 | 507.6314682 | 1.053335516 | 0.269245705 | 3.912172023 | 9.15E-05 | 0.002432215 | Farp1 |
| ENSMUSG00000034574.9 | 197.8102869 | -0.997931538 | 0.255104267 | -3.911857496 | 9.16E-05 | 0.002432215 | Daam1 |
| ENSMUSG00000037095.7 | 186.0039073 | 1.818156403 | 0.465023723 | 3.909814303 | 9.24E-05 | 0.002448171 | Lrg1 |
| ENSMUSG00000074272.6 | 97.48888079 | 1.675715938 | 0.428670074 | 3.909104083 | 9.26E-05 | 0.002450675 | Ceacam1 |
| ENSMUSG00000046591.9 | 401.5530956 | 0.932601755 | 0.238653864 | 3.907758874 | 9.32E-05 | 0.002459644 | Ticrr |
| ENSMUSG00000046402.10 | 14.81658282 | -1.95522765 | 0.500729098 | -3.904761395 | 9.43E-05 | 0.002485578 | Rbp1 |
| ENSMUSG00000076258.1 | 5716.761617 | 2.01944434 | 0.517310646 | 3.903736287 | 9.47E-05 | 0.00249138 | Gm23935 |
| ENSMUSG00000048612.11 | 110.5466805 | 1.811552417 | 0.464172946 | 3.902753125 | 9.51E-05 | 0.002496769 | Myof |
| ENSMUSG00000073062.3 | 90.88215021 | -1.134515139 | 0.29157737 | -3.89095745 | 9.98E-05 | 0.002616359 | Zxdb |
| ENSMUSG00000021236.12 | 675.9869386 | -0.765873797 | 0.197139858 | -3.884926197 | 0.000102361 | 0.002677089 | Entpd5 |
| ENSMUSG00000024769.7 | 122.8411692 | 1.367962212 | 0.352323241 | 3.882690818 | 0.000103307 | 0.00269672 | Cdc42bpg |
| ENSMUSG00000032398.6 | 112.8849479 | -1.181050976 | 0.304601578 | -3.877363289 | 0.000105595 | 0.002751238 | Snapc5 |
| ENSMUSG00000038884.10 | 148.0633704 | 1.285693774 | 0.33199406 | 3.872640896 | 0.000107662 | 0.002799831 | A230050P20Rik |
| ENSMUSG00000024222.12 | 1650.361298 | 0.67360049 | 0.174034677 | 3.870495828 | 0.000108614 | 0.002819274 | Fkbp5 |
| ENSMUSG00000035578.11 | 26.981652 | -1.769740229 | 0.4573223 | -3.869787743 | 0.00010893 | 0.00282217 | Iqcg |
| ENSMUSG00000070327.9 | 4310.834023 | 0.843903613 | 0.218162571 | 3.868232795 | 0.000109627 | 0.002834904 | Rnf213 |
| ENSMUSG00000038028.8 | 206.8568633 | -0.874879977 | 0.226293013 | -3.866137824 | 0.000110572 | 0.00285401 | 9630033F20Rik |
| ENSMUSG00000027864.8 | 79.06598426 | 1.276051224 | 0.330224521 | 3.86419282 | 0.000111457 | 0.002865393 | Ptgfrn |
| ENSMUSG00000046807.9 | 168.0066758 | -1.615309068 | 0.418032967 | -3.86407101 | 0.000111513 | 0.002865393 | Lrrc75b |
| ENSMUSG00000075591.2 | 24.52902784 | 1.881714389 | 0.487011158 | 3.863801392 | 0.000111636 | 0.002865393 | Gm10874 |
| ENSMUSG00000025648.13 | 93.00708967 | 1.71497087 | 0.444275712 | 3.860149959 | 0.000113317 | 0.002903156 | Pfkfb4 |
| ENSMUSG00000062866.11 | 350.255636 | 1.195195718 | 0.309669371 | 3.859586479 | 0.000113579 | 0.002904469 | Phactr2 |
| ENSMUSG00000070462.4 | 279.3043777 | 0.784350748 | 0.20343037 | 3.855622685 | 0.000115435 | 0.002946484 | Mesdc1 |
| ENSMUSG00000034252.10 | 1476.33411 | -0.672044831 | 0.174376409 | -3.853989405 | 0.000116209 | 0.002960747 | Senp6 |
| ENSMUSG00000038644.10 | 584.9145445 | 0.855318287 | 0.22203304 | 3.852211758 | 0.000117056 | 0.002976837 | Pold1 |
| ENSMUSG00000025464.10 | 74.66295307 | -1.324959912 | 0.344103123 | -3.850473375 | 0.00011789 | 0.002992536 | Paox |
| ENSMUSG00000040537.13 | 38.33677414 | -1.388605758 | 0.360903237 | -3.847584661 | 0.000119288 | 0.003022475 | Adam22 |
| ENSMUSG00000046598.10 | 201.4308447 | -0.834134418 | 0.216840935 | -3.846757158 | 0.000119691 | 0.003027143 | Bdh1 |
| ENSMUSG00000090100.3 | 210.7152257 | -0.85957483 | 0.223539618 | -3.845290768 | 0.00012041 | 0.003039736 | Ttbk2 |
| ENSMUSG00000034206.11 | 341.0041432 | 0.9224209 | 0.239956168 | 3.844122476 | 0.000120985 | 0.003048679 | Polq |
| ENSMUSG00000031530.6 | 202.0044742 | 1.556772483 | 0.405399519 | 3.840094546 | 0.000122987 | 0.003088805 | Dusp4 |
| ENSMUSG00000032184.4 | 131.2619912 | -1.276628318 | 0.332453562 | -3.840019972 | 0.000123024 | 0.003088805 | Lysmd2 |
| ENSMUSG00000028668.5 | 1478.147807 | -0.879963261 | 0.229403949 | -3.835867974 | 0.000125122 | 0.003135761 | Tceb3 |
| ENSMUSG00000042348.9 | 332.3901424 | -0.867016501 | 0.226106545 | -3.834548447 | 0.000125795 | 0.00314693 | Arl15 |
| ENSMUSG00000004709.10 | 68.60756339 | 1.37617969 | 0.358956429 | 3.833834918 | 0.000126161 | 0.00315037 | Cd244 |
| ENSMUSG00000039114.11 | 20.44427398 | 1.756925931 | 0.458490196 | 3.831981459 | 0.000127115 | 0.003168476 | Nrn1 |
| ENSMUSG00000005148.7 | 13.76248136 | 1.94744223 | 0.508316672 | 3.831159467 | 0.000127541 | 0.003173353 | Klf5 |
| ENSMUSG00000002100.11 | 18.19955477 | 2.023910652 | 0.528463955 | 3.829798856 | 0.000128248 | 0.003185211 | Mybpc3 |

| | | | | | | | |
|---|---|---|---|---|---|---|---|
| ENSMUSG00000055639.12 | 51.47166295 | 1.871480349 | 0.488725734 | 3.829305926 | 0.000128505 | 0.003185867 | Dach1 |
| ENSMUSG00000020422.9 | 119.4475637 | 1.755628809 | 0.458691471 | 3.827472103 | 0.000129466 | 0.003203123 | Tns3 |
| ENSMUSG00000026177.7 | 46.11291129 | 1.800513417 | 0.470464936 | 3.82709375 | 0.000129665 | 0.003203123 | Slc11a1 |
| ENSMUSG00000037572.12 | 518.8632932 | 0.700361386 | 0.183254268 | 3.82180122 | 0.00013248 | 0.003266826 | Wdhd1 |
| ENSMUSG00000015981.8 | 425.7288164 | 1.027233098 | 0.268846846 | 3.820885813 | 0.000132973 | 0.003273132 | Stk32c |
| ENSMUSG00000024885.8 | 70.44216095 | 1.827175384 | 0.478444492 | 3.818991368 | 0.000133998 | 0.003292499 | Aldh3b1 |
| ENSMUSG00000029490.3 | 15.07749363 | 1.931485951 | 0.50593136 | 3.817683792 | 0.00013471 | 0.003304114 | Mfsd7a |
| ENSMUSG00000031309.11 | 1257.500153 | -0.88807327 | 0.232780599 | -3.815065657 | 0.000136147 | 0.003333422 | Rps6ka3 |
| ENSMUSG00000059851.11 | 271.2245058 | 0.948023262 | 0.249073953 | 3.806191896 | 0.000141123 | 0.003449144 | Suv420h2 |
| ENSMUSG00000015053.10 | 16.2095184 | 1.894683957 | 0.497920128 | 3.805196558 | 0.000141692 | 0.003456925 | Gata2 |
| ENSMUSG00000004864.8 | 72.86381093 | 1.880744289 | 0.494831608 | 3.800776379 | 0.000144243 | 0.003512976 | Mapk13 |
| ENSMUSG00000063146.7 | 119.0787169 | 1.155954633 | 0.304196676 | 3.800023876 | 0.000144682 | 0.003517457 | Clip2 |
| ENSMUSG00000036067.8 | 58.02658916 | 1.28796325 | 0.339048424 | 3.798758989 | 0.000145422 | 0.003529241 | Slc2a6 |
| ENSMUSG00000000594.7 | 1102.861572 | -1.001238642 | 0.263618909 | -3.798053204 | 0.000145837 | 0.003533094 | Gm2a |
| ENSMUSG00000022637.10 | 1547.793354 | 0.999406997 | 0.263215687 | 3.796912743 | 0.000146509 | 0.003543165 | Cblb |
| ENSMUSG00000028362.2 | 218.3317491 | -1.055127274 | 0.277960394 | -3.795962652 | 0.000147072 | 0.003549054 | Tnfsf8 |
| ENSMUSG00000042272.13 | 198.353464 | -1.289655821 | 0.339773526 | -3.795633624 | 0.000147267 | 0.003549054 | Sestd1 |
| ENSMUSG00000064109.7 | 359.5026755 | -0.958792937 | 0.252699718 | -3.794198688 | 0.000148121 | 0.00356342 | Hcst |
| ENSMUSG00000040616.3 | 14.8667647 | 1.937515178 | 0.51100089 | 3.791608227 | 0.000149675 | 0.0035883 | Tmem51 |
| ENSMUSG00000052415.5 | 8.350506827 | 2.031460661 | 0.535742037 | 3.791863471 | 0.000149521 | 0.0035883 | Tchh |
| ENSMUSG00000104292.1 | 11.80572733 | 2.00622826 | 0.529187893 | 3.791145424 | 0.000149954 | 0.003588763 | Gm38042 |
| ENSMUSG00000019564.8 | 456.1104393 | 0.740237349 | 0.195402756 | 3.788264622 | 0.000151703 | 0.003620594 | Arid3a |
| ENSMUSG00000022686.10 | 363.8146733 | -0.81875557 | 0.216139293 | -3.788092196 | 0.000151809 | 0.003620594 | B3gnt5 |
| ENSMUSG00000003134.6 | 216.219345 | 1.741702356 | 0.460221127 | 3.784490226 | 0.000154024 | 0.003667098 | Tbc1d8 |
| ENSMUSG00000006736.8 | 290.7401049 | -0.872651484 | 0.230691794 | -3.782759099 | 0.0001551 | 0.00368635 | Tspan31 |
| ENSMUSG00000089672.4 | 2341.455016 | -0.940805462 | 0.248852137 | -3.780580204 | 0.000156463 | 0.003712374 | Gp49a |
| ENSMUSG00000022587.10 | 5127.005244 | -0.884844103 | 0.23419068 | -3.778306222 | 0.000157899 | 0.003740004 | Ly6e |
| ENSMUSG00000028843.8 | 1818.108834 | -0.884028928 | 0.234121828 | -3.775935524 | 0.000159408 | 0.003769295 | Sh3bgrl3 |
| ENSMUSG00000027605.14 | 193.3382267 | -1.257976929 | 0.33322765 | -3.775127692 | 0.000159926 | 0.003775067 | Acss2 |
| ENSMUSG00000036499.8 | 781.6537559 | 1.113386395 | 0.295033336 | 3.773764727 | 0.000160802 | 0.003789284 | Eea1 |
| ENSMUSG00000020423.6 | 1809.445041 | -0.834978115 | 0.221313549 | -3.772828719 | 0.000161407 | 0.003797054 | Btg2 |
| ENSMUSG00000020057.2 | 55.07240192 | 1.613259159 | 0.427768533 | 3.771336676 | 0.000162375 | 0.003811755 | Dram1 |
| ENSMUSG00000036875.11 | 318.0791325 | 0.944166371 | 0.250374519 | 3.771016216 | 0.000162584 | 0.003811755 | Dna2 |
| ENSMUSG00000024066.8 | 2361.416791 | 0.772321001 | 0.204893881 | 3.769370738 | 0.00016366 | 0.003819308 | Xdh |
| ENSMUSG00000031441.11 | 845.4100938 | 0.942323467 | 0.249959006 | 3.769912047 | 0.000163305 | 0.003819308 | Atp11a |
| ENSMUSG00000039994.11 | 475.8605883 | 0.866835813 | 0.229975412 | 3.76925431 | 0.000163736 | 0.003819308 | Timeless |
| ENSMUSG00000039960.5 | 61.91515572 | 1.71766261 | 0.455972037 | 3.767034972 | 0.000165198 | 0.00384691 | Rhou |
| ENSMUSG00000022270.11 | 519.6012975 | -1.048676594 | 0.278505912 | -3.765365647 | 0.000166306 | 0.003866184 | Fam134b |
| ENSMUSG00000032691.10 | 52.70296971 | 1.777072146 | 0.472916135 | 3.757689823 | 0.000171489 | 0.003978633 | Nlrp3 |
| ENSMUSG00000032750.9 | 208.0002574 | -0.958882895 | 0.255201573 | -3.757354958 | 0.000171719 | 0.003978633 | Gab3 |
| ENSMUSG00000026535.9 | 16.03549976 | -1.875151399 | 0.499747173 | -3.752200111 | 0.000175289 | 0.00405456 | Ifi202b |
| ENSMUSG00000083061.1 | 115.6027745 | -1.122466154 | 0.299762417 | -3.744519291 | 0.000180739 | 0.004173631 | Gm12191 |
| ENSMUSG00000008318.5 | 250.7352478 | 0.869864202 | 0.232386512 | 3.743178528 | 0.000181707 | 0.004188969 | Relt |
| ENSMUSG00000038665.11 | 29.52438847 | 1.670638098 | 0.446679667 | 3.740125691 | 0.000183928 | 0.004233109 | Dgki |
| ENSMUSG00000066952.7 | 35.02239649 | 1.710690398 | 0.457666625 | 3.737852631 | 0.000185599 | 0.004264446 | Myo1h |
| ENSMUSG00000030867.6 | 293.6762862 | 0.811768776 | 0.21720471 | 3.737344253 | 0.000185974 | 0.004265977 | Plk1 |
| ENSMUSG00000024513.12 | 1666.604758 | -0.685070086 | 0.183347535 | -3.736456477 | 0.000186632 | 0.00427396 | Mbd2 |
| ENSMUSG00000030302.12 | 15.94374598 | 1.998483517 | 0.535191001 | 3.734150077 | 0.00018835 | 0.004299054 | Atp2b2 |
| ENSMUSG00000045636.12 | 142.213056 | 1.761188877 | 0.471592117 | 3.734559617 | 0.000188044 | 0.004299054 | Mtus1 |
| ENSMUSG00000022425.11 | 25.04981803 | 1.6305303 | 0.437784022 | 3.724508479 | 0.000195696 | 0.004452006 | Enpp2 |
| ENSMUSG00000049608.8 | 169.243195 | -1.377229494 | 0.369765341 | -3.724604069 | 0.000195622 | 0.004452006 | Gpr55 |
| ENSMUSG00000046223.9 | 99.22492555 | 1.794893793 | 0.482565765 | 3.719480169 | 0.000199633 | 0.004534104 | Plaur |
| ENSMUSG00000022474.10 | 126.1722888 | 0.871765747 | 0.234412336 | 3.718941431 | 0.000200059 | 0.004536323 | Pmm1 |
| ENSMUSG00000046245.9 | 75.90383124 | 1.647892997 | 0.443273182 | 3.717556267 | 0.000201159 | 0.004553782 | Pilra |
| ENSMUSG00000038252.9 | 1861.26031 | 0.837516707 | 0.225353851 | 3.716451715 | 0.00020204 | 0.004566241 | Ncapd2 |
| ENSMUSG00000025558.11 | 524.8056423 | 1.061688632 | 0.286325786 | 3.70797421 | 0.000208924 | 0.004714102 | Dock9 |
| ENSMUSG00000064373.7 | 248.6176395 | -1.446247187 | 0.390291181 | -3.705559486 | 0.000210925 | 0.004751481 | Sepp1 |
| ENSMUSG00000008305.14 | 34.1044793 | -1.490663196 | 0.402562022 | -3.702940451 | 0.000213115 | 0.004793003 | Tle1 |
| ENSMUSG00000045502.5 | 21.85494025 | 1.93249395 | 0.522395084 | 3.699295817 | 0.000216198 | 0.004854447 | Hcar2 |
| ENSMUSG00000029455.10 | 246.462381 | 1.646924442 | 0.445350951 | 3.698037332 | 0.000217273 | 0.004870653 | Aldh2 |
| ENSMUSG00000050965.10 | 597.1340187 | 0.707311864 | 0.191341555 | 3.696593057 | 0.000218512 | 0.004890494 | Prkca |
| ENSMUSG00000029840.5 | 1986.139169 | -1.098627737 | 0.297942135 | -3.687386266 | 0.000226569 | 0.005062614 | Mtpn |

| | | | | | | | |
|---|---|---|---|---|---|---|---|
| ENSMUSG00000023067.9 | 196.2572566 | 1.064999191 | 0.288924811 | 3.686077314 | 0.000227737 | 0.005080489 | Cdkn1a |
| ENSMUSG00000038058.10 | 1006.813774 | 1.102410089 | 0.29912731 | 3.68542107 | 0.000228325 | 0.005085383 | Nod1 |
| ENSMUSG00000022971.14 | 293.7183015 | 0.850774974 | 0.231054658 | 3.682137303 | 0.000231287 | 0.00513479 | Ifnar2 |
| ENSMUSG00000068587.6 | 86.95625675 | 1.777164378 | 0.482635491 | 3.682208235 | 0.000231222 | 0.00513479 | Mgam |
| ENSMUSG00000068758.7 | 95.98360668 | 0.97664586 | 0.265568674 | 3.677564243 | 0.000235472 | 0.00521931 | Il3ra |
| ENSMUSG00000093843.1 | 5202.506681 | -1.215662997 | 0.330673482 | -3.676324425 | 0.000236619 | 0.005236324 | Gm25939 |
| ENSMUSG00000000555.6 | 41.91635664 | 1.756084542 | 0.478175686 | 3.672467244 | 0.00024022 | 0.005307516 | Itga5 |
| ENSMUSG00000020185.12 | 320.5456162 | 0.861569333 | 0.234648763 | 3.671740366 | 0.000240904 | 0.005314135 | E2f7 |
| ENSMUSG00000034957.9 | 45.19015328 | 1.752139201 | 0.477356925 | 3.67050127 | 0.000242075 | 0.005331447 | Cebpa |
| ENSMUSG00000037020.12 | 176.0752487 | 1.060817657 | 0.289195704 | 3.668165344 | 0.000244297 | 0.005371815 | Wdr62 |
| ENSMUSG00000000732.8 | 268.5845877 | -0.791305265 | 0.215791464 | -3.666990571 | 0.000245422 | 0.005373436 | Icosl |
| ENSMUSG00000016239.7 | 12.00889737 | 1.805682384 | 0.492431628 | 3.666869226 | 0.000245538 | 0.005373436 | Lonrf3 |
| ENSMUSG00000036622.11 | 267.1682745 | 0.844687929 | 0.230340459 | 3.667127926 | 0.00024529 | 0.005373436 | Atp13a2 |
| ENSMUSG00000006179.4 | 103.796217 | -1.154023737 | 0.314961821 | -3.664011511 | 0.000248296 | 0.005401673 | Prss16 |
| ENSMUSG00000028245.11 | 2951.657731 | 0.828326532 | 0.226045061 | 3.664431018 | 0.000247889 | 0.005401673 | Nsmaf |
| ENSMUSG00000034656.12 | 41.63901036 | 1.407915467 | 0.384204233 | 3.664497542 | 0.000247825 | 0.005401673 | Cacna1a |
| ENSMUSG00000049130.5 | 74.5757234 | 1.693579575 | 0.462232743 | 3.663910875 | 0.000248393 | 0.005401673 | C5ar1 |
| ENSMUSG00000056069.8 | 710.3069972 | -0.679282796 | 0.185464376 | -3.662605249 | 0.000249663 | 0.005420752 | Fam105a |
| ENSMUSG00000032281.7 | 1205.207872 | -0.883348355 | 0.241219857 | -3.662005129 | 0.000250249 | 0.005424939 | Acsbg1 |
| ENSMUSG00000073420.6 | 14.53537023 | 1.819767465 | 0.497144393 | 3.660440487 | 0.000251782 | 0.005449621 | Btnl5-ps |
| ENSMUSG00000031264.9 | 45.7658371 | 1.783651317 | 0.487482607 | 3.658902474 | 0.000253298 | 0.0054729 | Btk |
| ENSMUSG00000044629.5 | 14.01437365 | 1.836883184 | 0.502080115 | 3.658545978 | 0.00025365 | 0.0054729 | Cnrip1 |
| ENSMUSG00000097357.1 | 9.315763544 | -1.936330069 | 0.529355431 | -3.657901585 | 0.000254289 | 0.005478115 | Gm16793 |
| ENSMUSG00000021701.7 | 68.62021396 | 1.300349788 | 0.355846075 | 3.654247939 | 0.000257937 | 0.005548054 | Plk2 |
| ENSMUSG00000020042.11 | 403.1050982 | 1.201076828 | 0.328863562 | 3.652204035 | 0.000259999 | 0.005581583 | Btbd11 |
| ENSMUSG00000022015.8 | 61.05183993 | -1.267756794 | 0.347149626 | -3.651903091 | 0.000260304 | 0.005581583 | Tnfsf11 |
| ENSMUSG00000024669.7 | 2080.556149 | -0.693620227 | 0.190097482 | -3.648760728 | 0.000263508 | 0.005641531 | Cd5 |
| ENSMUSG00000039477.12 | 893.9403346 | 1.100807444 | 0.301746587 | 3.648118957 | 0.000264167 | 0.005646883 | Tnrc18 |
| ENSMUSG00000019866.9 | 2326.975746 | -0.836877462 | 0.229531584 | -3.646023119 | 0.00026633 | 0.005684314 | Aim1 |
| ENSMUSG00000038831.12 | 79.70179556 | 1.227160546 | 0.336873434 | 3.642794064 | 0.000269695 | 0.005747242 | Ralgps1 |
| ENSMUSG00000047766.11 | 54.6772072 | -1.160897487 | 0.318886693 | -3.640470146 | 0.000272141 | 0.005790432 | Lrrc49 |
| ENSMUSG00000023809.9 | 53.20246621 | 1.317307412 | 0.361960761 | 3.63936524 | 0.000273311 | 0.005804509 | Rps6ka2 |
| ENSMUSG00000031902.9 | 3820.970468 | -0.680908984 | 0.187128197 | -3.63872999 | 0.000273986 | 0.005804509 | Nfatc3 |
| ENSMUSG00000058587.7 | 1290.12276 | -0.846867909 | 0.232741876 | -3.638657219 | 0.000274063 | 0.005804509 | Tmod3 |
| ENSMUSG00000038173.10 | 13.94405704 | 1.890558091 | 0.519761693 | 3.637355576 | 0.000275452 | 0.005824978 | Enpp6 |
| ENSMUSG00000004032.6 | 24.3454754 | -1.649748571 | 0.453650347 | -3.636608199 | 0.000276252 | 0.005832965 | Gstm5 |
| ENSMUSG00000044345.9 | 294.0016038 | -1.033809185 | 0.284422096 | -3.634771003 | 0.000278228 | 0.005865719 | Marveld1 |
| ENSMUSG00000022351.10 | 261.4643048 | -0.887539076 | 0.244328453 | -3.632565368 | 0.000280617 | 0.005907083 | Sqle |
| ENSMUSG00000020846.6 | 246.7229214 | 1.393210682 | 0.383738899 | 3.630621457 | 0.00028274 | 0.005942696 | Fam101b |
| ENSMUSG00000052713.8 | 36.32977633 | 1.735251028 | 0.478379099 | 3.627355445 | 0.000286339 | 0.006009202 | Zfp608 |
| ENSMUSG00000004846.6 | 138.5408836 | 0.945707251 | 0.260743618 | 3.626962221 | 0.000286775 | 0.006009225 | Plod3 |
| ENSMUSG00000022636.9 | 280.3844864 | 1.100830628 | 0.303617637 | 3.625713703 | 0.000288164 | 0.006029187 | Alcam |
| ENSMUSG00000019734.12 | 39.86695428 | 1.536137782 | 0.423965175 | 3.623264065 | 0.000290909 | 0.006077392 | Tmc4 |
| ENSMUSG00000021596.12 | 72.73031921 | 1.69293033 | 0.467317963 | 3.622651952 | 0.000291598 | 0.006082595 | Mctp1 |
| ENSMUSG00000030339.7 | 51.35354513 | 1.794656204 | 0.49625041 | 3.616432689 | 0.000298691 | 0.006216684 | Ltbr |
| ENSMUSG00000031642.8 | 117.9223648 | 1.056784495 | 0.292233892 | 3.616228388 | 0.000298927 | 0.006216684 | Sh3rf1 |
| ENSMUSG00000031066.6 | 83.01801729 | 1.29289249 | 0.357580231 | 3.615671052 | 0.000299571 | 0.006220707 | Usp11 |
| ENSMUSG00000024187.10 | 166.8886394 | 1.004331221 | 0.277813367 | 3.61512922 | 0.000300198 | 0.006224372 | Itfg3 |
| ENSMUSG00000051278.8 | 349.5932845 | 0.819509085 | 0.226964969 | 3.610729397 | 0.000305337 | 0.006321439 | Zgrf1 |
| ENSMUSG00000030530.11 | 5473.961895 | -0.951886428 | 0.263793199 | -3.608457047 | 0.000308023 | 0.00636751 | Furin |
| ENSMUSG00000025701.8 | 117.1411686 | 1.690961246 | 0.468880572 | 3.606379425 | 0.000310499 | 0.00640909 | Alox5 |
| ENSMUSG00000008683.12 | 2841.521643 | -1.116456171 | 0.309679097 | -3.605203522 | 0.000311908 | 0.006424725 | Rps15a |
| ENSMUSG00000036291.5 | 284.6610371 | -0.770033957 | 0.213603333 | -3.60497164 | 0.000312187 | 0.006424725 | Ap5m1 |
| ENSMUSG00000004612.8 | 2152.934724 | 0.592340702 | 0.16438325 | 3.603412762 | 0.000314066 | 0.00645378 | Nkg7 |
| ENSMUSG00000051412.6 | 170.3963645 | -1.065604215 | 0.2957911 | -3.602556716 | 0.000315103 | 0.006465456 | Vamp7 |
| ENSMUSG00000022965.7 | 55.55629239 | 1.598847334 | 0.444133812 | 3.59992257 | 0.000318312 | 0.006515523 | Ifngr2 |
| ENSMUSG00000103869.1 | 167.6223068 | 1.242317922 | 0.345109389 | 3.599780129 | 0.000318486 | 0.006515523 | Gm37420 |
| ENSMUSG00000020715.5 | 982.3941736 | -0.909330724 | 0.252840586 | -3.596458694 | 0.000322579 | 0.006587706 | Ern1 |
| ENSMUSG00000058325.5 | 38.15243631 | 1.763312347 | 0.490334167 | 3.596144155 | 0.000322969 | 0.006587706 | Dock1 |
| ENSMUSG00000042064.9 | 109.5997529 | 1.266924174 | 0.352367175 | 3.595465939 | 0.000323811 | 0.006595152 | Myo3b |
| ENSMUSG00000035227.6 | 678.291521 | -0.946348823 | 0.263415728 | -3.592605618 | 0.000327388 | 0.006658173 | Spcs2 |
| ENSMUSG00000052572.11 | 20.33371685 | 1.680615363 | 0.467952668 | 3.591421697 | 0.000328879 | 0.006678662 | Dlg2 |

| | | | | | | | |
|---|---|---|---|---|---|---|---|
| ENSMUSG00000057378.10 | 56.04743922 | 1.175709371 | 0.327505125 | 3.5898961 | 0.00033081 | 0.006708007 | Ryr3 |
| ENSMUSG00000023274.10 | 5716.845455 | -0.738095107 | 0.205630096 | -3.589431316 | 0.0003314 | 0.006710124 | Cd4 |
| ENSMUSG00000020607.6 | 7.733518398 | 1.924131945 | 0.536626552 | 3.585607046 | 0.000336295 | 0.006799265 | Fam84a |
| ENSMUSG00000042745.9 | 23.75842806 | 1.859944846 | 0.51878995 | 3.585159749 | 0.000336872 | 0.006800972 | Id1 |
| ENSMUSG00000030747.4 | 47.72259507 | 1.78405838 | 0.497850673 | 3.583521078 | 0.000338993 | 0.00683381 | Dgat2 |
| ENSMUSG00000000127.10 | 12.61243851 | 1.722227785 | 0.480790128 | 3.582078095 | 0.000340872 | 0.006853654 | Fer |
| ENSMUSG00000048440.11 | 65.98750684 | -1.136694658 | 0.317334962 | -3.5820026 | 0.00034097 | 0.006853654 | Cyp4f16 |
| ENSMUSG00000021322.7 | 59.97167637 | 1.676613767 | 0.468390137 | 3.579524064 | 0.000344221 | 0.006908927 | Aoah |
| ENSMUSG00000056529.7 | 55.01756497 | 1.503968053 | 0.42043482 | 3.577172923 | 0.00034733 | 0.006961227 | Ptafr |
| ENSMUSG00000040488.12 | 32.32070051 | 1.723113802 | 0.481801703 | 3.576396247 | 0.000348363 | 0.006971814 | Ltbp4 |
| ENSMUSG00000031732.8 | 235.1448324 | 0.909312279 | 0.254420862 | 3.574047638 | 0.000351505 | 0.006996274 | Phlpp2 |
| ENSMUSG00000040264.9 | 406.4881092 | 1.019015247 | 0.285121558 | 3.573967726 | 0.000351612 | 0.006996274 | Gbp2b |
| ENSMUSG00000048264.11 | 62.38865913 | -1.353054033 | 0.378553658 | -3.57427277 | 0.000351203 | 0.006996274 | Dip2c |
| ENSMUSG00000071637.4 | 44.12029824 | 1.467904429 | 0.410615293 | 3.574889814 | 0.000350375 | 0.006996274 | Cebpd |
| ENSMUSG00000040327.12 | 68.56451575 | 1.272354238 | 0.356182318 | 3.572199333 | 0.000353996 | 0.007033565 | Cul9 |
| ENSMUSG00000024670.12 | 2970.736223 | -0.60780287 | 0.170212887 | -3.570839325 | 0.000355839 | 0.007039803 | Cd6 |
| ENSMUSG00000047250.9 | 70.63824795 | 1.665752768 | 0.466423211 | 3.571333344 | 0.000355169 | 0.007039803 | Ptgs1 |
| ENSMUSG00000048897.11 | 185.6898144 | 1.071581719 | 0.300083395 | 3.570946404 | 0.000355694 | 0.007039803 | Zfp710 |
| ENSMUSG00000019235.8 | 10.46150882 | 1.796085979 | 0.503190978 | 3.569392257 | 0.00035781 | 0.007068674 | Rps6kl1 |
| ENSMUSG00000098161.1 | 9.020688201 | 1.900346587 | 0.532650155 | 3.567719957 | 0.000360101 | 0.007103765 | Gm26975 |
| ENSMUSG00000034557.10 | 16.52747675 | 1.82991876 | 0.512974675 | 3.567269203 | 0.000360721 | 0.007105841 | Zfyve9 |
| ENSMUSG00000048249.10 | 808.1345666 | -0.687074342 | 0.192627495 | -3.566854993 | 0.000361291 | 0.007106939 | Crebrf |
| ENSMUSG00000028717.8 | 29.19324563 | 1.79239493 | 0.502885137 | 3.564223317 | 0.000364935 | 0.007168406 | Tal1 |
| ENSMUSG00000054889.5 | 19.46109848 | 1.586980946 | 0.445318248 | 3.563700686 | 0.000365663 | 0.007172498 | Dsp |
| ENSMUSG00000006567.7 | 14.21251706 | 1.82621455 | 0.513072547 | 3.559369059 | 0.000371747 | 0.007259382 | Atp7b |
| ENSMUSG00000024590.8 | 2203.980433 | 0.664451576 | 0.186664851 | 3.559596634 | 0.000371425 | 0.007259382 | Lmnb1 |
| ENSMUSG00000024910.4 | 1260.852502 | -0.98465638 | 0.276662505 | -3.559052503 | 0.000372195 | 0.007259382 | Ctsw |
| ENSMUSG00000086583.3 | 320.1100536 | -0.842595916 | 0.236713009 | -3.559567424 | 0.000371466 | 0.007259382 | Gm15500 |
| ENSMUSG00000024548.11 | 95.58108488 | 1.528577666 | 0.429903653 | 3.555628467 | 0.000377077 | 0.007344219 | Setbp1 |
| ENSMUSG00000048965.8 | 40.08066841 | 1.351547384 | 0.380250221 | 3.554363179 | 0.000378896 | 0.007369253 | Mrgpre |
| ENSMUSG00000027293.9 | 287.0669363 | 0.78249263 | 0.220214636 | 3.553318001 | 0.000380404 | 0.00738819 | Ehd4 |
| ENSMUSG00000097451.5 | 12.16767558 | 1.892366778 | 0.53271775 | 3.552287826 | 0.000381897 | 0.007406761 | Rian |
| ENSMUSG00000100150.1 | 217.131625 | -0.901703276 | 0.253914368 | -3.551210134 | 0.000383464 | 0.007426727 | Gm19585 |
| ENSMUSG00000073155.8 | 245.8952543 | -0.937593288 | 0.264058684 | -3.550700449 | 0.000384207 | 0.007430701 | 1810058I24Rik |
| ENSMUSG00000042043.5 | 210.4816633 | -1.039966572 | 0.292950906 | -3.54996878 | 0.000385277 | 0.007440962 | Tbca |
| ENSMUSG00000024677.9 | 4139.169532 | -0.754074939 | 0.2124898 | -3.548758292 | 0.000387052 | 0.00746481 | Ms4a6b |
| ENSMUSG00000000394.11 | 20.19536885 | 1.838414372 | 0.518273978 | 3.547186336 | 0.000389369 | 0.007499021 | Gcg |
| ENSMUSG00000049999.4 | 27.58637439 | 1.837970389 | 0.518335057 | 3.545911789 | 0.000391257 | 0.007523988 | Ppp1r3d |
| ENSMUSG00000055866.9 | 32.86378987 | 1.536022072 | 0.433222087 | 3.545576551 | 0.000391755 | 0.007523988 | Per2 |
| ENSMUSG00000009210.6 | 5.465592974 | -1.8993808 | 0.535873302 | -3.544458727 | 0.00039342 | 0.00754547 | 2310007L24Rik |
| ENSMUSG00000030187.11 | 57.95107348 | 1.599254433 | 0.451360707 | 3.543184881 | 0.000395325 | 0.007571497 | Klra2 |
| ENSMUSG00000048329.7 | 10.43993978 | 1.899020082 | 0.536519361 | 3.539518271 | 0.000400858 | 0.007666826 | Mfsd6l |
| ENSMUSG00000031278.8 | 1015.111575 | -0.901988657 | 0.254960614 | -3.537756842 | 0.000403541 | 0.007686213 | Acsl4 |
| ENSMUSG00000033306.10 | 765.0399824 | -0.714339177 | 0.201900418 | -3.538076759 | 0.000403053 | 0.007686213 | Lpp |
| ENSMUSG00000052504.6 | 14.72901264 | 1.794255103 | 0.507131155 | 3.538049445 | 0.000403095 | 0.007686213 | Epha3 |
| ENSMUSG00000033192.4 | 68.57384888 | 1.691165576 | 0.478746519 | 3.532486418 | 0.000411671 | 0.007830262 | Lpcat2 |
| ENSMUSG00000025348.8 | 58.44070049 | 1.229094236 | 0.34806594 | 3.531210885 | 0.000413662 | 0.007857298 | Itga7 |
| ENSMUSG00000024897.8 | 23.71567591 | 1.745135195 | 0.494562146 | 3.528646921 | 0.00041769 | 0.007922914 | Apba1 |
| ENSMUSG00000037902.14 | 329.9062813 | 1.55930429 | 0.442245591 | 3.525878657 | 0.00042208 | 0.007995209 | Sirpa |
| ENSMUSG00000036606.12 | 59.46679499 | 1.590574339 | 0.451171226 | 3.525433909 | 0.00042279 | 0.007997675 | Plxnb2 |
| ENSMUSG00000024660.8 | 984.2033979 | 0.598810254 | 0.169906362 | 3.524354513 | 0.000424516 | 0.008013691 | Incenp |
| ENSMUSG00000026579.8 | 104.5826261 | 1.679789493 | 0.476647026 | 3.524179115 | 0.000424797 | 0.008013691 | F5 |
| ENSMUSG00000026782.11 | 312.2120282 | -0.870757532 | 0.247179028 | -3.522780799 | 0.000427044 | 0.008045097 | Abi2 |
| ENSMUSG00000039286.8 | 50.25756928 | 1.481679024 | 0.420716119 | 3.521802368 | 0.000428624 | 0.008063846 | Fndc3b |
| ENSMUSG00000075602.6 | 2433.148526 | -0.9724395 | 0.276315886 | -3.519303621 | 0.000432681 | 0.008129109 | Ly6a |
| ENSMUSG00000029313.14 | 899.6080327 | 0.620244097 | 0.176304755 | 3.518022503 | 0.000434775 | 0.008157357 | Aff1 |
| ENSMUSG00000031231.4 | 412.6212929 | -1.112621938 | 0.317210763 | -3.507516352 | 0.000452311 | 0.008474839 | Cox7b |
| ENSMUSG00000051344.9 | 367.6433614 | -0.70298446 | 0.200607362 | -3.504280475 | 0.000457843 | 0.00855528 | Plekhm3 |
| ENSMUSG00000076498.2 | 2425.986164 | -0.854829063 | 0.24393191 | -3.50437572 | 0.000457679 | 0.00855528 | Trbc2 |
| ENSMUSG00000032198.8 | 51.34360878 | 1.715003095 | 0.48974958 | 3.50179595 | 0.000462133 | 0.008623784 | Dock6 |
| ENSMUSG00000028556.11 | 26.01044675 | 1.662694009 | 0.47490463 | 3.501111391 | 0.000463322 | 0.008634299 | Dock7 |
| ENSMUSG00000036299.4 | 219.0084978 | -0.917751912 | 0.26229214 | -3.498968402 | 0.000467062 | 0.008681043 | BC031181 |

| | | | | | | | |
|---|---|---|---|---|---|---|---|
| ENSMUSG00000042265.9 | 79.1857704 | 1.753905955 | 0.501265843 | 3.498953661 | 0.000467088 | 0.008681043 | Trem1 |
| ENSMUSG00000103734.1 | 23.63736601 | 1.602794804 | 0.458252221 | 3.497625828 | 0.000469419 | 0.008712648 | Gm37651 |
| ENSMUSG00000041324.9 | 20.45491831 | 1.696911366 | 0.485349435 | 3.496267315 | 0.000471816 | 0.008745375 | Inhba |
| ENSMUSG00000021375.9 | 41.36826368 | 1.556705716 | 0.44549543 | 3.49432477 | 0.000475262 | 0.008785676 | Kif13a |
| ENSMUSG00000070780.7 | 335.0513738 | 0.793139659 | 0.226971074 | 3.494452592 | 0.000475035 | 0.008785676 | Rbm47 |
| ENSMUSG00000001227.7 | 27.21657675 | 1.634390896 | 0.467968922 | 3.492520164 | 0.000478485 | 0.00883343 | Sema6b |
| ENSMUSG00000039109.11 | 220.5015096 | 1.637229022 | 0.468934279 | 3.491382688 | 0.000480527 | 0.008859284 | F13a1 |
| ENSMUSG00000019838.10 | 38.4398707 | 1.629702263 | 0.467235593 | 3.48796686 | 0.000486708 | 0.008961275 | Slc16a10 |
| ENSMUSG00000062456.3 | 81.65180911 | -1.26767578 | 0.363676489 | -3.485723767 | 0.000490808 | 0.009024715 | Rpl9-ps6 |
| ENSMUSG00000020641.11 | 62.63393176 | 1.337095212 | 0.383717613 | 3.484581287 | 0.000492908 | 0.00905128 | Rsad2 |
| ENSMUSG00000046743.6 | 10.51742177 | 1.80162547 | 0.517099288 | 3.48409969 | 0.000493796 | 0.009055541 | Fat4 |
| ENSMUSG00000019943.9 | 2787.986291 | -0.908644938 | 0.260838525 | -3.483553433 | 0.000494804 | 0.009062007 | Atp2b1 |
| ENSMUSG00000040653.5 | 5.887012988 | -1.861899328 | 0.535028872 | -3.479997855 | 0.000501418 | 0.009170965 | Ppp1r14c |
| ENSMUSG00000007080.10 | 774.4181328 | 0.780106668 | 0.224217036 | 3.479247972 | 0.000502823 | 0.009184503 | Pole |
| ENSMUSG00000026158.7 | 112.8837257 | 1.445163634 | 0.415587159 | 3.477402036 | 0.000506298 | 0.009211422 | Ogfrl1 |
| ENSMUSG00000032122.10 | 84.24568687 | 1.12724434 | 0.32416278 | 3.477402123 | 0.000506298 | 0.009211422 | Slc37a2 |
| ENSMUSG00000070923.4 | 635.816243 | -0.717986258 | 0.20645351 | -3.477713975 | 0.000505709 | 0.009211422 | Klhl9 |
| ENSMUSG00000020085.11 | 39.54728558 | 1.15345307 | 0.331737593 | 3.477004399 | 0.000507049 | 0.009212956 | Aifm2 |
| ENSMUSG00000036473.11 | 35.94179018 | 1.270884047 | 0.36569107 | 3.475294176 | 0.000510293 | 0.009259713 | Tbc1d24 |
| ENSMUSG00000103747.1 | 30.00187242 | 1.552984599 | 0.446978714 | 3.474403927 | 0.00051199 | 0.009278301 | Gm38236 |
| ENSMUSG00000037375.12 | 37.32085341 | 1.404580734 | 0.4043534 | 3.473646405 | 0.000513437 | 0.009292338 | Hhat |
| ENSMUSG00000071552.4 | 529.0070059 | 1.212133032 | 0.348992885 | 3.473231353 | 0.000514232 | 0.009294539 | Tigit |
| ENSMUSG00000008682.9 | 2425.71078 | -0.870322575 | 0.250995118 | -3.467488062 | 0.000525347 | 0.00948303 | Rpl10 |
| ENSMUSG00000032806.9 | 75.32051199 | -1.182636053 | 0.341206163 | -3.466045408 | 0.000528174 | 0.009521613 | Slc10a3 |
| ENSMUSG00000032328.8 | 1161.68317 | -0.744754011 | 0.214909153 | -3.465436448 | 0.000529372 | 0.009523665 | Tmem30a |
| ENSMUSG00000054843.8 | 619.4019422 | -0.745637429 | 0.215173397 | -3.465286322 | 0.000529667 | 0.009523665 | Atrnl1 |
| ENSMUSG00000084803.4 | 38.36184512 | -1.544147586 | 0.445758413 | -3.464090726 | 0.000532027 | 0.009553656 | 5830444B04Rik |
| ENSMUSG00000017499.11 | 347.4549488 | 0.734571923 | 0.212186709 | 3.461912981 | 0.000536351 | 0.009618785 | Cdc6 |
| ENSMUSG00000029213.7 | 492.0582679 | -0.969813374 | 0.280359519 | -3.459177625 | 0.000541827 | 0.009704404 | Commd8 |
| ENSMUSG00000018168.8 | 3103.543844 | 1.080675893 | 0.312449785 | 3.458718633 | 0.000542752 | 0.009708363 | Ikzf3 |
| ENSMUSG00000028252.16 | 326.2435553 | -0.860701524 | 0.24907177 | -3.455636598 | 0.000548995 | 0.009794666 | Ccnc |
| ENSMUSG00000079641.3 | 1171.430669 | -1.241055263 | 0.359123406 | -3.455790521 | 0.000548682 | 0.009794666 | Rpl39 |
| ENSMUSG00000047557.2 | 92.74863499 | -1.260468782 | 0.364861715 | -3.454647966 | 0.000551012 | 0.009817963 | Lxn |
| ENSMUSG00000040524.9 | 218.1868021 | 0.956414608 | 0.277184823 | 3.450458059 | 0.000559636 | 0.009958782 | Zfp609 |
| ENSMUSG00000013236.12 | 243.9020184 | 0.931030007 | 0.270260329 | 3.444937739 | 0.000571191 | 0.010135817 | Ptprs |
| ENSMUSG00000055733.6 | 6.550593502 | 1.825290504 | 0.529890509 | 3.444655971 | 0.000571787 | 0.010135817 | Nap1l3 |
| ENSMUSG00000078974.6 | 235.206283 | -1.22419735 | 0.35533656 | -3.445177019 | 0.000570686 | 0.010135817 | Sec61g |
| ENSMUSG00000059495.9 | 391.1191945 | -0.685507366 | 0.199047639 | -3.443936189 | 0.000573311 | 0.01014981 | Arhgef12 |
| ENSMUSG00000016758.3 | 72.36926556 | -1.364101917 | 0.396202485 | -3.442941349 | 0.000575424 | 0.010174176 | Bik |
| ENSMUSG00000032548.10 | 46.1924626 | 1.798905014 | 0.523212296 | 3.438193304 | 0.000585609 | 0.010341025 | Slco2a1 |
| ENSMUSG00000042377.8 | 16.82956249 | 1.494920337 | 0.434874812 | 3.437587772 | 0.00058692 | 0.010350938 | Fam83g |
| ENSMUSG00000017760.11 | 1551.228163 | -0.614241024 | 0.178735487 | -3.436592448 | 0.000589081 | 0.010369453 | Ctsa |
| ENSMUSG00000017774.15 | 473.7252038 | 0.680960413 | 0.198160249 | 3.436412777 | 0.000589472 | 0.010369453 | Myo1c |
| ENSMUSG00000037685.11 | 564.7585502 | 0.727754943 | 0.211838156 | 3.435428993 | 0.000591616 | 0.010393934 | Atp8a1 |
| ENSMUSG00000013974.2 | 111.8866353 | 1.572919085 | 0.45810984 | 3.433497708 | 0.000595847 | 0.010454961 | Mcemp1 |
| ENSMUSG00000049866.8 | 2566.172462 | -0.990429769 | 0.28855331 | -3.432397873 | 0.000598269 | 0.010484136 | Arl4c |
| ENSMUSG00000027009.14 | 13110.2772 | 0.678693739 | 0.197834486 | 3.430613908 | 0.000602217 | 0.010539944 | Itga4 |
| ENSMUSG00000055170.3 | 1111.216826 | -0.977881609 | 0.285278137 | -3.427818269 | 0.000608453 | 0.010635598 | Ifng |
| ENSMUSG00000046311.9 | 808.8445379 | -0.709947459 | 0.207189855 | -3.426555114 | 0.00061129 | 0.010671681 | Zfp62 |
| ENSMUSG00000059325.10 | 627.3037102 | -0.746066961 | 0.217762407 | -3.426059484 | 0.000612406 | 0.010677674 | Hopx |
| ENSMUSG00000029142.10 | 492.8188363 | -0.67833208 | 0.198249387 | -3.421609975 | 0.000622515 | 0.010840247 | ENSMUSG00000029142 |
| ENSMUSG00000009927.8 | 1819.366446 | -1.034429268 | 0.302408722 | -3.42063304 | 0.000624756 | 0.010851889 | Rps25 |
| ENSMUSG00000054934.6 | 59.54607787 | -1.331464831 | 0.389237269 | -3.420702321 | 0.000624597 | 0.010851889 | Kcnmb4 |
| ENSMUSG00000022450.5 | 241.9447894 | -0.848777807 | 0.248191171 | -3.419854955 | 0.000626545 | 0.010855664 | Ndufa6 |
| ENSMUSG00000048232.8 | 32.19382596 | 1.317900965 | 0.385361058 | 3.419912153 | 0.000626414 | 0.010855664 | Fbxo10 |
| ENSMUSG00000025993.6 | 105.26341 | 1.446356883 | 0.423084061 | 3.418604049 | 0.000629432 | 0.010892021 | Slc40a1 |
| ENSMUSG00000022500.10 | 320.4191391 | 0.997120604 | 0.291747298 | 3.417754373 | 0.000631401 | 0.010904399 | Litaf |
| ENSMUSG00000076462.2 | 87.82279988 | -1.575485045 | 0.4609898 | -3.417613672 | 0.000631727 | 0.010904399 | Trbv2 |
| ENSMUSG00000028648.9 | 80.23823123 | -1.210164504 | 0.354324401 | -3.415413959 | 0.000636851 | 0.010979128 | Ndufs5 |
| ENSMUSG00000023008.14 | 339.9856624 | 0.738369419 | 0.216300447 | 3.41362872 | 0.000641039 | 0.011037535 | Fmnl3 |
| ENSMUSG00000027133.3 | 139.6136634 | -1.107921708 | 0.324609535 | -3.413090468 | 0.000642306 | 0.011045586 | Nop10 |
| ENSMUSG00000009772.11 | 38.96740312 | 1.529405581 | 0.448276336 | 3.411747308 | 0.000645479 | 0.011084458 | Nuak2 |

| | | | | | | | |
|---|---|---|---|---|---|---|---|
| ENSMUSG00000035596.10 | 343.8549618 | 0.730268852 | 0.214063759 | 3.411454872 | 0.000646172 | 0.011084458 | Mboat7 |
| ENSMUSG00000024387.9 | 741.8259716 | -0.854571215 | 0.250529899 | -3.411054807 | 0.000647121 | 0.011086962 | Csnk2b |
| ENSMUSG00000062991.6 | 8.462771183 | 1.811771182 | 0.531542858 | 3.408513831 | 0.000653178 | 0.011176867 | Nrg1 |
| ENSMUSG00000021756.8 | 323.5729364 | 0.912020332 | 0.267691203 | 3.406986572 | 0.000656844 | 0.011225684 | Il6st |
| ENSMUSG00000101249.1 | 8.462783568 | -1.786029917 | 0.524358317 | -3.406124894 | 0.00065892 | 0.011247256 | Gm29216 |
| ENSMUSG00000037466.9 | 261.7296491 | 0.795182726 | 0.233584353 | 3.404263674 | 0.000663427 | 0.011310198 | 4930427A07Rik |
| ENSMUSG00000072214.6 | 78.7197399 | 1.379808187 | 0.405400442 | 3.403568535 | 0.000665117 | 0.011325036 | Sept5 |
| ENSMUSG00000090942.1 | 57.67096762 | 1.378494729 | 0.40521204 | 3.401909598 | 0.000669168 | 0.011379971 | F830016B08Rik |
| ENSMUSG00000079553.6 | 294.183264 | 0.949105522 | 0.279237573 | 3.398917674 | 0.000676531 | 0.011491037 | Kifc1 |
| ENSMUSG00000020250.9 | 1373.461182 | -0.575509868 | 0.169396228 | -3.397418432 | 0.000680249 | 0.011539992 | Txnrd1 |
| ENSMUSG00000041143.12 | 481.2847692 | 0.921483034 | 0.271258373 | 3.39706761 | 0.000681121 | 0.011540619 | Tmco4 |
| ENSMUSG00000018930.3 | 128.3133597 | 1.568523901 | 0.461988075 | 3.395161016 | 0.000685883 | 0.011592841 | Ccl4 |
| ENSMUSG00000020604.9 | 52.49238059 | 1.118865445 | 0.329519279 | 3.395447604 | 0.000685165 | 0.011592841 | Arsg |
| ENSMUSG00000060802.8 | 9934.974558 | -1.084778066 | 0.319691727 | -3.393200313 | 0.000690811 | 0.01166187 | B2m |
| ENSMUSG00000039521.8 | 11.14324741 | -1.730384919 | 0.510665351 | -3.388490949 | 0.000702783 | 0.011849496 | Foxp3 |
| ENSMUSG00000003810.8 | 465.1574497 | 0.671306642 | 0.198160852 | 3.387685489 | 0.00070485 | 0.011864661 | Mast2 |
| ENSMUSG00000026389.12 | 20.33818726 | 1.448384016 | 0.427570875 | 3.387471179 | 0.000705401 | 0.011864661 | Steap3 |
| ENSMUSG00000035840.6 | 275.2014058 | -0.86318396 | 0.254887258 | -3.386532409 | 0.000707819 | 0.011890845 | Lysmd3 |
| ENSMUSG00000003380.10 | 281.3069154 | -0.78430423 | 0.23173641 | -3.384466988 | 0.000713166 | 0.01196611 | Rabac1 |
| ENSMUSG00000003429.10 | 2936.364942 | -0.720825134 | 0.213009338 | -3.384007202 | 0.000714361 | 0.01197162 | Rps11 |
| ENSMUSG00000032501.8 | 365.4016094 | 0.677039714 | 0.200278427 | 3.380492463 | 0.000723561 | 0.012111089 | Trib1 |
| ENSMUSG00000071281.6 | 145.464274 | -0.860428671 | 0.254569845 | -3.379931634 | 0.000725039 | 0.012121136 | Zfp65 |
| ENSMUSG00000004085.10 | 49.87811242 | 1.624507679 | 0.480785591 | 3.378860992 | 0.000727868 | 0.012153724 | Zak |
| ENSMUSG00000026728.5 | 13407.18283 | -0.711831569 | 0.210727652 | -3.377969433 | 0.000730232 | 0.01217847 | Vim |
| ENSMUSG00000040212.7 | 806.024956 | -0.893261512 | 0.264755587 | -3.37390996 | 0.000741086 | 0.01234458 | Emp3 |
| ENSMUSG00000030144.2 | 23.86266089 | 1.632612813 | 0.484009347 | 3.373101832 | 0.000743265 | 0.012365953 | Clec4d |
| ENSMUSG00000055865.8 | 87.07453889 | -1.10939313 | 0.329110225 | -3.370886242 | 0.000749268 | 0.012450831 | Fam19a3 |
| ENSMUSG00000070031.7 | 412.4374448 | 0.604492857 | 0.179380885 | 3.369884459 | 0.000751997 | 0.012481163 | Sp140 |
| ENSMUSG00000018821.3 | 32.81518217 | -1.317490891 | 0.391083459 | -3.368822845 | 0.000754899 | 0.012514291 | Avpi1 |
| ENSMUSG00000035004.2 | 167.1854112 | 1.546640924 | 0.459322461 | 3.367222494 | 0.000759294 | 0.012559174 | Igsf6 |
| ENSMUSG00000067768.8 | 141.5742374 | -0.922269767 | 0.273900188 | -3.367174644 | 0.000759426 | 0.012559174 | Xlr4b |
| ENSMUSG00000036599.10 | 207.4493978 | 0.733663367 | 0.217978555 | 3.365759391 | 0.000763333 | 0.012608684 | Chst12 |
| ENSMUSG00000032514.7 | 12.8363345 | 1.784007283 | 0.53014285 | 3.365144477 | 0.000765036 | 0.012621721 | Ttc21a |
| ENSMUSG00000024480.7 | 452.3508143 | -0.905800672 | 0.269233303 | -3.364370829 | 0.000767184 | 0.012642055 | Ap3s1 |
| ENSMUSG00000086862.1 | 29.30992054 | -1.534986872 | 0.4565098 | -3.362440129 | 0.000772569 | 0.012715616 | Gm13546 |
| ENSMUSG00000017417.10 | 201.6682146 | 0.795628873 | 0.236718438 | 3.361076901 | 0.000776392 | 0.012748154 | Plxdc1 |
| ENSMUSG00000092622.4 | 50.99459648 | 1.213863121 | 0.361119305 | 3.361390829 | 0.00077551 | 0.012748154 | Khdc3 |
| ENSMUSG00000025477.9 | 56.55285147 | 1.174885534 | 0.349659301 | 3.360086603 | 0.00077918 | 0.012778744 | Inpp5a |
| ENSMUSG00000035517.13 | 51.15089757 | 1.386549837 | 0.412804981 | 3.358849581 | 0.000782677 | 0.012820855 | Tdrd7 |
| ENSMUSG00000060739.7 | 530.3896633 | -0.728064698 | 0.216876238 | -3.357051484 | 0.000787784 | 0.012889236 | Nsa2 |
| ENSMUSG00000029641.7 | 59.97345518 | -1.552863066 | 0.462637328 | -3.356545124 | 0.000789228 | 0.012893878 | Rasl11a |
| ENSMUSG00000039191.8 | 515.9842203 | -0.82806989 | 0.246721246 | -3.356297457 | 0.000789936 | 0.012893878 | Rbpj |
| ENSMUSG00000028681.7 | 6.047017825 | 1.790107059 | 0.533411593 | 3.355958296 | 0.000790905 | 0.012894459 | Ptch2 |
| ENSMUSG00000012519.10 | 75.3650509 | 1.220993921 | 0.364036217 | 3.354045189 | 0.000796394 | 0.012945715 | Mlkl |
| ENSMUSG00000041488.11 | 20.54044973 | 1.597972926 | 0.476454622 | 3.353882723 | 0.000796861 | 0.012945715 | Stx3 |
| ENSMUSG00000093803.2 | 78.76214699 | 1.096801849 | 0.326974683 | 3.354393798 | 0.000795391 | 0.012945715 | Ppp2r3d |
| ENSMUSG00000031822.14 | 852.5497166 | 1.049559248 | 0.313012927 | 3.353085951 | 0.000799159 | 0.012967784 | Gse1 |
| ENSMUSG00000001507.12 | 62.19665208 | -1.248326458 | 0.372356318 | -3.352505109 | 0.000800838 | 0.012976571 | Itga3 |
| ENSMUSG00000030427.13 | 11.42029259 | 1.672933465 | 0.499048137 | 3.352248695 | 0.00080158 | 0.012976571 | Lilra6 |
| ENSMUSG00000031950.7 | 388.7805376 | -0.839758707 | 0.250748097 | -3.349013285 | 0.000810999 | 0.013113684 | Gabarapl2 |
| ENSMUSG00000006360.7 | 1681.25363 | -1.164551884 | 0.347806007 | -3.348279958 | 0.000813148 | 0.013117715 | Crip1 |
| ENSMUSG00000037336.10 | 47.73085108 | 1.241796786 | 0.37084274 | 3.348580548 | 0.000812267 | 0.013117715 | Mfsd2b |
| ENSMUSG00000034868.8 | 1637.707247 | -0.868495568 | 0.259680109 | -3.344482449 | 0.000824362 | 0.013283105 | Myl12b |
| ENSMUSG00000094840.1 | 6.130731965 | 1.793504117 | 0.536451886 | 3.343271158 | 0.00082797 | 0.013325677 | A630081J09Rik |
| ENSMUSG00000032412.8 | 2561.353353 | -0.868176866 | 0.259755917 | -3.342279466 | 0.000830934 | 0.013357813 | Atp1b3 |
| ENSMUSG00000005580.7 | 9.760556411 | 1.678065157 | 0.502235611 | 3.341191106 | 0.000834198 | 0.013394695 | Adcy9 |
| ENSMUSG00000001666.8 | 72.31345827 | -0.983978362 | 0.294618568 | -3.339838242 | 0.000838272 | 0.013444481 | Ddt |
| ENSMUSG00000034438.12 | 742.0919897 | 1.036754221 | 0.310814729 | 3.335601966 | 0.000851149 | 0.013635175 | Gbp8 |
| ENSMUSG00000020189.9 | 1901.578079 | -0.615331139 | 0.184584068 | -3.333609148 | 0.00085727 | 0.013717318 | Osbpl8 |
| ENSMUSG00000032279.10 | 649.7627992 | -0.668878222 | 0.200669472 | -3.333233576 | 0.000858428 | 0.013719952 | Idh3a |
| ENSMUSG00000035638.10 | 19.80866449 | 1.649030531 | 0.494797065 | 3.332741136 | 0.000859949 | 0.013728367 | Muc20 |
| ENSMUSG00000000881.8 | 61.24740581 | 1.07636413 | 0.323000423 | 3.332392325 | 0.000861028 | 0.013729715 | Dlg3 |

| | | | | | | | |
|---|---|---|---|---|---|---|---|
| ENSMUSG00000025473.12 | 1622.432581 | -0.905268656 | 0.27176612 | -3.331057809 | 0.000865166 | 0.013779795 | Adam8 |
| ENSMUSG00000052749.8 | 41.75549181 | 1.430996345 | 0.430091855 | 3.327187738 | 0.000877272 | 0.013956516 | Trim30b |
| ENSMUSG00000038235.4 | 20.53171814 | 1.72822328 | 0.519572419 | 3.326241379 | 0.000880257 | 0.013987876 | F11r |
| ENSMUSG00000024206.10 | 41.79736105 | 1.129764863 | 0.339758644 | 3.325198298 | 0.000883557 | 0.014024177 | Rfx2 |
| ENSMUSG00000049775.12 | 20823.49912 | -0.912083456 | 0.274372518 | -3.324252234 | 0.00088656 | 0.014055687 | Tmsb4x |
| ENSMUSG00000027007.12 | 282.9572053 | 0.824325707 | 0.24809074 | 3.322678249 | 0.000891577 | 0.014102847 | Ssfa2 |
| ENSMUSG00000050912.11 | 1091.439032 | -0.914145799 | 0.275101905 | -3.322935187 | 0.000890756 | 0.014102847 | Tmem123 |
| ENSMUSG00000077394.1 | 211.9143535 | -0.878694547 | 0.264527221 | -3.32175473 | 0.000894533 | 0.014133415 | Gm24339 |
| ENSMUSG00000039377.7 | 43.24861372 | 1.552998745 | 0.467650736 | 3.320851704 | 0.000897432 | 0.014163017 | Hlx |
| ENSMUSG00000021879.8 | 15.07934575 | -1.609121692 | 0.485297489 | -3.31574288 | 0.000913999 | 0.014391573 | Dnah12 |
| ENSMUSG00000041895.11 | 39.84719976 | 1.523207286 | 0.459383386 | 3.315764854 | 0.000913927 | 0.014391573 | Wipi1 |
| ENSMUSG00000047187.9 | 648.8635522 | -0.820963125 | 0.247620319 | -3.315410984 | 0.000915085 | 0.014392262 | Rab2a |
| ENSMUSG00000029714.7 | 570.0855404 | 0.967340742 | 0.292083291 | 3.311866068 | 0.000926759 | 0.014559289 | Gigyf1 |
| ENSMUSG00000103968.1 | 5.785769085 | 1.776436906 | 0.536618204 | 3.310429823 | 0.000931528 | 0.014617581 | 2610509F24Rik |
| ENSMUSG00000001228.10 | 1135.557974 | 0.633504281 | 0.19149673 | 3.308172833 | 0.000939068 | 0.014719177 | Uhrf1 |
| ENSMUSG00000034361.5 | 116.6715898 | 1.110994404 | 0.335883355 | 3.307679248 | 0.000940725 | 0.014728424 | Cpne2 |
| ENSMUSG00000034773.12 | 58.57118853 | 0.984262469 | 0.297649528 | 3.306783233 | 0.000943739 | 0.014758879 | BC030867 |
| ENSMUSG00000041936.14 | 114.8128461 | 1.396235989 | 0.422304065 | 3.306233839 | 0.000945591 | 0.014771121 | Agrn |
| ENSMUSG00000032376.8 | 982.4588015 | -0.677365198 | 0.205036389 | -3.303634059 | 0.000954403 | 0.014891926 | Usp3 |
| ENSMUSG00000043230.2 | 336.0667578 | -0.942126357 | 0.285417366 | -3.300872578 | 0.000963846 | 0.015022298 | Fam124b |
| ENSMUSG00000040860.12 | 142.3986382 | 1.125117844 | 0.341031562 | 3.299160457 | 0.000969745 | 0.015097186 | Crocc |
| ENSMUSG00000050272.9 | 33.18482337 | 1.405153698 | 0.42626584 | 3.296425768 | 0.000979235 | 0.015227764 | Dscam |
| ENSMUSG00000020166.10 | 701.5398481 | -0.702262548 | 0.213185821 | -3.294133463 | 0.000987256 | 0.01533523 | Cnot2 |
| ENSMUSG00000025935.6 | 2271.714239 | -0.841698016 | 0.255572913 | -3.293377248 | 0.000989916 | 0.015342025 | Tram1 |
| ENSMUSG00000054065.7 | 720.2993551 | -0.770516146 | 0.233956593 | -3.293414964 | 0.000989783 | 0.015342025 | Pkp3 |
| ENSMUSG00000027002.9 | 237.7087484 | -0.874729495 | 0.265688937 | -3.292306802 | 0.000993691 | 0.015383279 | Nckap1 |
| ENSMUSG00000061859.12 | 40.47409639 | 1.229896676 | 0.373671566 | 3.291384164 | 0.000996957 | 0.015416545 | Inadl |
| ENSMUSG00000033762.7 | 96.87851095 | 0.906300102 | 0.275553037 | 3.289022362 | 0.00100536 | 0.015529106 | Recql4 |
| ENSMUSG00000018846.8 | 758.0576387 | -0.827269777 | 0.251561189 | -3.288542956 | 0.001007074 | 0.015536746 | Pank3 |
| ENSMUSG00000025743.10 | 77.5736909 | 1.347171743 | 0.409692005 | 3.2882549 | 0.001008105 | 0.015536746 | Sdc3 |
| ENSMUSG00000035311.12 | 1142.190635 | 0.666198106 | 0.202642024 | 3.28756145 | 0.001010591 | 0.015546541 | Gnptab |
| ENSMUSG00000040276.10 | 94.9694412 | 1.435397655 | 0.436629538 | 3.287449725 | 0.001010992 | 0.015546541 | Pacsin1 |
| ENSMUSG00000032349.9 | 821.5571599 | -0.72243375 | 0.219806302 | -3.286683513 | 0.001013747 | 0.015571559 | Elovl5 |
| ENSMUSG00000019843.10 | 3766.686676 | 0.724599799 | 0.220493922 | 3.286257468 | 0.001015282 | 0.015577804 | Fyn |
| ENSMUSG00000009376.11 | 30.96355598 | 1.644473418 | 0.500595517 | 3.285034252 | 0.0010197 | 0.015607698 | Met |
| ENSMUSG00000025236.10 | 520.3012265 | 1.105706595 | 0.336551573 | 3.285400173 | 0.001018376 | 0.015607698 | Adpgk |
| ENSMUSG00000053113.3 | 662.5206175 | 0.799218853 | 0.243309714 | 3.284779883 | 0.001020621 | 0.015607698 | Socs3 |
| ENSMUSG00000003032.8 | 57.25535751 | 1.534586094 | 0.467254731 | 3.284260152 | 0.001022505 | 0.015619215 | Klf4 |
| ENSMUSG00000056656.5 | 96.14755553 | 1.324722684 | 0.403825579 | 3.280432827 | 0.001036479 | 0.015815187 | Apol8 |
| ENSMUSG00000007097.10 | 17.45673403 | -1.514491832 | 0.46199222 | -3.278176056 | 0.001044802 | 0.015924583 | Atp1a2 |
| ENSMUSG00000028398.8 | 122.9314171 | -0.94367092 | 0.288111348 | -3.275368802 | 0.001055241 | 0.016065959 | Tmem261 |
| ENSMUSG00000031444.12 | 57.69718763 | 1.645794877 | 0.502548896 | 3.274895019 | 0.001057012 | 0.016075204 | F10 |
| ENSMUSG00000066798.3 | 261.6444771 | -0.806516745 | 0.246407581 | -3.273100366 | 0.001063747 | 0.016159826 | Zbtb6 |
| ENSMUSG00000020180.9 | 206.8837474 | -0.811330101 | 0.247993321 | -3.27158045 | 0.001069482 | 0.016226525 | Snrpd3 |
| ENSMUSG00000074918.4 | 220.9418343 | -0.694951477 | 0.212437986 | -3.271314559 | 0.001070488 | 0.016226525 | Inafm2 |
| ENSMUSG00000025930.6 | 42.66110235 | -1.686063915 | 0.515555165 | -3.270385071 | 0.001074012 | 0.016262092 | Msc |
| ENSMUSG00000027995.10 | 53.1333846 | 1.57924897 | 0.483422096 | 3.266811722 | 0.00108766 | 0.016435271 | Tlr2 |
| ENSMUSG00000050592.8 | 1267.583353 | -0.751678312 | 0.230098497 | -3.266767586 | 0.001087829 | 0.016435271 | Fam78a |
| ENSMUSG00000034041.4 | 46.29377553 | 1.598986223 | 0.489581711 | 3.266025237 | 0.001090685 | 0.016460408 | Lyl1 |
| ENSMUSG00000028211.7 | 2129.835861 | -0.809350233 | 0.247900803 | -3.26481489 | 0.001095356 | 0.016512856 | Trp53inp1 |
| ENSMUSG00000040033.11 | 649.978767 | 0.704886164 | 0.21599938 | 3.263371236 | 0.001100952 | 0.016543032 | Stat2 |
| ENSMUSG00000053870.7 | 153.1469211 | -0.95650075 | 0.29307575 | -3.263663921 | 0.001099815 | 0.016543032 | Fpgt |
| ENSMUSG00000094411.1 | 66.15974956 | -1.441501081 | 0.441654519 | -3.263865801 | 0.001099032 | 0.016543032 | Snord16a |
| ENSMUSG00000030522.10 | 173.9375797 | 1.012958687 | 0.310526093 | 3.262072692 | 0.001106008 | 0.016600936 | Mtmr10 |
| ENSMUSG00000025409.10 | 336.0841852 | 1.196049731 | 0.366842283 | 3.260392239 | 0.001112582 | 0.016681487 | Mbd6 |
| ENSMUSG00000005533.9 | 134.4646028 | 1.349553247 | 0.414033339 | 3.259527969 | 0.001115978 | 0.016714248 | Igf1r |
| ENSMUSG00000001128.7 | 85.38705873 | 1.07320324 | 0.329311547 | 3.25892988 | 0.001118333 | 0.016731377 | Cfp |
| ENSMUSG00000039953.9 | 262.8817402 | 0.965864546 | 0.29668934 | 3.255474378 | 0.001132031 | 0.016917984 | Clstn1 |
| ENSMUSG00000040785.13 | 545.0539223 | 0.750066678 | 0.230592725 | 3.252776858 | 0.001142832 | 0.017060938 | Ttc3 |
| ENSMUSG00000021678.8 | 116.7330847 | -0.899687365 | 0.276642812 | -3.252162443 | 0.001145305 | 0.017060974 | F2rl1 |
| ENSMUSG00000022957.15 | 65.45023398 | -1.250896667 | 0.3846092 | -3.252383634 | 0.001144414 | 0.017060974 | Itsn1 |
| ENSMUSG00000056399.1 | 77.53409939 | 1.613441737 | 0.496443304 | 3.25000201 | 0.001154042 | 0.017172592 | Prss34 |

| | | | | | | | |
|---|---|---|---|---|---|---|---|
| ENSMUSG00000054435.12 | 4972.674573 | -0.651144813 | 0.200411129 | -3.249045183 | 0.001157931 | 0.017211913 | Gimap4 |
| ENSMUSG00000000282.8 | 342.2852871 | 0.658726023 | 0.20283585 | 3.24758184 | 0.001163902 | 0.017282065 | Mnt |
| ENSMUSG00000035441.10 | 23.75647406 | 1.42595766 | 0.439170658 | 3.246932906 | 0.001166559 | 0.017302912 | Myo1d |
| ENSMUSG00000019782.9 | 335.1287501 | -0.809078414 | 0.249234809 | -3.246249657 | 0.001169362 | 0.017325886 | Rwdd1 |
| ENSMUSG00000030134.9 | 64.40447193 | 1.545566311 | 0.47618106 | 3.245753438 | 0.001171402 | 0.01733751 | Rasgef1a |
| ENSMUSG00000047996.12 | 79.5799155 | -1.31035365 | 0.403763615 | -3.245348519 | 0.00117307 | 0.017343595 | Prrg1 |
| ENSMUSG00000026532.7 | 53.92995357 | 1.625779374 | 0.502073768 | 3.238128493 | 0.001203166 | 0.01776954 | Spta1 |
| ENSMUSG00000045996.8 | 68.7283248 | -1.223051651 | 0.377743475 | -3.237783658 | 0.001204621 | 0.017772024 | Polr2k |
| ENSMUSG00000000552.9 | 16.65349345 | 1.562241481 | 0.48277685 | 3.235949448 | 0.001212389 | 0.017837371 | Zfp385a |
| ENSMUSG00000046169.9 | 104.2742601 | 0.962319806 | 0.297346693 | 3.236356174 | 0.001210662 | 0.017837371 | Adamts6 |
| ENSMUSG00000050600.5 | 800.5240011 | 1.22166599 | 0.377544134 | 3.235822995 | 0.001212926 | 0.017837371 | Zfp831 |
| ENSMUSG00000021733.9 | 1460.742391 | -0.744889516 | 0.230372482 | -3.233413599 | 0.001223203 | 0.017969377 | Slc4a7 |
| ENSMUSG00000019478.11 | 78.45657467 | -0.898946482 | 0.278077959 | -3.232713894 | 0.001226203 | 0.017975199 | Rab4a |
| ENSMUSG00000093930.1 | 551.6993049 | -0.801520777 | 0.247924133 | -3.232927621 | 0.001225286 | 0.017975199 | Hmgcs1 |
| ENSMUSG00000022946.8 | 993.4536704 | 0.834795386 | 0.258303474 | 3.231839564 | 0.001229961 | 0.018011167 | Dopey2 |
| ENSMUSG00000020432.8 | 68.04701442 | 1.419525898 | 0.439693729 | 3.228442447 | 0.001244663 | 0.018207152 | Tcn2 |
| ENSMUSG00000026727.6 | 351.7190133 | -0.840789755 | 0.260516763 | -3.227392146 | 0.001249241 | 0.018254786 | Rsu1 |
| ENSMUSG00000023043.6 | 13.84298923 | -1.580569003 | 0.489855296 | -3.226603888 | 0.001252687 | 0.018285794 | Krt18 |
| ENSMUSG00000030223.10 | 25.32821547 | 1.579728948 | 0.489690307 | 3.225975533 | 0.001255441 | 0.018306635 | Ptpro |
| ENSMUSG00000055782.8 | 83.09378599 | 1.638809233 | 0.508508383 | 3.222777221 | 0.001269543 | 0.018492735 | Abcd2 |
| ENSMUSG00000032745.13 | 1176.407946 | -0.606816381 | 0.188330761 | -3.222077896 | 0.001272645 | 0.018518397 | Gpbp1 |
| ENSMUSG00000024070.11 | 481.6143091 | -0.670032538 | 0.208006824 | -3.221204602 | 0.00127653 | 0.018555369 | Prkd3 |
| ENSMUSG00000026478.10 | 1240.903191 | -0.787850985 | 0.24463546 | -3.220510157 | 0.001279627 | 0.018580824 | Lamc1 |
| ENSMUSG00000042489.11 | 723.1942834 | 0.674895349 | 0.209705177 | 3.218305617 | 0.001289503 | 0.018704572 | Clspn |
| ENSMUSG00000022864.9 | 628.7852976 | -0.621845545 | 0.193291546 | -3.217137821 | 0.001294764 | 0.018761169 | D16Ertd472e |
| ENSMUSG00000027695.12 | 35.7985425 | 1.522989365 | 0.473467231 | 3.216673224 | 0.001296862 | 0.018771876 | Pld1 |
| ENSMUSG00000073409.8 | 3794.722313 | 0.707587935 | 0.220209273 | 3.213252223 | 0.00131241 | 0.018977034 | H2-Q6 |
| ENSMUSG00000026873.5 | 173.2025101 | 0.796357587 | 0.248174507 | 3.208861365 | 0.001332617 | 0.019249072 | Phf19 |
| ENSMUSG00000027313.3 | 7.951964488 | 1.720092948 | 0.536470588 | 3.206313607 | 0.001344474 | 0.01940004 | Chac1 |
| ENSMUSG00000050147.8 | 157.9868159 | -0.875819289 | 0.273393794 | -3.203508305 | 0.001357641 | 0.019569591 | F2rl3 |
| ENSMUSG00000032812.12 | 988.0339685 | 0.93896635 | 0.293194373 | 3.202538778 | 0.00136222 | 0.01961511 | Arap1 |
| ENSMUSG00000057315.10 | 11.9800249 | 1.659755932 | 0.518433921 | 3.2014802 | 0.001367235 | 0.019666817 | Arhgap24 |
| ENSMUSG00000093966.1 | 10.15450767 | -1.716968516 | 0.536371239 | -3.201082369 | 0.001369124 | 0.019673498 | Trav4-3 |
| ENSMUSG00000019461.8 | 59.0436868 | 1.035729626 | 0.323700695 | 3.199652155 | 0.001375935 | 0.019714554 | Plscr3 |
| ENSMUSG00000030165.12 | 170.288247 | -1.109237752 | 0.346681941 | -3.199583312 | 0.001376264 | 0.019714554 | Klrd1 |
| ENSMUSG00000040479.7 | 3715.397771 | 0.814617889 | 0.254558013 | 3.200126681 | 0.001373672 | 0.019714554 | Dgkz |
| ENSMUSG00000038764.10 | 355.3114282 | -0.904155823 | 0.282726761 | -3.197984578 | 0.001383917 | 0.019803634 | Ptpn3 |
| ENSMUSG00000072082.6 | 449.7644084 | 0.72378793 | 0.226384326 | 3.197164501 | 0.001387858 | 0.019839467 | Ccnf |
| ENSMUSG00000005696.7 | 409.7939606 | 0.689215698 | 0.215602121 | 3.196701841 | 0.001390085 | 0.019850764 | Sh2d1a |
| ENSMUSG00000038545.9 | 48.26928222 | 1.168518581 | 0.365917766 | 3.193391216 | 0.001406123 | 0.020059043 | Cul7 |
| ENSMUSG00000029484.8 | 72.81798981 | 1.591641677 | 0.49867278 | 3.191755679 | 0.001414109 | 0.020152147 | Anxa3 |
| ENSMUSG00000022534.9 | 54.95149567 | 1.504721532 | 0.471606133 | 3.190631813 | 0.001419621 | 0.020189023 | Mefv |
| ENSMUSG00000022623.11 | 9.486728617 | 1.703537692 | 0.533879288 | 3.190866796 | 0.001418467 | 0.020189023 | Shank3 |
| ENSMUSG00000028277.9 | 900.4216226 | -0.848612404 | 0.266036544 | -3.189833964 | 0.001423546 | 0.020224013 | Ube2j1 |
| ENSMUSG00000076471.3 | 6.874094259 | -1.704851079 | 0.53537918 | -3.184380612 | 0.001450641 | 0.020587777 | Trbv14 |
| ENSMUSG00000030162.10 | 15.32343394 | 1.560088978 | 0.490429017 | 3.181069887 | 0.001467322 | 0.020803135 | Olr1 |
| ENSMUSG00000019851.7 | 18.97221379 | 1.52382726 | 0.479359101 | 3.178884593 | 0.001478429 | 0.020874879 | Perp |
| ENSMUSG00000029413.10 | 149.2640815 | 1.081798451 | 0.340244297 | 3.179475627 | 0.001475418 | 0.020874879 | Naaa |
| ENSMUSG00000038384.12 | 863.7205267 | 1.184582211 | 0.372592172 | 3.179299784 | 0.001476313 | 0.020874879 | Setd1b |
| ENSMUSG00000038393.10 | 3505.721735 | -0.879140492 | 0.276552953 | -3.178922812 | 0.001478235 | 0.020874879 | Txnip |
| ENSMUSG00000038456.5 | 7.927595108 | 1.668819119 | 0.525179962 | 3.177613845 | 0.001484924 | 0.020945161 | Dennd2a |
| ENSMUSG00000087289.1 | 11.53559176 | -1.644394242 | 0.517669904 | -3.176530506 | 0.001490481 | 0.021002096 | 4933424M12Rik |
| ENSMUSG00000022422.9 | 73.27319289 | 0.966519201 | 0.304343063 | 3.175755653 | 0.001494468 | 0.021036804 | Dscc1 |
| ENSMUSG00000040152.8 | 42.18907522 | 1.61699125 | 0.509236184 | 3.175326699 | 0.001496679 | 0.021046476 | Thbs1 |
| ENSMUSG00000036398.9 | 230.8661587 | -0.892089935 | 0.281043452 | -3.174206434 | 0.001502468 | 0.021106388 | Ppp1r11 |
| ENSMUSG00000017764.2 | 98.33380789 | -0.902020917 | 0.284198573 | -3.17391079 | 0.001503999 | 0.021106426 | Zswim1 |
| ENSMUSG00000042581.10 | 5.441740947 | -1.701540169 | 0.536625168 | -3.170816935 | 0.001520109 | 0.021310849 | Thsd7b |
| ENSMUSG00000030055.12 | 749.2463362 | 0.679918895 | 0.214488014 | 3.169962192 | 0.001524588 | 0.021351959 | Rab43 |
| ENSMUSG00000068523.8 | 240.9412243 | -0.999580339 | 0.315515404 | -3.16808728 | 0.001534454 | 0.021468371 | Gng5 |
| ENSMUSG00000079614.3 | 329.6349567 | -0.625276693 | 0.19751465 | -3.165723111 | 0.00154698 | 0.021621704 | Seh1l |
| ENSMUSG00000038375.11 | 137.7701698 | 1.028831369 | 0.325329765 | 3.162426192 | 0.001564604 | 0.021845921 | Trp53inp2 |
| ENSMUSG00000076281.1 | 371.3278132 | 1.694388262 | 0.535876822 | 3.161898765 | 0.00156744 | 0.02186342 | Gm24270 |

| | | | | | | | |
|---|---|---|---|---|---|---|---|
| ENSMUSG00000029648.9 | 13.78653172 | 1.66808496 | 0.527865326 | 3.160057834 | 0.001577378 | 0.021957678 | Flt1 |
| ENSMUSG00000034275.13 | 115.7522112 | 1.334466265 | 0.422254651 | 3.160335269 | 0.001575877 | 0.021957678 | Igsf9b |
| ENSMUSG00000040732.14 | 27.20420584 | 1.610538155 | 0.50980949 | 3.15909803 | 0.001582582 | 0.021968648 | Erg |
| ENSMUSG00000056290.11 | 4417.278033 | -0.640249392 | 0.202658092 | -3.159258949 | 0.001581709 | 0.021968648 | Ms4a4b |
| ENSMUSG00000064023.3 | 175.1554738 | -0.995172477 | 0.315024462 | -3.159032383 | 0.001582939 | 0.021968648 | Klk8 |
| ENSMUSG00000054364.4 | 103.8705952 | 1.172867432 | 0.371376313 | 3.158164348 | 0.00158766 | 0.022012048 | Rhob |
| ENSMUSG00000036372.10 | 207.6521066 | -0.93827128 | 0.297240285 | -3.156608736 | 0.001596154 | 0.022107608 | Tmem258 |
| ENSMUSG00000032131.11 | 13.06758858 | 1.652058849 | 0.523458522 | 3.15604538 | 0.00159924 | 0.022128158 | Abcg4 |
| ENSMUSG00000035295.6 | 10.50208824 | -1.635360111 | 0.518421514 | -3.15449893 | 0.00160774 | 0.022223501 | Wdr38 |
| ENSMUSG00000090733.2 | 628.1787841 | -0.799256376 | 0.253394617 | -3.15419635 | 0.001609408 | 0.02222431 | Rps27 |
| ENSMUSG00000017314.8 | 40.81360142 | 1.088658604 | 0.345239864 | 3.153339802 | 0.001614138 | 0.022267364 | Mpp2 |
| ENSMUSG00000028965.9 | 178.1234496 | 1.1128204 | 0.352967411 | 3.15275678 | 0.001617365 | 0.022289614 | Tnfrsf9 |
| ENSMUSG00000018927.3 | 55.89372498 | 1.494270045 | 0.474051998 | 3.152122662 | 0.001620882 | 0.022315808 | Ccl6 |
| ENSMUSG00000037855.11 | 25.4615615 | 1.528137167 | 0.484994145 | 3.150836322 | 0.001628037 | 0.022391995 | Zfp365 |
| ENSMUSG00000063430.8 | 137.9528622 | 0.863699505 | 0.274303525 | 3.148699987 | 0.001639985 | 0.022533879 | Wscd2 |
| ENSMUSG00000024269.7 | 142.1907875 | -0.799602549 | 0.254050299 | -3.147418256 | 0.001647192 | 0.022606732 | Tpgs2 |
| ENSMUSG00000096678.2 | 7.817983456 | -1.679387392 | 0.533617369 | -3.147175278 | 0.001648561 | 0.022606732 | Trav9-4 |
| ENSMUSG00000024095.11 | 1289.65385 | -0.684059487 | 0.217410572 | -3.146394774 | 0.001652967 | 0.022644668 | Hnrnpll |
| ENSMUSG00000047242.10 | 171.1859741 | -0.876472343 | 0.278677666 | -3.14511154 | 0.001660235 | 0.022721692 | Taf9b |
| ENSMUSG00000078784.4 | 148.0751358 | -1.004282198 | 0.319419063 | -3.144089734 | 0.001666043 | 0.022778608 | 1810022K09Rik |
| ENSMUSG00000022575.4 | 509.8721418 | 0.761668752 | 0.24229852 | 3.143513846 | 0.001669325 | 0.022800902 | Gsdmd |
| ENSMUSG00000029401.7 | 390.1020819 | 0.609348075 | 0.193884707 | 3.142837223 | 0.001673189 | 0.022815222 | Rilpl2 |
| ENSMUSG00000029414.7 | 433.5140867 | 0.634392754 | 0.201859019 | 3.142751599 | 0.001673678 | 0.022815222 | Kntc1 |
| ENSMUSG00000087260.1 | 194.4758531 | -0.958987593 | 0.305372176 | -3.140389557 | 0.001687233 | 0.02297732 | Lamtor5 |
| ENSMUSG00000015944.8 | 24.71343492 | 1.255121634 | 0.400416387 | 3.134541125 | 0.001721231 | 0.023319915 | Gatsl2 |
| ENSMUSG00000017421.14 | 2004.93161 | -0.63682176 | 0.203202645 | -3.133924567 | 0.001724852 | 0.023319915 | Zfp207 |
| ENSMUSG00000021076.5 | 388.9846864 | -0.828217425 | 0.264193174 | -3.134893348 | 0.001719166 | 0.023319915 | Actr10 |
| ENSMUSG00000025903.10 | 469.8204857 | -0.808961386 | 0.258145142 | -3.133746306 | 0.0017259 | 0.023319915 | Lypla1 |
| ENSMUSG00000034321.9 | 120.3901962 | -0.787462983 | 0.251258828 | -3.134070906 | 0.001723992 | 0.023319915 | Exosc1 |
| ENSMUSG00000036309.10 | 547.5581957 | -0.86242258 | 0.275090685 | -3.13504828 | 0.001718258 | 0.023319915 | Skp1a |
| ENSMUSG00000043110.2 | 25.00825435 | 1.474436177 | 0.470247904 | 3.135444441 | 0.001715939 | 0.023319915 | Lrrn4 |
| ENSMUSG00000045165.5 | 923.5354311 | -0.779764628 | 0.248675806 | -3.135667443 | 0.001714635 | 0.023319915 | AI467606 |
| ENSMUSG00000000489.6 | 180.2067628 | 1.15483034 | 0.368580218 | 3.133185893 | 0.001729199 | 0.023341649 | Pdgfb |
| ENSMUSG00000036478.7 | 6249.673478 | -0.584399172 | 0.186539534 | -3.132843528 | 0.001731217 | 0.023343965 | Btg1 |
| ENSMUSG00000091694.4 | 23.66072767 | 1.569593641 | 0.50105405 | 3.132583485 | 0.001732751 | 0.023343965 | Apol11b |
| ENSMUSG00000032531.11 | 22.17012618 | 1.516831479 | 0.484500609 | 3.130711192 | 0.001743836 | 0.023470396 | Amotl2 |
| ENSMUSG00000037406.6 | 12.49718905 | 1.501343361 | 0.479876602 | 3.128602964 | 0.001756395 | 0.02361641 | Htra4 |
| ENSMUSG00000027835.7 | 550.0562577 | -0.856747795 | 0.273974929 | -3.127102898 | 0.001765381 | 0.023694759 | Pdcd10 |
| ENSMUSG00000067288.8 | 1151.235901 | -0.790623502 | 0.252833048 | -3.127057595 | 0.001765653 | 0.023694759 | Rps28 |
| ENSMUSG00000032735.10 | 16.60646322 | -1.52801138 | 0.489009813 | -3.124704943 | 0.001779835 | 0.023861884 | Ablim3 |
| ENSMUSG00000032103.6 | 134.7209641 | -0.795273964 | 0.254540419 | -3.124352382 | 0.001781969 | 0.023867324 | Pus3 |
| ENSMUSG00000030729.12 | 400.6177109 | -0.909994072 | 0.291494102 | -3.121826699 | 0.001797327 | 0.024049699 | Pgm2l1 |
| ENSMUSG00000023216.9 | 14.29750668 | 1.627235091 | 0.521616719 | 3.119599185 | 0.001810973 | 0.024208832 | Epb4.2 |
| ENSMUSG00000003545.2 | 321.0365059 | 0.838000267 | 0.268722491 | 3.118459737 | 0.00181799 | 0.024279131 | Fosb |
| ENSMUSG00000021108.13 | 2092.738366 | 0.741946796 | 0.237998962 | 3.117437109 | 0.001824309 | 0.024339757 | Prkch |
| ENSMUSG00000042121.12 | 1009.722687 | 0.771884959 | 0.247624815 | 3.117155114 | 0.001826055 | 0.024339757 | Ssh1 |
| ENSMUSG00000026566.11 | 7.44467737 | 1.668410344 | 0.535396031 | 3.116217245 | 0.001831873 | 0.02439376 | Mpzl1 |
| ENSMUSG00000004099.11 | 3187.059164 | 0.54646221 | 0.17543179 | 3.114955441 | 0.001839727 | 0.02447475 | Dnmt1 |
| ENSMUSG00000020732.9 | 417.0070792 | -0.648081078 | 0.208153869 | -3.113471212 | 0.001849005 | 0.024574511 | Rab37 |
| ENSMUSG00000100605.1 | 28.97845976 | -1.287868979 | 0.413896349 | -3.111573661 | 0.00186093 | 0.024709219 | Gm29243 |
| ENSMUSG00000033253.14 | 1415.003255 | 0.899654117 | 0.289220021 | 3.110621848 | 0.001866939 | 0.024765182 | Szt2 |
| ENSMUSG00000031760.8 | 10.54542809 | 1.624858425 | 0.522493311 | 3.109816702 | 0.001872035 | 0.024808952 | Mt3 |
| ENSMUSG00000033209.13 | 47.60644959 | 1.253653095 | 0.403215384 | 3.109140043 | 0.001876328 | 0.024842002 | Ttc28 |
| ENSMUSG00000028803.14 | 1736.08621 | -0.697824223 | 0.2245898 | -3.10710559 | 0.001889289 | 0.024989648 | Nipal3 |
| ENSMUSG00000007613.11 | 396.1572264 | -0.691662435 | 0.222734759 | -3.105318804 | 0.00190074 | 0.025117056 | Tgfbr1 |
| ENSMUSG00000094655.1 | 1049.585919 | -0.695658536 | 0.224080974 | -3.104496212 | 0.001906034 | 0.025162925 | Gm25360 |
| ENSMUSG00000074151.8 | 7343.729619 | 1.051951378 | 0.338957842 | 3.103487356 | 0.001912544 | 0.025224757 | Nlrc5 |
| ENSMUSG00000035673.9 | 1281.356152 | 0.701811858 | 0.22616827 | 3.103051807 | 0.001915361 | 0.025237806 | Sbno2 |
| ENSMUSG00000016028.9 | 1830.362346 | 0.792060655 | 0.255488499 | 3.100181249 | 0.001934023 | 0.02536437 | Celsr1 |
| ENSMUSG00000030774.9 | 46.87388386 | 1.520162694 | 0.490393961 | 3.099880538 | 0.001935987 | 0.02536437 | Pak1 |
| ENSMUSG00000037129.7 | 100.5450107 | 0.987291824 | 0.318470094 | 3.100108434 | 0.001934498 | 0.02536437 | Tmprss13 |
| ENSMUSG00000038264.7 | 219.5983443 | 1.045309141 | 0.337103459 | 3.100855578 | 0.001929624 | 0.02536437 | Sema7a |

| | | | | | | | |
|---|---|---|---|---|---|---|---|
| ENSMUSG00000047409.9 | 34.18803362 | 1.306421999 | 0.421362389 | 3.100471315 | 0.001932129 | 0.02536437 | Ctdspl |
| ENSMUSG00000094766.2 | 11.67820377 | -1.614348214 | 0.520615085 | -3.10084794 | 0.001929674 | 0.02536437 | Trav7-4 |
| ENSMUSG00000028088.10 | 38.37778037 | 1.12144387 | 0.361863918 | 3.099076237 | 0.001941251 | 0.025395011 | Fmo5 |
| ENSMUSG00000037013.11 | 945.1012916 | -0.54443219 | 0.175682152 | -3.098961296 | 0.001942004 | 0.025395011 | Ss18 |
| ENSMUSG00000016477.13 | 196.3218546 | 0.670974099 | 0.216672965 | 3.096713521 | 0.001956789 | 0.025492682 | E2f3 |
| ENSMUSG00000023951.12 | 25.90385331 | 1.608395761 | 0.519251139 | 3.097529578 | 0.001951409 | 0.025492682 | Vegfa |
| ENSMUSG00000051359.10 | 461.1978993 | -0.759408132 | 0.245199234 | -3.097106458 | 0.001954197 | 0.025492682 | Ncald |
| ENSMUSG00000070390.8 | 82.52763311 | 1.087819949 | 0.351283253 | 3.096703134 | 0.001956857 | 0.025492682 | Nlrp1b |
| ENSMUSG00000020647.9 | 625.4478698 | 0.753603549 | 0.243411187 | 3.096010325 | 0.001961435 | 0.025528238 | Ncoa1 |
| ENSMUSG00000021374.9 | 2449.457622 | -0.511550132 | 0.16529548 | -3.09476177 | 0.00196971 | 0.025611799 | Nup153 |
| ENSMUSG00000030872.10 | 36.82901617 | 1.267642671 | 0.409719572 | 3.093927548 | 0.001975257 | 0.025659761 | Gga2 |
| ENSMUSG00000027822.12 | 301.2655422 | -0.664019302 | 0.21466807 | -3.093237393 | 0.001979857 | 0.025695342 | Slc33a1 |
| ENSMUSG00000068394.4 | 491.4279086 | 0.647966206 | 0.209502523 | 3.092880196 | 0.001982241 | 0.025702133 | Cep152 |
| ENSMUSG00000041841.7 | 1062.94395 | -0.912713358 | 0.295268454 | -3.091130616 | 0.001993959 | 0.025808668 | Rpl37 |
| ENSMUSG00000042066.11 | 19.13480021 | 1.607936784 | 0.520183492 | 3.091095372 | 0.001994196 | 0.025808668 | Tmcc2 |
| ENSMUSG00000040751.8 | 61.82074857 | 1.284395065 | 0.41569029 | 3.089788468 | 0.002002991 | 0.025898223 | Lat2 |
| ENSMUSG00000075028.7 | 118.6654697 | 1.15300483 | 0.373286056 | 3.088796949 | 0.002009687 | 0.025960499 | Prdm11 |
| ENSMUSG00000036553.12 | 49.87019887 | 1.510033861 | 0.489020361 | 3.087875233 | 0.002015931 | 0.026016813 | Sh3tc1 |
| ENSMUSG00000030325.12 | 28.88645874 | -1.208078976 | 0.39131223 | -3.08725075 | 0.002020171 | 0.026047192 | Klrb1c |
| ENSMUSG00000037321.13 | 4108.498069 | 0.638715468 | 0.206909291 | 3.086934691 | 0.00202232 | 0.026050578 | Tap1 |
| ENSMUSG00000001506.10 | 14.6958486 | 1.544181912 | 0.500437673 | 3.085662802 | 0.00203099 | 0.026107876 | Col1a1 |
| ENSMUSG00000017830.11 | 148.6783724 | 0.767991816 | 0.248952114 | 3.084897744 | 0.002036221 | 0.026107876 | Dhx58 |
| ENSMUSG00000022336.2 | 1298.692211 | -0.852481856 | 0.276325887 | -3.085059697 | 0.002035113 | 0.026107876 | Eif3e |
| ENSMUSG00000028927.6 | 468.2111322 | 0.942287182 | 0.305426096 | 3.085156092 | 0.002034454 | 0.026107876 | Padi2 |
| ENSMUSG00000045328.7 | 1201.796217 | 0.571227115 | 0.185157461 | 3.085088295 | 0.002034917 | 0.026107876 | Cenpe |
| ENSMUSG00000027854.8 | 685.9257479 | -0.937766321 | 0.304037535 | -3.084376808 | 0.002039791 | 0.026129378 | Sike1 |
| ENSMUSG00000064844.1 | 22.87026532 | -1.494582183 | 0.48485523 | -3.082532867 | 0.002052471 | 0.026267439 | Gm26202 |
| ENSMUSG00000037815.6 | 545.9754776 | -0.949764635 | 0.308193864 | -3.081711699 | 0.002058141 | 0.026315616 | Ctnna1 |
| ENSMUSG00000046721.10 | 132.3189128 | -0.804996979 | 0.261278372 | -3.080993545 | 0.002063111 | 0.026354767 | Rpl14-ps1 |
| ENSMUSG00000018819.6 | 7138.440947 | -0.733687371 | 0.238360798 | -3.078053852 | 0.002083573 | 0.026579351 | Lsp1 |
| ENSMUSG00000022811.12 | 1102.729308 | -0.548929836 | 0.178376554 | -3.077365397 | 0.002088391 | 0.026579351 | Zfp148 |
| ENSMUSG00000037731.5 | 297.8456691 | 0.993023667 | 0.322684577 | 3.077381871 | 0.002088276 | 0.026579351 | Themis2 |
| ENSMUSG00000074170.4 | 186.0250906 | -1.044113697 | 0.339282249 | -3.077419172 | 0.002088015 | 0.026579351 | Plekhf1 |
| ENSMUSG00000013698.8 | 413.5648086 | -0.731213472 | 0.237790184 | -3.07503641 | 0.002104769 | 0.026763121 | Pea15a |
| ENSMUSG00000018916.5 | 11.97089067 | -1.52803028 | 0.497318208 | -3.072540388 | 0.002122451 | 0.026938352 | Csf2 |
| ENSMUSG00000090394.4 | 1465.150074 | -0.563272419 | 0.18330991 | -3.072787597 | 0.002120694 | 0.026938352 | 4930523C07Rik |
| ENSMUSG00000021520.4 | 175.2991946 | -1.160458841 | 0.377727763 | -3.072209549 | 0.002124805 | 0.026943465 | Uqcrb |
| ENSMUSG00000073902.5 | 2584.068638 | 0.545577024 | 0.177684382 | 3.070483839 | 0.002137123 | 0.027074793 | Gm1966 |
| ENSMUSG00000054598.9 | 11.67402074 | 1.580258433 | 0.514780512 | 3.069771283 | 0.002142228 | 0.027114592 | 9130230L23Rik |
| ENSMUSG00000025650.8 | 32.62378786 | 1.216681989 | 0.396521225 | 3.068390573 | 0.002152151 | 0.027215254 | Col7a1 |
| ENSMUSG00000103780.1 | 10.45533791 | 1.579237236 | 0.51522376 | 3.065148307 | 0.002175621 | 0.027486869 | Gm37524 |
| ENSMUSG00000028461.8 | 166.3587318 | -1.058211641 | 0.34532553 | -3.06438867 | 0.002181153 | 0.027525962 | Ccdc107 |
| ENSMUSG00000034709.8 | 416.9404338 | 0.650272256 | 0.212217631 | 3.064176395 | 0.002182702 | 0.027525962 | Ppp1r21 |
| ENSMUSG00000030629.10 | 524.8655034 | -0.682591496 | 0.222913806 | -3.062131988 | 0.002197665 | 0.027689379 | Zfand6 |
| ENSMUSG00000007670.9 | 1555.861738 | 0.634810331 | 0.207427378 | 3.060397988 | 0.00221043 | 0.027814223 | Khsrp |
| ENSMUSG00000040594.14 | 21.714137 | 1.458264357 | 0.476519721 | 3.060239258 | 0.002211602 | 0.027814223 | Ranbp17 |
| ENSMUSG00000022218.11 | 18.75217846 | 1.525600467 | 0.499044802 | 3.057041091 | 0.002235336 | 0.028087133 | Tgm1 |
| ENSMUSG00000026068.7 | 6139.829732 | -0.734279359 | 0.240395291 | -3.054466479 | 0.002254612 | 0.02830358 | Il18rap |
| ENSMUSG00000027433.5 | 2342.134228 | -0.514100995 | 0.168360817 | -3.053566768 | 0.002261384 | 0.028362807 | Xrn2 |
| ENSMUSG00000019872.9 | 686.3871727 | -0.897177326 | 0.293877779 | -3.052892699 | 0.00226647 | 0.028400798 | Smpdl3a |
| ENSMUSG00000021003.9 | 45.41686457 | 1.248737161 | 0.409201045 | 3.051647049 | 0.002275895 | 0.028470491 | Galc |
| ENSMUSG00000039542.11 | 27.11284981 | 1.560746472 | 0.511449702 | 3.051612831 | 0.002276155 | 0.028470491 | Ncam1 |
| ENSMUSG00000102974.1 | 241.163843 | 1.103333464 | 0.361623104 | 3.051059106 | 0.002280357 | 0.028497237 | ENSMUSG00000102974 |
| ENSMUSG00000075054.4 | 27.74478348 | -1.193785387 | 0.391437195 | -3.049749494 | 0.002290323 | 0.028595906 | Yae1d1 |
| ENSMUSG00000031879.9 | 99.48493831 | -0.898036647 | 0.294654078 | -3.047765884 | 0.002305495 | 0.028733372 | Fam96b |
| ENSMUSG00000058794.8 | 89.34557407 | 1.44987957 | 0.475694208 | 3.047923533 | 0.002304286 | 0.028733372 | Nfe2 |
| ENSMUSG00000035247.11 | 5170.581435 | -0.472107861 | 0.154935689 | -3.04712145 | 0.002310443 | 0.028769083 | Hectd1 |
| ENSMUSG00000022108.7 | 3820.946609 | -0.614413322 | 0.20178577 | -3.044879342 | 0.002327737 | 0.028958303 | Itm2b |
| ENSMUSG00000049751.6 | 561.7733557 | -0.724289806 | 0.237966057 | -3.043668564 | 0.002337125 | 0.029048924 | Rpl36al |
| ENSMUSG00000043323.12 | 362.4673076 | 0.780844985 | 0.25668727 | 3.042009003 | 0.002350049 | 0.029183293 | Fbrsl1 |
| ENSMUSG00000028937.10 | 501.9425942 | -0.877431411 | 0.288521353 | -3.041131627 | 0.002356908 | 0.029242173 | Acot7 |
| ENSMUSG00000023074.7 | 119.2804751 | -1.047444261 | 0.344663088 | -3.039038114 | 0.002373348 | 0.029419718 | Mospd1 |

| | | | | | | | |
|---|---|---|---|---|---|---|---|
| ENSMUSG00000022674.10 | 99.7555632 | -0.878371871 | 0.289183617 | -3.037419202 | 0.002386134 | 0.029551677 | Ube2v2 |
| ENSMUSG00000001750.11 | 866.5474628 | 0.757857746 | 0.249589372 | 3.036418343 | 0.002394069 | 0.029617753 | Tcirg1 |
| ENSMUSG00000034947.9 | 62.66595868 | 0.912501274 | 0.300540004 | 3.036205701 | 0.002395759 | 0.029617753 | Tmem106a |
| ENSMUSG00000029086.11 | 62.29171336 | 1.542193266 | 0.508059883 | 3.035455694 | 0.002401725 | 0.029664957 | Prom1 |
| ENSMUSG00000039001.8 | 1412.21318 | -0.691262672 | 0.227770197 | -3.034912742 | 0.002406053 | 0.029691854 | Rps21 |
| ENSMUSG00000043953.8 | 38.76711416 | 1.246200703 | 0.410681594 | 3.03446933 | 0.002409593 | 0.029708985 | Ccrl2 |
| ENSMUSG00000035835.10 | 18.2994988 | 1.600490386 | 0.527517673 | 3.034003351 | 0.002413317 | 0.029728369 | Lppr3 |
| ENSMUSG00000025289.11 | 133.3772597 | -1.12848883 | 0.372061627 | -3.033069655 | 0.002420797 | 0.029793929 | Prdx4 |
| ENSMUSG00000020823.12 | 399.3860009 | 0.783712785 | 0.258592203 | 3.030689925 | 0.002439957 | 0.029987848 | Sec14l1 |
| ENSMUSG00000034855.9 | 54.49275497 | 1.392714716 | 0.459554818 | 3.030573636 | 0.002440897 | 0.029987848 | Cxcl10 |
| ENSMUSG00000033863.1 | 52.66952937 | 1.165258243 | 0.384624825 | 3.02959707 | 0.002448802 | 0.030004886 | Klf9 |
| ENSMUSG00000036777.7 | 418.5810901 | 0.57562574 | 0.189984995 | 3.029848438 | 0.002446765 | 0.030004886 | Anln |
| ENSMUSG00000067017.5 | 62.72000557 | -1.129036031 | 0.372607507 | -3.030094699 | 0.002444771 | 0.030004886 | Gm3608 |
| ENSMUSG00000024614.6 | 810.7544835 | -0.736294873 | 0.243262317 | -3.026752689 | 0.002471961 | 0.030261801 | Tmx3 |
| ENSMUSG00000025868.6 | 157.0699763 | -0.927763294 | 0.306557327 | -3.026394126 | 0.002474895 | 0.030270878 | Higd2a |
| ENSMUSG00000046675.6 | 102.6567352 | -0.971196084 | 0.321007655 | -3.02546082 | 0.002482546 | 0.030313056 | Tmem251 |
| ENSMUSG00000079317.6 | 54.1345778 | -0.970130249 | 0.320657791 | -3.025437949 | 0.002482733 | 0.030313056 | Trappc2 |
| ENSMUSG00000055447.14 | 2575.930065 | -0.584319774 | 0.193184294 | -3.024675353 | 0.002489001 | 0.030362739 | Cd47 |
| ENSMUSG00000037463.10 | 54.10692452 | -1.362120119 | 0.450432118 | -3.024029733 | 0.002494319 | 0.030400754 | Fbxo27 |
| ENSMUSG00000026180.8 | 110.542351 | 1.390177616 | 0.459815122 | 3.023340356 | 0.002500009 | 0.03043553 | Cxcr2 |
| ENSMUSG00000067608.4 | 110.6071641 | -1.051279957 | 0.347743213 | -3.023150184 | 0.002501581 | 0.03043553 | Pcna-ps2 |
| ENSMUSG00000016529.5 | 81.81043756 | 1.609892836 | 0.532730929 | 3.0219624 | 0.002511417 | 0.030528313 | Il10 |
| ENSMUSG00000024401.10 | 408.4156762 | -0.587057808 | 0.194285198 | -3.021629101 | 0.002514184 | 0.030535064 | Tnf |
| ENSMUSG00000064437.1 | 51.19033081 | -1.394713107 | 0.461686161 | -3.020911658 | 0.002520149 | 0.03058061 | Snord49b |
| ENSMUSG00000034282.3 | 10.20206149 | 1.481448181 | 0.490595657 | 3.019692814 | 0.002530312 | 0.030676976 | Evpl |
| ENSMUSG00000025358.11 | 403.927655 | 0.615564886 | 0.203957687 | 3.018100935 | 0.002543642 | 0.030775193 | Cdk2 |
| ENSMUSG00000032301.9 | 591.2856831 | -0.887645117 | 0.294084575 | -3.0183328 | 0.002541696 | 0.030775193 | Psma4 |
| ENSMUSG00000038550.6 | 14.7499536 | 1.370084028 | 0.453981775 | 3.017927377 | 0.002545099 | 0.030775193 | Ciart |
| ENSMUSG00000027088.6 | 179.0620418 | -0.711755834 | 0.235869525 | -3.017582853 | 0.002547994 | 0.030783242 | Phospho2 |
| ENSMUSG00000011148.9 | 32.5550215 | 1.376816686 | 0.456361188 | 3.016945176 | 0.00255336 | 0.030817659 | Adssl1 |
| ENSMUSG00000024955.9 | 273.0669211 | 0.860910469 | 0.285380187 | 3.016714222 | 0.002555306 | 0.030817659 | Esrra |
| ENSMUSG00000050029.7 | 700.5660248 | -0.85586983 | 0.283872847 | -3.014976032 | 0.002569996 | 0.030967775 | Rap2c |
| ENSMUSG00000084786.5 | 355.3576914 | -0.875057499 | 0.290274019 | -3.014591189 | 0.002573258 | 0.030980057 | Ubl5 |
| ENSMUSG00000042312.9 | 442.2029173 | -0.873505044 | 0.289808205 | -3.014079757 | 0.0025776 | 0.031005299 | S100a13 |
| ENSMUSG00000020092.8 | 6.527308045 | 1.58885096 | 0.527862767 | 3.009969747 | 0.002612737 | 0.031373293 | Pald1 |
| ENSMUSG00000062006.8 | 1618.940497 | -0.913690933 | 0.303535332 | -3.010163351 | 0.002611072 | 0.031373293 | Rpl34 |
| ENSMUSG00000003068.11 | 707.3416345 | 0.737428945 | 0.245033409 | 3.009503679 | 0.002616749 | 0.031394169 | Stk11 |
| ENSMUSG00000020899.11 | 468.9073481 | 0.595511768 | 0.198001349 | 3.007614704 | 0.002633067 | 0.031557357 | Pfas |
| ENSMUSG00000026024.10 | 253.7983481 | 0.762365956 | 0.253496632 | 3.007400733 | 0.002634922 | 0.031557357 | Als2 |
| ENSMUSG00000039007.9 | 13.34219246 | 1.50346249 | 0.499965559 | 3.007132116 | 0.002637251 | 0.031557887 | Cpq |
| ENSMUSG00000039055.4 | 96.58846687 | 0.813267425 | 0.27057586 | 3.005690995 | 0.002649781 | 0.031680372 | Eme1 |
| ENSMUSG00000088252.1 | 160.1458249 | -1.358321679 | 0.452034675 | -3.004905941 | 0.00265663 | 0.031734777 | Snord13 |
| ENSMUSG00000029101.10 | 134.0062344 | 1.02403772 | 0.340886995 | 3.004038683 | 0.002664214 | 0.031770413 | Rgs12 |
| ENSMUSG00000032849.9 | 769.0083239 | 0.668493238 | 0.222512547 | 3.004294581 | 0.002661974 | 0.031770413 | Abcc4 |
| ENSMUSG00000020474.7 | 202.8898665 | 0.849453577 | 0.282875968 | 3.002918855 | 0.002674037 | 0.031843203 | Polm |
| ENSMUSG00000041528.11 | 962.542161 | 0.734807888 | 0.244706167 | 3.002817202 | 0.00267493 | 0.031843203 | Rnf123 |
| ENSMUSG00000036295.4 | 7.871106503 | 1.573236249 | 0.524012732 | 3.002286303 | 0.0026796 | 0.031871319 | Lrrn3 |
| ENSMUSG00000022604.14 | 835.1220436 | -0.64982413 | 0.21651255 | -3.001323161 | 0.002688091 | 0.031919723 | Cep97 |
| ENSMUSG00000026605.10 | 1152.570308 | 0.562465549 | 0.18742363 | 3.001038604 | 0.002690605 | 0.031919723 | Cenpf |
| ENSMUSG00000032224.10 | 5.169835436 | 1.606586431 | 0.535315529 | 3.001195264 | 0.002689221 | 0.031919723 | Fam81a |
| ENSMUSG00000025979.9 | 331.7727428 | -0.618536729 | 0.206371541 | -2.997199736 | 0.002724721 | 0.032296718 | Mob4 |
| ENSMUSG00000039176.13 | 712.3506797 | 0.714756283 | 0.238546236 | 2.996300827 | 0.002732767 | 0.032317278 | Polg |
| ENSMUSG00000040694.3 | 126.8820829 | 0.833171071 | 0.278063751 | 2.996331121 | 0.002732495 | 0.032317278 | Apobec2 |
| ENSMUSG00000071180.4 | 457.9790903 | -0.863052358 | 0.288046901 | -2.996221638 | 0.002733477 | 0.032317278 | Smim15 |
| ENSMUSG00000079491.5 | 411.2047978 | 0.608058814 | 0.203031137 | 2.994904243 | 0.00274531 | 0.032429416 | H2-T10 |
| ENSMUSG00000035929.11 | 3724.621595 | 0.820717046 | 0.274084955 | 2.994389268 | 0.002749949 | 0.032456444 | H2-Q4 |
| ENSMUSG00000022263.9 | 138.6388653 | 0.958750203 | 0.320590922 | 2.990571906 | 0.002784556 | 0.032813784 | Trio |
| ENSMUSG00000029554.11 | 373.4448921 | 0.893252256 | 0.298694064 | 2.990525632 | 0.002784978 | 0.032813784 | Mad1l1 |
| ENSMUSG00000038070.11 | 60.51125055 | 0.957117199 | 0.320146583 | 2.989621785 | 0.002793231 | 0.032882903 | Cntln |
| ENSMUSG00000039298.12 | 477.5219798 | 0.744288072 | 0.248978898 | 2.989362067 | 0.002795606 | 0.032882903 | Cdk5rap2 |
| ENSMUSG00000022180.6 | 8.445593648 | 1.581268695 | 0.529184858 | 2.988121582 | 0.002806979 | 0.032966136 | Slc7a8 |
| ENSMUSG00000092060.1 | 24.40735919 | 1.453537911 | 0.486447159 | 2.988069485 | 0.002807457 | 0.032966136 | Bend4 |

| | | | | | | | |
|---|---|---|---|---|---|---|---|
| ENSMUSG00000022489.5 | 286.1511282 | 0.855871827 | 0.286576766 | 2.986535992 | 0.002821576 | 0.033078642 | Pde1b |
| ENSMUSG00000023827.4 | 185.4502891 | 0.805345499 | 0.269661204 | 2.986508573 | 0.002821829 | 0.033078642 | Agpat4 |
| ENSMUSG00000020589.12 | 685.0024694 | -0.616641544 | 0.206501425 | -2.986136991 | 0.00282526 | 0.033090774 | Fam49a |
| ENSMUSG00000063193.6 | 13.42987006 | 1.538759608 | 0.515517577 | 2.984882917 | 0.002836869 | 0.033198578 | Cd300lb |
| ENSMUSG00000095597.2 | 190.7606286 | -0.715032784 | 0.239595047 | -2.984338755 | 0.002841919 | 0.033229522 | Gm6472 |
| ENSMUSG00000033088.14 | 726.3241778 | 0.676709026 | 0.226781653 | 2.983967258 | 0.002845372 | 0.033241747 | Triobp |
| ENSMUSG00000039046.11 | 214.7891052 | 0.902278878 | 0.3025124 | 2.982617832 | 0.002857946 | 0.033360421 | Usp6nl |
| ENSMUSG00000045679.9 | 472.4396307 | -0.701028748 | 0.235142434 | -2.98129409 | 0.00287033 | 0.033476678 | Pqlc3 |
| ENSMUSG00000026669.10 | 299.1131514 | 0.766587663 | 0.257164983 | 2.98091775 | 0.00287386 | 0.03348956 | Mcm10 |
| ENSMUSG00000027834.11 | 85.11737125 | -1.166423517 | 0.391631112 | -2.978372964 | 0.002897831 | 0.033740432 | Serpini1 |
| ENSMUSG00000001707.7 | 83.53175369 | -0.928890316 | 0.311922136 | -2.977955747 | 0.002901779 | 0.03375793 | Eef1e1 |
| ENSMUSG00000061414.4 | 258.5864026 | 1.10141977 | 0.369901559 | 2.977602402 | 0.002905126 | 0.033768419 | Cracr2a |
| ENSMUSG00000051506.12 | 330.7294784 | 1.192805079 | 0.400713747 | 2.976701171 | 0.002913679 | 0.03383935 | Wdfy4 |
| ENSMUSG00000029171.8 | 211.7567261 | -0.648265782 | 0.217879299 | -2.975343621 | 0.002926605 | 0.033897171 | Pgm1 |
| ENSMUSG00000029816.8 | 62.67033491 | -1.099793382 | 0.369660107 | -2.975147606 | 0.002928476 | 0.033897171 | Gpnmb |
| ENSMUSG00000058331.10 | 52.63795492 | -1.102969987 | 0.370676684 | -2.975558039 | 0.00292456 | 0.033897171 | Zfp85 |
| ENSMUSG00000063234.4 | 12.02273894 | 1.529859251 | 0.514208047 | 2.975175629 | 0.002928208 | 0.033897171 | Gpr84 |
| ENSMUSG00000029821.11 | 33.73978435 | 1.451136942 | 0.487808715 | 2.974807333 | 0.002931726 | 0.033906371 | Dfna5 |
| ENSMUSG00000026987.12 | 395.9428211 | 0.775868184 | 0.261056009 | 2.97203725 | 0.002958308 | 0.034185166 | Baz2b |
| ENSMUSG00000031497.8 | 22.84978754 | 1.200882875 | 0.404120283 | 2.971597631 | 0.002962547 | 0.034205524 | Tnfsf13b |
| ENSMUSG00000029030.10 | 808.6086013 | -0.657708925 | 0.221584039 | -2.968214357 | 0.002995354 | 0.03455542 | Tprgl |
| ENSMUSG00000076469.3 | 118.3654438 | -1.199179688 | 0.404057712 | -2.967842594 | 0.002998979 | 0.034568361 | Trbv13-2 |
| ENSMUSG00000022568.12 | 674.1952716 | 0.920803612 | 0.310390776 | 2.96659464 | 0.003011177 | 0.03459346 | Scrib |
| ENSMUSG00000037921.10 | 246.9987546 | 0.753170399 | 0.25387353 | 2.966714957 | 0.003009999 | 0.03459346 | Ddx60 |
| ENSMUSG00000042148.8 | 252.0601417 | -0.741666198 | 0.249942517 | -2.967347084 | 0.003003817 | 0.03459346 | Cox10 |
| ENSMUSG00000042331.9 | 51.81749986 | 1.276579446 | 0.430254754 | 2.967031585 | 0.003006901 | 0.03459346 | Specc1 |
| ENSMUSG00000015575.10 | 337.0973309 | -0.788470221 | 0.265862237 | -2.965709725 | 0.003019854 | 0.034648256 | Atp6v0e |
| ENSMUSG00000103358.1 | 48.9886407 | -1.313191829 | 0.442808651 | -2.965596595 | 0.003020965 | 0.034648256 | Gm37593 |
| ENSMUSG00000038633.5 | 658.7195008 | -0.910478624 | 0.307083044 | -2.964926397 | 0.003027554 | 0.03469501 | Degs1 |
| ENSMUSG00000015568.11 | 32.4482102 | 1.450362 | 0.489241115 | 2.964513722 | 0.003031617 | 0.034712771 | Lpl |
| ENSMUSG00000027583.9 | 27.55013565 | 1.14422965 | 0.386207317 | 2.962734259 | 0.003049197 | 0.03486144 | Zbtb46 |
| ENSMUSG00000044501.13 | 185.1715608 | -0.806563006 | 0.272275102 | -2.962309073 | 0.003053411 | 0.03486144 | Zfp758 |
| ENSMUSG00000054150.7 | 894.6099271 | 0.680151975 | 0.229612028 | 2.962179211 | 0.003054699 | 0.03486144 | Syne3 |
| ENSMUSG00000071528.3 | 165.6117612 | -1.133669068 | 0.382686482 | -2.962396428 | 0.003052545 | 0.03486144 | Usmg5 |
| ENSMUSG00000040268.12 | 231.2101647 | 0.849919114 | 0.286997314 | 2.96141836 | 0.003062257 | 0.034918829 | Plekha1 |
| ENSMUSG00000037936.11 | 99.83753491 | 1.158753339 | 0.39132695 | 2.961087498 | 0.003065548 | 0.034923544 | Scarb1 |
| ENSMUSG00000038024.13 | 1404.51222 | -0.589231389 | 0.199006263 | -2.96086857 | 0.003067728 | 0.034923544 | Dennd4c |
| ENSMUSG00000024048.10 | 1379.197613 | -0.661537203 | 0.223524407 | -2.9595748 | 0.003080639 | 0.035041635 | Myl12a |
| ENSMUSG00000000915.11 | 2396.755965 | 0.616713116 | 0.208435038 | 2.958778527 | 0.00308861 | 0.035087858 | Hip1r |
| ENSMUSG00000041064.9 | 192.8365811 | 0.780059165 | 0.263652739 | 2.958661336 | 0.003089785 | 0.035087858 | Pif1 |
| ENSMUSG00000034892.8 | 1816.119984 | -0.878465285 | 0.297031288 | -2.957484011 | 0.003101608 | 0.035193188 | Rps29 |
| ENSMUSG00000038717.7 | 297.1347164 | -0.997425697 | 0.337294886 | -2.957132576 | 0.003105146 | 0.035204399 | Atp5l |
| ENSMUSG00000028420.9 | 91.30669116 | 0.931891839 | 0.315201813 | 2.956492638 | 0.003111597 | 0.035248595 | Tmem38b |
| ENSMUSG00000015619.10 | 587.5490643 | -0.863843019 | 0.292236922 | -2.955968095 | 0.003116893 | 0.035279656 | Gata3 |
| ENSMUSG00000022094.11 | 215.8544726 | -0.701402988 | 0.237322087 | -2.955489722 | 0.003121731 | 0.035305473 | Slc39a14 |
| ENSMUSG00000000530.12 | 36.88579328 | 1.525753853 | 0.516749899 | 2.952596326 | 0.003151137 | 0.035608884 | Acvrl1 |
| ENSMUSG00000059839.8 | 92.52044308 | -0.8945835 | 0.30303343 | -2.952095085 | 0.003156257 | 0.035637577 | Zfp874b |
| ENSMUSG00000024248.9 | 1036.578521 | -0.62598147 | 0.212149127 | -2.950667204 | 0.003170884 | 0.035773474 | Cox7a2l |
| ENSMUSG00000029238.8 | 478.8779476 | -0.681160273 | 0.230902487 | -2.949991063 | 0.003177831 | 0.035804245 | Clock |
| ENSMUSG00000052144.6 | 498.078397 | -0.659645037 | 0.22361628 | -2.949897199 | 0.003178797 | 0.035804245 | Ppp4r2 |
| ENSMUSG00000031578.4 | 369.346246 | -0.772553012 | 0.261988029 | -2.948810358 | 0.003189997 | 0.035901111 | Mak16 |
| ENSMUSG00000002033.9 | 1882.392707 | -0.701147054 | 0.237853623 | -2.947809021 | 0.003200347 | 0.035988268 | Cd3g |
| ENSMUSG00000004668.10 | 927.1981418 | 1.526382726 | 0.517876716 | 2.947386276 | 0.003204726 | 0.036008187 | Abca13 |
| ENSMUSG00000031216.9 | 24.50039538 | 1.432767617 | 0.486533267 | 2.94485026 | 0.00323111 | 0.036274429 | Stard8 |
| ENSMUSG00000071350.8 | 438.2380699 | -0.573380397 | 0.194722368 | -2.944604685 | 0.003233675 | 0.036274429 | Setdb2 |
| ENSMUSG00000041926.11 | 417.3303183 | 0.696042487 | 0.236432091 | 2.943942532 | 0.003240602 | 0.03632262 | Rnpep |
| ENSMUSG00000037306.9 | 113.6479745 | 0.734233945 | 0.249683913 | 2.940653795 | 0.003275204 | 0.036680687 | Man1c1 |
| ENSMUSG00000030677.7 | 414.155814 | 0.563444419 | 0.191767139 | 2.938169821 | 0.003301561 | 0.036945915 | Kif22 |
| ENSMUSG00000024339.8 | 2331.203789 | 0.629639763 | 0.214331573 | 2.937690201 | 0.003306673 | 0.036973152 | Tap2 |
| ENSMUSG00000025894.10 | 110.1012694 | -0.758805695 | 0.258519387 | -2.935198416 | 0.003333345 | 0.037156614 | Aasdhppt |
| ENSMUSG00000029135.9 | 275.0422747 | 1.019366075 | 0.34730115 | 2.935107115 | 0.003334326 | 0.037156614 | Fosl2 |
| ENSMUSG00000034987.3 | 240.1332558 | 0.804537042 | 0.274041099 | 2.935826219 | 0.003326606 | 0.037156614 | Hrh2 |

| | | | | | | | |
|---|---|---|---|---|---|---|---|
| ENSMUSG00000041695.2 | 27.69335565 | 1.427840201 | 0.486507066 | 2.934880705 | 0.003336759 | 0.037156614 | Kcnj2 |
| ENSMUSG00000047181.8 | 27.303165 | 1.524732158 | 0.519561576 | 2.9346515 | 0.003339225 | 0.037156614 | Samd14 |
| ENSMUSG00000079293.7 | 46.63711464 | 1.433134218 | 0.488301902 | 2.934934744 | 0.003336178 | 0.037156614 | Clec7a |
| ENSMUSG00000038578.11 | 50.83320882 | 1.403237923 | 0.478522177 | 2.932440731 | 0.003363091 | 0.037392049 | Susd1 |
| ENSMUSG00000033249.6 | 35.95322871 | 1.158925145 | 0.395327978 | 2.931553572 | 0.003372712 | 0.037468848 | Hsf4 |
| ENSMUSG00000026696.11 | 348.1063639 | -0.661878105 | 0.225832666 | -2.930834218 | 0.003380531 | 0.037525528 | Vamp4 |
| ENSMUSG00000027720.7 | 8.132677082 | -1.541808486 | 0.526385595 | -2.929047643 | 0.003400023 | 0.037681315 | Il2 |
| ENSMUSG00000038527.9 | 38.21948519 | 1.46011139 | 0.49845837 | 2.929254433 | 0.003397762 | 0.037681315 | C1rl |
| ENSMUSG00000020038.9 | 605.5475932 | -0.630222779 | 0.21522007 | -2.928271418 | 0.003408524 | 0.037684789 | Cry1 |
| ENSMUSG00000031861.11 | 69.3280393 | 0.935942019 | 0.319580129 | 2.928661493 | 0.003404249 | 0.037684789 | Lpar2 |
| ENSMUSG00000034664.9 | 54.01768967 | 0.958536582 | 0.327336675 | 2.928289603 | 0.003408324 | 0.037684789 | Itga2b |
| ENSMUSG00000022969.9 | 246.8156379 | 0.725418504 | 0.247817854 | 2.927224543 | 0.003420019 | 0.03778163 | Il10rb |
| ENSMUSG00000028896.9 | 336.4236751 | 0.557662468 | 0.190657847 | 2.92493845 | 0.003445244 | 0.037999496 | Rcc1 |
| ENSMUSG00000040044.7 | 603.0972122 | -0.800097832 | 0.273526473 | -2.925120276 | 0.003443231 | 0.037999496 | Orc3 |
| ENSMUSG00000016427.7 | 190.8917778 | -0.885621025 | 0.303087629 | -2.921996607 | 0.003477953 | 0.038299089 | Ndufa1 |
| ENSMUSG00000057858.7 | 256.8156641 | -0.792329375 | 0.271159667 | -2.922003053 | 0.003477881 | 0.038299089 | Fam204a |
| ENSMUSG00000041453.8 | 2704.736894 | -0.652348606 | 0.223353906 | -2.920694869 | 0.003492517 | 0.038428819 | Rpl21 |
| ENSMUSG00000035275.10 | 70.40285723 | -1.219068167 | 0.417590784 | -2.919288967 | 0.003508308 | 0.03857184 | Raver2 |
| ENSMUSG00000037904.10 | 46.28689201 | 1.006639498 | 0.345068932 | 2.917212777 | 0.003531748 | 0.038737015 | Ankrd9 |
| ENSMUSG00000042726.10 | 847.4899755 | 0.585185504 | 0.200591806 | 2.91729516 | 0.003530815 | 0.038737015 | Trafd1 |
| ENSMUSG00000103753.1 | 34.55793452 | -1.136338451 | 0.38949798 | -2.917443762 | 0.003529133 | 0.038737015 | Gm6934 |
| ENSMUSG00000079845.4 | 150.5390202 | -1.057254787 | 0.362502413 | -2.916545513 | 0.003539311 | 0.038789161 | Xlr4a |
| ENSMUSG00000026313.11 | 813.388878 | 0.559246437 | 0.191775093 | 2.916157822 | 0.003543712 | 0.038806597 | Hdac4 |
| ENSMUSG00000092517.1 | 87.28788834 | 0.817458261 | 0.280350819 | 2.915840459 | 0.003547318 | 0.038815309 | Art2a-ps |
| ENSMUSG00000026981.11 | 22.14192477 | 1.465266264 | 0.502612507 | 2.915300046 | 0.003553467 | 0.038851805 | Il1rn |
| ENSMUSG00000049044.12 | 65.57900289 | -1.355093018 | 0.464898914 | -2.914812183 | 0.003559027 | 0.038881802 | Rapgef4 |
| ENSMUSG00000076499.3 | 167.7478362 | 1.386357109 | 0.475788465 | 2.913809835 | 0.003570473 | 0.03897602 | Trbv31 |
| ENSMUSG00000033318.6 | 134.2304427 | -0.699505141 | 0.240099736 | -2.913394049 | 0.003575231 | 0.038997133 | Gstt2 |
| ENSMUSG00000031197.7 | 402.353548 | -0.82410605 | 0.282962808 | -2.912418268 | 0.00358642 | 0.039088303 | Vbp1 |
| ENSMUSG00000005410.5 | 1264.242832 | 0.60576836 | 0.208149364 | 2.910258033 | 0.003611305 | 0.039328476 | Mcm5 |
| ENSMUSG00000024516.8 | 891.1915201 | -0.538643693 | 0.185179179 | -2.908770289 | 0.003628534 | 0.039403314 | Sec11c |
| ENSMUSG00000024589.11 | 17.40980877 | 1.320676645 | 0.454036391 | 2.908746239 | 0.003628813 | 0.039403314 | Nedd4l |
| ENSMUSG00000056515.8 | 106.1688269 | 1.271283104 | 0.437065417 | 2.908679237 | 0.003629591 | 0.039403314 | Rab31 |
| ENSMUSG00000072235.5 | 554.6798426 | -0.943434075 | 0.324305869 | -2.909087267 | 0.003624857 | 0.039403314 | Tuba1a |
| ENSMUSG00000025362.5 | 1196.664306 | -0.996610942 | 0.342740048 | -2.907774994 | 0.003640101 | 0.039486379 | Rps26 |
| ENSMUSG00000030291.8 | 79.95535063 | -0.908235325 | 0.312476745 | -2.90656933 | 0.003654159 | 0.039607758 | Med21 |
| ENSMUSG00000015314.6 | 972.0206177 | 0.56447936 | 0.194263824 | 2.905735869 | 0.003663906 | 0.039682256 | Slamf6 |
| ENSMUSG00000041515.5 | 472.1422312 | 1.13449547 | 0.390541455 | 2.904929693 | 0.003673356 | 0.039753427 | Irf8 |
| ENSMUSG00000030468.8 | 27.39592592 | 1.449681651 | 0.499308237 | 2.903380207 | 0.003691582 | 0.039919382 | Siglecg |
| ENSMUSG00000062328.7 | 3277.757203 | -0.720971947 | 0.248517472 | -2.901091594 | 0.003718652 | 0.040149227 | Rpl17 |
| ENSMUSG00000064356.3 | 2578.77831 | -0.732465286 | 0.252465957 | -2.901243778 | 0.003716846 | 0.040149227 | mt-Atp8 |
| ENSMUSG00000033508.6 | 64.16698404 | 1.361852638 | 0.46957253 | 2.900196565 | 0.003729287 | 0.040201191 | Asprv1 |
| ENSMUSG00000065126.1 | 103.1026316 | -1.316175533 | 0.453812819 | -2.900260804 | 0.003728523 | 0.040201191 | Snord104 |
| ENSMUSG00000046179.13 | 463.3035081 | 0.762614473 | 0.262993064 | 2.899751274 | 0.003734589 | 0.040226939 | E2f8 |
| ENSMUSG00000019214.9 | 154.3649969 | 0.813125992 | 0.280649604 | 2.897299629 | 0.003763901 | 0.040511073 | Chtf18 |
| ENSMUSG00000061684.5 | 36.71227617 | -1.224864494 | 0.422887838 | -2.896428753 | 0.003774363 | 0.040592043 | Rpl21-ps8 |
| ENSMUSG00000063849.5 | 185.7863362 | 0.85636307 | 0.295820487 | 2.894874112 | 0.003793106 | 0.04076187 | Ppcdc |
| ENSMUSG00000062075.9 | 246.8165484 | 0.673081527 | 0.232570341 | 2.894098727 | 0.003802486 | 0.040828062 | Lmnb2 |
| ENSMUSG00000087141.1 | 426.7480163 | -0.597101655 | 0.206332811 | -2.893876415 | 0.003805179 | 0.040828062 | Plcxd2 |
| ENSMUSG00000020303.2 | 40.6342108 | -1.203916282 | 0.416082134 | -2.893458244 | 0.003810249 | 0.040831916 | Stc2 |
| ENSMUSG00000060314.8 | 29.00933349 | -1.13205767 | 0.39126068 | -2.893359152 | 0.003811452 | 0.040831916 | Zfp941 |
| ENSMUSG00000063952.11 | 453.918532 | 0.969698341 | 0.335284862 | 2.892162612 | 0.003825999 | 0.040955981 | Brpf3 |
| ENSMUSG00000031387.10 | 16.75373028 | 1.330849284 | 0.4602286 | 2.891713562 | 0.003831471 | 0.04098279 | Renbp |
| ENSMUSG00000021824.8 | 447.6645334 | -0.664491122 | 0.229861596 | -2.890831409 | 0.003842242 | 0.041034429 | Ap3m1 |
| ENSMUSG00000038482.10 | 682.9946177 | 0.586820825 | 0.202992713 | 2.890846751 | 0.003842054 | 0.041034429 | Tfdp1 |
| ENSMUSG00000026489.9 | 73.25373621 | 0.881625258 | 0.305191949 | 2.888756602 | 0.003867683 | 0.041274217 | Adck3 |
| ENSMUSG00000063694.5 | 283.9245212 | -0.788963221 | 0.273484899 | -2.884851131 | 0.003915988 | 0.041757434 | Cycs |
| ENSMUSG00000017765.12 | 183.2145612 | 0.771644047 | 0.267505846 | 2.884587601 | 0.003919267 | 0.041760153 | Slc12a4 |
| ENSMUSG00000063406.7 | 727.081963 | -0.692350594 | 0.24018229 | -2.88260469 | 0.003944021 | 0.041991503 | Tmed5 |
| ENSMUSG00000018427.7 | 25.99899334 | 1.287107332 | 0.446584546 | 2.882113459 | 0.003950175 | 0.042020692 | Ypel2 |
| ENSMUSG00000046841.4 | 306.7448506 | 1.527097628 | 0.529892598 | 2.881900284 | 0.003952848 | 0.042020692 | Ckap4 |
| ENSMUSG00000022438.6 | 16.77115945 | 1.46174947 | 0.507313562 | 2.881353035 | 0.003959718 | 0.042029018 | Parvb |

| | | | | | | | |
|---|---|---|---|---|---|---|---|
| ENSMUSG00000076776.2 | 8.922101607 | -1.514275998 | 0.525513874 | -2.881514785 | 0.003957687 | 0.042029018 | ENSMUSG00000076776 |
| ENSMUSG00000022024.9 | 514.8302267 | -0.648111826 | 0.224983876 | -2.880703437 | 0.003967888 | 0.042083383 | Sugt1 |
| ENSMUSG00000025790.10 | 826.3966616 | -0.90639267 | 0.314716381 | -2.880030159 | 0.003976371 | 0.042108675 | Slco3a1 |
| ENSMUSG00000040624.13 | 38.45499282 | 1.476677364 | 0.512691512 | 2.880245391 | 0.003973658 | 0.042108675 | Plekhg1 |
| ENSMUSG00000063358.11 | 1558.451966 | -0.556531725 | 0.193285216 | -2.879328984 | 0.003985224 | 0.04217008 | Mapk1 |
| ENSMUSG00000068329.8 | 140.2478237 | 0.672232027 | 0.233586084 | 2.877877039 | 0.004003612 | 0.042332215 | Htra2 |
| ENSMUSG00000029478.12 | 924.4851963 | 0.915892759 | 0.318338153 | 2.877106464 | 0.004013402 | 0.042403262 | Ncor2 |
| ENSMUSG00000054469.9 | 168.9206533 | 0.969158247 | 0.336891031 | 2.876770698 | 0.004017674 | 0.042415951 | Lclat1 |
| ENSMUSG00000023892.7 | 212.5675544 | -0.751139433 | 0.261171077 | -2.876043711 | 0.004026939 | 0.042425776 | Zfp51 |
| ENSMUSG00000029915.10 | 87.21352216 | 1.430976754 | 0.497593518 | 2.875794604 | 0.004030119 | 0.042425776 | Clec5a |
| ENSMUSG00000064637.1 | 7.560754 | -1.531729552 | 0.532684089 | -2.875493345 | 0.004033966 | 0.042425776 | Snora20 |
| ENSMUSG00000069682.5 | 191.5611187 | -0.688268288 | 0.239305731 | -2.876104495 | 0.004026164 | 0.042425776 | Gm10275 |
| ENSMUSG00000069972.6 | 16.0161449 | -1.401684343 | 0.487426385 | -2.875684176 | 0.004031529 | 0.042425776 | Rps13-ps2 |
| ENSMUSG00000069206.9 | 69.39532889 | -0.905194699 | 0.314914621 | -2.874413054 | 0.004047792 | 0.042538782 | Zfp874a |
| ENSMUSG00000089706.3 | 11.33327988 | 1.501514386 | 0.522423623 | 2.874131873 | 0.004051398 | 0.042544296 | B230216N24Rik |
| ENSMUSG00000020257.8 | 1340.735191 | -0.508408392 | 0.176959772 | -2.873016768 | 0.004065725 | 0.042662311 | Wdr82 |
| ENSMUSG00000081769.4 | 231.2280618 | 1.13657824 | 0.395776143 | 2.871770469 | 0.004081793 | 0.042798392 | Gm12216 |
| ENSMUSG00000002996.13 | 499.4552653 | -0.851583146 | 0.296622214 | -2.870935172 | 0.004092594 | 0.042879087 | Hbp1 |
| ENSMUSG00000024663.13 | 14.14690892 | 1.51258975 | 0.526980159 | 2.870297346 | 0.00410086 | 0.042933109 | Rab3il1 |
| ENSMUSG00000021687.10 | 44.01862708 | 1.492382003 | 0.520324694 | 2.868174472 | 0.004128478 | 0.043156815 | Scamp1 |
| ENSMUSG00000051335.5 | 357.4843387 | 0.701822534 | 0.244688529 | 2.868228173 | 0.004127777 | 0.043156815 | Gfod1 |
| ENSMUSG00000028675.8 | 772.4599161 | -0.626914394 | 0.218608488 | -2.867749549 | 0.004134026 | 0.043182126 | Pnrc2 |
| ENSMUSG00000026977.13 | 1379.222774 | -0.561361601 | 0.195788664 | -2.867181329 | 0.004141457 | 0.04319439 | March7 |
| ENSMUSG00000035085.5 | 23.63577701 | 1.336713975 | 0.466183553 | 2.867355499 | 0.004139178 | 0.04319439 | 1700020L24Rik |
| ENSMUSG00000053080.10 | 31.96322306 | 1.347729331 | 0.470117079 | 2.866795086 | 0.004146514 | 0.043214499 | 2700081O15Rik |
| ENSMUSG00000021278.6 | 12.05387549 | 1.50543218 | 0.525318034 | 2.865753852 | 0.004160176 | 0.043324184 | Amn |
| ENSMUSG00000024287.7 | 344.1250372 | -0.750420643 | 0.26204262 | -2.863735083 | 0.00418678 | 0.043568385 | Thoc1 |
| ENSMUSG00000021537.8 | 165.9155131 | -0.950045888 | 0.331826665 | -2.863078792 | 0.004195462 | 0.043625856 | Cetn3 |
| ENSMUSG00000003031.10 | 1124.488556 | -0.618006122 | 0.215981702 | -2.861381853 | 0.004217987 | 0.043827073 | Cdkn1b |
| ENSMUSG00000052298.8 | 1399.125325 | -0.557161351 | 0.194763189 | -2.860711785 | 0.004226911 | 0.04388678 | Cdc42se2 |
| ENSMUSG00000029866.9 | 13.64194618 | 1.473929606 | 0.515434889 | 2.859584476 | 0.004241964 | 0.04400998 | Kel |
| ENSMUSG00000029814.8 | 132.3328622 | -0.759087964 | 0.265554009 | -2.858506893 | 0.004256399 | 0.044126583 | Igf2bp3 |
| ENSMUSG00000022574.6 | 24.43416129 | 1.164982821 | 0.407599014 | 2.858159079 | 0.004261067 | 0.044141843 | Naprt |
| ENSMUSG00000066362.5 | 42.35637733 | -1.115677865 | 0.390503234 | -2.857025929 | 0.004276309 | 0.04426653 | Rps13-ps1 |
| ENSMUSG00000020490.12 | 16.89629951 | 1.460413501 | 0.511423342 | 2.855586325 | 0.004295744 | 0.044429088 | Btnl10 |
| ENSMUSG00000035692.6 | 57.01845943 | 1.172960029 | 0.410788523 | 2.855386564 | 0.004298448 | 0.044429088 | Isg15 |
| ENSMUSG00000054582.4 | 26.74591627 | 1.181101077 | 0.413717642 | 2.854848225 | 0.00430574 | 0.044466065 | Pabpc1l |
| ENSMUSG00000060510.9 | 495.9392812 | -0.550773273 | 0.19293917 | -2.854647266 | 0.004308465 | 0.044466065 | Zfp266 |
| ENSMUSG00000066440.4 | 737.0103519 | 0.608118443 | 0.213127532 | 2.853307768 | 0.00432667 | 0.044620598 | Zfyve26 |
| ENSMUSG00000095892.1 | 17.77272743 | -1.349201202 | 0.473228115 | -2.851058844 | 0.00435739 | 0.044903884 | Rnu5g |
| ENSMUSG00000031377.7 | 21.07146118 | 1.447454621 | 0.507983726 | 2.849411402 | 0.00438002 | 0.04510343 | Bmx |
| ENSMUSG00000005986.11 | 104.5965094 | 1.092647403 | 0.383673788 | 2.847855227 | 0.004401494 | 0.045290785 | Ankrd13d |
| ENSMUSG00000018774.9 | 73.67990033 | 1.018820965 | 0.35785299 | 2.84703773 | 0.004412813 | 0.045307085 | Cd68 |
| ENSMUSG00000030214.4 | 167.1382709 | 1.30349202 | 0.457842751 | 2.84702994 | 0.004412921 | 0.045307085 | Plbd1 |
| ENSMUSG00000051256.9 | 84.50688146 | -0.849546122 | 0.298372702 | -2.847264899 | 0.004409665 | 0.045307085 | Jagn1 |
| ENSMUSG00000028438.12 | 132.0899159 | 0.717124225 | 0.251926547 | 2.846560766 | 0.004419429 | 0.045340193 | Kif24 |
| ENSMUSG00000028086.10 | 610.9117133 | -0.54280894 | 0.190706433 | -2.846306399 | 0.004422961 | 0.045342743 | Fbxw7 |
| ENSMUSG00000047880.10 | 60.16300328 | 1.078215945 | 0.378858149 | 2.845962128 | 0.004427746 | 0.04535812 | Cxcr5 |
| ENSMUSG00000034570.8 | 15.64801085 | 1.510058658 | 0.530718108 | 2.84531211 | 0.004436793 | 0.045417101 | Inpp5j |
| ENSMUSG00000020605.8 | 27.21981317 | 1.127229746 | 0.396393085 | 2.843716975 | 0.004459064 | 0.045611268 | Hs1bp3 |
| ENSMUSG00000029254.12 | 402.4549313 | -0.883534288 | 0.310813123 | -2.842654391 | 0.004473956 | 0.045729721 | Stap1 |
| ENSMUSG00000047945.6 | 175.7880113 | 0.905273558 | 0.318689686 | 2.84061141 | 0.004502714 | 0.045989631 | Marcksl1 |
| ENSMUSG00000028644.12 | 73.45680093 | 1.419973055 | 0.500054801 | 2.839634882 | 0.00451652 | 0.046062496 | Ermap |
| ENSMUSG00000071229.3 | 10.70823345 | 1.431792615 | 0.504209767 | 2.839676479 | 0.004515931 | 0.046062496 | Timm8a2 |
| ENSMUSG00000001175.9 | 5222.722995 | -0.661640398 | 0.23304828 | -2.839070076 | 0.004524522 | 0.046110054 | Calm1 |
| ENSMUSG00000045733.7 | 15.32991886 | -1.390863433 | 0.490278688 | -2.836883321 | 0.004555626 | 0.046392799 | Sprn |
| ENSMUSG00000022822.11 | 399.7066428 | 0.767696761 | 0.27074571 | 2.835490028 | 0.004575544 | 0.046527019 | Abcc5 |
| ENSMUSG00000051124.6 | 548.9812584 | -0.597524399 | 0.210715754 | -2.835689259 | 0.004572691 | 0.046527019 | Gimap9 |
| ENSMUSG00000050921.8 | 794.6218672 | -0.821293176 | 0.289697612 | -2.835001541 | 0.004582546 | 0.046563932 | P2ry10 |
| ENSMUSG00000015947.6 | 25.08573044 | 1.304042697 | 0.460044702 | 2.834599969 | 0.00458831 | 0.046588215 | Fcgr1 |
| ENSMUSG00000030340.12 | 8.217752562 | 1.516608726 | 0.535391262 | 2.832711014 | 0.004615509 | 0.046761237 | Scnn1a |
| ENSMUSG00000039187.12 | 189.4819099 | 0.642796533 | 0.226882687 | 2.833166964 | 0.004608931 | 0.046761237 | Fanci |

| | | | | | | | |
|---|---|---|---|---|---|---|---|
| ENSMUSG00000056888.7 | 610.6645521 | -0.789987932 | 0.278864254 | -2.832876287 | 0.004613124 | 0.046761237 | Glipr1 |
| ENSMUSG00000014504.12 | 261.3459955 | -0.661142504 | 0.233507156 | -2.831358638 | 0.004635072 | 0.046864528 | Srp19 |
| ENSMUSG00000015597.12 | 258.2521482 | -0.727478702 | 0.256962599 | -2.83106843 | 0.004639279 | 0.046864528 | Zfp318 |
| ENSMUSG00000024867.10 | 16.52811681 | 1.442413225 | 0.509385041 | 2.831675666 | 0.004630479 | 0.046864528 | Pip5k1b |
| ENSMUSG00000076487.1 | 72.12900938 | 1.091745924 | 0.38562835 | 2.831083149 | 0.004639066 | 0.046864528 | Trbj1-5 |
| ENSMUSG00000089844.3 | 12.61186401 | 1.392079511 | 0.492175194 | 2.828422739 | 0.004677799 | 0.047219099 | A530032D15Rik |
| ENSMUSG00000102727.1 | 4.936626569 | 1.506677961 | 0.532867297 | 2.827491894 | 0.00469142 | 0.047322004 | Gm37938 |
| ENSMUSG00000038412.7 | 170.4648055 | -0.769763221 | 0.272367264 | -2.826195808 | 0.004710446 | 0.047410021 | Higd1a |
| ENSMUSG00000051682.11 | 20.36834886 | 1.409832366 | 0.498813457 | 2.826371957 | 0.004707856 | 0.047410021 | Treml4 |
| ENSMUSG00000069873.3 | 65.85580998 | 1.447116983 | 0.512020794 | 2.826285574 | 0.004709126 | 0.047410021 | 4930438A08Rik |
| ENSMUSG00000020023.13 | 19.08037196 | 1.19063769 | 0.42134992 | 2.82576935 | 0.004716721 | 0.047438606 | Tmcc3 |
| ENSMUSG00000031370.8 | 323.1291537 | -0.582039104 | 0.206010763 | -2.825284932 | 0.004723859 | 0.047441285 | Zrsr2 |
| ENSMUSG00000038524.9 | 184.0429175 | 0.718409599 | 0.254260448 | 2.82548703 | 0.00472088 | 0.047441285 | Fchsd1 |
| ENSMUSG00000102752.1 | 23.02491579 | 1.487050194 | 0.526455676 | 2.82464462 | 0.004733308 | 0.047501638 | Gm7694 |
| ENSMUSG00000030142.6 | 52.62434345 | 1.34655059 | 0.476763309 | 2.824358678 | 0.004737534 | 0.047509514 | Clec4e |
| ENSMUSG00000004655.5 | 68.43171962 | 1.397401748 | 0.494812494 | 2.824103606 | 0.004741306 | 0.047512837 | Aqp1 |
| ENSMUSG00000024429.9 | 354.7975489 | 0.574405617 | 0.203461214 | 2.823170109 | 0.004755133 | 0.047616851 | Gnl1 |
| ENSMUSG00000073386.4 | 16.32512962 | 1.472360251 | 0.521653952 | 2.822484613 | 0.004765311 | 0.047684187 | 9830107B12Rik |
| ENSMUSG00000020897.8 | 388.9949851 | 0.628339107 | 0.222659087 | 2.821978277 | 0.004772841 | 0.047724954 | Aurkb |
| ENSMUSG00000020873.9 | 335.0516286 | -0.674098338 | 0.238914133 | -2.821508835 | 0.004779832 | 0.047760276 | Slc35b1 |
| ENSMUSG00000025702.11 | 37.85879768 | 1.38494786 | 0.49140728 | 2.818329961 | 0.004827417 | 0.048200871 | March8 |
| ENSMUSG00000037944.8 | 44.46541594 | -1.036502687 | 0.367859348 | -2.817660316 | 0.004837496 | 0.048266604 | Ccr7 |
| ENSMUSG00000029275.13 | 372.5543033 | 0.717160537 | 0.254615319 | 2.816643312 | 0.004852839 | 0.048384729 | Gfi1 |
| ENSMUSG00000100147.2 | 15.25519596 | 1.368240842 | 0.48604713 | 2.815037384 | 0.004877156 | 0.048592098 | 1700047M11Rik |
| ENSMUSG00000030103.7 | 4993.270812 | -0.66349537 | 0.235736656 | -2.814561729 | 0.00488438 | 0.048628982 | Bhlhe40 |
| ENSMUSG00000050856.12 | 259.2238288 | -0.85937709 | 0.305480888 | -2.813194286 | 0.0049052 | 0.048765956 | Atp5k |
| ENSMUSG00000051439.6 | 19.74552641 | 1.434243856 | 0.509819791 | 2.813236917 | 0.00490455 | 0.048765956 | Cd14 |
| ENSMUSG00000075590.2 | 24.62736445 | 1.186565869 | 0.421827183 | 2.812919403 | 0.004909395 | 0.048772548 | Nrbp2 |
| ENSMUSG00000002365.9 | 101.7263533 | 1.04287556 | 0.370810077 | 2.812425078 | 0.004916948 | 0.048812459 | Snx9 |
| ENSMUSG00000095415.1 | 7.527446449 | -1.507903977 | 0.536598605 | -2.810115349 | 0.004952375 | 0.049128838 | ENSMUSG00000095415 |
| ENSMUSG00000065037.1 | 87541.81559 | -0.693615079 | 0.246856074 | -2.809795469 | 0.004957299 | 0.049142387 | Rn7sk |
| ENSMUSG00000038128.6 | 1822.227986 | -0.629501878 | 0.22438031 | -2.805513003 | 0.005023655 | 0.049764461 | Camk4 |
| ENSMUSG00000037720.12 | 377.4874647 | -0.631606191 | 0.225225828 | -2.804323986 | 0.005042221 | 0.049912566 | Tmem33 |

| Comparison 2: EOMES$^{+}$ GFP$^{+}$ of *Eomes* $^{+/GFP}$ reporter mice vs EOMES$^{-/-}$ GFP$^{+}$ of *Eomes* $^{\Delta T/GFP}$ knock-out mice | | | | | | | |
|---|---|---|---|---|---|---|---|
| ENSEMBL ID | baseMean | log2FoldChange | lfcSE | stat | pvalue | padj | gene_names |
| ENSMUSG00000049410.8 | 167.9794509 | 4.003151195 | 0.29408753 | 13.6121079 | 3.39E-42 | 4.51E-38 | Zfp683 |
| ENSMUSG00000012123.11 | 135.7804192 | 2.919323967 | 0.256609707 | 11.37651417 | 5.47E-30 | 3.64E-26 | Aim1l |
| ENSMUSG00000057329.7 | 445.6630045 | -1.485237959 | 0.203061917 | -7.314212252 | 2.59E-13 | 1.15E-09 | Bcl2 |
| ENSMUSG00000018341.8 | 509.8425585 | 1.650913711 | 0.228181829 | 7.235079676 | 4.65E-13 | 1.55E-09 | Il12rb2 |
| ENSMUSG00000022900.10 | 86.72594163 | -2.055325605 | 0.293843648 | -6.994623237 | 2.66E-12 | 7.07E-09 | Ildr1 |
| ENSMUSG00000033066.11 | 308.1870303 | -1.536357584 | 0.223701766 | -6.867883118 | 6.52E-12 | 1.44E-08 | Gas7 |
| ENSMUSG00000074570.9 | 114.6363271 | 1.90641372 | 0.284926699 | 6.690891813 | 2.22E-11 | 4.21E-08 | Cass4 |
| ENSMUSG00000053113.3 | 297.5130136 | -1.692289387 | 0.269613283 | -6.276728538 | 3.46E-10 | 5.75E-07 | Socs3 |
| ENSMUSG00000030154.8 | 186.330221 | 1.564251071 | 0.267417462 | 5.84947242 | 4.93E-09 | 6.56E-06 | Klrb1f |
| ENSMUSG00000054672.8 | 33.17914872 | -1.860042733 | 0.317655825 | -5.855528494 | 4.75E-09 | 6.56E-06 | 5830411N06Rik |
| ENSMUSG00000006154.9 | 50.17569421 | -2.094140815 | 0.359452202 | -5.825922897 | 5.68E-09 | 6.87E-06 | Eps8l1 |
| ENSMUSG00000015709.8 | 80.46451577 | 1.598124694 | 0.276585238 | 5.778054921 | 7.56E-09 | 8.38E-06 | Arnt2 |
| ENSMUSG00000004633.13 | 181.4032358 | 1.447624608 | 0.251679105 | 5.751866482 | 8.83E-09 | 8.39E-06 | Chn2 |
| ENSMUSG00000037849.7 | 88.68125978 | -1.737124138 | 0.301872163 | -5.754502565 | 8.69E-09 | 8.39E-06 | Gm4955 |
| ENSMUSG00000030257.12 | 48.66161243 | -1.949886904 | 0.342802773 | -5.688072136 | 1.28E-08 | 1.14E-05 | Srgap3 |
| ENSMUSG00000045087.8 | 160.2022929 | 1.993323547 | 0.357833342 | 5.570536098 | 2.54E-08 | 2.11E-05 | S1pr5 |
| ENSMUSG00000030653.12 | 273.9528858 | -1.481918408 | 0.272267549 | -5.442875641 | 5.24E-08 | 4.10E-05 | Pde2a |
| ENSMUSG00000025348.8 | 27.30647216 | -1.823762053 | 0.344344632 | -5.296327813 | 1.18E-07 | 8.73E-05 | Itga7 |
| ENSMUSG00000020183.7 | 109.5423983 | -1.608258899 | 0.323365033 | -4.973508989 | 6.58E-07 | 0.000460262 | Cpm |
| ENSMUSG00000032446.10 | 411.7472269 | -1.180419956 | 0.241057198 | -4.896845924 | 9.74E-07 | 0.000647624 | Eomes |
| ENSMUSG00000096768.3 | 120.1948651 | 1.516266914 | 0.314898658 | 4.815094874 | 1.47E-06 | 0.000931823 | Erdr1 |
| ENSMUSG00000039410.12 | 29.81749815 | 1.583065859 | 0.329646104 | 4.802319331 | 1.57E-06 | 0.000948159 | Prdm16 |
| ENSMUSG00000075014.1 | 252.9337321 | 1.707002637 | 0.359244642 | 4.75164397 | 2.02E-06 | 0.001166754 | Gm10800 |
| ENSMUSG00000097312.1 | 248.3631448 | 1.689994318 | 0.359350541 | 4.702912965 | 2.56E-06 | 0.001421303 | Gm26870 |
| ENSMUSG00000032841.11 | 48.24454661 | -1.438340213 | 0.307550385 | -4.676762836 | 2.91E-06 | 0.001503183 | Prr5l |
| ENSMUSG00000087670.1 | 11.47904302 | 1.680479597 | 0.3594556 | 4.675068627 | 2.94E-06 | 0.001503183 | 9530036M11Rik |
| ENSMUSG00000021699.13 | 384.6570768 | -0.961334231 | 0.213035439 | -4.512555442 | 6.41E-06 | 0.003155114 | Pde4d |
| ENSMUSG00000033910.9 | 21.42831967 | 1.610740113 | 0.358842063 | 4.48871601 | 7.17E-06 | 0.003403553 | Gucy1a3 |
| ENSMUSG00000025997.9 | 510.104943 | -1.379672313 | 0.312440136 | -4.415797311 | 1.01E-05 | 0.004615483 | Ikzf2 |
| ENSMUSG00000020593.10 | 470.8420509 | -1.252062094 | 0.28416786 | -4.406065116 | 1.05E-05 | 0.004666763 | Lpin1 |
| ENSMUSG00000049608.8 | 26.88636698 | -1.563769401 | 0.359507155 | -4.349758768 | 1.36E-05 | 0.005847168 | Gpr55 |
| ENSMUSG00000041272.7 | 506.6761916 | -1.337575238 | 0.308746793 | -4.332272492 | 1.48E-05 | 0.006133716 | Tox |
| ENSMUSG00000001270.8 | 54.92700246 | 1.303371867 | 0.302022475 | 4.315479729 | 1.59E-05 | 0.006418526 | Ckb |
| ENSMUSG00000026872.12 | 907.1540831 | 1.160949898 | 0.269901957 | 4.301376363 | 1.70E-05 | 0.006639848 | Zeb2 |
| ENSMUSG00000004933.13 | 49.11386905 | 1.292085189 | 0.306418078 | 4.216739429 | 2.48E-05 | 0.009418684 | Matk |
| ENSMUSG00000024222.12 | 873.727804 | -0.843887905 | 0.202774338 | -4.16170957 | 3.16E-05 | 0.011669787 | Fkbp5 |
| ENSMUSG00000032221.10 | 45.0506928 | 1.267212375 | 0.305775393 | 4.144258838 | 3.41E-05 | 0.012254495 | Mns1 |
| ENSMUSG00000096385.3 | 34.25072515 | 1.472433707 | 0.356559063 | 4.129564663 | 3.63E-05 | 0.012720777 | Gm11168 |
| ENSMUSG00000046743.6 | 42.46299191 | 1.471214271 | 0.357426931 | 4.116125966 | 3.85E-05 | 0.013139506 | Fat4 |
| ENSMUSG00000017466.5 | 132.3016403 | 0.97615961 | 0.237924214 | 4.102817415 | 4.08E-05 | 0.013570966 | Timp2 |
| ENSMUSG00000078247.3 | 50.02460085 | -1.373090914 | 0.338525498 | -4.056093036 | 4.99E-05 | 0.0161872 | Airn |
| ENSMUSG00000025809.11 | 2110.515109 | 1.363138958 | 0.336962498 | 4.045372894 | 5.22E-05 | 0.016542636 | Itgb1 |
| ENSMUSG00000032356.8 | 29.30002967 | 1.444081898 | 0.359448687 | 4.017491093 | 5.88E-05 | 0.017006958 | Rasgrf1 |
| ENSMUSG00000040613.10 | 105.2613111 | -1.005065757 | 0.249488668 | -4.028502646 | 5.61E-05 | 0.017006958 | Apobec1 |
| ENSMUSG00000047281.3 | 155.1033698 | -1.074189552 | 0.267187554 | -4.020357737 | 5.81E-05 | 0.017006958 | Sfn |
| ENSMUSG00000097796.1 | 38.39427781 | -1.314306963 | 0.326812189 | -4.021597136 | 5.78E-05 | 0.017006958 | Gm16702 |
| ENSMUSG00000038354.9 | 19.63872184 | -1.352289559 | 0.340620982 | -3.970071222 | 7.19E-05 | 0.020332347 | Ankrd35 |
| ENSMUSG00000076479.3 | 15.64167048 | 1.349295364 | 0.34247827 | 3.939798467 | 8.16E-05 | 0.02259617 | Trbv26 |
| ENSMUSG00000016529.5 | 46.6369336 | -1.401190375 | 0.356151129 | -3.934257851 | 8.35E-05 | 0.022651842 | Il10 |
| ENSMUSG00000035275.10 | 59.66831142 | 1.369486825 | 0.351736411 | 3.89350315 | 9.88E-05 | 0.025351715 | Raver2 |
| ENSMUSG00000066026.10 | 90.44649663 | 1.020933332 | 0.262266181 | 3.892737249 | 9.91E-05 | 0.025351715 | Dhrs3 |
| ENSMUSG00000100815.1 | 38.11054796 | -1.343460533 | 0.344893772 | -3.895287893 | 9.81E-05 | 0.025351715 | Gm29112 |
| ENSMUSG00000026447.12 | 90.4055114 | 1.173548021 | 0.30196752 | 3.886338579 | 0.000101768 | 0.025537893 | Pik3c2b |
| ENSMUSG00000027073.5 | 16.82554881 | -1.366709487 | 0.352493563 | -3.877260841 | 0.000105639 | 0.026018515 | Prg2 |
| ENSMUSG00000003545.2 | 403.3259029 | 1.063274 | 0.275692103 | 3.856744498 | 0.000114907 | 0.027434376 | Fosb |
| ENSMUSG00000073491.6 | 69.68480401 | -1.378448742 | 0.357531775 | -3.855457996 | 0.000115513 | 0.027434376 | Pydc4 |

| | | | | | | | |
|---|---|---|---|---|---|---|---|
| ENSMUSG00000024401.10 | 337.2962545 | 1.122989583 | 0.29198289 | 3.846080101 | 0.000120023 | 0.027713121 | Tnf |
| ENSMUSG00000027546.11 | 70.53995973 | 1.199434896 | 0.312269314 | 3.84102709 | 0.000122521 | 0.027713121 | Atp9a |
| ENSMUSG00000047180.8 | 52.05629866 | -1.209198288 | 0.31495088 | -3.839323413 | 0.000123374 | 0.027713121 | Neurl3 |
| ENSMUSG00000047898.6 | 14.45734607 | -1.378723483 | 0.359410925 | -3.836064477 | 0.000125022 | 0.027713121 | Ccr4 |
| ENSMUSG00000015133.12 | 396.4379997 | -0.956911083 | 0.250676151 | -3.817319999 | 0.000134909 | 0.029414617 | Lrrk1 |
| ENSMUSG00000102550.1 | 9.844640054 | 1.357873689 | 0.357028919 | 3.803259671 | 0.000142805 | 0.030633875 | Gm8276 |
| ENSMUSG00000083950.1 | 17.25020726 | -1.35965309 | 0.359182997 | -3.78540494 | 0.000153458 | 0.032396792 | Gm14466 |
| ENSMUSG00000040537.13 | 24.13313039 | 1.267360266 | 0.336876234 | 3.762094621 | 0.000168496 | 0.034575069 | Adam22 |
| ENSMUSG00000074604.5 | 69.01995489 | -1.076393086 | 0.286169436 | -3.761383822 | 0.000168976 | 0.034575069 | Mgst2 |
| ENSMUSG00000028927.6 | 245.9071501 | -1.088084441 | 0.29291604 | -3.714663228 | 0.000203474 | 0.040033111 | Padi2 |
| ENSMUSG00000030365.7 | 235.0289459 | 0.831604963 | 0.224622562 | 3.702232553 | 0.000213711 | 0.040033111 | Clec2i |
| ENSMUSG00000038146.7 | 49.46809536 | 1.307916698 | 0.35312189 | 3.70386752 | 0.000212337 | 0.040033111 | Notch3 |
| ENSMUSG00000047586.3 | 21.0829035 | -1.283824142 | 0.346155306 | -3.708809659 | 0.000208236 | 0.040033111 | Nccrp1 |
| ENSMUSG00000075015.3 | 17.53866959 | 1.311159928 | 0.353787217 | 3.70606926 | 0.000210501 | 0.040033111 | Gm10801 |
| ENSMUSG00000086513.3 | 44.26020244 | -1.311107503 | 0.353085515 | -3.713286014 | 0.000204585 | 0.040033111 | 9130208D14Rik |
| ENSMUSG00000076490.2 | 826.5049948 | -1.143278877 | 0.309389595 | -3.695272542 | 0.000219651 | 0.040574429 | Trbc1 |
| ENSMUSG00000041762.12 | 77.2855414 | -0.947738153 | 0.258131524 | -3.671532014 | 0.000241101 | 0.043926597 | Gpr155 |
| ENSMUSG00000037868.11 | 183.0747048 | 1.15852037 | 0.31640582 | 3.66150145 | 0.000250741 | 0.044641567 | Egr2 |
| ENSMUSG00000061062.4 | 27.75667815 | 1.215087297 | 0.331947065 | 3.660485136 | 0.000251738 | 0.044641567 | Gm10093 |
| ENSMUSG00000079173.7 | 20.1115521 | 1.252312519 | 0.343568182 | 3.645018913 | 0.000267372 | 0.046790124 | Zan |
| ENSMUSG00000032265.10 | 200.0166249 | -0.919117698 | 0.253354941 | -3.627786744 | 0.000285861 | 0.049376029 | Fam46a |

| | Comparison 1 | | | | | Comparison 2 | | | | | |
|---|---|---|---|---|---|---|---|---|---|---|---|
| ENSEMBL ID | log2FoldChange | lfcSE | stat | pvalue | padj | log2FoldChange | lfcSE | stat | pvalue | padj | gene_names |
| ENSMUSG00000040537.13 | -1.388605758 | 0.360903237 | -3.847584661 | 0.000119288 | 0.003022475 | 1.267360266 | 0.336876234 | 3.762094621 | 0.000168496 | 0.034575069 | Adam22 |
| ENSMUSG00000012123.11 | -1.440487754 | 0.332397957 | -4.33362397 | 1.46675E-05 | 0.000580352 | 2.919323967 | 0.256609707 | 11.37651417 | 5.47E-30 | 3.64E-26 | Aim1l |
| ENSMUSG00000078247.3 | 2.135057137 | 0.419416893 | 5.090536821 | 3.57051E-07 | 2.59501E-05 | -1.373090914 | 0.338525498 | -4.056093036 | 4.99E-05 | 0.0161872 | Airn |
| ENSMUSG00000040613.10 | 1.592327262 | 0.279827805 | 5.690382555 | 1.26755E-08 | 1.54899E-06 | -1.005065757 | 0.249488668 | -4.028502646 | 5.61E-05 | 0.017006958 | Apobec1 |
| ENSMUSG00000015709.8 | -1.708286232 | 0.274201648 | -6.230036338 | 4.66327E-10 | 8.25578E-08 | 1.598124694 | 0.276585238 | 5.778054921 | 7.56E-09 | 8.38E-06 | Arnt2 |
| ENSMUSG00000074570.9 | -1.74457701 | 0.288632441 | -6.044285965 | 1.50073E-09 | 2.35495E-07 | 1.90641372 | 0.284926699 | 6.690891813 | 2.22E-11 | 4.21E-08 | Cass4 |
| ENSMUSG00000047898.6 | -1.720662892 | 0.394952805 | -4.35662912 | 1.32081E-05 | 0.000534594 | -1.378723483 | 0.359410925 | -3.836064477 | 0.000125022 | 0.027713121 | Ccr4 |
| ENSMUSG00000001270.8 | -2.730611721 | 0.279744392 | -9.761095469 | 1.65359E-22 | 2.28345E-19 | -2.730611721 | 0.279744392 | -9.761095469 | 1.65E-22 | 2.28E-19 | Ckb |
| ENSMUSG00000020183.7 | -2.131303995 | 0.269936239 | -7.89558305 | 2.88961E-15 | 1.70231E-12 | -1.608258899 | 0.323365033 | -4.973508989 | 6.58E-07 | 0.000460262 | Cpm |
| ENSMUSG00000066026.10 | -1.194342506 | 0.238065588 | -5.016863274 | 5.25219E-07 | 3.59047E-05 | 1.020933332 | 0.262266181 | 3.892737249 | 9.91E-05 | 0.025351715 | Dhrs3 |
| ENSMUSG00000032446.10 | 3.805099655 | 0.376946626 | 10.09453166 | 5.84092E-24 | 1.00822E-20 | 3.805099655 | 0.376946626 | 10.09453166 | 5.84E-24 | 1.01E-20 | Eomes |
| ENSMUSG00000046743.6 | 1.80162547 | 0.517099288 | 3.48409969 | 0.000493796 | 0.009055541 | 1.471214271 | 0.357426931 | 4.116125966 | 3.85E-05 | 0.013139506 | Fat4 |
| ENSMUSG00000024222.12 | 0.67360049 | 0.174034677 | 3.870495828 | 0.000108614 | 0.002819274 | -0.843887905 | 0.202774338 | -4.16170957 | 3.16E-05 | 0.011669787 | Fkbp5 |
| ENSMUSG00000003545.2 | 0.838000267 | 0.268722491 | 3.118459737 | 0.00181799 | 0.024279131 | 1.063274 | 0.275692103 | 3.856744498 | 0.000114907 | 0.027434376 | Fosb |
| ENSMUSG00000033066.11 | 1.645414623 | 0.195713565 | 8.407258976 | 4.19704E-17 | 3.21983E-14 | -1.536357584 | 0.223701766 | -6.867883118 | 6.52E-12 | 1.44E-08 | Gas7 |
| ENSMUSG00000083950.1 | 2.671189321 | 0.490843251 | 5.442041455 | 5.26734E-08 | 5.08648E-06 | -1.35965309 | 0.359182997 | -3.78540494 | 0.000153458 | 0.032396792 | Gm14466 |
| ENSMUSG00000097796.1 | 2.17601813 | 0.372892152 | 5.835516024 | 5.36243E-09 | 7.12018E-07 | -1.314306963 | 0.326812189 | -4.021597136 | 5.78E-05 | 0.017006958 | Gm16702 |
| ENSMUSG00000100815.1 | 2.661249313 | 0.418689379 | 6.356142404 | 2.06883E-10 | 3.96785E-08 | -1.343460533 | 0.344893772 | -3.895287893 | 9.81E-05 | 0.025351715 | Gm29112 |
| ENSMUSG00000049608.8 | -1.377229494 | 0.369765341 | -3.724604069 | 0.000195622 | 0.004452006 | -1.563769401 | 0.359507155 | -4.349758768 | 1.36E-05 | 0.005847168 | Gpr55 |
| ENSMUSG00000025997.9 | 2.402502984 | 0.335343205 | 7.16431092 | 7.81788E-13 | 2.51063E-10 | -1.379672313 | 0.312440136 | -4.415797311 | 1.01E-05 | 0.004615483 | Ikzf2 |
| ENSMUSG00000016529.5 | 1.609892836 | 0.532730929 | 3.0219624 | 0.002511417 | 0.030528313 | -1.401190375 | 0.356151129 | -3.934257851 | 8.35E-05 | 0.022651842 | Il10 |
| ENSMUSG00000022900.10 | 1.306197974 | 0.268983092 | 4.856059786 | 1.19745E-06 | 7.38193E-05 | -2.055325605 | 0.293843648 | -6.994623237 | 2.66E-12 | 7.07E-09 | Ildr1 |
| ENSMUSG00000025348.8 | 1.229094236 | 0.34806594 | 3.531210885 | 0.000413662 | 0.007857298 | -1.823762053 | 0.344344632 | -5.296327813 | 1.18E-07 | 8.73E-05 | Itga7 |
| ENSMUSG00000025809.11 | -1.701721465 | 0.352680231 | -4.825111576 | 1.39925E-06 | 8.47466E-05 | 1.363138958 | 0.336962498 | 4.045372894 | 5.22E-05 | 0.016542636 | Itgb1 |
| ENSMUSG00000020593.10 | 1.343555224 | 0.200349031 | 6.706072995 | 1.99932E-11 | 4.7601E-09 | -1.252062094 | 0.28416786 | -4.406065116 | 1.05E-05 | 0.004666763 | Lpin1 |
| ENSMUSG00000015133.12 | 2.226002211 | 0.282035199 | 7.892639701 | 2.95861E-15 | 1.70231E-12 | 2.226002211 | 0.282035199 | 7.892639701 | 2.96E-15 | 1.70E-12 | Lrrk1 |
| ENSMUSG00000074604.5 | 1.651090854 | 0.285381796 | 5.785550716 | 7.22752E-09 | 9.41554E-07 | -1.076393086 | 0.286169436 | -3.761383822 | 0.000168976 | 0.034575069 | Mgst2 |
| ENSMUSG00000028927.6 | 0.942287182 | 0.305426096 | 3.085156092 | 0.002034454 | 0.026107876 | -1.088084441 | 0.29291604 | -3.714663228 | 0.000203474 | 0.040033111 | Padi2 |
| ENSMUSG00000030653.12 | 1.277258155 | 0.228216271 | 5.596700663 | 2.18469E-08 | 2.37547E-06 | -1.481918408 | 0.272267549 | -5.442875641 | 5.24E-08 | 4.10E-05 | Pde2a |
| ENSMUSG00000027073.5 | 3.188574833 | 0.494270207 | 6.45107633 | 1.11059E-10 | 2.28897E-08 | -1.366709487 | 0.352493563 | -3.877260841 | 0.000105639 | 0.026018515 | Prg2 |
| ENSMUSG00000032841.11 | 1.512432352 | 0.358377324 | 4.220223352 | 2.4406E-05 | 0.000859753 | -1.438340213 | 0.307550385 | -4.676762836 | 2.91E-06 | 0.001503183 | Prr5l |
| ENSMUSG00000035275.10 | -1.219068167 | 0.417590784 | -2.919288967 | 0.003508308 | 0.03857184 | 1.369486825 | 0.351736411 | 3.89350315 | 9.88E-05 | 0.025351715 | Raver2 |
| ENSMUSG00000053113.3 | 0.799218853 | 0.243309714 | 3.284779883 | 0.001020621 | 0.015607698 | -1.692289387 | 0.269613283 | -6.276728538 | 3.46E-10 | 5.75E-07 | Socs3 |
| ENSMUSG00000017466.5 | -0.974314506 | 0.210643272 | -4.625424278 | 3.74E-06 | 0.000196283 | 0.97615961 | 0.237924214 | 4.102817415 | 4.08E-05 | 0.013570966 | Timp2 |
| ENSMUSG00000024401.10 | -0.587057808 | 0.194285198 | -3.021629101 | 0.002514184 | 0.030535064 | 1.122989583 | 0.29198289 | 3.846080101 | 0.000120023 | 0.027713121 | Tnf |
| ENSMUSG00000041272.7 | 3.79030092 | 0.387077992 | 9.792085826 | 1.22E-22 | 1.87E-19 | -1.337575238 | 0.308746793 | -4.332272492 | 1.48E-05 | 0.006133716 | Tox |
| ENSMUSG00000049410.8 | -2.392861254 | 0.388234495 | -6.163443188 | 7.12E-10 | 1.21E-07 | 4.003151195 | 0.29408753 | 13.6121079 | 3.39E-42 | 4.51E-38 | Zfp683 |

| Comparison 3: GFP+ vs GFP- of *Eomes* ΔT/GFP knock-out mice | | | | | | | |
|---|---|---|---|---|---|---|---|
| | baseMean | log2FoldChange | lfcSE | stat | pvalue | padj | gene_names |
| ENSMUSG00000029810.11 | 202.2869779 | -3.558747534 | 0.295223127 | -12.05443343 | 1.84E-33 | 2.29E-29 | Tmem176b |
| ENSMUSG00000023367.10 | 177.6119588 | -3.607329785 | 0.30905744 | -11.67203673 | 1.77E-31 | 1.10E-27 | Tmem176a |
| ENSMUSG00000025491.10 | 263.4325891 | -2.848056926 | 0.286206346 | -9.951061406 | 2.50E-23 | 1.04E-19 | Ifitm1 |
| ENSMUSG00000032446.10 | 202.0145646 | 3.077874663 | 0.312615021 | 9.8455751 | 7.16E-23 | 2.23E-19 | Eomes |
| ENSMUSG00000028150.10 | 99.47967111 | -3.480360782 | 0.357109744 | -9.745913803 | 1.92E-22 | 4.79E-19 | Rorc |
| ENSMUSG00000053965.6 | 137.8136253 | -2.809503864 | 0.28892849 | -9.72387274 | 2.39E-22 | 4.96E-19 | Pde5a |
| ENSMUSG00000060591.8 | 235.3156288 | -2.397661385 | 0.272076249 | -8.812461199 | 1.22E-18 | 2.18E-15 | Ifitm2 |
| ENSMUSG00000026826.9 | 190.1310263 | 1.796138816 | 0.206759246 | 8.687102759 | 3.72E-18 | 5.79E-15 | Nr4a2 |
| ENSMUSG00000033849.3 | 46.40904289 | -3.039983197 | 0.385018653 | -7.895677715 | 2.89E-15 | 4.00E-12 | B3galt2 |
| ENSMUSG00000043088.12 | 83.46258964 | -2.974428715 | 0.379173063 | -7.844514818 | 4.35E-15 | 5.42E-12 | Il17re |
| ENSMUSG00000047898.6 | 69.13291897 | -2.948920733 | 0.378432961 | -7.79245214 | 6.57E-15 | 7.45E-12 | Ccr4 |
| ENSMUSG00000021360.11 | 113.6741344 | -2.192668404 | 0.297356582 | -7.373868736 | 1.66E-13 | 1.72E-10 | Gcnt2 |
| ENSMUSG00000019256.13 | 123.4461495 | -2.005379521 | 0.288471081 | -6.951752357 | 3.61E-12 | 3.46E-09 | Ahr |
| ENSMUSG00000004655.5 | 132.1705665 | 2.366518045 | 0.342802817 | 6.903438143 | 5.08E-12 | 4.52E-09 | Aqp1 |
| ENSMUSG00000046743.6 | 65.11383478 | 2.35075612 | 0.378430359 | 6.21185924 | 5.24E-10 | 4.35E-07 | Fat4 |
| ENSMUSG00000046807.9 | 101.1844203 | -1.609941733 | 0.261385119 | -6.1592708 | 7.31E-10 | 5.69E-07 | Lrrc75b |
| ENSMUSG00000006574.11 | 247.9452257 | 2.190425152 | 0.363299239 | 6.029258852 | 1.65E-09 | 1.14E-06 | Slc4a1 |
| ENSMUSG00000026532.7 | 112.9728465 | 2.140258714 | 0.354566818 | 6.036263424 | 1.58E-09 | 1.14E-06 | Spta1 |
| ENSMUSG00000052374.10 | 138.0875268 | -1.538457223 | 0.262393424 | -5.863169882 | 4.54E-09 | 2.98E-06 | Actn2 |
| ENSMUSG00000076749.1 | 17.67249953 | -2.33643621 | 0.400781508 | -5.829700638 | 5.55E-09 | 3.46E-06 | Tcrg-C1 |
| ENSMUSG00000028332.9 | 51.88009352 | 1.966829344 | 0.345689171 | 5.689589115 | 1.27E-08 | 7.56E-06 | Hemgn |
| ENSMUSG00000006345.6 | 256.358635 | -1.511534837 | 0.266527763 | -5.671209703 | 1.42E-08 | 8.03E-06 | Ggt1 |
| ENSMUSG00000021728.7 | 1584.287655 | -1.153069189 | 0.203607125 | -5.663206464 | 1.49E-08 | 8.05E-06 | Emb |
| ENSMUSG00000028644.12 | 143.9883716 | 2.093617005 | 0.383489237 | 5.459389213 | 4.78E-08 | 2.38E-05 | Ermap |
| ENSMUSG00000049807.12 | 65.27816578 | 2.120526238 | 0.388394332 | 5.45972499 | 4.77E-08 | 2.38E-05 | Arhgap23 |
| ENSMUSG00000021831.8 | 707.0443623 | -1.134950324 | 0.208558335 | -5.441884288 | 5.27E-08 | 2.53E-05 | Ero1l |
| ENSMUSG00000033213.12 | 129.9658646 | -1.833395031 | 0.337455982 | -5.432990169 | 5.54E-08 | 2.56E-05 | AA467197 |
| ENSMUSG00000020490.12 | 40.74918531 | 2.093679917 | 0.387462955 | 5.403561534 | 6.53E-08 | 2.91E-05 | Btnl10 |
| ENSMUSG00000003882.4 | 1542.417909 | -1.684175716 | 0.312522124 | -5.388980766 | 7.09E-08 | 3.05E-05 | Il7r |
| ENSMUSG00000070407.5 | 200.9179448 | -1.596703821 | 0.298623295 | -5.346883007 | 8.95E-08 | 3.72E-05 | Hs3st3b1 |
| ENSMUSG00000020617.9 | 26.59540456 | -2.089526249 | 0.393531329 | -5.309682091 | 1.10E-07 | 4.42E-05 | 1700012B07Rik |
| ENSMUSG00000031543.14 | 194.0862229 | 2.085341819 | 0.393453293 | 5.300100048 | 1.16E-07 | 4.51E-05 | Ank1 |
| ENSMUSG00000021061.11 | 214.5433398 | 2.055741842 | 0.389178971 | 5.282253139 | 1.28E-07 | 4.82E-05 | Sptb |
| ENSMUSG00000089672.4 | 1605.360222 | -1.210074528 | 0.2295673 | -5.271110157 | 1.36E-07 | 4.97E-05 | Gp49a |
| ENSMUSG00000027398.9 | 53.04233206 | 2.05066321 | 0.391902119 | 5.232590264 | 1.67E-07 | 5.95E-05 | Il1b |
| ENSMUSG00000040249.11 | 179.5326873 | 1.855963523 | 0.357745729 | 5.187940408 | 2.13E-07 | 7.36E-05 | Lrp1 |
| ENSMUSG00000038725.7 | 60.70287868 | 2.063076892 | 0.399251976 | 5.167355492 | 2.37E-07 | 8.00E-05 | Pkhd1l1 |
| ENSMUSG00000025473.12 | 1143.15024 | -1.177465905 | 0.228340347 | -5.15662659 | 2.51E-07 | 8.25E-05 | Adam8 |
| ENSMUSG00000006342.10 | 350.0049921 | -1.342056021 | 0.263130608 | -5.100341727 | 3.39E-07 | 0.000105654 | Susd2 |
| ENSMUSG00000056737.10 | 655.2825861 | -0.87435842 | 0.171273279 | -5.105048637 | 3.31E-07 | 0.000105654 | Capg |
| ENSMUSG00000028825.7 | 41.68088816 | 1.923851209 | 0.38380029 | 5.012636148 | 5.37E-07 | 0.000163229 | Rhd |
| ENSMUSG00000076752.1 | 67.42125566 | -1.590977833 | 0.318012287 | -5.002881638 | 5.65E-07 | 0.000167623 | Tcrg-C2 |
| ENSMUSG00000037944.8 | 31.26094199 | -1.801663559 | 0.362371906 | -4.971863247 | 6.63E-07 | 0.000192229 | Ccr7 |
| ENSMUSG00000027562.8 | 201.7182709 | 1.603005105 | 0.322804332 | 4.965872346 | 6.84E-07 | 0.000193754 | Car2 |
| ENSMUSG00000006567.7 | 30.43676161 | 1.95572057 | 0.396858611 | 4.928003362 | 8.31E-07 | 0.000230116 | Atp7b |
| ENSMUSG00000041272.7 | 244.7499702 | 1.438615427 | 0.2922969 | 4.921760801 | 8.58E-07 | 0.000232415 | Tox |
| ENSMUSG00000001763.10 | 54.05309103 | 1.93771349 | 0.395582093 | 4.898385256 | 9.66E-07 | 0.000256268 | Tspan33 |
| ENSMUSG00000017707.9 | 1458.07406 | -0.683493507 | 0.142580609 | -4.793733948 | 1.64E-06 | 0.000425122 | Serinc3 |
| ENSMUSG00000023216.9 | 39.81405927 | 1.915586015 | 0.400175838 | 4.786860753 | 1.69E-06 | 0.000430959 | Epb4.2 |
| ENSMUSG00000030281.12 | 19.41326743 | -1.92546841 | 0.40307118 | -4.776993506 | 1.78E-06 | 0.000434895 | Il17rc |
| ENSMUSG00000051212.7 | 265.0477083 | -1.558515608 | 0.325987111 | -4.780911747 | 1.75E-06 | 0.000434895 | Gpr183 |
| ENSMUSG00000038058.10 | 1069.528245 | 0.837999631 | 0.175737879 | 4.768463333 | 1.86E-06 | 0.000444992 | Nod1 |
| ENSMUSG00000026628.9 | 24.08959098 | 1.799312913 | 0.37964733 | 4.739432551 | 2.14E-06 | 0.000504051 | Atf3 |
| ENSMUSG00000064225.6 | 34.26907372 | 1.821110893 | 0.385136566 | 4.728480889 | 2.26E-06 | 0.000522159 | Paqr9 |
| ENSMUSG00000020027.14 | 143.5154679 | -1.42293642 | 0.301174841 | -4.724619143 | 2.31E-06 | 0.000522503 | Socs2 |
| ENSMUSG00000030000.8 | 60.85907174 | 1.875189384 | 0.402347332 | 4.660623381 | 3.15E-06 | 0.00070172 | Add2 |
| ENSMUSG00000020865.12 | 24.86836142 | -1.732128552 | 0.375137846 | -4.617312203 | 3.89E-06 | 0.000850118 | Abcc3 |
| ENSMUSG00000022099.12 | 28.95592242 | 1.858861847 | 0.403099164 | 4.611425702 | 4.00E-06 | 0.000859476 | Dmtn |
| ENSMUSG00000028435.8 | 47.56723154 | -1.692700769 | 0.367445704 | -4.606669097 | 4.09E-06 | 0.000864459 | Aqp3 |
| ENSMUSG00000000982.5 | 133.1239663 | 1.234199291 | 0.269595948 | 4.577959356 | 4.70E-06 | 0.000956113 | Ccl3 |

| | | | | | | | |
|---|---|---|---|---|---|---|---|
| ENSMUSG00000024621.11 | 109.7478397 | 1.634594226 | 0.357265799 | 4.575288844 | 4.76E-06 | 0.000956113 | Csf1r |
| ENSMUSG00000073418.4 | 27.64091047 | 1.827078135 | 0.398950932 | 4.579706393 | 4.66E-06 | 0.000956113 | C4b |
| ENSMUSG00000042476.8 | 63.56729265 | 1.795071026 | 0.395009757 | 4.544371352 | 5.51E-06 | 0.001082672 | Abcb4 |
| ENSMUSG00000043252.8 | 229.8761089 | -1.24985617 | 0.275344731 | -4.539241286 | 5.65E-06 | 0.001082672 | Tmem64 |
| ENSMUSG00000097971.3 | 343059.1617 | 1.758631244 | 0.387274781 | 4.541042506 | 5.60E-06 | 0.001082672 | Gm26917 |
| ENSMUSG00000000732.8 | 199.9115381 | -1.045958148 | 0.230784351 | -4.53218836 | 5.84E-06 | 0.001089995 | Icosl |
| ENSMUSG00000038264.7 | 294.6427867 | 1.42666952 | 0.314839202 | 4.531422733 | 5.86E-06 | 0.001089995 | Sema7a |
| ENSMUSG00000022270.11 | 376.0759145 | -0.962630474 | 0.21297449 | -4.51993323 | 6.19E-06 | 0.001125051 | Fam134b |
| ENSMUSG00000040899.9 | 49.69159271 | -1.802866015 | 0.398995953 | -4.518507026 | 6.23E-06 | 0.001125051 | Ccr6 |
| ENSMUSG00000048521.7 | 2330.743337 | -1.639896711 | 0.364070808 | -4.504334523 | 6.66E-06 | 0.001162551 | Cxcr6 |
| ENSMUSG00000062593.11 | 1119.724655 | -0.833552492 | 0.184958189 | -4.506707678 | 6.58E-06 | 0.001162551 | Lilrb4 |
| ENSMUSG00000078942.6 | 127.0147342 | 1.20969859 | 0.26867118 | 4.502524573 | 6.72E-06 | 0.001162551 | Naip6 |
| ENSMUSG00000096385.3 | 55.33238278 | 1.806545968 | 0.40312576 | 4.481345895 | 7.42E-06 | 0.001266543 | Gm11168 |
| ENSMUSG00000029516.15 | 1146.694529 | 1.274669391 | 0.285415244 | 4.4660172 | 7.97E-06 | 0.001273495 | Cit |
| ENSMUSG00000040229.7 | 32.43576564 | -1.667460678 | 0.37324977 | -4.467412471 | 7.92E-06 | 0.001273495 | Gpr34 |
| ENSMUSG00000040703.7 | 33.70239953 | -1.585716346 | 0.354459779 | -4.473614326 | 7.69E-06 | 0.001273495 | Cyp2s1 |
| ENSMUSG00000049608.8 | 51.00569829 | -1.796764184 | 0.402215624 | -4.467166558 | 7.93E-06 | 0.001273495 | Gpr55 |
| ENSMUSG00000051910.9 | 33.72624787 | 1.776011389 | 0.397666322 | 4.466084481 | 7.97E-06 | 0.001273495 | Sox6 |
| ENSMUSG00000045502.5 | 14.36948327 | 1.782670482 | 0.400420795 | 4.451992762 | 8.51E-06 | 0.001342386 | Hcar2 |
| ENSMUSG00000023274.10 | 5292.051337 | -0.624046914 | 0.140692097 | -4.435550602 | 9.18E-06 | 0.001430939 | Cd4 |
| ENSMUSG00000019838.10 | 49.56715068 | 1.738982109 | 0.392809226 | 4.427039884 | 9.55E-06 | 0.00144938 | Slc16a10 |
| ENSMUSG00000047415.7 | 834.3197808 | -0.775149521 | 0.175180951 | -4.42485051 | 9.65E-06 | 0.00144938 | Gpr68 |
| ENSMUSG00000048440.11 | 34.93052799 | -1.52528298 | 0.34451788 | -4.427297008 | 9.54E-06 | 0.00144938 | Cyp4f16 |
| ENSMUSG00000003032.8 | 45.56631923 | 1.594970476 | 0.360755959 | 4.421189547 | 9.82E-06 | 0.00145661 | Klf4 |
| ENSMUSG00000032021.9 | 113.8027481 | 1.246049404 | 0.282242495 | 4.414818557 | 1.01E-05 | 0.00146959 | Crtam |
| ENSMUSG00000052821.3 | 18.89414431 | -1.778563217 | 0.402920044 | -4.414184014 | 1.01E-05 | 0.00146959 | Cysltr1 |
| ENSMUSG00000026019.11 | 187.4412389 | -0.878245336 | 0.199138306 | -4.410228016 | 1.03E-05 | 0.001479493 | Wdr12 |
| ENSMUSG00000028717.8 | 40.8279565 | 1.714942829 | 0.390118133 | 4.395957743 | 1.10E-05 | 0.001544615 | Tal1 |
| ENSMUSG00000037936.11 | 97.74211941 | 1.393318573 | 0.316897753 | 4.396744879 | 1.10E-05 | 0.001544615 | Scarb1 |
| ENSMUSG00000020841.5 | 371.4879459 | -1.409573888 | 0.321085923 | -4.390020821 | 1.13E-05 | 0.001552506 | Cpd |
| ENSMUSG00000024052.13 | 890.5238818 | 0.664055193 | 0.151183398 | 4.392381714 | 1.12E-05 | 0.001552506 | Lpin2 |
| ENSMUSG00000024793.10 | 684.5084782 | -0.929526405 | 0.212060123 | -4.383315418 | 1.17E-05 | 0.0015696 | Tnfrsf25 |
| ENSMUSG00000031877.8 | 30.78975324 | 1.658293868 | 0.378354799 | 4.382906925 | 1.17E-05 | 0.0015696 | Ces2g |
| ENSMUSG00000053702.12 | 484.1962494 | -1.433671721 | 0.327739713 | -4.374421722 | 1.22E-05 | 0.001614546 | Nebl |
| ENSMUSG00000024646.9 | 381.3498474 | -0.86212688 | 0.197793741 | -4.35871669 | 1.31E-05 | 0.001716592 | Cyb5a |
| ENSMUSG00000025427.10 | 32.10845638 | 1.644403018 | 0.377689991 | 4.353843247 | 1.34E-05 | 0.001736938 | Rnf165 |
| ENSMUSG00000041237.8 | 34.11228071 | 1.692228338 | 0.388878845 | 4.351556686 | 1.35E-05 | 0.00173706 | Pklr |
| ENSMUSG00000032131.11 | 32.98863984 | 1.72748463 | 0.397501654 | 4.345855202 | 1.39E-05 | 0.001764612 | Abcg4 |
| ENSMUSG00000031170.10 | 14.3729425 | 1.727754423 | 0.39842304 | 4.336482207 | 1.45E-05 | 0.001822926 | Slc38a5 |
| ENSMUSG00000003469.5 | 8.839702257 | 1.726412054 | 0.400683287 | 4.308669994 | 1.64E-05 | 0.002047242 | Phyhip |
| ENSMUSG00000043807.6 | 78.38713908 | -1.534022306 | 0.356900727 | -4.298176466 | 1.72E-05 | 0.002117905 | Ly6g5b |
| ENSMUSG00000044345.9 | 238.8663933 | -0.848907547 | 0.197568851 | -4.296768157 | 1.73E-05 | 0.002117905 | Marveld1 |
| ENSMUSG00000041515.5 | 441.3140235 | 1.016822626 | 0.237057562 | 4.289349049 | 1.79E-05 | 0.002168638 | Irf8 |
| ENSMUSG00000054889.5 | 40.26076736 | 1.542298611 | 0.360394198 | 4.279476799 | 1.87E-05 | 0.002243377 | Dsp |
| ENSMUSG00000086968.4 | 40.24433186 | -1.627758623 | 0.380536443 | -4.277536759 | 1.89E-05 | 0.002243377 | 4933431E20Rik |
| ENSMUSG00000058427.7 | 10.02984777 | 1.722100546 | 0.402805145 | 4.275269484 | 1.91E-05 | 0.002244949 | Cxcl2 |
| ENSMUSG00000028078.10 | 241.1844124 | 1.046525887 | 0.245040671 | 4.27082525 | 1.95E-05 | 0.002268758 | Dclk2 |
| ENSMUSG00000026072.8 | 196.9934097 | -1.677368423 | 0.393861032 | -4.258782384 | 2.06E-05 | 0.002329181 | Il1r1 |
| ENSMUSG00000028456.13 | 53.27328814 | 1.711060467 | 0.401711305 | 4.259428212 | 2.05E-05 | 0.002329181 | Unc13b |
| ENSMUSG00000028977.12 | 72.74470896 | 1.434878619 | 0.336622853 | 4.262570427 | 2.02E-05 | 0.002329181 | Casz1 |
| ENSMUSG00000034156.12 | 771.3319209 | 1.202023599 | 0.282599921 | 4.253446335 | 2.11E-05 | 0.002363919 | Bzrap1 |
| ENSMUSG00000028132.11 | 24.73720151 | 1.646997389 | 0.387772172 | 4.247332606 | 2.16E-05 | 0.002407645 | Tmem56 |
| ENSMUSG00000027995.10 | 31.4350471 | 1.565172559 | 0.369226101 | 4.239062605 | 2.24E-05 | 0.002475958 | Tlr2 |
| ENSMUSG00000000555.6 | 25.66631204 | 1.642760546 | 0.388662718 | 4.22669958 | 2.37E-05 | 0.002592982 | Itga5 |
| ENSMUSG00000029866.9 | 40.7005409 | 1.680363797 | 0.39886625 | 4.212850291 | 2.52E-05 | 0.002733284 | Kel |
| ENSMUSG00000042035.7 | 22.14728972 | 1.691292913 | 0.403039063 | 4.196349852 | 2.71E-05 | 0.002889868 | Igsf3 |
| ENSMUSG00000047293.6 | 14.85865206 | -1.6893979 | 0.402474218 | -4.197530733 | 2.70E-05 | 0.002889868 | Gpr15 |
| ENSMUSG00000005640.7 | 66.92441179 | 1.529422013 | 0.366826442 | 4.169334158 | 3.05E-05 | 0.003173286 | Insrr |
| ENSMUSG00000022126.6 | 27.32545948 | 1.652326008 | 0.396102024 | 4.171465701 | 3.03E-05 | 0.003173286 | Irg1 |
| ENSMUSG00000024533.11 | 49.17874549 | 1.574646233 | 0.377630683 | 4.169804793 | 3.05E-05 | 0.003173286 | Spire1 |
| ENSMUSG00000021253.6 | 45.21057511 | -1.408501463 | 0.338439675 | -4.161750436 | 3.16E-05 | 0.003226774 | Tgfb3 |
| ENSMUSG00000026815.10 | 20.45985215 | 1.632805339 | 0.392218817 | 4.162995929 | 3.14E-05 | 0.003226774 | Gfi1b |
| ENSMUSG00000076745.1 | 8.993142243 | -1.661432512 | 0.399471302 | -4.159078517 | 3.20E-05 | 0.003238205 | Tcrg-V4 |

| | | | | | | | |
|---|---|---|---|---|---|---|---|
| ENSMUSG00000042262.4 | 69.70866769 | -1.306882386 | 0.315750435 | -4.138972563 | 3.49E-05 | 0.003478878 | Ccr8 |
| ENSMUSG00000060012.7 | 1115.211849 | 1.191969707 | 0.287919037 | 4.139947531 | 3.47E-05 | 0.003478878 | Kif13b |
| ENSMUSG00000030302.12 | 20.56983355 | 1.6447682 | 0.397762397 | 4.135052009 | 3.55E-05 | 0.00351072 | Atp2b2 |
| ENSMUSG00000036606.12 | 42.68085899 | 1.598532916 | 0.387410348 | 4.126200871 | 3.69E-05 | 0.003619808 | Plxnb2 |
| ENSMUSG00000030157.5 | 1176.20471 | -0.733515492 | 0.178008338 | -4.120680516 | 3.78E-05 | 0.003678684 | Clec2d |
| ENSMUSG00000039021.11 | 53.05843947 | 1.228370235 | 0.299873091 | 4.096300312 | 4.20E-05 | 0.004056491 | Ttc16 |
| ENSMUSG00000028937.10 | 413.2234645 | -0.68344522 | 0.167494079 | -4.08041422 | 4.50E-05 | 0.004310542 | Acot7 |
| ENSMUSG00000046908.5 | 104.0295693 | -1.262043052 | 0.309997599 | -4.07113815 | 4.68E-05 | 0.004451622 | Ltb4r1 |
| ENSMUSG00000033910.9 | 36.64876438 | 1.601915876 | 0.394225513 | 4.063450541 | 4.84E-05 | 0.004566021 | Gucy1a3 |
| ENSMUSG00000021236.12 | 495.0966791 | -0.776689875 | 0.191464825 | -4.056566919 | 4.98E-05 | 0.00466728 | Entpd5 |
| ENSMUSG00000054672.8 | 46.98435953 | -1.528069492 | 0.377376919 | -4.049186406 | 5.14E-05 | 0.004780978 | 5830411N06Rik |
| ENSMUSG00000023926.7 | 36.28402933 | 1.565834729 | 0.38708015 | 4.045246781 | 5.23E-05 | 0.004817135 | Rhag |
| ENSMUSG00000032548.10 | 49.78893139 | 1.618498915 | 0.400226993 | 4.043952412 | 5.26E-05 | 0.004817135 | Slco2a1 |
| ENSMUSG00000029716.9 | 27.4982121 | 1.627790659 | 0.40275643 | 4.041625501 | 5.31E-05 | 0.004829683 | Tfr2 |
| ENSMUSG00000004085.10 | 37.38810576 | 1.574625034 | 0.389987396 | 4.037630574 | 5.40E-05 | 0.004877048 | Zak |
| ENSMUSG00000037463.10 | 38.65071108 | -1.302295022 | 0.32324633 | -4.028800649 | 5.61E-05 | 0.005027442 | Fbxo27 |
| ENSMUSG00000035064.12 | 38.19085901 | 1.553118939 | 0.385933031 | 4.024322394 | 5.71E-05 | 0.005087465 | Eef2k |
| ENSMUSG00000021109.9 | 2527.467319 | -0.679702384 | 0.169078349 | -4.0200439 | 5.82E-05 | 0.005144005 | Hif1a |
| ENSMUSG00000021057.11 | 35.25564431 | 1.52970922 | 0.381273012 | 4.012109886 | 6.02E-05 | 0.005245627 | Akap5 |
| ENSMUSG00000045991.14 | 21.74155842 | -1.472995165 | 0.367131142 | -4.012177118 | 6.02E-05 | 0.005245627 | Onecut2 |
| ENSMUSG00000026475.7 | 208.6590831 | 1.343024606 | 0.334964842 | 4.00944946 | 6.09E-05 | 0.005268235 | Rgs16 |
| ENSMUSG00000037706.12 | 175.77261 | 1.021308673 | 0.254839831 | 4.007649317 | 6.13E-05 | 0.005271929 | Cd81 |
| ENSMUSG00000026480.8 | 201.2013756 | 1.368743887 | 0.34182057 | 4.004275951 | 6.22E-05 | 0.005304633 | Ncf2 |
| ENSMUSG00000051839.6 | 36.16580412 | 1.520904839 | 0.379946041 | 4.002949563 | 6.26E-05 | 0.005304633 | Gypa |
| ENSMUSG00000041329.9 | 24.18739252 | 1.609928267 | 0.402638235 | 3.99844855 | 6.38E-05 | 0.005369976 | Atp1b2 |
| ENSMUSG00000081453.1 | 38.60707388 | -1.294569987 | 0.324558064 | -3.98871614 | 6.64E-05 | 0.005557538 | Gm6767 |
| ENSMUSG00000022657.9 | 1018.053093 | -1.125178012 | 0.282216669 | -3.986929673 | 6.69E-05 | 0.005562203 | Cd96 |
| ENSMUSG00000029101.10 | 184.1294574 | 1.088055175 | 0.273923313 | 3.972116005 | 7.12E-05 | 0.005880592 | Rgs12 |
| ENSMUSG00000012123.11 | 316.5501839 | 0.770727461 | 0.194313514 | 3.96641203 | 7.30E-05 | 0.00598342 | Aim1l |
| ENSMUSG00000020009.8 | 3157.66473 | -1.227616498 | 0.309843617 | -3.962051924 | 7.43E-05 | 0.006053947 | Ifngr1 |
| ENSMUSG00000032691.10 | 27.25264185 | 1.591943285 | 0.402670707 | 3.953461866 | 7.70E-05 | 0.00623481 | Nlrp3 |
| ENSMUSG00000024140.9 | 294.92997 | -0.74424647 | 0.188454416 | -3.949212159 | 7.84E-05 | 0.006296413 | Epas1 |
| ENSMUSG00000025492.6 | 295.7037713 | -1.191088574 | 0.301692557 | -3.948021079 | 7.88E-05 | 0.006296413 | Ifitm3 |
| ENSMUSG00000001020.7 | 3388.309343 | -1.375423372 | 0.34883188 | -3.942940569 | 8.05E-05 | 0.00630565 | S100a4 |
| ENSMUSG00000033446.7 | 242.3777644 | -1.006737968 | 0.255214878 | -3.944668016 | 7.99E-05 | 0.00630565 | Lpar6 |
| ENSMUSG00000045362.7 | 159.4361175 | -1.071243395 | 0.271697769 | -3.942775816 | 8.05E-05 | 0.00630565 | Tnfrsf26 |
| ENSMUSG00000047139.8 | 191.3459923 | 1.430402948 | 0.362898809 | 3.941602757 | 8.09E-05 | 0.00630565 | Cd24a |
| ENSMUSG00000020614.9 | 34.32723225 | -1.45866188 | 0.37086238 | -3.933162165 | 8.38E-05 | 0.006490753 | Fam20a |
| ENSMUSG00000059674.6 | 103.0068344 | 1.335669746 | 0.339742286 | 3.931420372 | 8.44E-05 | 0.00649761 | Cdh24 |
| ENSMUSG00000061533.11 | 359.0240672 | 1.149175196 | 0.292536473 | 3.928314252 | 8.55E-05 | 0.006541707 | Cep128 |
| ENSMUSG00000002997.11 | 56.96110274 | 1.067739604 | 0.272517647 | 3.918056747 | 8.93E-05 | 0.006784737 | Prkar2b |
| ENSMUSG00000068566.8 | 303.3088137 | -0.925920914 | 0.236940186 | -3.907825562 | 9.31E-05 | 0.007035586 | Myadm |
| ENSMUSG00000053310.7 | 108.0654689 | 1.45026793 | 0.371301456 | 3.90590424 | 9.39E-05 | 0.007049009 | Nrgn |
| ENSMUSG00000035891.12 | 128.046794 | -1.022468089 | 0.261987681 | -3.902733455 | 9.51E-05 | 0.00709926 | Cerk |
| ENSMUSG00000024053.10 | 104.3589743 | 1.54351448 | 0.395740848 | 3.900316302 | 9.61E-05 | 0.007127835 | Emilin2 |
| ENSMUSG00000051439.6 | 10.62131762 | 1.569454069 | 0.403036775 | 3.894071625 | 9.86E-05 | 0.007270674 | Cd14 |
| ENSMUSG00000030283.5 | 131.8879303 | -1.341914871 | 0.344803746 | -3.891822194 | 9.95E-05 | 0.007295262 | St8sia1 |
| ENSMUSG00000032011.4 | 3813.19599 | -0.679912637 | 0.175959004 | -3.864040042 | 0.000111527 | 0.008129726 | Thy1 |
| ENSMUSG00000020644.8 | 2979.674813 | -0.989748688 | 0.256559421 | -3.857775652 | 0.000114424 | 0.008244452 | Id2 |
| ENSMUSG00000045071.9 | 171.4140312 | 1.22456195 | 0.317381743 | 3.858325111 | 0.000114167 | 0.008244452 | E130308A19Rik |
| ENSMUSG00000034792.8 | 708.1374716 | -0.707449771 | 0.183969913 | -3.845464518 | 0.000120324 | 0.008619782 | Gna15 |
| ENSMUSG00000021281.11 | 168.7857805 | 1.527041821 | 0.397788142 | 3.838831933 | 0.000123621 | 0.008767596 | Tnfaip2 |
| ENSMUSG00000022724.11 | 565.2254138 | -0.704158246 | 0.183446786 | -3.838487782 | 0.000123794 | 0.008767596 | Mina |
| ENSMUSG00000079186.2 | 16.24709748 | -1.522921113 | 0.398085992 | -3.825608394 | 0.00013045 | 0.009186742 | Gzmc |
| ENSMUSG00000032089.12 | 1065.866149 | 0.863620594 | 0.22623346 | 3.81738666 | 0.000134873 | 0.009425317 | Il10ra |
| ENSMUSG00000037020.12 | 189.5559773 | 0.964791603 | 0.252793831 | 3.816515614 | 0.00013535 | 0.009425317 | Wdr62 |
| ENSMUSG00000026657.12 | 29.17770985 | 1.493823666 | 0.391833125 | 3.812397599 | 0.000137625 | 0.009530549 | Frmd4a |
| ENSMUSG00000020863.11 | 930.2235777 | -0.652805117 | 0.171564159 | -3.805020352 | 0.000141793 | 0.00971123 | Luc7l3 |
| ENSMUSG00000042745.9 | 10.00990393 | 1.502192603 | 0.394755393 | 3.805375757 | 0.000141589 | 0.00971123 | Id1 |
| ENSMUSG00000020689.4 | 439.0309902 | -0.606157698 | 0.159598135 | -3.798024948 | 0.000145854 | 0.009934785 | Itgb3 |
| ENSMUSG00000020108.3 | 994.0107736 | -0.555679116 | 0.146431333 | -3.794810206 | 0.000147756 | 0.010009696 | Ddit4 |
| ENSMUSG00000019943.9 | 1998.992488 | -0.779420724 | 0.205526126 | -3.792319447 | 0.000149247 | 0.010056004 | Atp2b1 |
| ENSMUSG00000097240.1 | 35.88697106 | -1.404256972 | 0.370844325 | -3.786648135 | 0.000152693 | 0.010232899 | Gm26614 |

| | | | | | | | |
|---|---|---|---|---|---|---|---|
| ENSMUSG00000051339.8 | 174.9076785 | 1.485667686 | 0.392754641 | 3.782686522 | 0.000155145 | 0.010341601 | 2900026A02Rik |
| ENSMUSG00000061175.7 | 94.9128033 | 1.216550358 | 0.321976451 | 3.778383034 | 0.00015785 | 0.010465956 | Fnip2 |
| ENSMUSG00000058056.11 | 11.02981127 | 1.518537137 | 0.403040916 | 3.767699697 | 0.000164759 | 0.010866231 | Palld |
| ENSMUSG00000033767.10 | 904.1756064 | 0.591444593 | 0.157398873 | 3.757616446 | 0.00017154 | 0.011253897 | D930015E06Rik |
| ENSMUSG00000000594.7 | 935.1344719 | -0.856961168 | 0.228253326 | -3.754430137 | 0.000173736 | 0.011279286 | Gm2a |
| ENSMUSG00000034265.7 | 12.33404343 | 1.479752634 | 0.394104314 | 3.754723262 | 0.000173533 | 0.011279286 | Zdhhc14 |
| ENSMUSG00000046591.9 | 456.7255373 | 0.849911592 | 0.226748844 | 3.748251045 | 0.000178072 | 0.011500866 | Ticrr |
| ENSMUSG00000003380.10 | 247.9054193 | -0.942435278 | 0.251620889 | -3.745457228 | 0.000180066 | 0.011569674 | Rabac1 |
| ENSMUSG00000027368.6 | 1462.804714 | 0.597058882 | 0.159633628 | 3.740182373 | 0.000183887 | 0.011635273 | Dusp2 |
| ENSMUSG00000040466.11 | 59.80226882 | 1.307489326 | 0.349417688 | 3.741909381 | 0.000182627 | 0.011635273 | Blvrb |
| ENSMUSG00000041912.8 | 24.24552315 | -1.339990282 | 0.358231423 | -3.740571587 | 0.000183602 | 0.011635273 | Tdrkh |
| ENSMUSG00000025702.11 | 41.37577611 | 1.417921011 | 0.380092151 | 3.730466434 | 0.000191126 | 0.012032226 | March8 |
| ENSMUSG00000054640.10 | 20.67433342 | 1.499788819 | 0.40276346 | 3.723745994 | 0.000196288 | 0.012295151 | Slc8a1 |
| ENSMUSG00000040102.9 | 279.886892 | -0.686039794 | 0.184320243 | -3.721999195 | 0.000197652 | 0.012318639 | Klhl42 |
| ENSMUSG00000034107.6 | 44.87557467 | 1.476708306 | 0.397822461 | 3.711978207 | 0.000205646 | 0.012753103 | Ano7 |
| ENSMUSG00000015846.10 | 285.4451721 | -0.67627202 | 0.182346093 | -3.708727781 | 0.000208303 | 0.012853959 | Rxra |
| ENSMUSG00000022377.12 | 1273.988519 | 1.141051716 | 0.307845405 | 3.706573811 | 0.000210082 | 0.012899864 | Asap1 |
| ENSMUSG00000031015.7 | 230.5229612 | -1.015646581 | 0.274216083 | -3.703818432 | 0.000212378 | 0.012976938 | Swap70 |
| ENSMUSG00000015533.8 | 289.1984166 | -0.956962206 | 0.258778642 | -3.697995323 | 0.000217309 | 0.013113314 | Itga2 |
| ENSMUSG00000023927.11 | 2207.737853 | -0.816006624 | 0.22059883 | -3.699052373 | 0.000216406 | 0.013113314 | Satb1 |
| ENSMUSG00000036718.13 | 24.76689238 | 1.489451339 | 0.402830789 | 3.697461511 | 0.000217766 | 0.013113314 | Micall2 |
| ENSMUSG00000029408.9 | 232.4752201 | 0.875002167 | 0.236747999 | 3.695922119 | 0.00021909 | 0.013129608 | Abcb9 |
| ENSMUSG00000073758.6 | 16.95343238 | 1.445506438 | 0.391833752 | 3.689080968 | 0.000225066 | 0.01342317 | Sh3d21 |
| ENSMUSG00000029641.7 | 32.68396822 | -1.389518843 | 0.376794742 | -3.687734163 | 0.00022626 | 0.013430138 | Rasl11a |
| ENSMUSG00000002365.9 | 94.45329751 | 1.057700729 | 0.28749685 | 3.67899937 | 0.000234151 | 0.013767407 | Snx9 |
| ENSMUSG00000049410.8 | 410.0552477 | 0.811792911 | 0.220586503 | 3.680156771 | 0.000233091 | 0.013767407 | Zfp683 |
| ENSMUSG00000091650.1 | 11.4849621 | 1.449755692 | 0.394614059 | 3.67385718 | 0.000238916 | 0.013981652 | Apol11a |
| ENSMUSG00000026070.11 | 3976.475248 | -0.618389755 | 0.168596003 | -3.6678791 | 0.000244571 | 0.014179417 | Il18r1 |
| ENSMUSG00000032690.12 | 63.73725551 | 1.475929666 | 0.402284737 | 3.668868172 | 0.000243627 | 0.014179417 | Oas2 |
| ENSMUSG00000028460.6 | 322.3438651 | -1.081779981 | 0.295888603 | -3.65603802 | 0.000256143 | 0.014781608 | Sit1 |
| ENSMUSG00000020900.11 | 36.14806575 | 1.459033498 | 0.400145837 | 3.646254345 | 0.000266091 | 0.015284882 | Myh10 |
| ENSMUSG00000024247.10 | 56.04417311 | 1.40470215 | 0.385786295 | 3.641140623 | 0.000271433 | 0.01552023 | Pkdcc |
| ENSMUSG00000055013.10 | 45.77835299 | 1.465313101 | 0.402691648 | 3.63879685 | 0.000273915 | 0.015590628 | Agap1 |
| ENSMUSG00000075590.2 | 63.8551014 | 1.364085196 | 0.375439857 | 3.633298832 | 0.000279821 | 0.015854382 | Nrbp2 |
| ENSMUSG00000060419.8 | 83.73878669 | -1.220548898 | 0.336244918 | -3.629940065 | 0.000283487 | 0.015989439 | Rps16-ps2 |
| ENSMUSG00000029603.11 | 1028.747051 | -0.590752795 | 0.162837561 | -3.627865646 | 0.000285774 | 0.016045818 | Dtx1 |
| ENSMUSG00000051278.8 | 355.6176845 | 0.799423421 | 0.221277255 | 3.612768159 | 0.000302946 | 0.016933705 | Zgrf1 |
| ENSMUSG00000032815.11 | 349.9024955 | 1.157936731 | 0.320880063 | 3.60862785 | 0.000307821 | 0.017053273 | Fanca |
| ENSMUSG00000071226.7 | 13.38592625 | 1.454808962 | 0.403063843 | 3.609376003 | 0.000306934 | 0.017053273 | Cecr2 |
| ENSMUSG00000006235.5 | 18.30236589 | 1.45014891 | 0.40252117 | 3.602664949 | 0.000314971 | 0.017154624 | Epor |
| ENSMUSG00000036882.6 | 118.2740052 | 1.260026198 | 0.349662099 | 3.603553833 | 0.000313896 | 0.017154624 | Arhgap33 |
| ENSMUSG00000054191.7 | 33.22730554 | 1.434296312 | 0.398137675 | 3.602513409 | 0.000315155 | 0.017154624 | Klf1 |
| ENSMUSG00000054404.8 | 142.3256811 | -1.178544419 | 0.326944516 | -3.604723006 | 0.000312486 | 0.017154624 | Slfn5 |
| ENSMUSG00000028874.10 | 111.2086746 | 1.306544458 | 0.363289045 | 3.596432303 | 0.000322611 | 0.01748414 | Fgr |
| ENSMUSG00000097636.3 | 218.0063066 | 0.732626271 | 0.20390789 | 3.592927525 | 0.000326984 | 0.017644372 | 5830416P10Rik |
| ENSMUSG00000020889.11 | 14.14674329 | -1.445617801 | 0.402907572 | -3.587963844 | 0.00033327 | 0.017753063 | Nr1d1 |
| ENSMUSG00000076754.1 | 34.22643511 | -1.27264225 | 0.354476704 | -3.590199958 | 0.000330424 | 0.017753063 | Trgv2 |
| ENSMUSG00000102151.1 | 262.8685752 | 1.099384299 | 0.306370132 | 3.588418663 | 0.00033269 | 0.017753063 | Gm37472 |
| ENSMUSG00000022438.6 | 18.07069451 | 1.435641893 | 0.401502175 | 3.575676502 | 0.000349323 | 0.018450492 | Parvb |
| ENSMUSG00000034220.7 | 578.8700995 | -0.742078382 | 0.207483199 | -3.576570951 | 0.000348131 | 0.018450492 | Gpc1 |
| ENSMUSG00000049866.8 | 1944.985621 | -0.963022831 | 0.26944192 | -3.574138834 | 0.000351382 | 0.018480938 | Arl4c |
| ENSMUSG00000028525.12 | 663.0630926 | -0.609282031 | 0.170565415 | -3.572131137 | 0.000354088 | 0.018544988 | Pde4b |
| ENSMUSG00000015950.9 | 465.7587881 | 1.286970761 | 0.360752896 | 3.56745788 | 0.000360461 | 0.018798579 | Ncf1 |
| ENSMUSG00000036086.12 | 108.4545297 | 0.853026173 | 0.239185428 | 3.566380191 | 0.000361946 | 0.018798579 | Zranb3 |
| ENSMUSG00000030134.9 | 173.2262402 | 1.189430333 | 0.3340686 | 3.560437384 | 0.000370238 | 0.019109803 | Rasgef1a |
| ENSMUSG00000031444.12 | 20.16585212 | 1.431216496 | 0.402039083 | 3.559893939 | 0.000371005 | 0.019109803 | F10 |
| ENSMUSG00000035493.9 | 156.8205598 | 1.428011334 | 0.401817652 | 3.553879047 | 0.000379594 | 0.019471753 | Tgfbi |
| ENSMUSG00000019843.10 | 3905.858998 | 0.504053712 | 0.142037497 | 3.548736926 | 0.000387084 | 0.01950077 | Fyn |
| ENSMUSG00000022021.9 | 656.026487 | 0.628243228 | 0.17699308 | 3.54953555 | 0.000385911 | 0.01950077 | Diap3 |
| ENSMUSG00000026880.10 | 87.57129904 | 1.044901129 | 0.294198837 | 3.551683411 | 0.000382775 | 0.01950077 | Stom |
| ENSMUSG00000027075.12 | 40.34798693 | 1.417871194 | 0.399561365 | 3.548569305 | 0.00038733 | 0.01950077 | Slc43a1 |
| ENSMUSG00000050621.6 | 371.3689133 | -1.259935691 | 0.35509885 | -3.548126645 | 0.000387982 | 0.01950077 | Rps27rt |
| ENSMUSG00000001741.7 | 1690.348091 | -0.458884275 | 0.129822424 | -3.534707349 | 0.000408227 | 0.020435942 | Il16 |

| | | | | | | | |
|---|---|---|---|---|---|---|---|
| ENSMUSG00000033209.13 | 58.84788577 | 1.165340118 | 0.329912863 | 3.532266385 | 0.000412014 | 0.020543022 | Ttc28 |
| ENSMUSG00000040675.13 | 602.9075684 | 0.742177956 | 0.210480547 | 3.526111866 | 0.000421709 | 0.020886602 | Mthfd1l |
| ENSMUSG00000061414.4 | 261.3678329 | 0.993868818 | 0.281887161 | 3.525768299 | 0.000422256 | 0.020886602 | Cracr2a |
| ENSMUSG00000025931.11 | 40.59125279 | -1.228468594 | 0.348897341 | -3.52100303 | 0.000429918 | 0.021129567 | Paqr8 |
| ENSMUSG00000037992.12 | 292.6215242 | -0.722802436 | 0.20530613 | -3.520608155 | 0.000430558 | 0.021129567 | Rara |
| ENSMUSG00000102189.1 | 11.26914611 | 1.408281898 | 0.400644071 | 3.515044902 | 0.00043968 | 0.021492575 | Gm37194 |
| ENSMUSG00000061878.11 | 19.73653 | 1.405773184 | 0.400253846 | 3.512204065 | 0.000444407 | 0.021638784 | Sphk1 |
| ENSMUSG00000043940.10 | 71.06550584 | 1.415001585 | 0.403122958 | 3.510099229 | 0.000447939 | 0.021725937 | Wdfy3 |
| ENSMUSG00000064267.9 | 554.4612761 | -0.548310648 | 0.156575522 | -3.501892512 | 0.000461966 | 0.022319401 | Hvcn1 |
| ENSMUSG00000028668.5 | 1243.411273 | -0.546826205 | 0.156344806 | -3.497565525 | 0.000469525 | 0.022524881 | Tceb3 |
| ENSMUSG00000038384.12 | 634.0267907 | 1.273643678 | 0.364189568 | 3.497199784 | 0.00047017 | 0.022524881 | Setd1b |
| ENSMUSG00000054555.7 | 61.80049819 | -1.336751451 | 0.382325875 | -3.496366679 | 0.00047164 | 0.022524881 | Adam12 |
| ENSMUSG00000029406.11 | 652.5416757 | 1.132644366 | 0.32428998 | 3.492689982 | 0.000478181 | 0.022671277 | Pitpnm2 |
| ENSMUSG00000039994.11 | 518.3064514 | 0.795402583 | 0.227739408 | 3.492599675 | 0.000478343 | 0.022671277 | Timeless |
| ENSMUSG00000032198.8 | 46.5265291 | 1.367620707 | 0.391850552 | 3.490158938 | 0.000482733 | 0.022792694 | Dock6 |
| ENSMUSG00000001506.10 | 23.26801122 | 1.39120869 | 0.399037777 | 3.486408484 | 0.000489553 | 0.022906112 | Col1a1 |
| ENSMUSG00000003545.2 | 655.2756421 | 1.073704891 | 0.308021586 | 3.485810544 | 0.000490648 | 0.022906112 | Fosb |
| ENSMUSG00000042265.9 | 27.29484864 | 1.388743335 | 0.398267767 | 3.486958903 | 0.000488546 | 0.022906112 | Trem1 |
| ENSMUSG00000059326.6 | 107.9308675 | 1.394285258 | 0.400317608 | 3.482947618 | 0.000495925 | 0.023066082 | Csf2ra |
| ENSMUSG00000038267.10 | 20.45402107 | 1.389443135 | 0.399502792 | 3.477930981 | 0.0005053 | 0.023414739 | Slc22a23 |
| ENSMUSG00000019302.12 | 40.92388564 | 1.394065566 | 0.40105888 | 3.475962352 | 0.000509024 | 0.023499929 | Atp6v0a1 |
| ENSMUSG00000020601.7 | 596.1014099 | -0.871433779 | 0.250935923 | -3.472734273 | 0.000515185 | 0.023633114 | Trib2 |
| ENSMUSG00000071203.6 | 76.10130369 | 1.334160063 | 0.384211134 | 3.4724659 | 0.000515701 | 0.023633114 | Naip5 |
| ENSMUSG00000064437.1 | 46.6545078 | -1.269799402 | 0.366176898 | -3.467721228 | 0.000524891 | 0.0239662 | Snord49b |
| ENSMUSG00000042284.9 | 170.2024137 | -1.242530071 | 0.358561169 | -3.465322458 | 0.000529596 | 0.024092756 | Itga1 |
| ENSMUSG00000051354.9 | 65.55272733 | 1.315691151 | 0.379876818 | 3.463467864 | 0.00053326 | 0.024171235 | Samd3 |
| ENSMUSG00000017861.7 | 361.1085266 | 0.94155858 | 0.272721636 | 3.452452811 | 0.000555515 | 0.025088731 | Mybl2 |
| ENSMUSG00000015981.8 | 516.1097752 | 0.854336037 | 0.24773644 | 3.448568307 | 0.000563567 | 0.025088791 | Stk32c |
| ENSMUSG00000022416.11 | 91.49600617 | -0.948823578 | 0.275094391 | -3.449083689 | 0.000562492 | 0.025088791 | Cacna1i |
| ENSMUSG00000029530.11 | 40.56838065 | -1.224632338 | 0.35500005 | -3.449668073 | 0.000561276 | 0.025088791 | Ccr9 |
| ENSMUSG00000043230.2 | 267.2444284 | -0.675964574 | 0.195877806 | -3.450950304 | 0.000558616 | 0.025088791 | Fam124b |
| ENSMUSG00000032528.4 | 47.66718935 | -1.202540213 | 0.34902122 | -3.445464472 | 0.000570079 | 0.0251987 | Vipr1 |
| ENSMUSG00000044340.7 | 164.6899048 | 0.925018535 | 0.268402269 | 3.446388661 | 0.000568133 | 0.0251987 | Phlpp1 |
| ENSMUSG00000037346.4 | 10.54875417 | -1.364814273 | 0.396290091 | -3.443977791 | 0.000573223 | 0.025248134 | Hrh4 |
| ENSMUSG00000025993.6 | 73.06141709 | 1.247676285 | 0.362511816 | 3.441753426 | 0.000577957 | 0.025278006 | Slc40a1 |
| ENSMUSG00000053716.9 | 276.4403495 | -0.792237508 | 0.230135272 | -3.442486237 | 0.000576393 | 0.025278006 | Dusp7 |
| ENSMUSG00000016087.9 | 1178.844799 | -0.801732374 | 0.233346213 | -3.43580624 | 0.000590793 | 0.025639075 | Fli1 |
| ENSMUSG00000035547.10 | 16.03478819 | 1.360636327 | 0.396100545 | 3.435078153 | 0.000592383 | 0.025639075 | Capn5 |
| ENSMUSG00000079553.6 | 328.4755263 | 0.86880486 | 0.252814003 | 3.436537727 | 0.0005892 | 0.025639075 | Kifc1 |
| ENSMUSG00000104418.1 | 8.993759972 | -1.378100619 | 0.402540103 | -3.423511372 | 0.000618177 | 0.026662879 | Gm37070 |
| ENSMUSG00000036503.9 | 377.2992376 | -0.909210532 | 0.265660079 | -3.422458258 | 0.000620576 | 0.026674079 | Rnf13 |
| ENSMUSG00000009731.4 | 13.85507869 | 1.366406999 | 0.399806694 | 3.417669135 | 0.000631598 | 0.027053524 | Kcnd1 |
| ENSMUSG00000021701.7 | 88.79744807 | 1.059157082 | 0.31022446 | 3.414163674 | 0.000639781 | 0.027053524 | Plk2 |
| ENSMUSG00000039512.11 | 126.9467295 | 1.131023405 | 0.331293531 | 3.413961641 | 0.000640256 | 0.027053524 | Uhrf1bp1 |
| ENSMUSG00000049103.9 | 3303.256366 | -1.287605767 | 0.37704398 | -3.415001529 | 0.000637817 | 0.027053524 | Ccr2 |
| ENSMUSG00000103057.1 | 41.55991233 | 1.367551295 | 0.40040577 | 3.415413561 | 0.000636852 | 0.027053524 | ENSMUSG00000103057 |
| ENSMUSG00000064043.9 | 2232.277183 | 1.051282373 | 0.308040616 | 3.412804409 | 0.000642981 | 0.027076874 | Trerf1 |
| ENSMUSG00000024867.10 | 19.62971605 | 1.363528928 | 0.400551469 | 3.40412914 | 0.000663754 | 0.027626532 | Pip5k1b |
| ENSMUSG00000032434.8 | 1072.915299 | -0.588733112 | 0.172951826 | -3.404029471 | 0.000663996 | 0.027626532 | Cmtm6 |
| ENSMUSG00000037815.6 | 557.5393287 | -0.537824051 | 0.158055741 | -3.402749228 | 0.000667115 | 0.027626532 | Ctnna1 |
| ENSMUSG00000075015.3 | 29.85649937 | 1.372251706 | 0.403109339 | 3.404167484 | 0.00066366 | 0.027626532 | Gm10801 |
| ENSMUSG00000079563.5 | 464.9072879 | -0.601481381 | 0.176741411 | -3.403171769 | 0.000666084 | 0.027626532 | Pglyrp2 |
| ENSMUSG00000023903.7 | 102.4152681 | -1.217939598 | 0.35842792 | -3.398004261 | 0.000678794 | 0.028017095 | Mmp25 |
| ENSMUSG00000078995.5 | 102.096881 | -0.803098508 | 0.23702617 | -3.388227173 | 0.00070346 | 0.028939359 | Zfp456 |
| ENSMUSG00000003031.10 | 959.4381572 | -0.602442067 | 0.17795554 | -3.385351567 | 0.000710871 | 0.029131768 | Cdkn1b |
| ENSMUSG00000079298.5 | 33.9948459 | -1.334784037 | 0.394369379 | -3.384603645 | 0.000712811 | 0.029131768 | Klrb1b |
| ENSMUSG00000024206.10 | 57.80926122 | 0.984541257 | 0.291738699 | 3.374736573 | 0.000738864 | 0.0300824 | Rfx2 |
| ENSMUSG00000037826.3 | 299.3987496 | -0.766581454 | 0.227253981 | -3.373236627 | 0.000742901 | 0.0300824 | Ppm1k |
| ENSMUSG00000045414.7 | 240.6437051 | -0.649297495 | 0.192493699 | -3.373084412 | 0.000743312 | 0.0300824 | 1190002N15Rik |
| ENSMUSG00000002985.11 | 63.80055236 | 1.290327967 | 0.383446442 | 3.365080033 | 0.000765215 | 0.030414207 | Apoe |
| ENSMUSG00000022218.11 | 10.91849611 | 1.356973931 | 0.402792606 | 3.368914698 | 0.000754648 | 0.030414207 | Tgm1 |
| ENSMUSG00000029287.10 | 372.2361173 | 0.900176134 | 0.267443471 | 3.365855706 | 0.000763066 | 0.030414207 | Tgfbr3 |
| ENSMUSG00000030339.7 | 20.76328879 | 1.352404271 | 0.401933909 | 3.364742912 | 0.00076615 | 0.030414207 | Ltbr |

| | | | | | | | |
|---|---|---|---|---|---|---|---|
| ENSMUSG00000044026.2 | 274.5220195 | -0.719803875 | 0.213773146 | -3.3671389 | 0.000759524 | 0.030414207 | Slc35g1 |
| ENSMUSG00000047821.12 | 160.6853223 | -0.877073014 | 0.260650594 | -3.364937715 | 0.000765609 | 0.030414207 | Trim16 |
| ENSMUSG00000034855.9 | 79.06431275 | 1.153447483 | 0.344913774 | 3.344161844 | 0.000825316 | 0.032658923 | Cxcl10 |
| ENSMUSG00000052013.10 | 326.9174169 | -0.747190216 | 0.223573006 | -3.342041279 | 0.000831647 | 0.032805313 | Btla |
| ENSMUSG00000025044.11 | 12.13550062 | 1.335845469 | 0.400267796 | 3.337379333 | 0.000845724 | 0.033255376 | Msr1 |
| ENSMUSG00000015133.12 | 314.6004984 | 0.782260903 | 0.23452892 | 3.335456046 | 0.000851596 | 0.033380962 | Lrrk1 |
| ENSMUSG00000037902.14 | 172.2004596 | 1.332077099 | 0.400870317 | 3.322962668 | 0.000890668 | 0.034493167 | Sirpa |
| ENSMUSG00000045165.5 | 889.4507591 | -0.78848469 | 0.237353724 | -3.321981534 | 0.000893806 | 0.034493167 | AI467606 |
| ENSMUSG00000059033.7 | 58.24033036 | -1.00731353 | 0.30320861 | -3.322179834 | 0.000893171 | 0.034493167 | Rpl18a-ps1 |
| ENSMUSG00000063810.5 | 395.9664463 | 0.711789015 | 0.214065791 | 3.325094643 | 0.000883885 | 0.034493167 | Alms1 |
| ENSMUSG00000091694.4 | 77.08580135 | 1.337822114 | 0.402613087 | 3.322848057 | 0.000891034 | 0.034493167 | Apol11b |
| ENSMUSG00000058818.9 | 111.8065419 | 1.337623809 | 0.402851383 | 3.320390264 | 0.000898917 | 0.034583331 | Pirb |
| ENSMUSG00000026009.10 | 949.9558529 | -0.881768776 | 0.266070885 | -3.314037066 | 0.000919593 | 0.035269935 | Icos |
| ENSMUSG00000037110.15 | 1179.32492 | 0.609746109 | 0.184394515 | 3.306747531 | 0.000943859 | 0.036089585 | Ralgapa2 |
| ENSMUSG00000019087.9 | 508.1372404 | -0.662950132 | 0.200648255 | -3.304041348 | 0.000953018 | 0.036328337 | Atp6ap1 |
| ENSMUSG00000085603.2 | 161.93303 | -0.888046281 | 0.269016598 | -3.301083608 | 0.000963122 | 0.036601564 | Gm11346 |
| ENSMUSG00000037563.10 | 1265.966503 | -0.790383937 | 0.239924485 | -3.294302938 | 0.000986661 | 0.037382156 | Rps16 |
| ENSMUSG00000022102.9 | 946.016193 | -0.461440566 | 0.140445077 | -3.285558861 | 0.001017803 | 0.038329037 | Dok2 |
| ENSMUSG00000059040.4 | 55.89234864 | -0.946530102 | 0.288070538 | -3.285758093 | 0.001017083 | 0.038329037 | Eno1b |
| ENSMUSG00000022602.10 | 109.8733459 | 0.840626193 | 0.25606876 | 3.282814324 | 0.001027763 | 0.038587561 | Arc |
| ENSMUSG00000027456.8 | 113.2290649 | -0.789239888 | 0.240486556 | -3.281846186 | 0.001031298 | 0.038604009 | Sdcbp2 |
| ENSMUSG00000026399.8 | 50.81969118 | -1.114636293 | 0.339843936 | -3.279847527 | 0.001038632 | 0.03876212 | Cd55 |
| ENSMUSG00000057729.8 | 70.860543 | 1.321261232 | 0.403062161 | 3.278058228 | 0.001045238 | 0.038892226 | Prtn3 |
| ENSMUSG00000030142.6 | 23.962315 | 1.317465063 | 0.402269158 | 3.275083451 | 0.001056308 | 0.038907942 | Clec4e |
| ENSMUSG00000031698.10 | 11.62034791 | 1.31874932 | 0.402452766 | 3.276780362 | 0.00104998 | 0.038907942 | Mylk3 |
| ENSMUSG00000031785.11 | 25.22433052 | 1.29622036 | 0.395841772 | 3.274592153 | 0.001058146 | 0.038907942 | Adgrg1 |
| ENSMUSG00000061684.5 | 33.15310224 | -1.176972932 | 0.35928995 | -3.275830371 | 0.001053518 | 0.038907942 | Rpl21-ps8 |
| ENSMUSG00000047181.8 | 37.45976436 | 1.308722585 | 0.400155258 | 3.27053702 | 0.001073435 | 0.039354018 | Samd14 |
| ENSMUSG00000032265.10 | 298.6627577 | -0.803744223 | 0.246158229 | -3.265152767 | 0.001094051 | 0.039992199 | Fam46a |
| ENSMUSG00000026447.12 | 156.3604431 | 0.97682431 | 0.29926559 | 3.26407159 | 0.001098234 | 0.040027744 | Pik3c2b |
| ENSMUSG00000034135.10 | 885.8168575 | 0.777602673 | 0.238390784 | 3.261882276 | 0.001106751 | 0.040220556 | Sik3 |
| ENSMUSG00000042417.4 | 11.10073989 | 1.293429259 | 0.396645657 | 3.260918748 | 0.001110519 | 0.040240157 | Ccno |
| ENSMUSG00000040964.12 | 15.63488246 | 1.313046087 | 0.40297104 | 3.258413028 | 0.001120372 | 0.040479532 | Arhgef10l |
| ENSMUSG00000021614.12 | 104.1071354 | 1.283999473 | 0.394184117 | 3.257359745 | 0.001124538 | 0.040512625 | Vcan |
| ENSMUSG00000014498.5 | 1164.645722 | 0.818779393 | 0.251556492 | 3.254852967 | 0.001134511 | 0.040754115 | Ankrd52 |
| ENSMUSG00000073490.6 | 23.66010116 | 1.200767756 | 0.369147229 | 3.2528153 | 0.001142677 | 0.040929524 | AI607873 |
| ENSMUSG00000020893.13 | 464.1762747 | 0.976878775 | 0.300690039 | 3.248789949 | 0.00115897 | 0.041394167 | Per1 |
| ENSMUSG00000020185.12 | 363.1816188 | 0.863059689 | 0.265797593 | 3.247056072 | 0.001166054 | 0.041528185 | E2f7 |
| ENSMUSG00000020125.6 | 28.82736115 | 1.282126332 | 0.395309697 | 3.243346529 | 0.001181344 | 0.04195287 | Elane |
| ENSMUSG00000009687.10 | 2051.033536 | -0.810543451 | 0.250204615 | -3.239522387 | 0.001197301 | 0.042040431 | Fxyd5 |
| ENSMUSG00000028228.5 | 410.109778 | -0.825721777 | 0.254878021 | -3.23967431 | 0.001196663 | 0.042040431 | Cpne3 |
| ENSMUSG00000039477.12 | 767.6208994 | 1.019833739 | 0.31459234 | 3.241762783 | 0.001187929 | 0.042040431 | Tnrc18 |
| ENSMUSG00000061589.10 | 678.9513901 | 1.028944698 | 0.317589944 | 3.239852893 | 0.001195914 | 0.042040431 | Dot1l |
| ENSMUSG00000019467.9 | 12.32609367 | 1.27782229 | 0.394592323 | 3.238335406 | 0.001202294 | 0.042097166 | Arhgef25 |
| ENSMUSG00000018476.7 | 523.5455237 | 1.106757899 | 0.341897907 | 3.237100534 | 0.001207509 | 0.042161331 | Kdm6b |
| ENSMUSG00000021457.10 | 137.0152323 | 1.302345407 | 0.402439687 | 3.236125681 | 0.00121164 | 0.042187418 | Syk |
| ENSMUSG00000028618.7 | 497.1706783 | -0.891261504 | 0.275519817 | -3.234836295 | 0.001217125 | 0.042260341 | Tmem59 |
| ENSMUSG00000029797.8 | 68.46705564 | 1.279183934 | 0.396574781 | 3.225580636 | 0.001257174 | 0.043125547 | Sspo |
| ENSMUSG00000032440.8 | 2472.486011 | -0.65474183 | 0.20301522 | -3.225087415 | 0.001259342 | 0.043125547 | Tgfbr2 |
| ENSMUSG00000038679.12 | 599.8986248 | 1.013349285 | 0.314129058 | 3.225901138 | 0.001255767 | 0.043125547 | Trps1 |
| ENSMUSG00000040451.13 | 931.2300693 | -0.556051471 | 0.172373634 | -3.225849913 | 0.001255992 | 0.043125547 | Sgms1 |
| ENSMUSG00000071252.5 | 28.69643118 | 1.129707267 | 0.350255214 | 3.225383158 | 0.001258042 | 0.043125547 | 2210408I21Rik |
| ENSMUSG00000004032.6 | 19.98768231 | -1.201003953 | 0.372719345 | -3.222274259 | 0.001271773 | 0.043372951 | Gstm5 |
| ENSMUSG00000023995.7 | 17.47808262 | 1.288280022 | 0.399853544 | 3.221879717 | 0.001273526 | 0.043372951 | Tspo2 |
| ENSMUSG00000023886.9 | 15.3479503 | -1.296821875 | 0.402779395 | -3.219682762 | 0.001283325 | 0.043587601 | Smoc2 |
| ENSMUSG00000049999.4 | 14.66351581 | 1.290499955 | 0.40116773 | 3.216858831 | 0.001296024 | 0.043899275 | Ppp1r3d |
| ENSMUSG00000056656.5 | 54.88576794 | 1.272617027 | 0.396274002 | 3.211457271 | 0.001320636 | 0.044611731 | Apol8 |
| ENSMUSG00000081769.4 | 195.7853627 | 0.90475798 | 0.282068858 | 3.207578418 | 0.001338576 | 0.045095527 | Gm12216 |
| ENSMUSG00000031137.13 | 66.0770876 | 1.058062198 | 0.329968123 | 3.206558828 | 0.001343328 | 0.04512662 | Fgf13 |
| ENSMUSG00000033174.13 | 73.29248509 | -0.957887054 | 0.298795412 | -3.205829185 | 0.001346739 | 0.04512662 | Mgll |
| ENSMUSG00000090290.1 | 332.5207342 | 0.851127534 | 0.265691424 | 3.203443758 | 0.001357946 | 0.045258802 | Gm17296 |
| ENSMUSG00000090942.1 | 99.29820964 | 1.277468622 | 0.398690838 | 3.204158466 | 0.001354579 | 0.045258802 | F830016B08Rik |
| ENSMUSG00000099954.1 | 25.01688652 | 1.244341325 | 0.388539212 | 3.202614525 | 0.001361861 | 0.045268272 | Gm28112 |

| | | | | | | | |
|---|---|---|---|---|---|---|---|
| ENSMUSG00000043017.9 | 26.3416053 | -1.138648424 | 0.355854256 | -3.199760588 | 0.001375418 | 0.045597297 | Ptgir |
| ENSMUSG00000003348.9 | 1455.359713 | -0.58271596 | 0.18257817 | -3.191597108 | 0.001414885 | 0.046657531 | Mob3a |
| ENSMUSG00000099250.1 | 1408.213345 | -1.182337839 | 0.370377526 | -3.192250488 | 0.001411689 | 0.046657531 | Rn7s2 |
| ENSMUSG00000023571.4 | 71.13832268 | 1.095635467 | 0.343692391 | 3.187837429 | 0.001433411 | 0.047143716 | Fam132a |
| ENSMUSG00000064023.3 | 206.1368031 | -0.720078271 | 0.225995876 | -3.186245179 | 0.001441324 | 0.047279218 | Klk8 |
| ENSMUSG00000039959.9 | 2187.671399 | 0.646986755 | 0.203177295 | 3.184345751 | 0.001450816 | 0.047465676 | Hip1 |
| ENSMUSG00000032028.11 | 22.7443674 | 1.273536189 | 0.400299521 | 3.181458185 | 0.001465357 | 0.047815895 | Nxpe2 |
| ENSMUSG00000096751.3 | 233.8627531 | 1.210040124 | 0.380732017 | 3.178193764 | 0.001481957 | 0.048231308 | Gm28373 |
| ENSMUSG00000057329.7 | 536.3831371 | -0.878531374 | 0.276568462 | -3.176542146 | 0.001490421 | 0.048380473 | Bcl2 |
| ENSMUSG00000061848.5 | 8.589356577 | -1.275554383 | 0.401758462 | -3.174928475 | 0.001498734 | 0.048523959 | Gm5805 |
| ENSMUSG00000020718.8 | 253.8770192 | 0.679320529 | 0.214344754 | 3.169289267 | 0.001528122 | 0.049347264 | Polg2 |
| ENSMUSG00000030589.11 | 85.44841504 | 1.26862675 | 0.400563303 | 3.167106773 | 0.001539638 | 0.049590658 | Rasgrp4 |
