## Supplementary Table 5 for "EOMES and IL-10 regulate anti-tumor activity of PD-1^+^ CD4^+^ T-cells in B-cell Non-Hodgkin lymphoma"

| Genes filtered from GO cytokine activity (GO:0005125) |
| --- |
| Areg |
| Csf1 |
| Csf2 |
| Csf3 |
| Ctf1 |
| Ctf2 |
| Ebi3 |
| Flt3l |
| Ifna1 |
| Ifna11 |
| Ifna12 |
| Ifna13 |
| Ifna14 |
| Ifna15 |
| Ifna16 |
| Ifna2 |
| Ifna4 |
| Ifna5 |
| Ifna6 |
| Ifna7 |
| Ifna9 |
| Ifnab |
| Ifnb1 |
| Ifne |
| Ifng |
| Ifnk |
| Ifnl2 |
| Ifnl3 |
| Ifnz |
| Il10 |
| Il11 |
| Il12a |
| Il12b |
| Il13 |
| Il15 |
| Il16 |
| Il17a |
| Il17b |
| Il17c |
| Il17f |
| Il18 |
| Il19 |

Il1a
Il1b
Il1f10
Il1f5
Il1f6
Il1f8
Il1f9
Il1rn
Il2
Il20
Il21
Il22
Il23a
Il24
Il25
Il27
Il3
Il31
Il33
Il34
Il4
Il5
Il6
Il7
Il9
Lif
Lta
Ltb
Osm
Tgfb1
Tgfb2
Tgfb3
Tnf
Tnfsf10
Tnfsf11
Tnfsf12
Tnfsf13
Tnfsf13b
Tnfsf14
Tnfsf15
Tnfsf18
Tnfsf4
Tnfsf8

Tnfsf9
Tslp

| Genes filtered from GO cytokine receptor activity (GO:0004896) |
|---|
| Csf1r |
| Csf2ra |
| Csf2rb |
| Csf2rb2 |
| Csf3r |
| Ifnar1 |
| Ifnar2 |
| Ifngr1 |
| Ifngr2 |
| Ifnlr1 |
| Il10ra |
| Il10rb |
| Il11ra1 |
| Il11ra2 |
| Il12b |
| Il12rb1 |
| Il12rb2 |
| Il13ra1 |
| Il13ra2 |
| Il15ra |
| Il17ra |
| Il17rb |
| Il17rc |
| Il17rd |
| Il17re |
| Il18r1 |
| Il18rap |
| Il1r1 |
| Il1r2 |
| Il1rap |
| Il1rapl2 |
| Il1rl1 |
| Il1rl2 |
| Il1rn |
| Il20ra |
| Il20rb |
| Il21r |
| Il22ra1 |
| Il22ra2 |
| Il23r |
| Il27ra |
| Il2ra |

Il2rb
Il2rg
Il31ra
Il3ra
Il4ra
Il5ra
Il6ra
Il6st
Il7r
Il9r
Lifr
Osmr
Prlr

| Genes filtered of Crawford et al. and GO cell surface (GO:0009986) |
| --- |
| Alcam |
| Cacna1d |
| Ccr2 |
| Ccrl2 |
| Cd44 |
| Cd80 |
| Cxcl10 |
| Cxcr5 |
| Cxcr6 |
| Entpd1 |
| H2-Q6 |
| Icos |
| Ifng |
| Il12rb2 |
| Itga4 |
| Itgb8 |
| Klrc1 |
| Lag3 |
| Lgals1 |
| Ly6a |
| Nt5e |
| Pdcd1 |
| Tigit |
| Timp2 |
| Tjp2 |
